# Chaperone-Mediated Reflux of Secretory Proteins to the Cytosol During Endoplasmic Reticulum Stress

**DOI:** 10.1101/562306

**Authors:** Aeid Igbaria, Philip I. Merksamer, Ala Trusina, Firehiwot Tilahun, Jefferey R. Johnson, Onn Brandman, Nevan J. Krogan, Jonathan S. Weissman, Feroz R. Papa

**Author notes:** Equal Contribution (Aeid Igbaria, Philip I. Merksamer).

## Abstract

Diverse perturbations to endoplasmic reticulum (ER) functions compromise the proper folding and structural maturation of secretory proteins. To study secretory pathway physiology during such “ER stress”, we employed an ER-targeted, redox-responsive, green fluorescent protein—eroGFP—that reports on ambient changes in oxidizing potential. Here we find that diverse ER stress agents cause properly folded, ER-resident eroGFP (and other ER luminal proteins) to “reflux” back to the reducing environment of the cytosol as intact, folded proteins. By utilizing eroGFP in a comprehensive genetic screen in *S. cerevisiae*, we show that ER protein reflux during ER stress requires specific chaperones and co-chaperones residing in both the ER and the cytosol. Chaperone-mediated ER protein reflux does not require E3 ligase activity, and proceeds even more vigorously when these ER-associated degradation (ERAD) factors are crippled, suggesting that reflux may work in parallel with ERAD. In summary, chaperone-mediated ER-protein reflux may be a conserved protein quality control process that evolved to maintain secretory pathway homeostasis during ER protein-folding stress.

**SIGNIFICANCE:** Approximately one third of eukaryotic proteins are synthesized on ribosomes attached to the endoplasmic reticulum (ER) membrane. Many of these polypeptides co- or post-translationally translocate into the ER, wherein they fold and mature. An ER quality-control system proofreads these proteins by facilitating their folding and modification, while eliminating misfolded proteins through ER-associated degradation (ERAD). Yet, the fate of many secretory proteins during ER stress is not completely understood. Here, we uncovered an ER-stress induced “protein reflux” system that delivers intact, folded ER luminal proteins back to the cytosol without degrading them. We found that ER protein reflux works in parallel to ERAD and requires distinct ER-resident and cytosolic chaperones and co-chaperones.

## INTRODUCTION

In eukaryotic cells, secretory and membrane proteins begin translation in the cytoplasm and are then either co-or post-translationally translocated through the Sec61 translocon channel into the endoplasmic reticulum (ER) [1]. The ER is crowded with molecular chaperones and protein modifying enzymes that promote folding and structural maturation of these nascent, maturing secretory pathway client proteins as they traverse the early secretory pathway [2]. To ensure stringent quality control over these secretory cargo, those proteins that fail to correctly fold and mature are retrieved from the ER, ubiquitylated, and degraded by the 26S proteasome in the cytosol in a process termed ER-associated degradation (ERAD) [3].

Diverse environmental perturbations or genetic mutations can elevate misfolding of maturing proteins in the ER. During such “ER stress,” cells trigger an intracellular signaling pathway called the unfolded protein response (UPR) that augments protein folding reactions through transcriptional upregulation of genes encoding ER chaperones, oxidoreductases, lipid biosynthetic enzymes, and ERAD components [4]. If these adaptive UPR outputs prove successful in reducing the concentration of unfolded proteins in the ER, cells become restored to a homeostatic state [5].

However, because the UPR’s activating inputs— (i.e. unfolded proteins)—are unfeasible to monitor *in vivo*, it is often unclear if and when the UPR has successfully restored homeostasis. To address this problem orthogonally, we previously developed an ER-targeted redox-sensitive green fluorescent protein (GFP)—called eroGFP—to follow oxidative protein folding in the ER, reasoning that this essential ER physiological function may deviate during ER stress and thereby provide an independent measure of ER health (i.e., that is distinct from solely measuring UPR activation). We previously showed that differential, real-time, quantitative eroGFP changes occurred dynamically upon general loss of ER protein folding homeostasis in wild-type cells, and in a small, select group of yeast mutants [6]. Here, using high-throughput flow cytometry, we have extended this analysis to the entire yeast genome to query nearly all non-essential and essential genes. Through this screen, we have identified and characterized a process by which eroGFP, and a number of ER-resident luminal proteins, are “refluxed” back to the cytosol as intact folded proteins during ER stress. The protein reflux process occurs independently of Hrd1 and Doa10 E3-ligases and does not require poly-ubiquitinylation. Instead, ER protein reflux requires specific chaperones and co-chaperones both in the ER and cytosol, and is reminiscent of a molecular ratchet that promotes translocation, but proceeding vectorially in the opposite direction [7, 8].

## RESULTS

### ER to Cytosol Reflux of ER-targeted eroGFP

Designed to be a reporter of ambient redox potential, eroGFP has an engineered reversible disulfide bond that alters fluorescence excitability from its two maxima of 490 and 400nm, such that reduction of the disulfide increases fluorescence from 490nm excitation, at the expense of that from 400nm [6, 9]. Thus, eroGFP is ratiometric by excitation, which facilitates internally controlled measurement of its oxidation state. Through flow cytometry, the eroGFP ratio—defined as fluorescence from excitation at 488 versus 405 nm in *log*_2_ space—can be measured in single yeast cells growing in populations (Figure 1A). Targeted to the oxidizing environment of the ER though an N-terminal Kar2 signal peptide sequence (and retained in the organelle through a C-terminal HDEL sequence), eroGFP (which has a redox midpoint potential of -282 mV) is nearly completely oxidized in the oxidizing environment of the ER at baseline. Treatment with hydrogen peroxide (H2O2) only slightly further decreases the eroGFP ratio [6]. However, a wide dynamic range exists for reduction since titration with increasing amounts of the reductant dithiothreitol (DTT)—an ER stress agent—dose-dependently increases the eroGFP ratio until the reporter becomes fully reduced (Figure 1B). As previously shown, acute treatment of cells with (saturating) DTT causes rapid elevation of the eroGFP ratio to its new steady-state level due to *in situ* and complete reduction of the reporter [6] (Figure S1A-C). Tunicamycin (Tm), which impairs N-linked protein glycosylation in the ER, also led to partial reduction of the eroGFP reporter, but with slower dynamics compared to treatment with DTT (as previously shown [6]) (Figure S1A-C). Indeed, the general utility of the eroGFP tool is that it deflects differentially (by reduction) in response to diverse ER stress agents (including forced expression of unfolded secretory proteins) [6].

**Figure 1.**
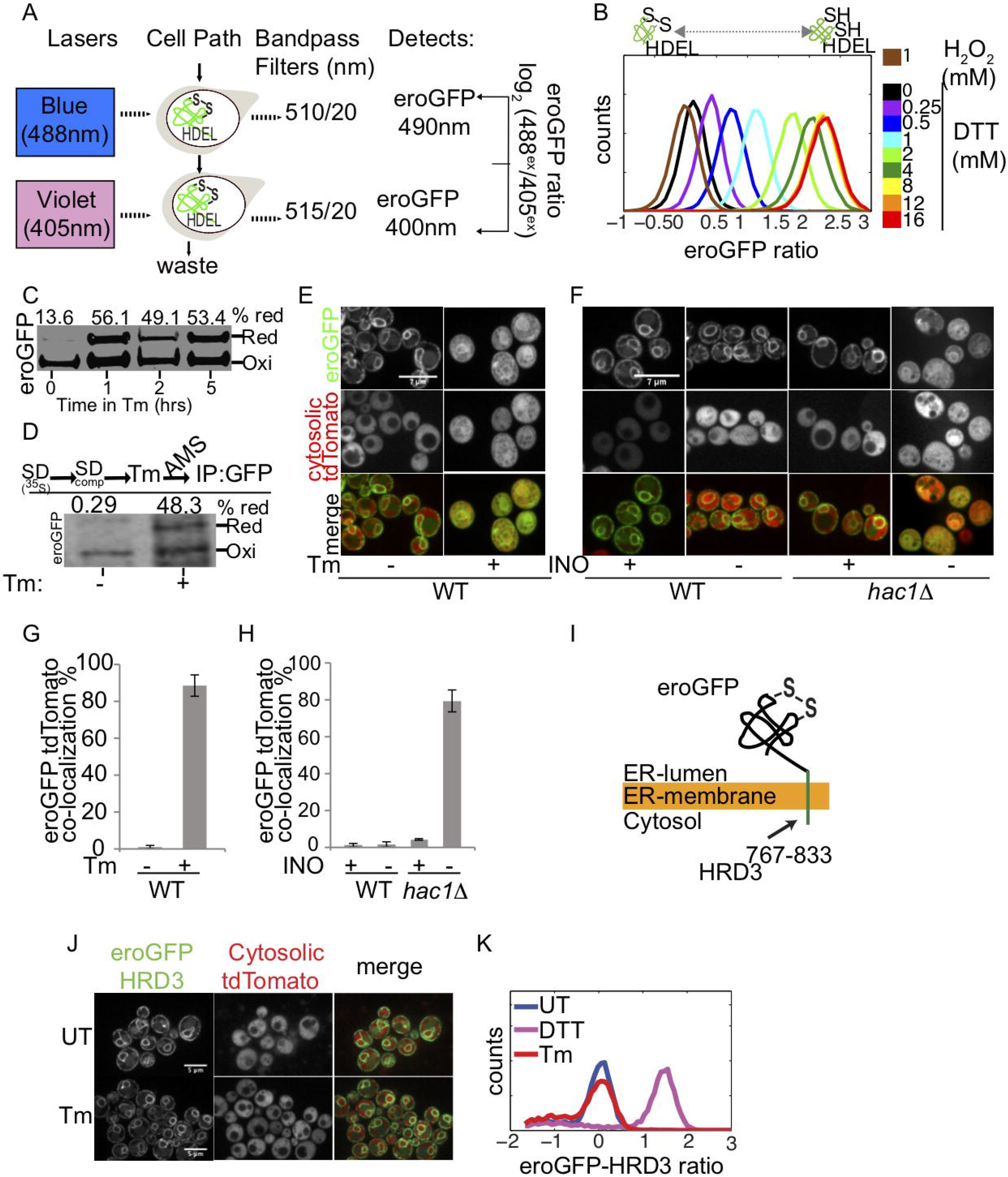
ER-targeted eroGFP localizes to the cytosol during ER stress. (A) Schematic showing configuration of flow cytometer laser lines and filters used to measure eroGFP fluorescence excitation and emission. (B) eroGFP ratios for populations of wild-type cells treated with the indicated concentration of DTT or H2O2 for 20 min. (C) eroGFP redox state in WT cells treated with (6μg/mL) Tm for the indicated time points. (D) eroGFP redox state after ^35^S pulse chase in WT cells treated with Tm for 2hrs. Extracts were treated with AMS, immunoprecipitated with Anti-GFP and resolved on nonreducing SDS-PAGE. (E) Confocal images of wild-type cells expressing eroGFP and cytosolic tdTomato treated with Tm for 2 h. (F) Confocal images of wild-type and *hac1Δ* yeast expressing eroGFP and cytosolic tdTomato starved for inositol for 8 hours. Quantification of Tm confocal images (G) and inositol confocal images (H), Error bars represent S.E.M of two independent experiments. (I) Schematic of HRD3-eroGFP. eroGFP was translationally fused to residues 767-833 of HRD3 to imbed eroGFP in the ER membrane. (J) Confocal images of WT cells expressing HRD3-eroGFP and cytosolic tdTomato treated with Tm for 2 hours. (K) Histograms of HRD3-eroGFP ratios for wild-type cells treated with 2mM DTT for 20 minutes and Tm for 5 hours.

Since our original study, which was based on single-cell interrogations using real-time flow cytometry, it was also reported that partial cytoplasmic localization of eroGFP occurs during expression of a mutant of the UPR master regulator, Ire1 that cannot deactivate the UPR [10]; this result implied that cytoplasmic localization of the eroGFP reporter may occur due to a translocation defect under unresolved ER stress signaling. But in our original study, through using a yeast strain expressing a GAL1/10 promoter-driven eroGFP construct gene that no longer expresses new eroGFP after glucose shutoff, we had established that eroGFP reduction due to Tm provision occurred *after* the glucose shutoff -(Figure S7 in [6]). Thus, we had reasonably concluded that ER stress induced by Tm caused reduction of *pre-existing* eroGFP that was already residing in the ER lumen.

We revisited these experimental systems by showing again that provision of Tm to wild-type yeast cells dynamically caused eroGFP reporter reduction (to ~50% of basal levels) (detectable in a time course of AMS modification, resolution on non-reducing SDS-PAGE, then followed by immunoblotting against GFP) (Figure 1C). Also, in a pulse-chase regime to label *pre-existing* reporter, reduction was detectable within two hours after Tm provision (i.e., when eroGFP reduction reaches its new steady state) (Figure 1D). Next, to specifically track the cytoplasmic compartment with high sensitivity, we integrated a cytosolically-disposed tdTomato reporter into eroGFP-expressing yeast and followed both reporters by fluorescence microscopy. Using this double reporter system and a high dose regime of Tm (6 μg/ml) provided for 2 hours, we visually scored any cell showing merged eroGFP/cytosolic tdTomato as being “colocalized” (yellow overlay). This regime revealed significant numbers of cells with some eroGFP signal localized to the cytosol (Figure 1E, G; Figure S1D). However, we still observed significant eroGFP signal retained within the ER in these co-localized cells, which is consistent with the incomplete reduction of eroGFP that we had observed after Tm treatment by flow cytometry.

Because Tm is a severe non-physiological stress, we next asked whether eroGFP localizes to the cytosol under a more physiologic stress. To this end, we starved UPR-deficient yeast mutants (which are inositol auxotrophs) for inositol, which we previously found to cause reduction of eroGFP in sub-populations when measured by flow cytometry [6, 10]. Relative to the Tm regime, we observed a smaller fraction of cells with eroGFP localized to the cytosol in inositol-starved *hac1*Δ mutants, but not in control wild-type cells, which are prototrophs for inositol and retain eroGFP in the ER upon inositol starvation (Figures 1F, H)).

To further test whether Tm-induced reduction of eroGFP occurred due to its exposure to the reducing cytosol, we constructed a variant of eroGFP that was fused to the transmembrane domain of the single-pass ER membrane protein Hrd3 such that the GFP domain remained topologically disposed inside the ER (Figure 1I). This Hrd3-eroGFP reporter’s fluorescence signal did not overlap with cytosolic tdTomato following Tm (Figure 1J), and its oxidation state remained unperturbed under this treatment, while it could still be reduced *in situ* with DTT (Figure 1K). Thus, altered (cytosolic) localization during ER stress is confined to a soluble form of eroGFP, and the oxidation change of this reporter occurs upon exposure to the reducing environment of the cytosol.

Two mechanisms could account *a priori* for localization of ER-targeted eroGFP in the cytosol during ER stress: 1) eroGFP that was *en route* to the ER may have become averted due to disabled translocation into the organelle during ER stress, as was shown for other client proteins in a “pre-emptive” quality control pathway [11]; or 2) eroGFP already in the ER lumen may have been returned back to the cytosol during stress. To distinguish between these two possibilities, we engineered an N-linked glycosylation site into eroGFP in a nine amino acid linker region between the GFP coding sequence and the C-terminal HDEL retrieval sequence to follow the fate of the reporter after its glycosylation in the ER. We termed this variant eroGFP-Glyc (Figure S2A). As expected, in unstressed cells eroGFP-Glyc migrates slower on SDS-PAGE compared to eroGFP, consistent with its glycosylation. Confirming this, treatment with the deglycosylase, EndoH, increased eroGFP-Glyc mobility such that it now co-migrated with eroGFP (Figure S2B).

To then directly follow this reporter’s fate after it was already resident in the ER lumen, we placed eroGFP-Glyc under control of the GAL1/10 promoter to rapidly cease *de novo* production with glucose. Quantitative PCR and pulse-label analysis confirmed that glucose provision halted transcription within the first 30 minutes and new protein synthesis by 2 hours (Figures S2C). We estimated the half-life of eroGFP-Glyc based on a ^35^S methionine pulse-label experiment (Figure S2D, S2E) and used this information to determine that the fraction of newly synthesized eroGFP is ~ 6% of total eroGFP at 2 hours from the shift to glucose (Figure S2F); thus, the majority of the reporter should be pre-existing. As with eroGFP, we again confirmed that this pre-existing eroGFP-Glyc (i.e., post glucose addition) became localized to the cytosol during the Tm treatment of 2 hours (Figure S2G). Moreover, we reasoned that if this pool of pre-existing eroGFP-Glyc had already entered and then later exited the ER, the beta-aspartylglycosylamine bond at its N-glycan tree should become cleaved in the cytosol by deglycosylating enzyme peptide N-glycanase— PNGase—thus converting the asparagine at the glycosylation site to an aspartate residue [12]. To test this, we used two-dimensional gel electrophoresis to monitor isoelectric shifts that would indicate an asparagine to aspartate conversion. After treatment with Tm, we observed that the faster migrating deglycosylated species have a lower isolectric point (i.e., more acidic, as expected) than the slower migrating glycosylated species, suggesting an asparagine to aspartate conversion (Figure S2H). Confirming this, the shift with Tm is superimposable when eroGFP-Glyc is treated enzymatically with PNGase, which cleaves the beta aspartylglycosylamine bond, and also with an eroGFP-Glyc variant in which an aspartate has replaced the asparagine (eroGFP^ND^) at the glycosylation site. In addition, we used mass spectrometry to measure deamidation of the deglycosylated asparagine. While a cytosolic version of the reporter lacking the signal peptide, but bearing the C-terminal glycosylation signal, termed cytoGFP-Glyc, displayed low levels of deamidation under Tm, eroGFP-Glyc showed a 200-fold relative increase in deamidation of the Asn residue in the relevant peptide (Figure S2I). For comparison, eroGFP-Glyc treated with PNGase, which should yield the theoretical maximum level of deamidation, displayed a 1000-fold relative increase.

Together, the aforementioned data implied that a significant fraction of eroGFP (and eroGFP-Glyc) was already resident and properly folded in the ER, whereupon the reporter was subsequently removed back to the reducing environment of the cytosol during ER stress. To confirm this finding visually (i.e., directly), we constructed an ER-targeted photoactivatable fluorescent protein (ER-mEos3.2)-expressing yeast strain to specifically follow the fate of an ER-resident reporter while ignoring contributions from new protein synthesis and translocation (Figure 2A). mEos3.2 is a monomeric photoactivatable fluorescent protein that has an excitation maxima at 507nm (Green)[13]. A UV pulse will optically highlight the existing pool of folded reporter by shifting the excitation maxima to 573nm (Red) for pre-existing mEos3.2. Thus, the ER-mEos3.2 detected in the 573nm channel (Red) represents pre-existing reporter, while newly synthesized ER-mEos3.2 will be detected in the 507nm channel (Green). In the absence of stress, ER-mEos3.2 localizes to the ER, both before and after photoconversion (Figures 2B and 2C; Movie S1). But when treated with Tm after photoconversion, ER-mEos3.2 (Red) localized to the cytosol in a time course consistent with that seen for eroGFP reduction (Figure-2D; Movie S2). These last results using ER-mEos3.2 have been independently corroborated by Erik Snapp’s group (see biorxiv manuscript by Lajoie and Snapp). Furthermore, we found that ER-mEos3.2 accumulates in the cytosol in its fluorescent (i.e. correctly folded) state for up to 8 hours after Tm treatment without being degraded (Movie S3). Thus, remarkably, ER stress resulted in pre-existing ER-mEos3.2 protein returning back from the ER to the cytosol in an intact, folded state (as with the two other reporters described above). We termed this retrograde trafficking process “ER protein reflux” and decided to study the phenomenon further through genetics.

**Figure 2.**
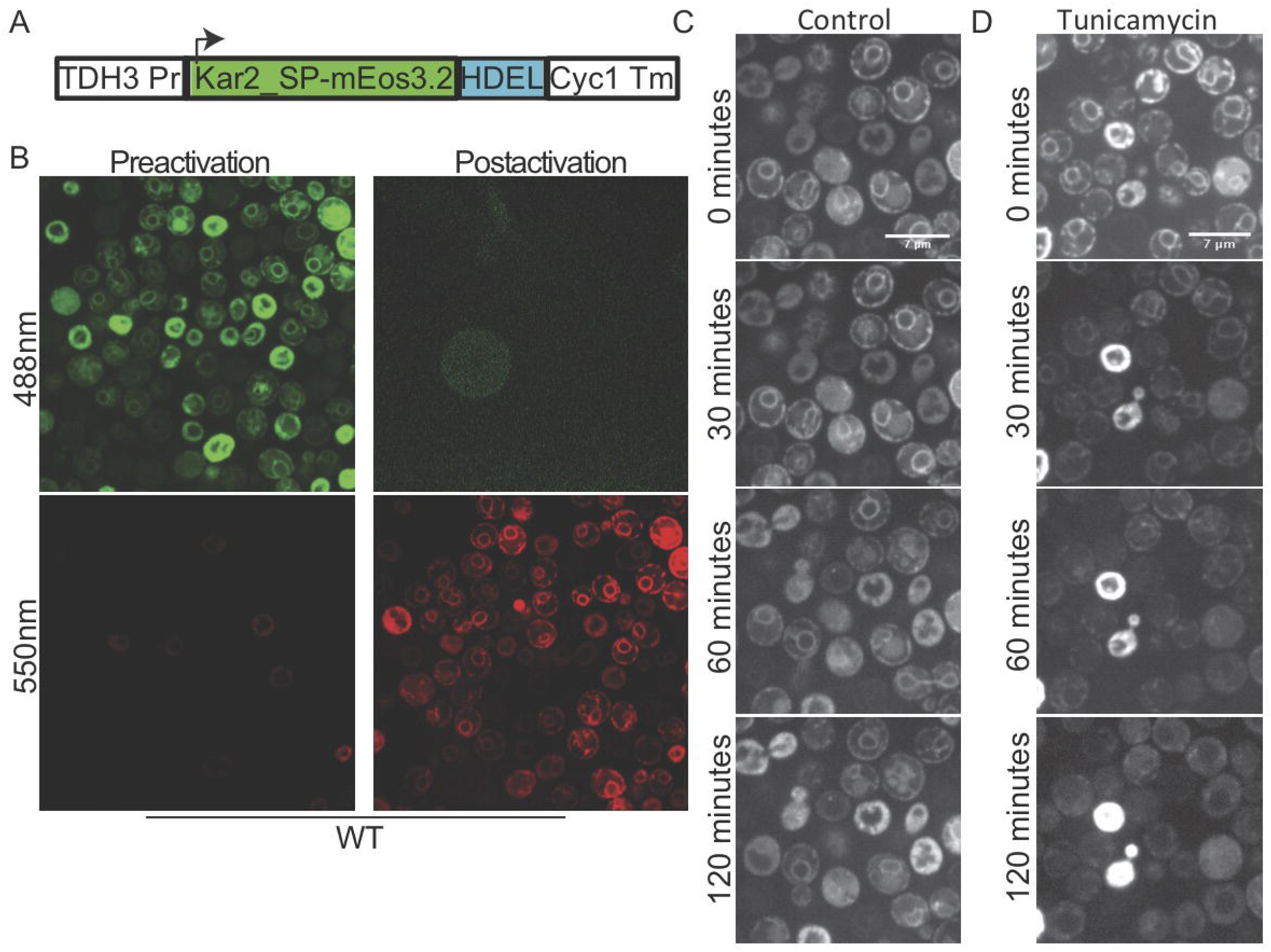
Pre-existing yemEos3 reporter is refluxed from the ER to the cytosol during ER stress. (A) Schematic of the ER-targeted yemEos3.2 construct. (B) Microscopy images of WT cells expressing ER-targeted yemEos3.2, before and after photoconversion in both wavelengths (488nm, 550nm). ER-targeted yemEos3.2 was first photoconverted by UV light in WT cells, then images were taken (550nm) after conversion for the indicated time points, in the absence (C) or presence (D) of Tm.

### Comprehensive Identification of Genes Affecting Reflux of eroGFP

What factors could promote the reflux of eroGFP (and ER-mEos3.2) already targeted to, residing in, and properly folded (i.e., fluorescent) in the ER back to the cytosol in an intact state? To identify genes mediating this reflux phenomenon, we integrated the eroGFP reporter into the *S. cerevisiae* non-essential gene deletion collection [14] and the essential gene DAmP library [15] using synthetic genetic array techniques (Figure S3A)[16]. Using high-throughput flow cytometry [17, 18], we measured the eroGFP ratios in approximately 6000 mutant strains during exposure to Tm and compared the ratios to untreated controls (Tables S1, S2). We observed a range of ratios, consistent with the idea that the mutants can modulate reflux of eroGFP during ER stress. To define hits, we fitted a curve to the difference between replicate measurements and obtained threshold eroGFP ratio values corresponding to p<0.001 (Figure S3B-G and Methods). After treatment with Tm, eroGFP ratios increased to a mean value of 0.57 in wild-type cells, normalized to untreated (Figure 3A; Table S2). This value corresponds approximately to the eroGFP ratio of wild-type cells treated with 0.5 mM DTT (Figure 1B). Exploiting the fact that this regime of Tm treatment caused an intermediate (sub-complete) level of reduction (see Figure 1B), we could identify hundreds of mutants with eroGFP ratios that are both higher (sensitive hits), and lower (resistant hits), than the deflection experienced by wild-type cells (Figure 3A).

**Figure 3.**
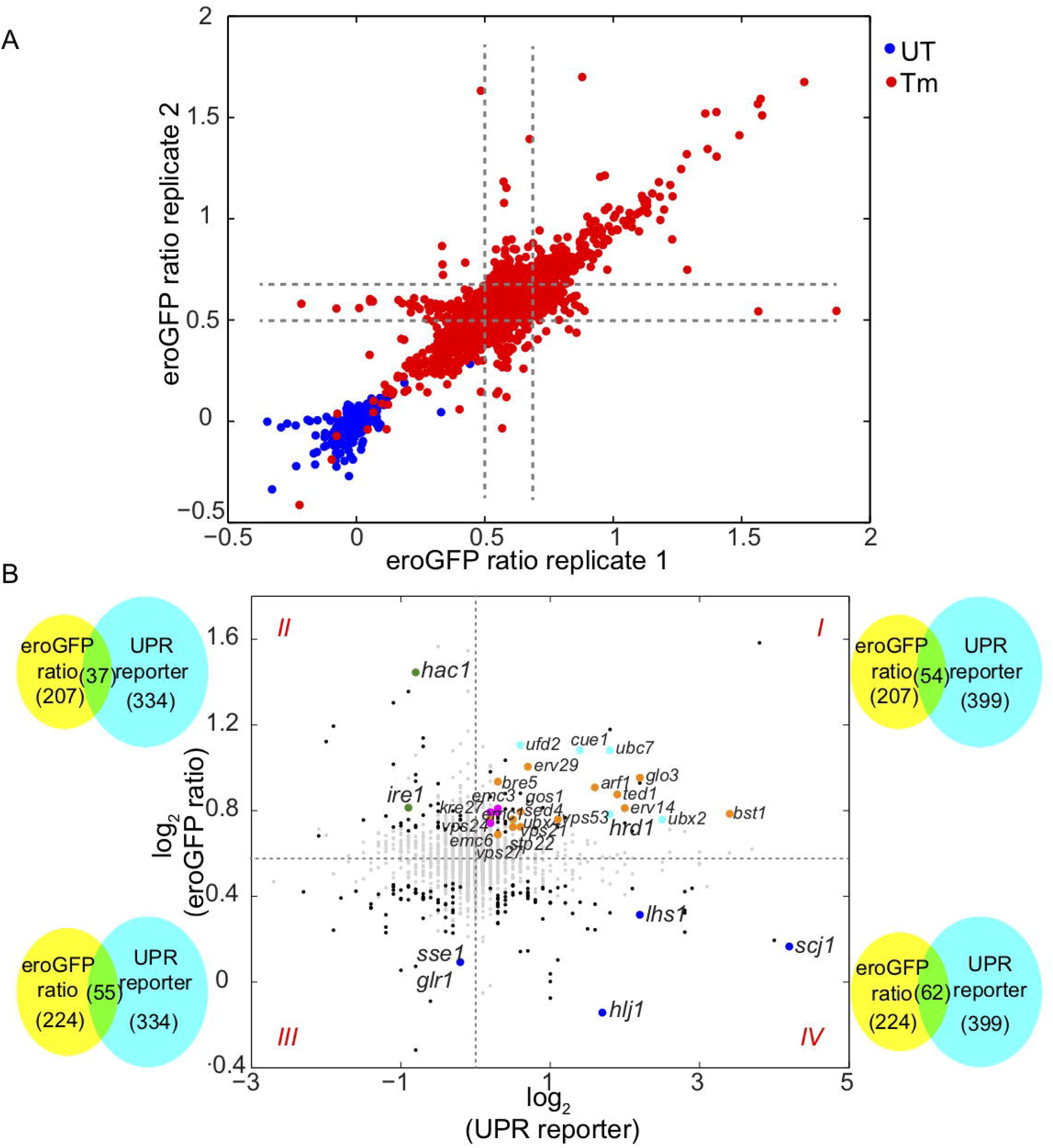
Quantitative screen for genes whose deletion or down-regulation affect eroGFP oxidation. (A) Median eroGFP ratios of the deletion and DAmP libraries with and without Tm (6 μg/mL for 5 h). (B) Scatter plot of median eroGFP ratios after Tm-treatment vs. UPR reporter levels for the non-essential yeast deletion library (Jonikas et al., 2009). The following gene categories are indicated by color: cyan (ERAD), orange (trafficking), magenta (EMC) (Jonikas et al., 2009), blue (ER-resident chaperones), and green (UPR). Venn diagrams indicate overlap for each quadrant.

In a previous genomic screen, basal UPR activity resulting from mutation of most nonessential genes in *S. cerevisiae* was comprehensively measured [19]. Using these datasets, we compared each mutant’s UPR activity to its eroGFP ratio under Tm-induced stress. Unexpectedly, we found that there is minimal global overlap between gene deletions that constitutively induce the UPR and those that significantly perturb eroGFP oxidation during Tm treatment (see Venn diagrams in Figure 3B). However, when we examined mutant subsets grouped by their common molecular functions, we identified three subgroups in which correlations are evident between the two reporters (Figure 3B). Mutants displaying *both* increased UPR activity and increased eroGFP ratios (compared to wild-type) are found in quadrant I; these sub-groups have mutations in genes encoding subunits of the ER membrane complex (EMC), components of the ER-associated degradation (ERAD) system, and activities needed for trafficking throughout the secretory pathway (Figure S3H). Mutants in quadrant IV display decreased eroGFP ratios despite increased UPR activity; these sub-groups have mutations in genes encoding activities supporting N-linked glycosylation in the ER and many ER and cytosolic chaperones and co-chaperones. Identification of quadrant IV mutant chaperone/co-chaperone groups was unexpected and will be addressed in the following section. The UPR-deficient mutants, *ire1*Δ and *hac1*Δ, had greater eroGFP ratio deflection than wild-type upon Tm exposure (quadrant II—as we previously showed in the real-time flow cytometry studies [6]), supporting the expectation that these mutants experience more ER stress relative to wild-type because they cannot trigger a protective UPR.

### Reflux of ER resident proteins requires Cytosolic and ER chaperones

To identify components mediating ER protein reflux, we first focused on ERAD components since many ERAD genes are upregulated during ER stress [4], and their encoded products exert quality control by removing misfolded secretory proteins to the cytosol for subsequent ubiquitylation and degradation by the 26S proteasome. However, as mentioned above, genes associated with the canonical ERAD pathway were not among the resistant hits. Instead, several ERAD-defective mutants are found in quadrant I (increased eroGFP ratios during Tm—i.e., sensitive hits) (Figure 3B, Table S2). In *S. cerevisiae*, the membrane proteins HRD1 and D0A10 are the predominant ERAD ubiquitin-protein E3-ligases whose cytoplasmically-oriented RING domains recruit distinct ubiquitin-conjugating enzymes to cause substrate ubiquitylation [20–23]. Confirming the finding from the screen that these ERAD components are not required for ER protein reflux, we found that eroGFP refluxes to the cytosol after treatment with Tm in both single and double mutants of HRD1 and DOA10 (Figure 4A). To follow the dynamics of the reflux process we used ER-mEos3.2 and found that not only were HRD1 and D0A10 unnecessary for ER reporter reflux to the cytosol, but that in the absence of these ERAD components reflux even occurred in the absence of Tm, with some basal level of reporter seen in the cytosol in untreated cells, and more rapidly in the presence of Tm (Figure 4A-E, Movie S4 and Movie S5).

**Figure 4.**
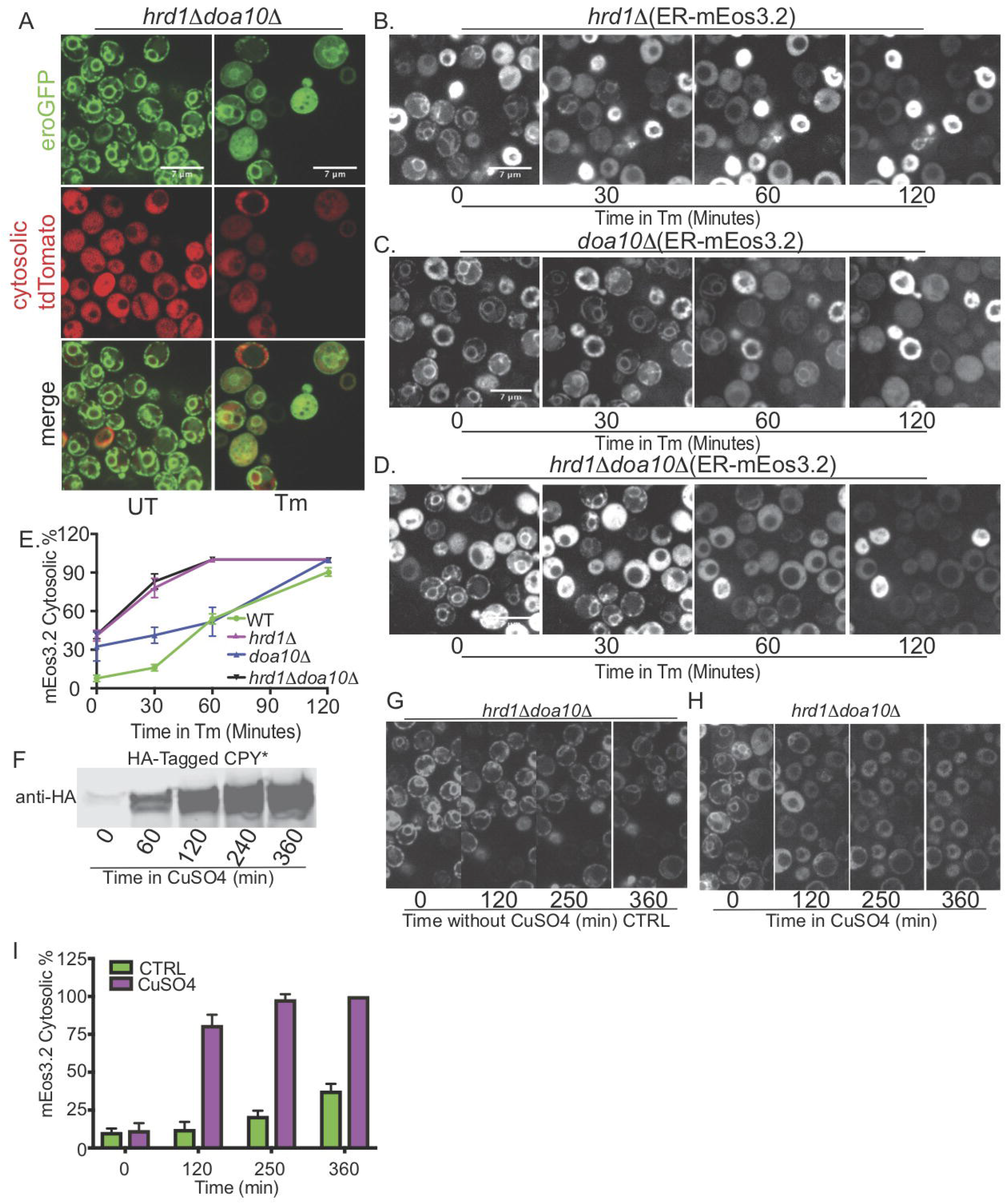
ER protein reflux is not reliant on canonical ERAD machinery. (A) Confocal images for the *hrd1*Δ*doa10*Δ double mutant expressing eroGFP treated with Tm (6μg/mL) for 2 h. Confocal images for *hrd1Δ* (B) *doa10Δ* (C) and *hrd1Δdoa10Δ* double mutant (D) expressing the ER-targeted yemEos3.2 and treated with Tm for the indicated time points. (E) Quantification of images in (B, C and D) of *hrd1Δ*, *doa10Δ* and *hrd1Δdoa10Δ* with ER targeted yemEos3.2. (F) Immunoblot (anti-HA) of protein extracts from wild-type cells expressing HA-tagged CPY* under the cup1 promoter after addition of copper sulfate for the indicated time points. Confocal microscopy images of *hrd1*Δ*doa10*Δ double mutant cells expressing ER targeted mEos3.2 and CPY* under the Cup1 promoter in the absence (G) AND presence of Copper sulfate (H) after photoconversion for the indicated time points. (I) Quantification of images of *hrd1Δdoa10Δ* with ER targeted yemEos3.2 Control or after CuSO4 treatment.

In our previous study, we found that expressing the constitutively misfolded secretory protein CPY* under the Cup1 promoter in ERAD mutants increased the ratio of reduced eroGFP upon addition of copper [6]. Here, we tested whether ER protein reflux could account for the increased eroGFP ratio that we had previously observed. To this end, we monitored the localization of the ER-targeted mEos3.2 in a *hrdlΔdoal0Δ* double mutant strain that expresses CPY* under the Cup1 promoter. We found that in cells expressing CPY*, photoconverted ER-targeted mEos3.2 robustly localized to the cytosol and stayed fluorescent in the cytosol for up to 6 hrs after copper was added (Movie S6, Figure 4F-I).

In sum, it appeared that ER to cytosol reflux of eroGFP (or ER-mEos3.2) during ER stress not only does not rely on ERAD function, but furthermore, mutations in many ERAD components, or the proteasome, appear to *compensatorily* increase the reflux process and recovery of intact reduced eroGFP in the cytosol (i.e., these are all sensitive hits with eroGFP ratios > WT). Moreover, in support of the notion that ER reflux may work in parallel with (or even substitute for) ERAD, the forced overexpression of CPY* was sufficient to cause spontaneous reflux of ER-targeted mEos3.2 in the *hrd1Δdoa10Δ* double mutant. Finally, supporting the notion that eroGFP reflux is not dependent on ERAD, eroGFP appeared to not be polyubiquitinated during Tm provision (unlike CPY*) (Figure S4A,B).

Next, to identify and characterize genes that may mediate the ER protein reflux process, we focused on ER-resident proteins from our screen that showed minimal eroGFP ratio changes during Tm treatment (i.e., resistant hits in quadrant IV, Figure 3B). Through this analysis, HLJ1, an ER-resident tail-anchored co-chaperone, containing a cytosolically-disposed DnaJ domain, stood out as the single, strongest resistant hit among ER-resident proteins. We confirmed that the *hlj1*Δ mutant did not alter eroGFP oxidation after Tm treatment (Figure 5A). Furthermore, in *hlj1*Δ mutants, eroGFP remained in reticular structures that did not overlap with cytosolic tdTomato during Tm treatment, though a small fraction of eroGFP localized to the vacuole even under unstressed conditions (Figure 5B, 5D). As with eroGFP, photoconverted (pre-existing) ER-mEos3.2 remained localized to the ER in the *hlj1*Δ mutant upon Tm treatment (Figure S5A-B). Thus, as predicted, the resistant hits from the genetic screen successfully identified a gene whose product promoted the reflux process. However, in each case the combination of fluorescence microscopy and flow cytometry for eroGFP ratio value need to be compared. For example, a special exception for a resistant hit that leaves reflux unaffected is the *glr1*Δ glutathione oxidoreductase deletion mutant in which Tm still caused eroGFP to reflux to the cytosol (as with WT) but without appreciably changing the eroGFP ratio from untreated (i.e., 0.09; Table S2). Glr1 is responsible for maintaining a reduced cytosol by converting oxidized to reduced glutathione [24, 25] and its absence even causes oxidation of cyto-roGFP (Figure S5C and S5D).

**Figure 5.**
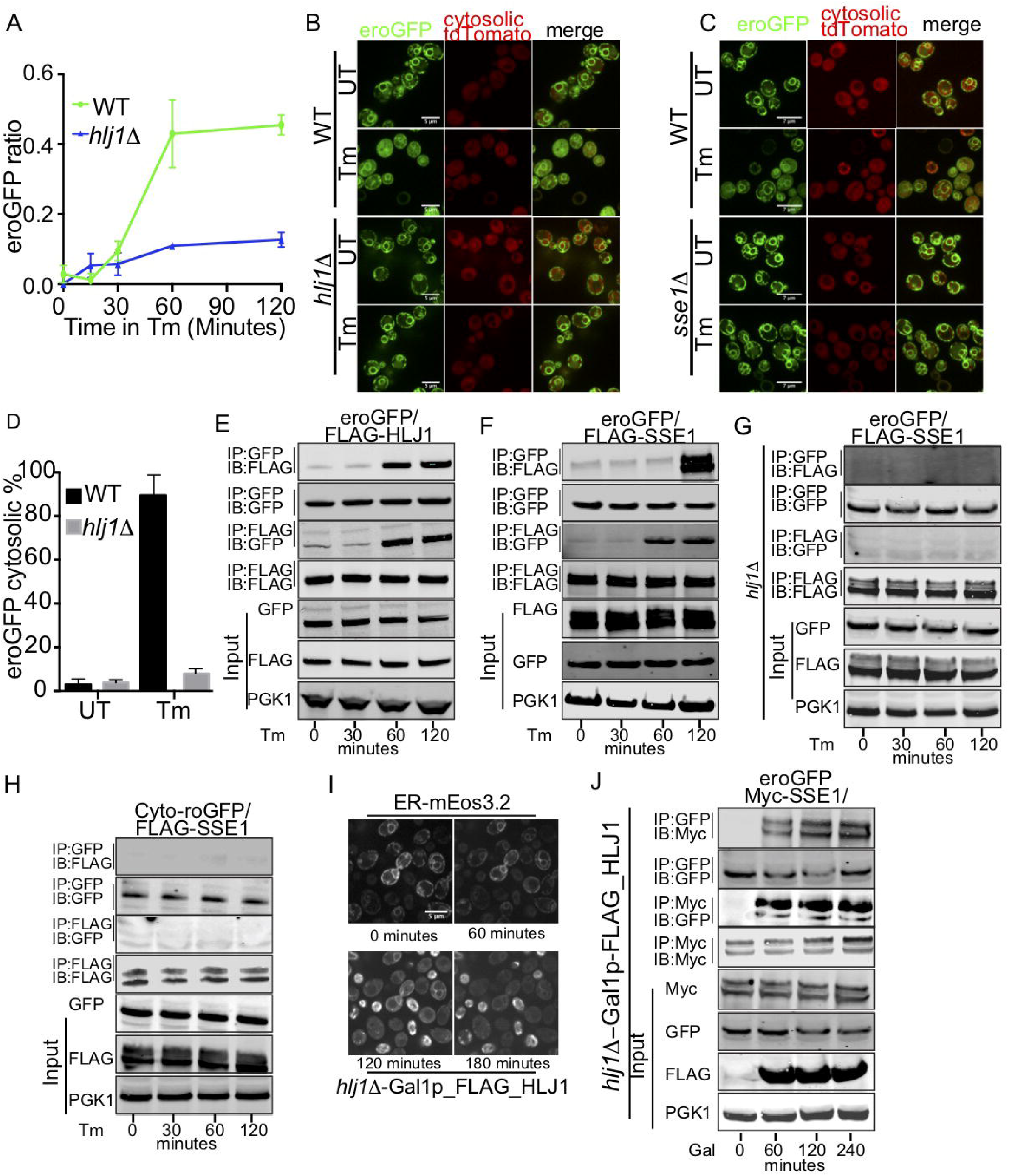
Reflux of ER proteins requires HLJ1 and SSE1. (A) Time course of eroGFP ratios in wild-type and *hlj1*Δ cells expressing eroGFP treated with 6μg/mL Tm. (B) Confocal images of eroGFP in wild-type and *hlj1Δ* treated with Tm (6μg/mL) for 2hrs. (C) Confocal images of WT and *sse1*Δ treated with Tm (6μg/mL) for 2hrs. (D) Quantification of WT and *hlj1*Δ images. Error bars represent S.E.M of two independent experiments. Immunoprecipitation of C-terminal FLAG-tagged HLJ1(E) or FLAG tagged SSE1 (F) and eroGFP in WT cells after treatment with Tm (6μg/mL). (G) Immunoprecipitation of C-terminal FLAG-tagged SSE1 and ER-targeted eroGFP in *hlj1*Δ cells. (H) Immunoprecipitation of C-terminal FLAG-tagged SSE1 and cytosolic roGFP in WT cells treated with Tm (6μg/mL). (I) Confocal images of ER-targeted mEos3.2 in *hlj1*Δ cells overexpressing FLAG-tagged HLJ1 under the Gal1/10 promoter after shifting to galactose-containing media. (J) Immunoprecipitation of C-terminal Myc-tagged SSE1 and ER-targeted eroGFP in *hlj1*Δ cells overexpressing WT-HLJ1 after shifting to galactose-containing media.

In yeast, the ER-resident chaperone KAR2 acts as an *anterograde* molecular ratchet during translocation of secretory proteins through the Sec61 translocon. The binding of KAR2 to a translocating polypeptide on the luminal side of the Sec61 channel prevents it from moving backward [7]. Successful protein translocation requires interaction between KAR2 and the Sec complex via the J domain of SEC63p [21, 26–31]. Perhaps the resistant mutants identified from our screen may promote *retrograde* movement of eroGFP during ER stress-induced reflux? For instance, we found that besides HLJ1, mutations of genes encoding other ER and cytosolic chaperones, co-chaperones, and nucleotide exchange factors (e.g. *Ihs1*Δ, kar2-DAmP, sec66 and *ssel*Δ) also resisted eroGFP reduction and cytosolic relocalization upon Tm treatment (Figure 3B and Table S2). Thus, we hypothesized that these chaperones and co-chaperones may assist eroGFP reflux and should therefore bind this client protein during ER stress. To test this notion, we immunoprecipitated C-terminal FLAG-tagged HLJ1 in WT cells expressing eroGFP and found increasing interactions between HLJ1 and eroGFP over the time course of Tm treatment (Figure 5E); this increase in the interaction between HLJ1 and eroGFP correlated with the kinetics of eroGFP protein reflux in these cells (Figure S1D).

A null mutation in SSE1 (also a resistant hit), which encodes a cytosolic nucleotide exchange factor and acts as a “holdase” activity, also strongly resisted eroGFP reflux during ER stress [32–36] (Figure 5C). Moreover, we found that eroGFP progressively interacts over time with FLAG-tagged SSE1 under Tm treatment in WT cells (Figure 5F); this interaction between SSE1 and eroGFP became abrogated in *hlj1*Δ cells (Figure 5G). Furthermore, to eliminate the possibility that SSE1 may interact with eroGFP *after* it has already translocated to the cytosol (i.e., in a non-reflux manner), we employed the cytosolic version of roGFP (cyto-roGFP) described above and did not observe an interaction between SSE1 and this cytosolic roGFP variant (Figure 5H). Importantly, this last result supports the conclusion that under ER stress the reporter has to originate from *inside* the ER lumen to interact later with cytosolic SSE1.

Because all preceding data supported the notion that HLJ1 is *necessary* for ER protein reflux during ER stress, we next asked whether HLJ1 is *sufficient* to cause reflux of ER proteins to the cytosol upon forced expression. Because HLJ1 is reported to be lethal when constitutively overexpressed in *S. cerevisiae* (Stepanek P, et al. (1995), SGD paper) we placed a FLAG-tagged-HLJ1 under the control of the inducible GAL1/10 promoter and followed its expression for several hours (Figure S5E). We found that 2-4 hours after shifting cells to galactose a significant fraction of ER-targeted eroGFP spontaneously accumulated in the cytosol, without the need for exogenous ER stress agents (Figure S5F). To confirm the ER origin of the refluxed reporter, we expressed ER-targeted-mEos3.2 in the inducible FLAG-HLJ1 strain and followed the reporter’s fate during growth on galactose; at 3 hours after shift to galactose media the majority of cells displayed a spontaneous cytosolic localization of the photo-converted mEos3.2 (Figure 5I). Finally, we observed that in cells overexpressing HLJ1p there was an increased reciprocal physical interaction (through co-immunoprecipitation) between eroGFP and the cytosolic chaperone SSE1p, but without the need for exogenous ER stress agents (Figure 5J). Thus, HLJ1p was both necessary and sufficient to promote ER reflux.

Finally, we inquired into the scope of the reflux process—i.e., does it extend to other ER proteins (besides the FP reporters used in this study)? In theory, other ER-targeted proteins may also reflux to the cytosol during ER stress. To test this notion, we monitored the localization of ER endogenous protein disulfide isomerase (PDI1) and peptidyl-prolyl cis-trans isomerase (CPR5) during ER stress. Following treatment with Tm, we found that both CPR5 and PDI1 became progressively enriched over time in the cytosolic fraction (i.e. S100) in WT cells, with similar kinetics to eroGFP, while the ER membrane protein, Spf1 (predictably) remained in the membrane fraction (i.e.-P100)(as with Hrd1-eroGFP) (Figure 1). Importantly, in *hlj1*Δ cells, both CPR5 and PDI1 remained stably in the membrane fraction (as did eroGFP)(Figure S5G). Finally, applying the inducible ER stress regime of expressing the CPY* mutant using the CUP promoter in the *hrd1*Δ*doa10*Δ double mutant, we found increased recovery (compared to expression of CPY) of CPR5, PDI1, and eroGFP in the cytosolic fraction (Figure S5H).

To conclude our study, we also asked whether cells resistant to ER protein expulsion may perhaps be more susceptible to ER stress than wildtype cells. To this end, we performed a yeast viability assay by treating WT and *hlj1*Δ cells with Tm for 4hrs and then spread the same optical density (O.D) of cells on YPD plates and then counted the viable colonies on the plate. We found that in this regime WT cells were almost ~75% viable, while in *hlj1*Δ and *sse1*Δ mutants only 53% and 50% of the cells were viable respectively (Figure S5I). Plate sensitivity assay also confirmed that *hlj1*Δ and *sse1*Δ are more susceptible to ER stress (Figure S5J). These data were consistent with the possibility that blocking ER protein expulsion may cause yeast cells to become more sensitive to acute ER stress.

## DISCUSSION

In response to ER stress, several cellular stress response pathways, such as ERAD and the UPR, become activated to restore protein-folding homeostasis in the ER. Here, using a combination of fluorescence microscopy and high-throughput flow cytometry of yeast gene deletion libraries to measure fluorescent changes in an ER-targeted redox-sensitive GFP (eroGFP), we identify a new cellular stress response pathway in which ER-resident proteins are removed in an intact state back into the cytosol through a chaperone-mediated manner. We have termed this process “ER protein reflux”.

We had originally adapted eroGFP to track ambient redox state as a proxy measure of ER physiological health and previously shown that reduction of the normally oxidized eroGFP occurred in individual cells (wildtype and a few select mutants) during ER stress induced by inhibiting N-linked glycosylation, inositol deprivation (in UPR mutants), and by expressing misfolded secretory proteins (e.g., CPY*)[6]. Here we have confirmed our previous conclusion that reduction of eroGFP occurs after its translocation and maturation in the ER [6](i.e., in pre-existing, ER-resident reporter) by using an eroGFP variant in which an N-linked glycosylation signal is engineered at the C-terminus of the reporter (called eroGFP-Glyc). When expressed through a GAL 1/10 inducible system in which new transcription and translation is shut off through application of glucose followed by induction of ER stress, pre-existing eroGFP-Glyc became deglycosylated in the cytosol and eroGFP-Glyc’s asparagine-linked glycan converted into an aspartate residue as measured using 2D electrophoresis and mass spectrometry. In direct support of these biochemical data, we observed using live cell imaging, a photoconverted ER-resident mEos3.2 reflux into the cytosol during ER stress, with similar kinetics to the reduction of eroGFP observed during similar modes of ER stress.

By measuring eroGFP ratios in yeast gene deletion libraries using high-throughput flow-cytometry, we first identified both sensitive and resistant mutants that precisely change eroGFP oxidation levels to varying degrees during ER stress. eroGFP ratio changes in the mutants showed limited overlap with changes measured by a UPR reporter, validating the use—at the genomic scale—of the utility of a redox-responsive reporter to provide orthogonal information about a physiological secretory pathway function to a corrective signaling pathway (the UPR). Then by validating the hits from the screen using fluorescence microscopy, we uncovered the chaperone-mediated basis of the ER-protein reflux phenomenon. This reflux phenomenon was missed in our prior eroGFP work (that only utilized flow cytometry of a small group of mutants) in which we had proposed an unnecessarily constrained interpretation of eroGFP oxidation changes occurring within the ER [6]. The central feature of ER-protein reflux appears to that soluble, folded proteins (i.e. fluorescent in the case of the reporters) are removed from the ER without becoming degraded (unlike in the ERAD process). Furthermore, upon their reflux to the cytosol, the pre-existing reporters that originated from the ER remain fluorescent (i.e. folded) for at least 8 hours. Further distinguishing reflux from ERAD, we found that eroGFP, eroGFP-Glyc, ER-mEos3.2 and endogenous ER proteins are returned to the cytosol in the absence of the canonical ERAD-associated E3 ubiquitin-protein ligases, HRD1 and D0A10.

ER reflux, however, is strikingly abrogated in the absence of HLJ1, a tail-anchored ER-membrane co-chaperone with a cytosolically-disposed J domain. HLJ1 is known to have overlapping roles with another co-chaperone containing a cytosolic J-domain, YDJ1, in promoting the removal and ubiquitin-dependent degradation of the ER membrane protein CFTR[37]. Intriguingly, *ydj1*Δ is a sensitive hit in our screen (Table S2), supporting the notion that it does not play an overlapping role with YDJ1 in reflux. It is thus conceivable that HLJ1 has auxiliary role(s) in ER protein quality control beyond those previously ascribed to it (e.g., through mediating reflux). More generally, besides *hlj1*Δ, mutations in other ER-resident chaperones and co-chaperones (e.g. *lhs1*Δ, *kar2*-DAmP, *scj1*Δ)(Figure 3B Quadrant IV, Table S2) also resist eroGFP oxidation changes during ER stress to varying degrees. Thus, these gene products may also play critical roles in mediating and regulating ER protein reflux; future investigations into the functions of these gene products may provide a more comprehensive understanding of ER protein reflux. For instance, besides HLJ1, we found notably that cytosolic SSE1, a Hsp110 chaperone/nucleotide-exchange factor/”holdase”, which also emerged from the resistant hits in our screen can bind eroGFP in an HLJ1-dependent manner under ER stress (and importantly only if the reporter is first targeted to the ER rather than originating from the cytosol).

Finally, we note that we do not yet fully understand how folded ER-resident proteins can re-emerge in a folded state (i.e., fluorescent in the case of eroGFP, eroGFP-Glyc, and mEos3.2) in the cytosol. While it is conceivable that reflux substrates stay folded during this retrograde transit across the ER membrane (perhaps transiting through an ER channel), no obvious candidate genes meeting the expected criteria of a putative channel were identified as (resistant) hits in our screens. Therefore, while remaining agnostic to the involvement of a putative channel, we suggest that a relay shuttle of ER-resident and cytosolic chaperones (e.g. KAR2, HLJ1, SSE1) may allow for partial unfolding in the ER, transit across the ER, and refolding in the cytosol of ER luminal proteins during ER stress. Such a process is intriguingly reminiscent of, but directionally opposite, to that of the chaperone-assisted molecular ratchet that promotes translocation of nascent secretory proteins [7]. Furthermore, putative functional homologs of HLJ1 in humans, DnaJB12 and DnaJB14, which were shown to mediate retrograde trafficking/entry into the cytosol from the ER of non-enveloped viruses [38], may also promote ER stress-induced protein reflux in mammalian cells. In this speculative view, the ER reflux machinery may perhaps be usurped by some viruses to gain entry to the cytosol.

While the larger scope of ER reflux remains to be defined, the process could conceivably extend to many endogenous ER-resident proteins (and maturing secretory cargo). Indeed, our finding that PDI1 and CPR5 proteins are recovered (through an HLJ1-dependent mechanism) in the cytosol during ER stress is consistent with the possibility that these proteins are also reflux substrates (and provides an explanation for previous enigmatic reports of how ER chaperones could be found in the cytosol)[39–41]. Physiologically, it is conceivable that ER protein reflux, by clearing the ER of luminal proteins during ER stress, may have an adaptive benefit and may work in parallel with other ER protein quality control mechanisms such as ERAD and protein translocational attenuation referred to as pre-emptive quality control (pre-QC) [11]. Indeed, the decreased viability of *sse1* and *hlj1* mutants under ER stress is consistent with this possibility. It is also conceivable that ER protein reflux may be integrated into the binary cell fate decisions made by cells of higher eukaryotes once ER stress levels reach critical thresholds (manuscript in preparation). Future studies will address such mechanistic and physiological questions.

## MATERIALS AND METHODS

### Plasmid Construction

eroGFP-Glyc was constructed using the Quikchange lightning kit (Agilent Technologies). The glycosylation site was engineered in the second residue of the 9 amino acid linker between the C-terminus of GFP and the HDEL retrieval sequence. eroGFP^ND^ was also constructed using Quikchange lightning kit to change the asparagine in eroGFP-Glyc to an aspartate. Hrd3-eroGFP was constructed using synthetic DNA from Gene Art (Life Technologies) containing eroGFP followed by residues 767-833 of HRD3. ER-yemEos3.2 was constructed by PCR amplification of yemEos3.2 (a gift from Erik Snapp) using forward oligos containing the first 20 amino acids of yemEos3.2 and reverse oligos of the last 20 amino acids with addition of a HDEL sequence. This construct was then cloned in pRS416 carrying the Kar2 signal peptide by BamHI and XbaI. HLJ1, and SSE1 were PCR-tagged in the chromosome under their native promoter with either 3XFLAG-tag or 3XHA-tag using pFA6a-3XFLAG-NAT or pFA6a-3HA-His3MX6 plasmids.

### High-Throughput Flow Cytometry

For all growth conditions described below, yeast strains were grown in 80 μL of SD complete media supplemented with myo-inositol (Sigma-Aldrich) at 100 μg/ml. For tunicamycin (Tm) experiments, Tm was added to the media at 6 μg/mL. Strains were inoculated from 384-colony agar plates to 384-well liquid cultures using a RoToR HDA robot (Singer Instrument Company Limited). The cultures were grown for 36 hours to saturation in a DTS-4 microplate thermoshaker (Appropriate Technical Resources). They were then diluted 1:400 using a BioMek liquid-handling robot (Beckman Coulter, Inc.) and grown to mid-log phase for 10 hours after which they were diluted 1:10 into media with or without Tm. After 5 hours of growth, cultures were loaded on Becton Dickinson High Throughput Sampler (BD), which injected cells from each well into a LSRII flow cytometer (BD). eroGFP fluorescence was measured according to [6].

### Light Microscopy

Yeast were imaged as previously described [6] with the exception that a 561nm laser line was used to excite tdTomato. Cells carrying the ER-yemEos3.2 were growing for mid log phase and then immobilized on concanavalin A glass bottoms. Cells then were imaged using the 488-channel followed by 1 minute with DAPI for photoconversion. Tm were then added and cells were imaged for 2hrs using the High Speed Widefield microscope. All other images were captured using a spinning disk confocal at the Nikon Imaging Center (UCSF).

### Immunoblots and immunoprecipitation

For yeast immunoprecipitation studies, we followed the exact protocol as in [42] with one modification: after collecting the cells, proteins were extracted by disrupting the cells twice with glass beads in the lysis buffer for 45 seconds in a TissueLyser-II (QIAGEN). Immunoblots were performed as previously described [6]. Antibodies used included: rabbit anti GFP; mouse anti PGK1 (Thermo Fischer), Monoclonal ANTI-FLAG^®^ M2 antibody (Sigma-Aldrich) and HA Tag Antibody (Ptglabs). Antibody binding was detected by using near-infrared-dye-conjugated secondary antibodies (LiCor) on the LI-COR Odyssey scanner.

## ACKNOWLEDGEMENTS

We thank Erik Snapp for the yemEos3.2 plasmid, stimulating discussions, and exchanging data; Jeff Brodsky for providing the *hrd1*Δ, *doa10*Δ double mutant strain; and David Breslow for the cytosolic tdTomato plasmid. AI was supported by a JDRF postdoctoral fellowship and P.I.M. was supported by a National Science Foundation Graduate Research Fellowship and a Ruth L. Kirschstein National Research Service Award. F.R.P was supported by grants from the NIH (Director’s New Innovator Award DP2 0D001925 and R01DK095306) and a Career Award in the Biomedical Sciences from the Burroughs Wellcome Foundation.

