## Supplemental table 1 for "Chaperone-Mediated Reflux of Secretory Proteins to the Cytosol During Endoplasmic Reticulum Stress"

**Table S1** eroGFP ratios for the gene libraries during normal growth conditions

UT indicates untreated
Rep indicates individual replicates

Hit Legend 0: not a hit
1: hit, eroGFP more reduced
-1: hit, eroGFP less reduced

| ORF | UT Rep 1 | UT Rep 2 | UT ave | Hit |
|---|---|---|---|---|
| VMA9 | 0.007 | 0.039 | 0.023 | 0 |
| SRN2 | -0.018 | 0.024 | 0.003 | 0 |
| FLC2 | 0.010 | 0.016 | 0.013 | 0 |
| YML084W | -0.002 | 0.004 | 0.001 | 0 |
| RBG1 | -0.002 | 0.016 | 0.007 | 0 |
| SML1 | 0.010 | 0.017 | 0.014 | 0 |
| ATS1 | 0.032 | 0.042 | 0.037 | 1 |
| YML037C | -0.010 | 0.011 | 0.000 | 0 |
| YAL004W | 0.000 | -0.001 | -0.001 | 0 |
| OST6 | 0.017 | 0.031 | 0.024 | 0 |
| YAR029W | -0.012 | 0.007 | -0.002 | 0 |
| GIS4 | 0.005 | 0.011 | 0.008 | 0 |
| SPO75 | -0.009 | 0.008 | 0.000 | 0 |
| YMR010W | -0.041 | -0.035 | -0.038 | -1 |
| SPA2 | -0.053 | -0.097 | -0.075 | -1 |
| MSS1 | 0.009 | -0.001 | 0.004 | 0 |
| YLR118C | 0.003 | 0.027 | 0.015 | 0 |
| CHA4 | 0.024 | 0.022 | 0.023 | 1 |
| NDE1 | 0.010 | -0.001 | 0.004 | 0 |
| RAX2 | -0.001 | 0.011 | 0.005 | 0 |
| ATG16 | 0.001 | 0.002 | 0.002 | 0 |
| FYV7 | 0.009 | 0.039 | 0.024 | 0 |
| SIP18 | -0.016 | -0.015 | -0.016 | 0 |
| OSW2 | -0.008 | 0.003 | -0.002 | 0 |
| GCV2 | 0.012 | 0.017 | 0.015 | 0 |
| UBR2 | 0.031 | 0.022 | 0.027 | 1 |
| ERG2 | 0.033 | 0.032 | 0.032 | 1 |
| LOT6 | 0.002 | 0.004 | 0.003 | 0 |
| UBP8 | 0.015 | 0.022 | 0.019 | 0 |
| YLL054C | -0.010 | -0.010 | -0.010 | 0 |
| YHM2 | 0.003 | -0.007 | -0.002 | 0 |
| YAL068C | 0.002 | 0.015 | 0.008 | 0 |
| YPS1 | -0.013 | 0.023 | 0.005 | 0 |
| OAF1 | -0.004 | 0.010 | 0.003 | 0 |
| YML083C | -0.002 | 0.013 | 0.006 | 0 |
| FUN12 | -0.007 | -0.003 | -0.005 | 0 |
| CMP2 | -0.014 | 0.001 | -0.007 | 0 |
| FUN30 | 0.006 | 0.024 | 0.015 | 0 |
| AMD1 | 0.000 | 0.003 | 0.001 | 0 |
| SSA1 | 0.006 | 0.011 | 0.008 | 0 |
| YML018C | 0.006 | 0.011 | 0.009 | 0 |
| PRM9 | -0.017 | -0.001 | -0.009 | 0 |
| TRM12 | -0.007 | -0.001 | -0.004 | 0 |
| MMM1 | 0.024 | 0.032 | 0.028 | 1 |
| HXT2 | 0.000 | -0.001 | -0.001 | 0 |
| his3D::KAN | -0.002 | 0.000 | -0.001 | 0 |

| | | | | |
|---|---|---|---|---|
| AVL9 | 0.018 | 0.034 | 0.026 | 0 |
| MIH1 | -0.001 | 0.019 | 0.009 | 0 |
| HRT3 | -0.007 | 0.005 | -0.001 | 0 |
| YMR147W | -0.005 | -0.003 | -0.004 | 0 |
| EMP70 | -0.015 | -0.023 | -0.019 | 0 |
| HLJ1 | -0.020 | -0.013 | -0.016 | 0 |
| ECM5 | -0.011 | 0.008 | -0.002 | 0 |
| YLR053C | -0.001 | -0.002 | -0.002 | 0 |
| SGS1 | -0.003 | -0.002 | -0.003 | 0 |
| IZH3 | -0.019 | -0.012 | -0.016 | 0 |
| INP1 | -0.008 | -0.029 | -0.019 | 0 |
| THI73 | -0.003 | 0.003 | 0.000 | 0 |
| MRE11 | 0.023 | 0.033 | 0.028 | 1 |
| YLL053C | -0.003 | -0.007 | -0.005 | 0 |
| YCR095W-A | -0.007 | -0.011 | -0.009 | 0 |
| SEO1 | -0.005 | 0.025 | 0.010 | 0 |
| YAL049C | 0.000 | 0.022 | 0.011 | 0 |
| YML082W | -0.006 | 0.004 | -0.001 | 0 |
| FUN19 | 0.005 | 0.014 | 0.009 | 0 |
| YML058C-A | -0.001 | -0.008 | -0.005 | 0 |
| YAL018C | -0.001 | 0.012 | 0.005 | 0 |
| SRC1 | 0.001 | 0.000 | 0.000 | 0 |
| VPS8 | -0.017 | -0.018 | -0.018 | 0 |
| PSP2 | -0.012 | 0.007 | -0.003 | 0 |
| YAR030C | -0.010 | 0.004 | -0.003 | 0 |
| GLO1 | 0.005 | 0.008 | 0.007 | 0 |
| COX17 | -0.003 | 0.006 | 0.002 | 0 |
| CLU1 | 0.005 | 0.010 | 0.008 | 0 |
| YLL023C | 0.003 | -0.002 | 0.000 | 0 |
| MRPL3 | 0.026 | 0.002 | 0.014 | 0 |
| HOG1 | -0.014 | 0.003 | -0.006 | 0 |
| SUB1 | -0.022 | 0.018 | -0.002 | 0 |
| KIN2 | -0.011 | 0.007 | -0.002 | 0 |
| YMR148W | 0.009 | 0.021 | 0.015 | 0 |
| SRL2 | -0.003 | 0.004 | 0.001 | 0 |
| DNF3 | -0.007 | 0.008 | 0.001 | 0 |
| YLR065C | -0.001 | 0.008 | 0.003 | 0 |
| MMT1 | -0.006 | 0.003 | -0.002 | 0 |
| YLR049C | 0.008 | 0.007 | 0.007 | 0 |
| SPG5 | 0.005 | 0.009 | 0.007 | 0 |
| IRC25 | 0.059 | 0.083 | 0.071 | 1 |
| PFK2 | -0.003 | 0.002 | -0.001 | 0 |
| CMS1 | 0.006 | 0.008 | 0.007 | 0 |
| MRPL44 | 0.007 | 0.000 | 0.003 | 0 |
| AQY2 | 0.000 | -0.007 | -0.004 | 0 |
| RPL20A | 0.013 | 0.009 | 0.011 | 0 |
| YAL066W | -0.002 | 0.020 | 0.009 | 0 |
| YPS3 | -0.006 | 0.014 | 0.004 | 0 |
| GEM1 | 0.028 | 0.033 | 0.031 | 1 |
| YML081W | 0.042 | 0.038 | 0.040 | 1 |
| GIP4 | -0.006 | -0.007 | -0.007 | 0 |
| IMD4 | 0.004 | 0.005 | 0.004 | 0 |
| PSK1 | -0.006 | 0.009 | 0.001 | 0 |

| | | | | |
|---|---|---|---|---|
| YML035C-A | -0.005 | 0.006 | 0.000 | 0 |
| NUP60 | 0.036 | 0.032 | 0.034 | 1 |
| PPZ1 | -0.006 | -0.006 | -0.006 | 0 |
| YAT1 | -0.004 | 0.005 | 0.001 | 0 |
| YML003W | -0.003 | 0.000 | -0.002 | 0 |
| PSR1 | -0.010 | 0.001 | -0.005 | 0 |
| BUD22 | -0.007 | 0.002 | -0.003 | 0 |
| SSA2 | 0.000 | -0.007 | -0.003 | 0 |
| CSI1 | 0.003 | -0.010 | -0.004 | 0 |
| YLR112W | 0.004 | 0.029 | 0.016 | 0 |
| IOC2 | -0.019 | -0.005 | -0.012 | 0 |
| YIM2 | 0.005 | 0.010 | 0.007 | 0 |
| GAL2 | 0.009 | 0.017 | 0.013 | 0 |
| INP2 | 0.010 | 0.013 | 0.012 | 0 |
| YLR064W | -0.013 | 0.001 | -0.006 | 0 |
| YMR178W | -0.006 | -0.004 | -0.005 | 0 |
| RPS0B | 0.003 | 0.029 | 0.016 | 0 |
| GYL1 | 0.007 | 0.001 | 0.004 | 0 |
| YEH2 | -0.023 | -0.003 | -0.013 | 0 |
| YMR206W | -0.003 | 0.000 | -0.002 | 0 |
| YLR001C | -0.006 | 0.010 | 0.002 | 0 |
| YMR226C | 0.005 | 0.000 | 0.002 | 0 |
| FRE6 | 0.002 | 0.009 | 0.006 | 0 |
| ZRC1 | -0.004 | -0.020 | -0.012 | 0 |
| YAL065C | 0.005 | 0.027 | 0.016 | 0 |
| YLR122C | -0.003 | 0.021 | 0.009 | 0 |
| YAL046C | -0.006 | 0.007 | 0.001 | 0 |
| DUS1 | -0.019 | -0.003 | -0.011 | 0 |
| SNC1 | -0.005 | 0.007 | 0.001 | 0 |
| SPC2 | 0.002 | 0.000 | 0.001 | 0 |
| NTG1 | -0.002 | -0.001 | -0.001 | 0 |
| YML033W | -0.004 | 0.008 | 0.002 | 0 |
| SWD1 | -0.024 | -0.045 | -0.035 | -1 |
| TRM9 | 0.060 | 0.112 | 0.086 | 1 |
| YAR037W | -0.018 | -0.002 | -0.010 | 0 |
| YML002W | 0.004 | 0.001 | 0.002 | 0 |
| YEH1 | -0.004 | -0.001 | -0.003 | 0 |
| ERG5 | 0.009 | 0.005 | 0.007 | 0 |
| PAU17 | 0.005 | -0.004 | 0.001 | 0 |
| PEX12 | -0.004 | -0.020 | -0.012 | 0 |
| YLR111W | 0.003 | 0.007 | 0.005 | 0 |
| YET2 | -0.008 | 0.009 | 0.000 | 0 |
| GIS3 | -0.012 | 0.001 | -0.006 | 0 |
| IMP1 | 0.023 | -0.113 | -0.045 | 0 |
| EMP46 | -0.008 | 0.011 | 0.001 | 0 |
| MSS11 | 0.002 | -0.006 | -0.002 | 0 |
| YLR063W | -0.007 | 0.011 | 0.002 | 0 |
| SPT21 | 0.036 | 0.050 | 0.043 | 1 |
| FRE8 | 0.014 | 0.012 | 0.013 | 0 |
| MRPL24 | -0.004 | -0.004 | -0.004 | 0 |
| PSR2 | -0.005 | 0.003 | -0.001 | 0 |
| HFA1 | -0.007 | 0.000 | -0.003 | 0 |
| AYT1 | -0.006 | 0.006 | 0.000 | 0 |

| | | | | |
|---|---|---|---|---|
| MTF1 | -0.078 | -0.086 | -0.082 | -1 |
| YLL047W | 0.009 | -0.002 | 0.004 | 0 |
| YMR244W | 0.006 | -0.009 | -0.001 | 0 |
| GDH3 | 0.011 | 0.026 | 0.018 | 0 |
| YLR123C | -0.009 | 0.028 | 0.010 | 0 |
| YAL045C | -0.002 | 0.019 | 0.009 | 0 |
| YML079W | -0.014 | 0.009 | -0.003 | 0 |
| MYO4 | 0.008 | 0.002 | 0.005 | 0 |
| CYB2 | -0.004 | -0.004 | -0.004 | 0 |
| SYN8 | -0.026 | -0.012 | -0.019 | 0 |
| RAD52 | 0.022 | 0.024 | 0.023 | 1 |
| BUD14 | 0.019 | 0.022 | 0.020 | 0 |
| UBX2 | -0.001 | 0.011 | 0.005 | 0 |
| YAR040C | -0.015 | 0.000 | -0.007 | 0 |
| YPT7 | -0.012 | -0.008 | -0.010 | 0 |
| PUF3 | 0.003 | 0.010 | 0.006 | 0 |
| SOK2 | 0.014 | 0.015 | 0.015 | 0 |
| HSP104 | 0.010 | -0.002 | 0.004 | 0 |
| YMR027W | 0.000 | -0.009 | -0.004 | 0 |
| AHP1 | 0.005 | 0.043 | 0.024 | 0 |
| ARA2 | 0.001 | 0.024 | 0.012 | 0 |
| NYV1 | -0.004 | 0.004 | 0.000 | 0 |
| YIM1 | -0.014 | 0.008 | -0.003 | 0 |
| SIC1 | 0.026 | 0.026 | 0.026 | 1 |
| YMR166C | 0.007 | 0.022 | 0.015 | 0 |
| BUD28 | 0.028 | 0.053 | 0.041 | 1 |
| CTL1 | 0.000 | 0.014 | 0.007 | 0 |
| YLR046C | -0.001 | 0.001 | 0.000 | 0 |
| RPL36A | 0.029 | 0.039 | 0.034 | 1 |
| POM34 | 0.001 | 0.015 | 0.008 | 0 |
| YMR210W | -0.001 | 0.006 | 0.003 | 0 |
| MHT1 | -0.004 | 0.002 | -0.001 | 0 |
| RPS10B | 0.004 | 0.012 | 0.008 | 0 |
| RNP1 | -0.003 | 0.001 | -0.001 | 0 |
| YMR245W | -0.015 | -0.012 | -0.013 | 0 |
| BDH2 | 0.004 | 0.017 | 0.011 | 0 |
| YLR124W | -0.008 | 0.016 | 0.004 | 0 |
| GCV3 | -0.029 | -0.031 | -0.030 | -1 |
| CPR3 | -0.006 | 0.005 | 0.000 | 0 |
| FRT2 | -0.002 | -0.002 | -0.002 | 0 |
| YML053C | 0.001 | -0.002 | 0.000 | 0 |
| DEP1 | -0.032 | -0.024 | -0.028 | -1 |
| YML030W | 0.002 | 0.024 | 0.013 | 0 |
| ADE1 | 0.023 | 0.015 | 0.019 | 0 |
| YML013C-A | -0.055 | -0.036 | -0.046 | -1 |
| SWH1 | -0.004 | 0.003 | 0.000 | 0 |
| MIC17 | -0.004 | 0.006 | 0.001 | 0 |
| YLL014W | 0.018 | 0.027 | 0.022 | 0 |
| SPO20 | -0.002 | 0.002 | 0.000 | 0 |
| TPO1 | 0.008 | -0.003 | 0.002 | 0 |
| FAR8 | 0.006 | -0.001 | 0.002 | 0 |
| YLR108C | 0.003 | 0.045 | 0.024 | 0 |
| ARG80 | -0.006 | 0.023 | 0.008 | 0 |

| | | | | |
|---|---|---|---|---|
| SUL2 | 0.006 | 0.016 | 0.011 | 0 |
| NUP53 | 0.000 | 0.005 | 0.003 | 0 |
| FMP25 | 0.008 | 0.007 | 0.007 | 0 |
| MLH1 | 0.013 | 0.016 | 0.014 | 0 |
| RPL22A | 0.033 | 0.005 | 0.019 | 0 |
| RGM1 | 0.004 | 0.017 | 0.011 | 0 |
| PDC1 | -0.010 | 0.004 | -0.003 | 0 |
| YMR193C-A | 0.010 | 0.035 | 0.022 | 0 |
| MEU1 | -0.002 | 0.005 | 0.002 | 0 |
| SCJ1 | -0.094 | -0.105 | -0.099 | -1 |
| MMP1 | -0.015 | 0.010 | -0.003 | 0 |
| PEP5 | -0.014 | -0.023 | -0.019 | 0 |
| RPL8B | 0.019 | 0.014 | 0.017 | 0 |
| YMR244C-A | 0.013 | -0.003 | 0.005 | 0 |
| BDH1 | 0.006 | 0.030 | 0.018 | 0 |
| YLR125W | -0.011 | 0.014 | 0.002 | 0 |
| ERV46 | -0.007 | 0.006 | -0.001 | 0 |
| RPS1B | -0.004 | 0.002 | -0.001 | 0 |
| SAW1 | -0.003 | 0.011 | 0.004 | 0 |
| SUR7 | 0.014 | 0.022 | 0.018 | 0 |
| SWC3 | -0.060 | -0.044 | -0.052 | -1 |
| USA1 | -0.003 | 0.006 | 0.002 | 0 |
| KIN3 | 0.019 | 0.026 | 0.022 | 0 |
| ERV25 | -0.124 | -0.126 | -0.125 | -1 |
| YAR043C | 0.001 | 0.004 | 0.002 | 0 |
| YMR003W | 0.000 | 0.006 | 0.003 | 0 |
| BPT1 | 0.005 | 0.004 | 0.005 | 0 |
| YMR018W | 0.002 | -0.005 | -0.001 | 0 |
| FRA1 | -0.009 | -0.012 | -0.010 | 0 |
| RSF1 | -0.006 | -0.011 | -0.008 | 0 |
| REX3 | 0.004 | 0.031 | 0.017 | 0 |
| IOC4 | -0.004 | -0.004 | -0.004 | 0 |
| YLR091W | 0.026 | -0.097 | -0.036 | 0 |
| YMR153C-A | 0.002 | 0.017 | 0.010 | 0 |
| BUD20 | -0.058 | -0.054 | -0.056 | -1 |
| ALD3 | 0.003 | 0.001 | 0.002 | 0 |
| REX2 | -0.021 | -0.014 | -0.018 | 0 |
| SSO2 | 0.003 | 0.017 | 0.010 | 0 |
| TRX1 | 0.003 | -0.002 | 0.000 | 0 |
| ICY1 | -0.004 | 0.001 | -0.001 | 0 |
| PML1 | -0.005 | 0.000 | -0.003 | 0 |
| GAS3 | -0.001 | 0.007 | 0.003 | 0 |
| GTT2 | -0.003 | 0.003 | 0.000 | 0 |
| FUS2 | -0.011 | 0.000 | -0.005 | 0 |
| FPS1 | -0.014 | -0.081 | -0.047 | 0 |
| FAA4 | 0.003 | -0.009 | -0.003 | 0 |
| ECM1 | -0.007 | 0.020 | 0.006 | 0 |
| YML089C | -0.008 | 0.020 | 0.006 | 0 |
| YAL043C-a | -0.014 | 0.004 | -0.005 | 0 |
| MFT1 | 0.005 | 0.021 | 0.013 | 0 |
| DRS2 | 0.033 | 0.013 | 0.023 | 0 |
| GAL80 | -0.002 | 0.009 | 0.004 | 0 |
| MDM10 | -0.004 | 0.003 | 0.000 | 0 |

| | | | | |
|---|---|---|---|---|
| TSA1 | 0.024 | 0.051 | 0.037 | 1 |
| PAU7 | -0.010 | 0.005 | -0.003 | 0 |
| RAD33 | -0.004 | 0.007 | 0.002 | 0 |
| YAR044W | -0.003 | 0.012 | 0.005 | 0 |
| PLB2 | -0.011 | 0.006 | -0.003 | 0 |
| SDC25 | 0.000 | 0.006 | 0.003 | 0 |
| STB4 | -0.007 | 0.008 | 0.001 | 0 |
| YLL032C | -0.018 | -0.010 | -0.014 | 0 |
| 'MR031W-A | 0.006 | -0.007 | 0.000 | 0 |
| YLR104W | -0.029 | 0.009 | -0.010 | 0 |
| SIP5 | -0.017 | 0.001 | -0.008 | 0 |
| XDJ1 | -0.007 | 0.011 | 0.002 | 0 |
| YMR155W | -0.010 | 0.005 | -0.003 | 0 |
| YLR073C | 0.009 | 0.009 | 0.009 | 0 |
| ALD2 | -0.005 | 0.001 | -0.002 | 0 |
| SHM2 | -0.047 | -0.019 | -0.033 | 0 |
| ADD37 | 0.017 | 0.034 | 0.025 | 0 |
| YLR042C | 0.004 | 0.016 | 0.010 | 0 |
| YMR196W | -0.010 | 0.004 | -0.003 | 0 |
| BRE2 | -0.041 | -0.022 | -0.031 | -1 |
| SKY1 | -0.004 | 0.001 | -0.001 | 0 |
| YLL058W | -0.004 | 0.012 | 0.004 | 0 |
| TRI1 | 0.000 | -0.006 | -0.003 | 0 |
| ATG10 | 0.006 | -0.007 | 0.000 | 0 |
| RKR1 | 0.001 | 0.000 | 0.001 | 0 |
| CNE1 | 0.024 | 0.050 | 0.037 | 1 |
| UFO1 | 0.004 | 0.024 | 0.014 | 0 |
| CLN3 | 0.009 | 0.016 | 0.013 | 0 |
| PIF1 | 0.001 | 0.011 | 0.006 | 0 |
| PMT2 | 0.027 | 0.036 | 0.032 | 1 |
| YML050W | 0.002 | -0.004 | -0.001 | 0 |
| SPO7 | -0.008 | 0.008 | 0.000 | 0 |
| RPS18B | -0.037 | 0.017 | -0.010 | 0 |
| YAR023C | 0.003 | 0.002 | 0.003 | 0 |
| MRPL39 | 0.002 | 0.006 | 0.004 | 0 |
| YAR047C | -0.009 | 0.004 | -0.002 | 0 |
| YMR007W | 0.014 | 0.013 | 0.013 | 0 |
| YLL017W | -0.001 | 0.000 | 0.000 | 0 |
| FMS1 | -0.004 | -0.005 | -0.005 | 0 |
| IRC19 | 0.027 | 0.022 | 0.025 | 1 |
| YMR031C | -0.004 | -0.005 | -0.004 | 0 |
| APC9 | 0.003 | 0.026 | 0.014 | 0 |
| YMR141C | -0.016 | 0.023 | 0.004 | 0 |
| ALT1 | 0.067 | 0.068 | 0.068 | 1 |
| TPP1 | -0.008 | 0.001 | -0.003 | 0 |
| YLR072W | -0.001 | 0.008 | 0.003 | 0 |
| 'MR172C-A | 0.004 | 0.010 | 0.007 | 0 |
| YLR057W | 0.002 | 0.013 | 0.007 | 0 |
| HSC82 | -0.013 | -0.001 | -0.007 | 0 |
| ADE16 | -0.009 | -0.006 | -0.007 | 0 |
| CIK1 | 0.028 | 0.034 | 0.031 | 1 |
| PPR1 | 0.005 | 0.004 | 0.005 | 0 |
| ESC1 | 0.003 | 0.020 | 0.011 | 0 |

| | | | | |
|---|---|---|---|---|
| JLP1 | -0.002 | -0.008 | -0.005 | 0 |
| RNH1 | 0.010 | 0.001 | 0.005 | 0 |
| SDH2 | 0.006 | 0.000 | 0.003 | 0 |
| GAD1 | 0.008 | -0.002 | 0.003 | 0 |
| GPB2 | -0.035 | 0.002 | -0.017 | 0 |
| YML087C | -0.011 | 0.022 | 0.006 | 0 |
| CYC3 | -0.006 | 0.019 | 0.006 | 0 |
| OGG1 | 0.006 | 0.016 | 0.011 | 0 |
| FUN26 | -0.005 | 0.001 | -0.002 | 0 |
| GSF2 | -0.051 | -0.044 | -0.047 | -1 |
| FUN14 | -0.001 | 0.003 | 0.001 | 0 |
| RPS17A | 0.018 | 0.021 | 0.020 | 0 |
| UIP3 | 0.003 | 0.010 | 0.006 | 0 |
| ERG6 | -0.015 | 0.005 | -0.005 | 0 |
| DNM1 | -0.003 | 0.008 | 0.003 | 0 |
| PLB1 | -0.005 | 0.005 | 0.000 | 0 |
| KNS1 | 0.001 | 0.000 | 0.000 | 0 |
| MAC1 | 0.004 | 0.023 | 0.014 | 0 |
| ENT4 | 0.022 | 0.019 | 0.020 | 0 |
| HOF1 | 0.008 | -0.010 | -0.001 | 0 |
| RPS16A | 0.011 | 0.012 | 0.012 | 0 |
| CSF1 | 0.025 | 0.039 | 0.032 | 1 |
| YMR157C | 0.012 | 0.023 | 0.017 | 0 |
| XYL2 | -0.004 | 0.015 | 0.005 | 0 |
| YMR173W-A | 0.001 | 0.012 | 0.006 | 0 |
| ERG3 | 0.005 | 0.000 | 0.003 | 0 |
| YMR187C | -0.003 | 0.013 | 0.005 | 0 |
| AAT2 | 0.019 | -0.056 | -0.018 | 0 |
| CLN1 | -0.005 | 0.002 | -0.001 | 0 |
| GAT3 | -0.001 | 0.004 | 0.001 | 0 |
| YMR221C | -0.166 | -0.158 | -0.162 | -1 |
| YLL056C | 0.007 | -0.002 | 0.002 | 0 |
| BCH1 | 0.015 | 0.014 | 0.015 | 0 |
| GTO3 | 0.000 | -0.006 | -0.003 | 0 |
| PEX22 | -0.007 | -0.001 | -0.004 | 0 |
| ALO1 | 0.003 | 0.019 | 0.011 | 0 |
| YAL037W | -0.011 | -0.003 | -0.007 | 0 |
| NTE1 | 0.004 | 0.016 | 0.010 | 0 |
| CCR4 | -0.019 | -0.006 | -0.013 | 0 |
| YML048W-A | 0.007 | 0.009 | 0.008 | 0 |
| ERP2 | -0.038 | -0.023 | -0.030 | -1 |
| YML020W | -0.006 | 0.004 | -0.001 | 0 |
| YAR028W | -0.002 | 0.007 | 0.003 | 0 |
| YAP1 | 0.001 | 0.007 | 0.004 | 0 |
| RTT109 | 0.059 | 0.063 | 0.061 | 1 |
| ADI1 | -0.002 | 0.010 | 0.004 | 0 |
| YLL020C | -0.012 | 0.004 | -0.004 | 0 |
| UBC7 | 0.058 | 0.065 | 0.062 | 1 |
| UBI4 | 0.001 | -0.007 | -0.003 | 0 |
| YMR034C | -0.006 | -0.010 | -0.008 | 0 |
| ICT1 | 0.008 | 0.052 | 0.030 | 0 |
| YMR144W | 0.008 | 0.019 | 0.014 | 0 |
| ARP6 | -0.069 | -0.042 | -0.056 | -1 |

| | | | | |
|---|---|---|---|---|
| (MR158W-A | -0.003 | 0.002 | -0.001 | 0 |
| MEF1 | -0.019 | -0.073 | -0.046 | 0 |
| PAI3 | -0.005 | 0.001 | -0.002 | 0 |
| SPT8 | 0.008 | 0.033 | 0.020 | 0 |
| MRPS17 | 0.010 | 0.010 | 0.010 | 0 |
| SNF7 | -0.013 | -0.188 | -0.101 | 0 |
| RAD14 | -0.007 | 0.015 | 0.004 | 0 |
| YLR012C | 0.002 | -0.007 | -0.003 | 0 |
| FSH2 | -0.013 | 0.004 | -0.004 | 0 |
| YCT1 | -0.018 | -0.002 | -0.010 | 0 |
| DFG5 | 0.023 | 0.013 | 0.018 | 0 |
| VPS13 | -0.028 | -0.023 | -0.025 | -1 |
| HOR7 | 0.002 | -0.005 | -0.001 | 0 |
| YMR252C | -0.002 | 0.017 | 0.007 | 0 |
| YOR088W | 0.002 | 0.007 | 0.005 | 0 |
| CUE1 | 0.039 | 0.048 | 0.043 | 1 |
| RPS10A | 0.054 | 0.058 | 0.056 | 1 |
| AEP2 | -0.008 | -0.006 | -0.007 | 0 |
| SLY41 | -0.010 | 0.006 | -0.002 | 0 |
| YMR295C | 0.002 | -0.001 | 0.001 | 0 |
| PRO2 | 0.000 | 0.002 | 0.001 | 0 |
| YRF1-6 | 0.011 | -0.006 | 0.002 | 0 |
| TYE7 | 0.004 | 0.007 | 0.006 | 0 |
| YNL324W | -0.002 | 0.000 | -0.001 | 0 |
| HAP5 | 0.005 | 0.002 | 0.004 | 0 |
| YNL305C | -0.009 | -0.004 | -0.007 | 0 |
| YOR378W | -0.009 | 0.003 | -0.003 | 0 |
| YNL089C | -0.010 | -0.017 | -0.014 | 0 |
| TOP1 | 0.007 | 0.002 | 0.005 | 0 |
| YNL276C | -0.012 | 0.008 | -0.002 | 0 |
| YOL019W | 0.001 | -0.003 | -0.001 | 0 |
| SIP3 | -0.017 | -0.012 | -0.014 | 0 |
| SGT2 | -0.001 | -0.006 | -0.003 | 0 |
| YOL048C | -0.005 | -0.001 | -0.003 | 0 |
| YOR019W | -0.010 | -0.001 | -0.005 | 0 |
| GPD2 | -0.007 | -0.015 | -0.011 | 0 |
| HMS1 | 0.010 | -0.002 | 0.004 | 0 |
| YOL075C | -0.008 | -0.021 | -0.015 | 0 |
| IRC23 | -0.004 | -0.007 | -0.006 | 0 |
| ATP15 | -0.033 | 0.081 | 0.024 | 0 |
| CKA2 | 0.022 | 0.024 | 0.023 | 1 |
| YPL257W | -0.003 | -0.006 | -0.004 | 0 |
| SKI7 | 0.004 | 0.004 | 0.004 | 0 |
| HSP82 | -0.014 | -0.001 | -0.008 | 0 |
| YMR253C | -0.005 | 0.011 | 0.003 | 0 |
| VPS21 | -0.018 | -0.020 | -0.019 | 0 |
| YMR265C | -0.002 | 0.007 | 0.003 | 0 |
| UAF30 | -0.012 | 0.049 | 0.018 | 0 |
| RIT1 | 0.012 | 0.012 | 0.012 | 0 |
| SNU66 | 0.007 | 0.001 | 0.004 | 0 |
| PRC1 | -0.019 | -0.011 | -0.015 | 0 |
| FRT1 | -0.018 | -0.003 | -0.011 | 0 |
| YNL338W | 0.012 | 0.004 | 0.008 | 0 |

| | | | | |
|---|---|---|---|---|
| REV1 | -0.008 | -0.005 | -0.006 | 0 |
| FIG4 | -0.002 | -0.001 | -0.002 | 0 |
| VTS1 | -0.030 | -0.020 | -0.025 | -1 |
| YPT11 | -0.014 | -0.004 | -0.009 | 0 |
| RDR1 | -0.009 | 0.003 | -0.003 | 0 |
| BOR1 | -0.007 | 0.018 | 0.006 | 0 |
| TAT2 | -0.007 | 0.005 | -0.001 | 0 |
| GIS2 | -0.006 | -0.004 | -0.005 | 0 |
| YOL035C | 0.012 | 0.012 | 0.012 | 0 |
| SLG1 | 0.013 | 0.009 | 0.011 | 0 |
| GSH2 | -0.018 | 0.063 | 0.022 | 0 |
| YOR021C | 0.004 | 0.007 | 0.005 | 0 |
| MAM3 | -0.005 | 0.001 | -0.002 | 0 |
| EXO1 | 0.007 | 0.002 | 0.004 | 0 |
| MDM20 | -0.006 | 0.006 | 0.000 | 0 |
| TOM6 | 0.011 | 0.011 | 0.011 | 0 |
| MDL2 | 0.023 | 0.016 | 0.019 | 0 |
| YOR062C | 0.004 | 0.004 | 0.004 | 0 |
| CLN2 | 0.013 | 0.009 | 0.011 | 0 |
| FAA2 | 0.006 | 0.001 | 0.004 | 0 |
| YMR254C | -0.014 | -0.004 | -0.009 | 0 |
| PTC5 | -0.002 | 0.011 | 0.004 | 0 |
| RSN1 | -0.009 | 0.003 | -0.003 | 0 |
| YOR296W | 0.004 | 0.004 | 0.004 | 0 |
| YKU70 | 0.010 | 0.011 | 0.011 | 0 |
| DGK1 | 0.007 | 0.011 | 0.009 | 0 |
| DYN3 | -0.012 | -0.002 | -0.007 | 0 |
| SNC2 | -0.005 | 0.000 | -0.003 | 0 |
| COS1 | 0.007 | -0.002 | 0.003 | 0 |
| PYK2 | -0.006 | -0.001 | -0.003 | 0 |
| LEM3 | -0.057 | -0.030 | -0.043 | -1 |
| PDE2 | -0.023 | -0.012 | -0.018 | 0 |
| YNL303W | -0.001 | 0.001 | 0.000 | 0 |
| FRE3 | -0.011 | -0.001 | -0.006 | 0 |
| MID1 | -0.010 | -0.010 | -0.010 | 0 |
| CSI2 | 0.010 | -0.003 | 0.003 | 0 |
| TOF1 | -0.008 | 0.016 | 0.004 | 0 |
| IFM1 | 0.044 | 0.040 | 0.042 | 1 |
| RTC4 | 0.000 | 0.004 | 0.002 | 0 |
| YOL036W | 0.003 | 0.006 | 0.005 | 0 |
| TIR4 | 0.006 | 0.003 | 0.004 | 0 |
| YOL050C | -0.116 | -0.118 | -0.117 | -1 |
| YOR022C | 0.007 | 0.019 | 0.013 | 0 |
| PRS5 | 0.023 | 0.019 | 0.021 | 0 |
| AKR2 | 0.009 | -0.005 | 0.002 | 0 |
| YOL079W | 0.001 | -0.005 | -0.002 | 0 |
| STD1 | -0.004 | 0.001 | -0.002 | 0 |
| KAR9 | 0.002 | 0.007 | 0.004 | 0 |
| YNG1 | 0.001 | -0.002 | -0.001 | 0 |
| HFI1 | -0.030 | -0.006 | -0.018 | 0 |
| BUD21 | -0.053 | -0.045 | -0.049 | -1 |
| YAR1 | 0.016 | 0.013 | 0.014 | 0 |
| GFD1 | -0.002 | 0.014 | 0.006 | 0 |

| | | | | |
|---|---|---|---|---|
| TMA46 | -0.006 | 0.011 | 0.003 | 0 |
| PPA2 | -0.008 | 0.040 | 0.016 | 0 |
| TIM18 | 0.016 | 0.018 | 0.017 | 0 |
| NGL2 | -0.005 | 0.001 | -0.002 | 0 |
| RPL20B | 0.046 | 0.070 | 0.058 | 1 |
| ADE4 | 0.001 | 0.004 | 0.002 | 0 |
| PDR10 | -0.003 | -0.001 | -0.002 | 0 |
| DDI3 | 0.003 | 0.001 | 0.002 | 0 |
| PUT4 | 0.001 | -0.001 | 0.000 | 0 |
| KRE1 | 0.024 | 0.019 | 0.021 | 0 |
| PIP2 | 0.001 | 0.003 | 0.002 | 0 |
| RPS19B | 0.001 | 0.009 | 0.005 | 0 |
| FIT2 | -0.004 | -0.001 | -0.002 | 0 |
| PCL1 | -0.011 | -0.026 | -0.019 | 0 |
| COQ10 | 0.021 | 0.018 | 0.019 | 0 |
| BNI1 | 0.014 | 0.017 | 0.016 | 0 |
| YOL024W | -0.008 | 0.002 | -0.003 | 0 |
| TEX1 | 0.006 | 0.010 | 0.008 | 0 |
| YOL037C | 0.001 | 0.005 | 0.003 | 0 |
| TIR2 | -0.007 | -0.003 | -0.005 | 0 |
| GAL11 | -0.005 | 0.004 | 0.000 | 0 |
| AHC1 | -0.016 | -0.008 | -0.012 | 0 |
| APM4 | -0.004 | -0.001 | -0.002 | 0 |
| SHE4 | 0.017 | -0.027 | -0.005 | 0 |
| REX4 | 0.000 | -0.008 | -0.004 | 0 |
| RSB1 | -0.006 | 0.003 | -0.001 | 0 |
| ACM1 | 0.005 | -0.001 | 0.002 | 0 |
| VIK1 | -0.006 | 0.004 | -0.001 | 0 |
| ATX2 | -0.002 | -0.015 | -0.008 | 0 |
| YPL236C | -0.018 | -0.015 | -0.017 | 0 |
| GEA2 | -0.004 | 0.030 | 0.013 | 0 |
| ECM3 | -0.011 | -0.004 | -0.008 | 0 |
| TMA23 | -0.001 | -0.003 | -0.002 | 0 |
| MUM3 | -0.008 | -0.002 | -0.005 | 0 |
| MRPL33 | 0.029 | 0.057 | 0.043 | 1 |
| SPS4 | -0.011 | -0.001 | -0.006 | 0 |
| YME2 | 0.000 | -0.007 | -0.003 | 0 |
| MIP1 | 0.000 | -0.003 | -0.001 | 0 |
| SNO2 | 0.008 | -0.002 | 0.003 | 0 |
| CIN1 | -0.008 | 0.003 | -0.003 | 0 |
| VNX1 | -0.001 | 0.001 | 0.000 | 0 |
| YOR365C | 0.012 | 0.000 | 0.006 | 0 |
| RPL18B | -0.006 | -0.002 | -0.004 | 0 |
| FIT3 | 0.000 | -0.004 | -0.002 | 0 |
| CAF40 | 0.039 | 0.034 | 0.037 | 1 |
| MDM12 | 0.020 | 0.025 | 0.022 | 0 |
| ALP1 | 0.006 | 0.003 | 0.004 | 0 |
| LAG2 | -0.002 | 0.009 | 0.003 | 0 |
| MPA43 | 0.010 | 0.007 | 0.009 | 0 |
| RPP2A | 0.001 | 0.013 | 0.007 | 0 |
| AUS1 | -0.002 | 0.002 | 0.000 | 0 |
| SPE2 | 0.037 | 0.030 | 0.033 | 1 |
| YOR024W | -0.011 | -0.008 | -0.009 | 0 |

| | | | | |
|---|---|---|---|---|
| CRT10 | -0.012 | -0.004 | -0.008 | 0 |
| PEP12 | -0.002 | -0.022 | -0.012 | 0 |
| IRA2 | -0.003 | -0.019 | -0.011 | 0 |
| YOR050C | -0.004 | 0.004 | 0.000 | 0 |
| DIP5 | -0.020 | -0.011 | -0.015 | 0 |
| MSA1 | -0.018 | -0.010 | -0.014 | 0 |
| ICY2 | -0.005 | -0.009 | -0.007 | 0 |
| DIA2 | 0.043 | 0.046 | 0.044 | 1 |
| TFP3 | 0.010 | 0.078 | 0.044 | 0 |
| COX7 | -0.018 | -0.008 | -0.013 | 0 |
| YOR093C | -0.008 | 0.000 | -0.004 | 0 |
| SCS7 | -0.006 | -0.012 | -0.009 | 0 |
| BUD7 | -0.001 | -0.001 | -0.001 | 0 |
| DSS1 | -0.013 | 0.057 | 0.022 | 0 |
| YOR314W | -0.011 | -0.001 | -0.006 | 0 |
| ADH2 | -0.003 | 0.009 | 0.003 | 0 |
| VMA4 | -0.014 | -0.019 | -0.016 | 0 |
| SNZ2 | 0.011 | 0.003 | 0.007 | 0 |
| MNE1 | 0.021 | 0.015 | 0.018 | 0 |
| YNL320W | -0.006 | 0.009 | 0.002 | 0 |
| SCP1 | 0.002 | 0.003 | 0.003 | 0 |
| TRF5 | -0.006 | 0.001 | -0.003 | 0 |
| FRE5 | -0.007 | 0.002 | -0.003 | 0 |
| CUS2 | 0.008 | 0.000 | 0.004 | 0 |
| PLB3 | -0.001 | -0.001 | -0.001 | 0 |
| MDM38 | 0.019 | 0.015 | 0.017 | 0 |
| RPA49 | 0.030 | 0.044 | 0.037 | 1 |
| NOP12 | 0.015 | 0.041 | 0.028 | 0 |
| YOR012W | -0.004 | 0.002 | -0.001 | 0 |
| YOL053C-A | 0.002 | 0.000 | 0.001 | 0 |
| HST3 | -0.009 | -0.007 | -0.008 | 0 |
| MET22 | 0.056 | 0.061 | 0.058 | 1 |
| CYC2 | 0.013 | 0.012 | 0.012 | 0 |
| ATG19 | 0.010 | 0.000 | 0.005 | 0 |
| YOR051C | 0.003 | 0.008 | 0.006 | 0 |
| YPL264C | 0.002 | 0.003 | 0.003 | 0 |
| ALG8 | 0.007 | 0.002 | 0.004 | 0 |
| GYP5 | -0.001 | 0.006 | 0.002 | 0 |
| TGL5 | 0.007 | 0.000 | 0.004 | 0 |
| SSO1 | -0.024 | -0.013 | -0.018 | 0 |
| PET111 | 0.052 | 0.053 | 0.052 | 1 |
| ZDS1 | 0.012 | 0.019 | 0.016 | 0 |
| RAX1 | -0.013 | -0.006 | -0.009 | 0 |
| ABZ2 | -0.012 | -0.002 | -0.007 | 0 |
| SFG1 | -0.003 | 0.003 | 0.000 | 0 |
| UBP15 | -0.099 | -0.067 | -0.083 | -1 |
| MRS2 | -0.007 | 0.002 | -0.002 | 0 |
| THI12 | 0.010 | 0.000 | 0.005 | 0 |
| MEK1 | 0.011 | 0.003 | 0.007 | 0 |
| YNL319W | -0.007 | 0.006 | -0.001 | 0 |
| RAD17 | 0.008 | 0.003 | 0.006 | 0 |
| CLA4 | -0.084 | -0.100 | -0.092 | -1 |
| YOR385W | -0.012 | 0.002 | -0.005 | 0 |

| | | | | |
|---|---|---|---|---|
| YNL285W | 0.012 | 0.009 | 0.011 | 0 |
| HTZ1 | -0.065 | -0.062 | -0.063 | -1 |
| BSC4 | -0.010 | 0.000 | -0.005 | 0 |
| YAP7 | -0.003 | 0.003 | 0.000 | 0 |
| VPS75 | 0.011 | 0.021 | 0.016 | 0 |
| NGL1 | 0.006 | 0.006 | 0.006 | 0 |
| IRC11 | -0.013 | 0.001 | -0.006 | 0 |
| YOL053W | -0.002 | 0.008 | 0.003 | 0 |
| BUB3 | 0.016 | 0.009 | 0.013 | 0 |
| INP54 | -0.001 | -0.006 | -0.003 | 0 |
| HIR2 | -0.003 | 0.001 | -0.001 | 0 |
| YOL083W | 0.010 | 0.000 | 0.005 | 0 |
| YOR052C | -0.013 | -0.006 | -0.010 | 0 |
| KEL3 | -0.002 | 0.002 | 0.000 | 0 |
| VAM10 | -0.021 | -0.008 | -0.014 | 0 |
| GAL4 | 0.001 | 0.006 | 0.004 | 0 |
| YOR082C | 0.005 | -0.001 | 0.002 | 0 |
| USV1 | 0.000 | -0.009 | -0.005 | 0 |
| YMR258C | -0.001 | 0.009 | 0.004 | 0 |
| ARF3 | 0.002 | 0.002 | 0.002 | 0 |
| RCE1 | -0.053 | -0.058 | -0.056 | -1 |
| YOR302W | 0.001 | 0.023 | 0.012 | 0 |
| YMR291W | -0.007 | 0.004 | -0.001 | 0 |
| COT1 | 0.006 | 0.001 | 0.004 | 0 |
| YMR304C-A | 0.004 | 0.005 | 0.004 | 0 |
| TEA1 | -0.002 | 0.008 | 0.003 | 0 |
| RPD3 | -0.005 | -0.005 | -0.005 | 0 |
| YOR352W | 0.010 | -0.001 | 0.004 | 0 |
| HXT14 | -0.009 | 0.000 | -0.005 | 0 |
| GPB1 | 0.002 | 0.001 | 0.001 | 0 |
| YNL086W | -0.021 | 0.000 | -0.010 | 0 |
| PHR1 | 0.007 | 0.001 | 0.004 | 0 |
| WSC2 | 0.010 | 0.007 | 0.008 | 0 |
| HRD1 | -0.009 | 0.002 | -0.003 | 0 |
| LYP1 | -0.014 | -0.003 | -0.009 | 0 |
| RRP6 | -0.013 | 0.003 | -0.005 | 0 |
| NTG2 | -0.006 | 0.005 | 0.000 | 0 |
| RTS1 | 0.022 | 0.028 | 0.025 | 1 |
| PSH1 | 0.010 | 0.012 | 0.011 | 0 |
| STI1 | -0.038 | -0.025 | -0.031 | -1 |
| RTG1 | -0.025 | 0.075 | 0.025 | 0 |
| CKB2 | -0.008 | -0.009 | -0.008 | 0 |
| PHM7 | 0.012 | 0.009 | 0.010 | 0 |
| YOR053W | -0.007 | 0.006 | -0.001 | 0 |
| FUM1 | 0.022 | 0.012 | 0.017 | 0 |
| VPS5 | -0.019 | -0.016 | -0.018 | 0 |
| YPL247C | -0.006 | 0.003 | -0.001 | 0 |
| WHI5 | -0.016 | -0.008 | -0.012 | 0 |
| YPL229W | 0.001 | -0.011 | -0.005 | 0 |
| YMR259C | -0.009 | 0.008 | -0.001 | 0 |
| YOR289W | 0.004 | 0.010 | 0.007 | 0 |
| BUL1 | -0.035 | -0.026 | -0.031 | -1 |
| CPA1 | -0.018 | -0.035 | -0.027 | 0 |

| | | | | |
|---|---|---|---|---|
| GOT1 | -0.018 | -0.024 | -0.021 | 0 |
| YOR318C | 0.003 | 0.002 | 0.003 | 0 |
| SCW10 | 0.013 | 0.009 | 0.011 | 0 |
| YOR338W | -0.005 | -0.002 | -0.003 | 0 |
| PEX6 | -0.004 | -0.010 | -0.007 | 0 |
| MSC6 | 0.011 | 0.006 | 0.008 | 0 |
| DAL82 | -0.011 | 0.006 | -0.002 | 0 |
| ALD4 | 0.009 | -0.002 | 0.004 | 0 |
| MON2 | -0.009 | 0.008 | 0.000 | 0 |
| PHO80 | 0.031 | 0.046 | 0.038 | 1 |
| HCH1 | 0.012 | 0.003 | 0.007 | 0 |
| YOL014W | 0.005 | 0.004 | 0.004 | 0 |
| YNL266W | -0.012 | 0.006 | -0.003 | 0 |
| YOL029C | -0.002 | 0.010 | 0.004 | 0 |
| ALG6 | 0.005 | 0.028 | 0.017 | 0 |
| PEX15 | 0.011 | 0.002 | 0.006 | 0 |
| YOR015W | 0.003 | 0.006 | 0.005 | 0 |
| THI20 | 0.005 | 0.002 | 0.003 | 0 |
| CIN5 | 0.003 | -0.001 | 0.001 | 0 |
| HST1 | -0.027 | -0.016 | -0.021 | 0 |
| GLO4 | 0.000 | 0.000 | 0.000 | 0 |
| YOL085C | 0.005 | 0.003 | 0.004 | 0 |
| VHS3 | -0.010 | 0.004 | -0.003 | 0 |
| YPL260W | 0.003 | -0.005 | -0.001 | 0 |
| GYP1 | 0.009 | 0.018 | 0.013 | 0 |
| RBD2 | 0.018 | 0.010 | 0.014 | 0 |
| LPX1 | 0.001 | 0.006 | 0.004 | 0 |
| ALG5 | -0.010 | -0.001 | -0.006 | 0 |
| TPS3 | 0.010 | 0.029 | 0.020 | 0 |
| SNF2 | 0.006 | 0.021 | 0.014 | 0 |
| DSK2 | 0.012 | 0.025 | 0.018 | 0 |
| YOR304C-A | -0.004 | 0.004 | 0.000 | 0 |
| HER2 | -0.021 | 0.063 | 0.021 | 0 |
| GNT1 | 0.008 | 0.008 | 0.008 | 0 |
| YMR306C-A | 0.000 | -0.003 | -0.001 | 0 |
| UBC11 | 0.004 | -0.002 | 0.001 | 0 |
| MDJ2 | -0.008 | -0.006 | -0.007 | 0 |
| GDS1 | 0.011 | 0.012 | 0.011 | 0 |
| YNL311C | -0.013 | -0.005 | -0.009 | 0 |
| GDH1 | 0.007 | -0.004 | 0.001 | 0 |
| YNL295W | 0.001 | 0.003 | 0.002 | 0 |
| IZH2 | 0.000 | 0.000 | 0.000 | 0 |
| ERG24 | -0.121 | -0.058 | -0.090 | -1 |
| IRC10 | -0.001 | -0.002 | -0.002 | 0 |
| IST1 | -0.005 | 0.007 | 0.001 | 0 |
| GAS5 | 0.005 | 0.008 | 0.006 | 0 |
| YSP3 | -0.011 | 0.010 | 0.000 | 0 |
| PSK2 | -0.032 | -0.019 | -0.026 | 0 |
| ERP4 | -0.013 | -0.012 | -0.013 | 0 |
| GPM3 | 0.004 | 0.011 | 0.008 | 0 |
| YOR029W | 0.003 | 0.005 | 0.004 | 0 |
| NBA1 | 0.006 | -0.001 | 0.002 | 0 |
| YOR041C | -0.002 | -0.003 | -0.002 | 0 |

| | | | | |
|---|---|---|---|---|
| SAM3 | 0.002 | -0.004 | -0.001 | 0 |
| YOR055W | -0.010 | 0.002 | -0.004 | 0 |
| YPL261C | 0.026 | 0.012 | 0.019 | 0 |
| NRT1 | -0.001 | 0.011 | 0.005 | 0 |
| YPL245W | -0.006 | -0.001 | -0.003 | 0 |
| OST3 | -0.012 | -0.010 | -0.011 | 0 |
| NEW1 | -0.008 | 0.000 | -0.004 | 0 |
| YMR262W | -0.004 | 0.017 | 0.006 | 0 |
| YOR291W | 0.000 | 0.008 | 0.004 | 0 |
| PGM3 | -0.003 | 0.002 | -0.001 | 0 |
| ISW2 | 0.001 | 0.006 | 0.004 | 0 |
| JNM1 | -0.008 | -0.002 | -0.005 | 0 |
| PMT3 | 0.014 | -0.004 | 0.005 | 0 |
| GAS1 | 0.040 | 0.039 | 0.039 | 1 |
| YOR342C | 0.012 | 0.000 | 0.006 | 0 |
| EGT2 | -0.009 | -0.001 | -0.005 | 0 |
| YOR356W | 0.008 | 0.003 | 0.005 | 0 |
| STB1 | 0.001 | 0.008 | 0.004 | 0 |
| YOR376W | 0.007 | 0.000 | 0.004 | 0 |
| RIM21 | -0.018 | -0.035 | -0.027 | 0 |
| PFA4 | 0.044 | 0.042 | 0.043 | 1 |
| CAF120 | 0.010 | 0.009 | 0.009 | 0 |
| ESC8 | -0.006 | -0.001 | -0.004 | 0 |
| PDR17 | -0.009 | 0.040 | 0.016 | 0 |
| SIL1 | -0.090 | -0.089 | -0.089 | -1 |
| DNL4 | -0.005 | -0.015 | -0.010 | 0 |
| YOL046C | -0.040 | -0.023 | -0.032 | -1 |
| PET127 | 0.003 | 0.008 | 0.005 | 0 |
| YOL057W | -0.004 | -0.012 | -0.008 | 0 |
| DFG16 | -0.042 | -0.041 | -0.041 | -1 |
| EMI5 | -0.003 | 0.003 | 0.000 | 0 |
| CUE5 | -0.005 | -0.004 | -0.005 | 0 |
| SAM4 | 0.000 | -0.005 | -0.003 | 0 |
| ASE1 | -0.003 | -0.002 | -0.003 | 0 |
| APM1 | 0.016 | 0.009 | 0.013 | 0 |
| YOR072W | -0.005 | -0.003 | -0.004 | 0 |
| HUT1 | -0.010 | -0.003 | -0.006 | 0 |
| TCB1 | 0.006 | -0.008 | -0.001 | 0 |
| YPL225W | 0.017 | 0.010 | 0.013 | 0 |
| SLM2 | -0.003 | 0.012 | 0.005 | 0 |
| YOR292C | -0.015 | 0.007 | -0.004 | 0 |
| CAT8 | -0.008 | -0.003 | -0.005 | 0 |
| YOR305W | 0.029 | 0.053 | 0.041 | 1 |
| YMR294W-A | -0.003 | -0.001 | -0.002 | 0 |
| LDB19 | 0.011 | 0.008 | 0.010 | 0 |
| YMR310C | -0.011 | -0.012 | -0.011 | 0 |
| YOR343C | 0.011 | 0.011 | 0.011 | 0 |
| PFA3 | 0.008 | 0.023 | 0.016 | 0 |
| SNX3 | -0.018 | -0.017 | -0.017 | 0 |
| MSG5 | 0.009 | 0.007 | 0.008 | 0 |
| ATF1 | 0.001 | 0.005 | 0.003 | 0 |
| MSB3 | -0.016 | 0.000 | -0.008 | 0 |
| SIN3 | 0.047 | 0.039 | 0.043 | 1 |

| | | | | |
|---|---|---|---|---|
| RPS7B | 0.008 | 0.014 | 0.011 | 0 |
| TLG2 | -0.008 | -0.007 | -0.008 | 0 |
| ATX1 | -0.002 | 0.009 | 0.004 | 0 |
| OPI10 | -0.017 | 0.001 | -0.008 | 0 |
| YOR006C | 0.001 | 0.019 | 0.010 | 0 |
| YOL047C | 0.004 | 0.005 | 0.005 | 0 |
| ROD1 | -0.001 | 0.005 | 0.002 | 0 |
| ARG1 | 0.009 | 0.065 | 0.037 | 0 |
| CRS5 | 0.004 | 0.006 | 0.005 | 0 |
| THP1 | 0.033 | 0.060 | 0.047 | 1 |
| WHI2 | -0.009 | -0.037 | -0.023 | 0 |
| YPL272C | -0.001 | -0.001 | -0.001 | 0 |
| YOR059C | -0.004 | -0.003 | -0.004 | 0 |
| THI21 | -0.003 | 0.004 | 0.000 | 0 |
| SGO1 | -0.006 | -0.001 | -0.003 | 0 |
| CIN2 | -0.002 | 0.015 | 0.007 | 0 |
| YVC1 | 0.001 | 0.000 | 0.000 | 0 |
| GRE1 | 0.006 | 0.001 | 0.004 | 0 |
| FMP40 | -0.011 | -0.039 | -0.025 | 0 |
| DOS2 | -0.004 | -0.030 | -0.017 | 0 |
| PGC1 | -0.004 | -0.001 | -0.003 | 0 |
| RRP8 | 0.015 | 0.014 | 0.015 | 0 |
| RSA1 | 0.013 | 0.014 | 0.014 | 0 |
| BMH2 | -0.002 | 0.010 | 0.004 | 0 |
| PPQ1 | -0.007 | -0.002 | -0.005 | 0 |
| IRC2 | 0.009 | 0.008 | 0.009 | 0 |
| SET6 | -0.001 | -0.007 | -0.004 | 0 |
| SWF1 | -0.020 | -0.050 | -0.035 | 0 |
| ATG5 | -0.002 | 0.009 | 0.003 | 0 |
| AGC1 | 0.004 | 0.007 | 0.005 | 0 |
| TAF14 | -0.055 | -0.027 | -0.041 | -1 |
| MRP1 | -0.036 | -0.035 | -0.035 | -1 |
| PEX25 | 0.010 | 0.000 | 0.005 | 0 |
| CDC40 | 0.001 | 0.003 | 0.002 | 0 |
| YPL102C | 0.046 | 0.063 | 0.055 | 1 |
| ARO10 | -0.009 | -0.004 | -0.007 | 0 |
| EHT1 | -0.009 | -0.004 | -0.006 | 0 |
| SHE9 | 0.015 | 0.009 | 0.012 | 0 |
| YBR194W | 0.001 | -0.004 | -0.002 | 0 |
| DFM1 | 0.046 | 0.056 | 0.051 | 1 |
| YBR209W | -0.010 | -0.019 | -0.014 | 0 |
| CYM1 | -0.003 | -0.006 | -0.004 | 0 |
| PCS60 | 0.010 | 0.003 | 0.006 | 0 |
| YEL008W | 0.002 | 0.007 | 0.004 | 0 |
| YBR238C | 0.052 | 0.042 | 0.047 | 1 |
| RIP1 | 0.004 | 0.005 | 0.005 | 0 |
| MTC4 | -0.025 | -0.049 | -0.037 | -1 |
| YEF1 | 0.002 | 0.006 | 0.004 | 0 |
| TPI1 | 0.006 | 0.006 | 0.006 | 0 |
| RPL12A | 0.043 | 0.027 | 0.035 | 1 |
| FLC1 | 0.000 | 0.001 | 0.000 | 0 |
| DOA4 | -0.025 | -0.021 | -0.023 | -1 |
| YPL205C | 0.017 | 0.029 | 0.023 | 0 |

| | | | | |
|---|---|---|---|---|
| TVP23 | -0.001 | 0.001 | 0.000 | 0 |
| PRM3 | -0.004 | -0.001 | -0.002 | 0 |
| TVP15 | -0.010 | -0.001 | -0.005 | 0 |
| CBC2 | 0.017 | 0.012 | 0.015 | 0 |
| YDR114C | 0.013 | 0.011 | 0.012 | 0 |
| MLH3 | 0.000 | -0.003 | -0.002 | 0 |
| ARO1 | 0.009 | -0.016 | -0.004 | 0 |
| PXA1 | 0.003 | -0.019 | -0.008 | 0 |
| RUB1 | 0.004 | 0.024 | 0.014 | 0 |
| HHO1 | 0.003 | 0.006 | 0.005 | 0 |
| YDR348C | 0.023 | 0.024 | 0.024 | 1 |
| ATG21 | 0.004 | 0.012 | 0.008 | 0 |
| RPP2B | 0.050 | 0.045 | 0.048 | 1 |
| YBR178W | -0.009 | -0.003 | -0.006 | 0 |
| SXM1 | -0.017 | -0.016 | -0.017 | 0 |
| MSI1 | -0.009 | 0.000 | -0.005 | 0 |
| ERD1 | -0.060 | -0.037 | -0.048 | -1 |
| ERV15 | -0.004 | 0.005 | 0.000 | 0 |
| YDR431W | -0.009 | -0.029 | -0.019 | 0 |
| TDP1 | -0.006 | 0.000 | -0.003 | 0 |
| GCN4 | -0.078 | -0.068 | -0.073 | -1 |
| YBR239C | -0.014 | 0.003 | -0.006 | 0 |
| YEL025C | 0.010 | 0.010 | 0.010 | 0 |
| SHG1 | -0.018 | -0.006 | -0.012 | 0 |
| GDA1 | 0.012 | 0.017 | 0.014 | 0 |
| YER076C | -0.005 | -0.003 | -0.004 | 0 |
| RPL1A | -0.008 | 0.000 | -0.004 | 0 |
| FMP16 | -0.005 | 0.000 | -0.003 | 0 |
| TPK2 | -0.010 | -0.004 | -0.007 | 0 |
| AFR1 | -0.003 | 0.007 | 0.002 | 0 |
| YPL191C | -0.006 | -0.002 | -0.004 | 0 |
| ARX1 | 0.033 | 0.033 | 0.033 | 1 |
| CUP9 | -0.016 | -0.025 | -0.020 | 0 |
| YDR115W | 0.003 | 0.004 | 0.003 | 0 |
| SVS1 | -0.009 | 0.001 | -0.004 | 0 |
| MTC5 | -0.014 | -0.028 | -0.021 | 0 |
| KES1 | -0.005 | -0.024 | -0.015 | 0 |
| MTQ2 | -0.008 | 0.000 | -0.004 | 0 |
| KAP120 | 0.002 | 0.001 | 0.001 | 0 |
| YPS7 | 0.009 | -0.032 | -0.011 | 0 |
| CAR1 | -0.002 | 0.011 | 0.005 | 0 |
| YPR1 | 0.010 | 0.010 | 0.010 | 0 |
| YPL099C | -0.006 | 0.000 | -0.003 | 0 |
| NKP1 | 0.001 | 0.001 | 0.001 | 0 |
| FZO1 | 0.055 | 0.050 | 0.052 | 1 |
| HPT1 | 0.069 | 0.074 | 0.072 | 1 |
| YBR197C | -0.017 | -0.016 | -0.016 | 0 |
| YDR415C | -0.005 | 0.001 | -0.002 | 0 |
| NGR1 | 0.011 | 0.012 | 0.011 | 0 |
| NPL3 | 0.037 | 0.029 | 0.033 | 1 |
| YBR224W | 0.006 | 0.009 | 0.007 | 0 |
| YEL010W | 0.000 | -0.001 | -0.001 | 0 |
| THI2 | -0.003 | 0.008 | 0.002 | 0 |

| | | | | |
|---|---|---|---|---|
| CUP5 | 0.000 | 0.000 | 0.000 | 0 |
| YBR259W | -0.001 | 0.004 | 0.002 | 0 |
| YEL043W | 0.041 | 0.033 | 0.037 | 1 |
| DET1 | 0.009 | 0.010 | 0.010 | 0 |
| HAT2 | 0.011 | 0.012 | 0.011 | 0 |
| PCL8 | -0.011 | -0.004 | -0.007 | 0 |
| IPT1 | -0.010 | -0.010 | -0.010 | 0 |
| AFT2 | -0.003 | 0.004 | 0.001 | 0 |
| YDR089W | 0.001 | -0.010 | -0.004 | 0 |
| GUP2 | -0.009 | 0.005 | -0.002 | 0 |
| YDR102C | -0.008 | -0.008 | -0.008 | 0 |
| TRE1 | 0.000 | -0.010 | -0.005 | 0 |
| MRPL1 | 0.001 | 0.007 | 0.004 | 0 |
| YPL162C | -0.001 | -0.007 | -0.004 | 0 |
| SAC6 | -0.001 | -0.014 | -0.007 | 0 |
| POC4 | 0.075 | 0.079 | 0.077 | 1 |
| PEX7 | 0.006 | 0.009 | 0.008 | 0 |
| RNY1 | 0.010 | 0.004 | 0.007 | 0 |
| ATP22 | -0.034 | -0.032 | -0.033 | -1 |
| GDE1 | -0.010 | -0.007 | -0.008 | 0 |
| XRS2 | 0.047 | 0.060 | 0.053 | 1 |
| MGR2 | 0.029 | 0.031 | 0.030 | 1 |
| ATO3 | 0.001 | 0.003 | 0.002 | 0 |
| DTR1 | 0.001 | 0.002 | 0.001 | 0 |
| URH1 | -0.016 | 0.005 | -0.005 | 0 |
| KTR4 | -0.013 | -0.022 | -0.018 | 0 |
| RPL12B | 0.037 | 0.044 | 0.040 | 1 |
| MET8 | -0.006 | -0.003 | -0.005 | 0 |
| YDR433W | -0.011 | -0.009 | -0.010 | 0 |
| YBR225W | 0.011 | 0.010 | 0.010 | 0 |
| VAC8 | -0.021 | -0.009 | -0.015 | 0 |
| YBR241C | -0.002 | -0.009 | -0.005 | 0 |
| YEL028W | -0.001 | 0.003 | 0.001 | 0 |
| RGD1 | -0.006 | -0.007 | -0.006 | 0 |
| YEL045C | 0.000 | -0.054 | -0.027 | 0 |
| PST1 | -0.004 | 0.006 | 0.001 | 0 |
| YEL057C | -0.010 | -0.006 | -0.008 | 0 |
| YPL216W | -0.005 | 0.001 | -0.002 | 0 |
| SNF11 | 0.010 | -0.007 | 0.002 | 0 |
| YIG1 | -0.006 | 0.001 | -0.003 | 0 |
| YDR090C | -0.010 | -0.011 | -0.011 | 0 |
| POS5 | -0.011 | -0.018 | -0.015 | 0 |
| STE5 | 0.018 | 0.014 | 0.016 | 0 |
| NIP100 | 0.017 | 0.011 | 0.014 | 0 |
| TMA64 | -0.012 | -0.001 | -0.006 | 0 |
| BEM4 | 0.026 | 0.009 | 0.018 | 0 |
| FIN1 | -0.010 | -0.008 | -0.009 | 0 |
| YPL141C | 0.001 | -0.004 | -0.001 | 0 |
| SAN1 | 0.013 | 0.017 | 0.015 | 0 |
| MEI5 | -0.010 | -0.007 | -0.009 | 0 |
| SBE2 | -0.012 | 0.007 | -0.002 | 0 |
| YPL109C | 0.003 | -0.001 | 0.001 | 0 |
| YDR370C | -0.001 | 0.003 | 0.001 | 0 |

| | | | | |
|---|---|---|---|---|
| MSY1 | 0.034 | 0.024 | 0.029 | 1 |
| EFT2 | 0.003 | 0.010 | 0.007 | 0 |
| RPS6B | 0.007 | -0.004 | 0.001 | 0 |
| YDR401W | -0.005 | 0.000 | -0.002 | 0 |
| BEM1 | 0.008 | -0.015 | -0.003 | 0 |
| RAD30 | -0.010 | 0.007 | -0.001 | 0 |
| SDS24 | -0.004 | 0.004 | 0.000 | 0 |
| PPM1 | 0.015 | 0.014 | 0.015 | 0 |
| YBR226C | 0.011 | 0.009 | 0.010 | 0 |
| YEL014C | 0.006 | 0.013 | 0.009 | 0 |
| YBR242W | 0.010 | 0.004 | 0.007 | 0 |
| ECM10 | -0.008 | 0.001 | -0.004 | 0 |
| TAE1 | -0.005 | 0.003 | -0.001 | 0 |
| GLY1 | -0.004 | -0.001 | -0.003 | 0 |
| YDR056C | 0.002 | 0.008 | 0.005 | 0 |
| YEL059W | 0.000 | -0.001 | -0.001 | 0 |
| CBP3 | -0.007 | -0.004 | -0.006 | 0 |
| PPH3 | -0.015 | -0.010 | -0.013 | 0 |
| CSM4 | -0.004 | -0.002 | -0.003 | 0 |
| UBC13 | -0.008 | -0.033 | -0.020 | 0 |
| MF(ALPHA) | -0.007 | -0.001 | -0.004 | 0 |
| SPO71 | -0.005 | -0.010 | -0.007 | 0 |
| MRPL40 | 0.000 | -0.004 | -0.002 | 0 |
| VBA4 | 0.003 | 0.008 | 0.005 | 0 |
| PET20 | -0.007 | -0.001 | -0.004 | 0 |
| YDR131C | 0.005 | 0.007 | 0.006 | 0 |
| MKK2 | -0.016 | -0.014 | -0.015 | 0 |
| MKC7 | -0.008 | -0.010 | -0.009 | 0 |
| VPS30 | -0.043 | -0.035 | -0.039 | -1 |
| YDR352W | 0.003 | -0.019 | -0.008 | 0 |
| YPL108W | -0.003 | 0.002 | 0.000 | 0 |
| CTS2 | -0.008 | 0.008 | 0.000 | 0 |
| PNG1 | -0.012 | -0.005 | -0.008 | 0 |
| MUS81 | 0.006 | 0.009 | 0.007 | 0 |
| SMP1 | -0.005 | 0.012 | 0.003 | 0 |
| DIT2 | 0.007 | -0.006 | 0.001 | 0 |
| DER1 | -0.011 | -0.013 | -0.012 | 0 |
| HKR1 | -0.005 | -0.001 | -0.003 | 0 |
| HPC2 | -0.014 | 0.000 | -0.007 | 0 |
| PPZ2 | 0.010 | -0.006 | 0.002 | 0 |
| MCX1 | 0.008 | -0.006 | 0.001 | 0 |
| EDC3 | 0.002 | 0.002 | 0.002 | 0 |
| GPX2 | -0.001 | 0.001 | 0.000 | 0 |
| SPF1 | 1.126 | 1.093 | 1.109 | 1 |
| YBR262C | 0.010 | 0.003 | 0.006 | 0 |
| YEL047C | 0.005 | 0.005 | 0.005 | 0 |
| YOS9 | 0.007 | -0.002 | 0.003 | 0 |
| PRB1 | 0.005 | -0.005 | 0.000 | 0 |
| THI6 | 0.020 | 0.025 | 0.022 | 0 |
| RAD55 | 0.008 | 0.024 | 0.016 | 0 |
| YPL199C | -0.009 | -0.005 | -0.007 | 0 |
| DNF2 | -0.014 | -0.008 | -0.011 | 0 |
| YPL185W | -0.011 | -0.005 | -0.008 | 0 |

| | | | | |
|---|---|---|---|---|
| TMS1 | -0.012 | -0.005 | -0.008 | 0 |
| COX10 | 0.001 | 0.002 | 0.001 | 0 |
| TRM1 | -0.031 | -0.043 | -0.037 | -1 |
| TGS1 | 0.051 | 0.054 | 0.052 | 1 |
| YDR132C | 0.008 | 0.006 | 0.007 | 0 |
| UME1 | -0.018 | -0.004 | -0.011 | 0 |
| SWI5 | -0.005 | -0.034 | -0.019 | 0 |
| DBP1 | -0.002 | 0.004 | 0.001 | 0 |
| TRP4 | 0.003 | 0.004 | 0.003 | 0 |
| YPL107W | -0.030 | -0.013 | -0.021 | 0 |
| VPS74 | 0.005 | 0.003 | 0.004 | 0 |
| EEB1 | -0.008 | -0.008 | -0.008 | 0 |
| YDR387C | 0.000 | 0.006 | 0.003 | 0 |
| YPC1 | 0.004 | 0.011 | 0.007 | 0 |
| DIT1 | -0.014 | 0.000 | -0.007 | 0 |
| COS111 | -0.002 | -0.001 | -0.001 | 0 |
| ARO80 | -0.015 | 0.000 | -0.008 | 0 |
| YBP1 | -0.001 | -0.002 | -0.002 | 0 |
| IRC22 | 0.006 | 0.003 | 0.005 | 0 |
| SLX1 | 0.004 | 0.007 | 0.005 | 0 |
| NPP2 | 0.001 | 0.005 | 0.003 | 0 |
| ISW1 | -0.050 | -0.029 | -0.039 | -1 |
| MTC7 | 0.013 | 0.019 | 0.016 | 0 |
| SHM1 | -0.004 | -0.008 | -0.006 | 0 |
| YEL048C | 0.018 | 0.009 | 0.013 | 0 |
| UBC5 | 0.001 | 0.008 | 0.004 | 0 |
| CIN8 | 0.035 | 0.047 | 0.041 | 1 |
| LEA1 | 0.017 | 0.023 | 0.020 | 0 |
| SED1 | -0.005 | -0.020 | -0.013 | 0 |
| RPL7B | -0.001 | -0.009 | -0.005 | 0 |
| YDR094W | 0.008 | 0.009 | 0.008 | 0 |
| UIP4 | 0.001 | 0.008 | 0.004 | 0 |
| TMN2 | -0.019 | -0.012 | -0.016 | 0 |
| OYE3 | -0.006 | 0.000 | -0.003 | 0 |
| DPB4 | 0.030 | 0.033 | 0.032 | 1 |
| PRM4 | 0.002 | 0.004 | 0.003 | 0 |
| YDR133C | -0.010 | 0.000 | -0.005 | 0 |
| SPP1 | -0.015 | -0.040 | -0.027 | 0 |
| YDR338C | 0.001 | 0.000 | 0.000 | 0 |
| MRP51 | -0.004 | -0.001 | -0.002 | 0 |
| YDR357C | -0.009 | 0.005 | -0.002 | 0 |
| SSE1 | 0.072 | 0.081 | 0.077 | 1 |
| YDR374C | 0.004 | 0.002 | 0.003 | 0 |
| SSU1 | -0.008 | -0.001 | -0.005 | 0 |
| RVS167 | 0.009 | 0.016 | 0.013 | 0 |
| YBR184W | -0.003 | 0.001 | -0.001 | 0 |
| YBR204C | 0.003 | 0.010 | 0.006 | 0 |
| SIP1 | -0.032 | -0.004 | -0.018 | 0 |
| ATG12 | 0.007 | 0.004 | 0.005 | 0 |
| GIM4 | 0.002 | 0.004 | 0.003 | 0 |
| ROT2 | 0.025 | 0.023 | 0.024 | 1 |
| PMP2 | -0.011 | -0.025 | -0.018 | 0 |
| YBR246W | -0.037 | -0.029 | -0.033 | -1 |

| | | | | |
|---|---|---|---|---|
| YPR091C | 0.013 | 0.016 | 0.014 | 0 |
| YPT10 | 0.001 | 0.000 | 0.001 | 0 |
| PAU2 | -0.001 | 0.009 | 0.004 | 0 |
| YDR061W | 0.005 | 0.007 | 0.006 | 0 |
| NPR2 | -0.006 | -0.002 | -0.004 | 0 |
| CHO1 | -0.023 | -0.005 | -0.014 | 0 |
| SHU2 | 0.001 | -0.022 | -0.010 | 0 |
| YPL197C | -0.013 | -0.007 | -0.010 | 0 |
| YDR095C | -0.025 | -0.029 | -0.027 | -1 |
| YPL184C | -0.008 | -0.011 | -0.009 | 0 |
| GSG1 | -0.035 | -0.023 | -0.029 | -1 |
| DAP1 | 0.001 | -0.005 | -0.002 | 0 |
| KIN1 | -0.003 | 0.001 | -0.001 | 0 |
| KIP2 | -0.007 | -0.008 | -0.007 | 0 |
| YDR134C | -0.005 | -0.002 | -0.004 | 0 |
| YPL136W | -0.024 | -0.044 | -0.034 | -1 |
| YDR340W | 0.002 | 0.001 | 0.001 | 0 |
| HOS3 | 0.001 | -0.004 | -0.002 | 0 |
| GGA1 | -0.005 | -0.011 | -0.008 | 0 |
| SYH1 | -0.004 | 0.002 | -0.001 | 0 |
| BCS1 | -0.006 | -0.001 | -0.004 | 0 |
| YBR174C | -0.067 | -0.073 | -0.070 | -1 |
| SAC7 | 0.012 | 0.013 | 0.013 | 0 |
| MBA1 | -0.004 | 0.004 | 0.000 | 0 |
| PDR15 | -0.003 | 0.003 | 0.000 | 0 |
| KTR3 | 0.016 | 0.016 | 0.016 | 0 |
| CAD1 | -0.004 | 0.003 | -0.001 | 0 |
| PYC2 | -0.018 | -0.026 | -0.022 | 0 |
| YEA4 | 0.009 | 0.008 | 0.009 | 0 |
| OM14 | 0.000 | 0.003 | 0.002 | 0 |
| GTT3 | 0.000 | -0.004 | -0.002 | 0 |
| HIS7 | 0.013 | 0.004 | 0.009 | 0 |
| RAD23 | 0.027 | 0.032 | 0.030 | 1 |
| SLM6 | 0.035 | 0.035 | 0.035 | 1 |
| RML2 | -0.034 | -0.081 | -0.058 | -1 |
| YDR063W | 0.000 | -0.002 | -0.001 | 0 |
| CAN1 | -0.003 | 0.002 | 0.000 | 0 |
| PUS1 | -0.004 | -0.007 | -0.005 | 0 |
| PET100 | 0.030 | 0.026 | 0.028 | 1 |
| OXR1 | -0.050 | -0.076 | -0.063 | -1 |
| GIS1 | -0.004 | -0.011 | -0.007 | 0 |
| CTI6 | 0.010 | 0.008 | 0.009 | 0 |
| YDR109C | -0.005 | 0.007 | 0.001 | 0 |
| YPL168W | -0.003 | 0.005 | 0.001 | 0 |
| INO2 | 0.080 | 0.057 | 0.069 | 1 |
| PEP4 | -0.004 | 0.006 | 0.001 | 0 |
| YCF1 | 0.003 | 0.004 | 0.003 | 0 |
| ISU1 | -0.004 | 0.007 | 0.001 | 0 |
| YDR344C | 0.002 | 0.007 | 0.004 | 0 |
| BEM3 | 0.008 | 0.011 | 0.009 | 0 |
| EAF1 | 0.008 | 0.013 | 0.011 | 0 |
| MSD1 | 0.012 | -0.030 | -0.009 | 0 |
| ATP17 | 0.031 | 0.037 | 0.034 | 1 |

| | | | | |
|---|---|---|---|---|
| YDR391C | -0.007 | -0.009 | -0.008 | 0 |
| PCH2 | -0.014 | -0.012 | -0.013 | 0 |
| ADE8 | 0.005 | 0.009 | 0.007 | 0 |
| YBR206W | 0.008 | 0.016 | 0.012 | 0 |
| SNX41 | 0.001 | -0.004 | -0.002 | 0 |
| YBR219C | -0.009 | -0.009 | -0.009 | 0 |
| VAB2 | -0.011 | 0.001 | -0.005 | 0 |
| SWC5 | -0.067 | -0.047 | -0.057 | -1 |
| MDM36 | 0.005 | 0.010 | 0.008 | 0 |
| ARO4 | -0.007 | 0.000 | -0.003 | 0 |
| UTR4 | 0.014 | 0.028 | 0.021 | 0 |
| REI1 | 0.018 | 0.021 | 0.019 | 0 |
| VMA8 | -0.006 | 0.000 | -0.003 | 0 |
| AVT2 | 0.003 | -0.004 | 0.000 | 0 |
| RKM1 | -0.005 | -0.001 | -0.003 | 0 |
| APL5 | -0.029 | -0.270 | -0.149 | -1 |
| MSH6 | 0.007 | 0.009 | 0.008 | 0 |
| YPL182C | 0.007 | 0.006 | 0.007 | 0 |
| FOB1 | -0.009 | -0.030 | -0.019 | 0 |
| REV3 | -0.003 | 0.002 | -0.001 | 0 |
| YDR124W | -0.003 | 0.003 | 0.000 | 0 |
| RRD2 | 0.013 | 0.012 | 0.013 | 0 |
| VPS61 | 0.042 | -0.017 | 0.013 | 0 |
| RDS2 | -0.013 | 0.007 | -0.003 | 0 |
| HXT3 | -0.007 | 0.002 | -0.002 | 0 |
| YPL114W | 0.001 | -0.009 | -0.004 | 0 |
| OPI7 | -0.004 | -0.001 | -0.003 | 0 |
| FMP30 | 0.020 | 0.025 | 0.022 | 0 |
| LSM6 | -0.005 | 0.011 | 0.003 | 0 |
| SWD3 | 0.044 | 0.030 | 0.037 | 1 |
| SPT3 | 0.009 | 0.029 | 0.019 | 0 |
| GDT1 | 0.011 | 0.003 | 0.007 | 0 |
| SIZ1 | 0.004 | 0.002 | 0.003 | 0 |
| FTH1 | -0.016 | -0.010 | -0.013 | 0 |
| YDR426C | 0.018 | 0.031 | 0.025 | 0 |
| YBR220C | -0.002 | 0.004 | 0.001 | 0 |
| YEA6 | 0.000 | 0.018 | 0.009 | 0 |
| PBP2 | 0.015 | -0.009 | 0.003 | 0 |
| YEL020C | -0.009 | -0.003 | -0.006 | 0 |
| SPO23 | -0.004 | 0.011 | 0.003 | 0 |
| CYC7 | -0.001 | -0.009 | -0.005 | 0 |
| MRPL37 | -0.005 | 0.004 | -0.001 | 0 |
| AFG1 | 0.000 | 0.001 | 0.001 | 0 |
| RTR2 | -0.002 | 0.008 | 0.003 | 0 |
| SIT1 | -0.003 | -0.006 | -0.005 | 0 |
| TYW1 | -0.005 | -0.002 | -0.003 | 0 |
| VPS41 | -0.024 | -0.086 | -0.055 | -1 |
| DDC1 | -0.003 | -0.026 | -0.014 | 0 |
| GRX3 | -0.012 | -0.065 | -0.039 | 0 |
| TCO89 | -0.021 | -0.106 | -0.064 | -1 |
| ALT2 | -0.004 | -0.001 | -0.002 | 0 |
| ATG29 | 0.002 | -0.057 | -0.027 | 0 |
| ECM18 | -0.013 | 0.014 | 0.000 | 0 |

| | | | | |
|---|---|---|---|---|
| YPL150W | 0.011 | -0.010 | 0.001 | 0 |
| RGP1 | -0.044 | -0.109 | -0.076 | -1 |
| SPO19 | -0.006 | -0.012 | -0.009 | 0 |
| SVF1 | 0.014 | 0.027 | 0.021 | 0 |
| YPL113C | -0.004 | -0.028 | -0.016 | 0 |
| ESC2 | 0.001 | 0.008 | 0.005 | 0 |
| ELP4 | 0.055 | 0.046 | 0.050 | 1 |
| RGA2 | 0.006 | 0.004 | 0.005 | 0 |
| ECM31 | -0.010 | -0.002 | -0.006 | 0 |
| NTC20 | -0.010 | 0.001 | -0.005 | 0 |
| STE14 | 0.023 | 0.019 | 0.021 | 0 |
| DUR1,2 | 0.005 | 0.009 | 0.007 | 0 |
| BNA7 | 0.003 | 0.006 | 0.005 | 0 |
| PDB1 | 0.015 | 0.013 | 0.014 | 0 |
| YEL007W | -0.087 | -0.074 | -0.080 | -1 |
| YBR235W | 0.003 | -0.005 | -0.001 | 0 |
| YEL023C | -0.015 | -0.012 | -0.014 | 0 |
| MRPS5 | 0.000 | 0.011 | 0.006 | 0 |
| UTR2 | 0.004 | 0.018 | 0.011 | 0 |
| YDR049W | 0.010 | -0.031 | -0.010 | 0 |
| MAK10 | -0.043 | -0.067 | -0.055 | -1 |
| OCA6 | 0.032 | 0.036 | 0.034 | 1 |
| HPA3 | -0.003 | 0.011 | 0.004 | 0 |
| YEL067C | -0.007 | -0.009 | -0.008 | 0 |
| RRM3 | 0.024 | 0.027 | 0.026 | 1 |
| YHR049C-A | -0.009 | -0.007 | -0.008 | 0 |
| YER038W-A | -0.001 | 0.006 | 0.003 | 0 |
| YHR078W | -0.012 | -0.010 | -0.011 | 0 |
| JHD1 | -0.003 | -0.002 | -0.003 | 0 |
| HXT5 | -0.004 | -0.002 | -0.003 | 0 |
| HOR2 | -0.014 | -0.002 | -0.008 | 0 |
| YHR113W | 0.006 | 0.005 | 0.005 | 0 |
| RPS24A | 0.004 | -0.004 | 0.000 | 0 |
| YHR130C | 0.005 | 0.002 | 0.004 | 0 |
| YGR125W | 0.017 | 0.013 | 0.015 | 0 |
| MRPL6 | -0.086 | -0.052 | -0.069 | -1 |
| YGR139W | 0.002 | 0.003 | 0.003 | 0 |
| YAP1801 | 0.000 | 0.000 | 0.000 | 0 |
| YGR153W | -0.011 | -0.006 | -0.008 | 0 |
| NVJ1 | -0.005 | 0.002 | -0.002 | 0 |
| PUS6 | 0.007 | 0.001 | 0.004 | 0 |
| YCL002C | -0.006 | -0.011 | -0.008 | 0 |
| QCR9 | 0.016 | 0.014 | 0.015 | 0 |
| YCL023C | 0.011 | 0.003 | 0.007 | 0 |
| YGR203W | -0.005 | 0.001 | -0.002 | 0 |
| SRO9 | -0.011 | -0.006 | -0.009 | 0 |
| YHL046C | -0.004 | -0.001 | -0.002 | 0 |
| KAR4 | 0.005 | 0.005 | 0.005 | 0 |
| RPL8A | 0.018 | 0.025 | 0.021 | 0 |
| YCP4 | -0.005 | -0.001 | -0.003 | 0 |
| APM2 | -0.002 | -0.004 | -0.003 | 0 |
| MAK32 | -0.005 | 0.000 | -0.002 | 0 |
| LAG1 | 0.011 | 0.014 | 0.012 | 0 |

| | | | | |
|---|---|---|---|---|
| CKI1 | 0.008 | 0.011 | 0.009 | 0 |
| YEL068C | 0.000 | -0.001 | 0.000 | 0 |
| PIH1 | 0.023 | 0.022 | 0.022 | 1 |
| YPR011C | 0.000 | 0.005 | 0.002 | 0 |
| SMF2 | -0.012 | -0.012 | -0.012 | 0 |
| HVG1 | -0.003 | 0.002 | 0.000 | 0 |
| IRE1 | 0.016 | 0.013 | 0.015 | 0 |
| HOM3 | -0.002 | 0.003 | 0.000 | 0 |
| YHR097C | 0.005 | 0.003 | 0.004 | 0 |
| ICL1 | -0.005 | -0.001 | -0.003 | 0 |
| BZZ1 | 0.001 | 0.001 | 0.001 | 0 |
| PTP3 | 0.005 | 0.010 | 0.008 | 0 |
| ECM14 | 0.008 | 0.001 | 0.005 | 0 |
| YGR126W | 0.004 | 0.003 | 0.003 | 0 |
| PEX28 | -0.003 | 0.006 | 0.002 | 0 |
| GTO1 | 0.001 | -0.002 | 0.000 | 0 |
| YHR198C | 0.000 | 0.002 | 0.001 | 0 |
| YGR161W-C | 0.002 | 0.003 | 0.002 | 0 |
| LDB16 | -0.029 | -0.026 | -0.027 | -1 |
| UBR1 | 0.006 | 0.011 | 0.008 | 0 |
| KCC4 | -0.004 | 0.000 | -0.002 | 0 |
| YGR205W | 0.000 | 0.002 | 0.001 | 0 |
| GID7 | -0.007 | 0.015 | 0.004 | 0 |
| YHL045W | -0.005 | 0.002 | -0.001 | 0 |
| YCL056C | 0.003 | -0.007 | -0.002 | 0 |
| GUT1 | -0.012 | 0.000 | -0.006 | 0 |
| CIT2 | -0.048 | -0.051 | -0.049 | -1 |
| YHL017W | -0.025 | -0.010 | -0.017 | 0 |
| PET18 | 0.001 | 0.001 | 0.001 | 0 |
| RPL14B | -0.003 | -0.005 | -0.004 | 0 |
| DLD3 | -0.002 | -0.003 | -0.003 | 0 |
| YHR035W | -0.013 | 0.007 | -0.003 | 0 |
| AFG3 | -0.003 | 0.000 | -0.002 | 0 |
| GLN3 | 0.022 | 0.017 | 0.020 | 0 |
| YHR080C | -0.006 | 0.003 | -0.002 | 0 |
| PIC2 | 0.004 | 0.001 | 0.002 | 0 |
| YHR100C | 0.049 | 0.041 | 0.045 | 1 |
| YER066C-A | -0.004 | -0.004 | -0.004 | 0 |
| DMA1 | -0.003 | 0.000 | -0.002 | 0 |
| YER079W | -0.010 | -0.008 | -0.009 | 0 |
| NSG1 | 0.010 | 0.004 | 0.007 | 0 |
| YGR127W | -0.003 | 0.001 | -0.001 | 0 |
| MTC6 | 0.000 | -0.005 | -0.002 | 0 |
| VPS62 | 0.007 | 0.001 | 0.004 | 0 |
| SOL3 | 0.014 | 0.007 | 0.010 | 0 |
| CHO2 | 0.137 | 0.132 | 0.135 | 1 |
| YHR199C | 0.001 | -0.001 | 0.000 | 0 |
| PSD2 | -0.011 | -0.013 | -0.012 | 0 |
| YMR118C | 0.013 | 0.006 | 0.009 | 0 |
| HGH1 | -0.028 | -0.028 | -0.028 | -1 |
| AGP1 | -0.001 | 0.001 | 0.000 | 0 |
| MVB12 | -0.012 | -0.012 | -0.012 | 0 |
| GLK1 | -0.002 | 0.006 | 0.002 | 0 |

| | | | | |
|---|---|---|---|---|
| YHL044W | 0.005 | -0.004 | 0.000 | 0 |
| PRD1 | 0.002 | 0.001 | 0.001 | 0 |
| GOS1 | 0.007 | 0.018 | 0.013 | 0 |
| YCR006C | -0.048 | -0.066 | -0.057 | -1 |
| DUR3 | -0.013 | -0.001 | -0.007 | 0 |
| MAK31 | 0.017 | 0.008 | 0.012 | 0 |
| QCR10 | 0.000 | 0.008 | 0.004 | 0 |
| PDC5 | 0.007 | 0.003 | 0.005 | 0 |
| RMD6 | -0.003 | 0.001 | -0.001 | 0 |
| PUT2 | -0.006 | 0.000 | -0.003 | 0 |
| ISC1 | -0.023 | -0.027 | -0.025 | -1 |
| COX6 | -0.006 | 0.000 | -0.003 | 0 |
| YEN1 | -0.002 | 0.000 | -0.001 | 0 |
| LRP1 | 0.026 | 0.019 | 0.023 | 0 |
| GIP2 | -0.002 | -0.018 | -0.010 | 0 |
| SBE22 | -0.005 | -0.001 | -0.003 | 0 |
| YER067W | 0.004 | 0.004 | 0.004 | 0 |
| COX23 | 0.019 | 0.031 | 0.025 | 0 |
| AIM9 | -0.008 | -0.005 | -0.007 | 0 |
| WSS1 | 0.012 | 0.017 | 0.014 | 0 |
| SYF2 | -0.007 | 0.003 | -0.002 | 0 |
| SPO12 | 0.007 | 0.005 | 0.006 | 0 |
| BTN2 | 0.001 | 0.002 | 0.001 | 0 |
| THP2 | 0.003 | 0.009 | 0.006 | 0 |
| NSR1 | 0.008 | 0.008 | 0.008 | 0 |
| RPN10 | 0.073 | 0.080 | 0.076 | 1 |
| MSM1 | -0.015 | -0.014 | -0.015 | 0 |
| CRH1 | -0.002 | -0.003 | -0.003 | 0 |
| YCL026C | -0.004 | 0.000 | -0.002 | 0 |
| YGR207C | 0.004 | 0.005 | 0.004 | 0 |
| YCL042W | -0.009 | -0.006 | -0.007 | 0 |
| ECM34 | 0.000 | -0.007 | -0.003 | 0 |
| YCL060C | 0.021 | 0.024 | 0.023 | 1 |
| ECM29 | 0.002 | 0.011 | 0.007 | 0 |
| YCR007C | -0.007 | -0.001 | -0.004 | 0 |
| YLF2 | -0.005 | 0.000 | -0.002 | 0 |
| HSP30 | 0.007 | 0.006 | 0.006 | 0 |
| DIA4 | -0.051 | -0.039 | -0.045 | -1 |
| SLX4 | -0.001 | 0.000 | -0.001 | 0 |
| MNN1 | -0.002 | -0.004 | -0.003 | 0 |
| RRF1 | -0.005 | -0.007 | -0.006 | 0 |
| SBH2 | 0.007 | 0.009 | 0.008 | 0 |
| CPR2 | 0.002 | 0.007 | 0.005 | 0 |
| MXR1 | 0.001 | -0.003 | -0.001 | 0 |
| KSP1 | -0.006 | -0.005 | -0.005 | 0 |
| FCY2 | 0.059 | 0.057 | 0.058 | 1 |
| GRE3 | -0.005 | -0.002 | -0.004 | 0 |
| YER067C-A | 0.001 | 0.000 | 0.000 | 0 |
| TOM71 | -0.013 | -0.025 | -0.019 | 0 |
| SER3 | -0.045 | -0.053 | -0.049 | -1 |
| YCK1 | -0.020 | -0.018 | -0.019 | 0 |
| YGR130C | -0.010 | 0.000 | -0.005 | 0 |
| SPO16 | 0.006 | -0.002 | 0.002 | 0 |

| | | | | |
|---|---|---|---|---|
| SKN1 | 0.001 | -0.007 | -0.003 | 0 |
| FMO1 | 0.001 | 0.005 | 0.003 | 0 |
| YGR160W | 0.008 | 0.011 | 0.009 | 0 |
| YHR202W | -0.014 | -0.004 | -0.009 | 0 |
| RBG2 | -0.008 | 0.001 | -0.003 | 0 |
| STP22 | -0.024 | -0.025 | -0.025 | -1 |
| TDH3 | 0.002 | -0.004 | -0.001 | 0 |
| FUS1 | -0.020 | 0.000 | -0.010 | 0 |
| SER2 | -0.016 | -0.006 | -0.011 | 0 |
| MGR1 | 0.004 | 0.009 | 0.007 | 0 |
| YHL042W | -0.003 | 0.001 | -0.001 | 0 |
| MRC1 | 0.027 | 0.033 | 0.030 | 1 |
| OCA5 | 0.024 | 0.025 | 0.025 | 1 |
| SAT4 | -0.003 | 0.006 | 0.001 | 0 |
| OTU2 | 0.001 | 0.003 | 0.002 | 0 |
| YCR022C | 0.007 | -0.009 | -0.001 | 0 |
| VPS29 | -0.051 | -0.033 | -0.042 | -1 |
| TIS11 | -0.006 | -0.003 | -0.005 | 0 |
| NOP16 | -0.003 | 0.000 | -0.001 | 0 |
| MSC7 | -0.022 | 0.000 | -0.011 | 0 |
| GPA2 | 0.002 | 0.007 | 0.004 | 0 |
| VMA22 | -0.009 | -0.001 | -0.005 | 0 |
| MEI4 | 0.000 | 0.005 | 0.002 | 0 |
| NAM8 | 0.020 | 0.021 | 0.020 | 0 |
| RPL34A | 0.011 | 0.007 | 0.009 | 0 |
| YPT35 | -0.054 | -0.005 | -0.030 | 0 |
| MOT2 | 0.073 | 0.074 | 0.074 | 1 |
| MSH1 | 0.024 | 0.036 | 0.030 | 1 |
| GET2 | 0.015 | 0.008 | 0.012 | 0 |
| SPL2 | 0.003 | -0.001 | 0.001 | 0 |
| YGR131W | 0.002 | 0.004 | 0.003 | 0 |
| RTT107 | 0.016 | 0.016 | 0.016 | 0 |
| THI4 | -0.001 | 0.009 | 0.004 | 0 |
| YHR177W | 0.009 | 0.006 | 0.008 | 0 |
| RTS3 | -0.004 | -0.006 | -0.005 | 0 |
| RPS4B | 0.001 | 0.011 | 0.006 | 0 |
| CBP4 | 0.008 | 0.017 | 0.012 | 0 |
| ILV6 | -0.013 | -0.004 | -0.008 | 0 |
| PDX1 | 0.004 | 0.007 | 0.006 | 0 |
| RNQ1 | -0.003 | -0.004 | -0.003 | 0 |
| TRX2 | -0.005 | -0.006 | -0.006 | 0 |
| YCL045C | 0.018 | 0.018 | 0.018 | 0 |
| YHL041W | -0.001 | -0.006 | -0.003 | 0 |
| YCL062W | -0.001 | 0.012 | 0.005 | 0 |
| WSC4 | 0.006 | -0.004 | 0.001 | 0 |
| RVS161 | -0.002 | 0.010 | 0.004 | 0 |
| YHL012W | -0.001 | 0.002 | 0.001 | 0 |
| YCR023C | -0.007 | 0.000 | -0.003 | 0 |
| ARD1 | 0.017 | 0.036 | 0.027 | 0 |
| YLR137W | 0.003 | 0.008 | 0.005 | 0 |
| FMP52 | -0.015 | -0.008 | -0.011 | 0 |
| DOG2 | -0.004 | -0.005 | -0.004 | 0 |
| YAT2 | -0.005 | -0.001 | -0.003 | 0 |

| | | | | |
|---|---|---|---|---|
| GIC1 | -0.013 | 0.002 | -0.005 | 0 |
| ACA1 | 0.005 | -0.005 | 0.000 | 0 |
| RTC3 | 0.002 | -0.001 | 0.001 | 0 |
| HMF1 | 0.000 | 0.009 | 0.004 | 0 |
| TRR2 | -0.001 | -0.004 | -0.002 | 0 |
| YER068C-A | -0.010 | -0.008 | -0.009 | 0 |
| LSM12 | -0.006 | 0.001 | -0.003 | 0 |
| YER084W | -0.013 | -0.022 | -0.018 | 0 |
| ARO9 | -0.001 | -0.002 | -0.001 | 0 |
| PHB1 | 0.006 | 0.011 | 0.008 | 0 |
| YSP1 | -0.002 | 0.004 | 0.001 | 0 |
| YGR146C | -0.003 | -0.002 | -0.002 | 0 |
| STB5 | 0.004 | 0.002 | 0.003 | 0 |
| GTR2 | 0.000 | -0.006 | -0.003 | 0 |
| MNL1 | 0.001 | 0.000 | 0.001 | 0 |
| YGR176W | 0.003 | 0.011 | 0.007 | 0 |
| SGF29 | 0.014 | 0.019 | 0.017 | 0 |
| XKS1 | -0.007 | 0.000 | -0.004 | 0 |
| BIK1 | -0.027 | -0.027 | -0.027 | -1 |
| SLI1 | 0.000 | 0.006 | 0.003 | 0 |
| YCL046W | 0.018 | 0.022 | 0.020 | 0 |
| ARN1 | -0.008 | 0.001 | -0.004 | 0 |
| VAC17 | 0.000 | 0.002 | 0.001 | 0 |
| RIM101 | -0.006 | -0.009 | -0.008 | 0 |
| ADY2 | -0.005 | -0.009 | -0.007 | 0 |
| YHL010C | 0.002 | -0.003 | 0.000 | 0 |
| URA4 | -0.027 | -0.004 | -0.015 | 0 |
| SPO13 | 0.000 | -0.008 | -0.004 | 0 |
| NHA1 | 0.002 | -0.006 | -0.002 | 0 |
| YND1 | -0.060 | -0.042 | -0.051 | -1 |
| DOG1 | -0.002 | 0.001 | -0.001 | 0 |
| CHZ1 | 0.005 | 0.012 | 0.008 | 0 |
| SSF1 | 0.017 | 0.027 | 0.022 | 0 |
| YER046W-A | 0.003 | -0.004 | 0.000 | 0 |
| MSR1 | -0.008 | -0.004 | -0.006 | 0 |
| GGA2 | 0.016 | 0.011 | 0.013 | 0 |
| ARG5,6 | 0.050 | 0.028 | 0.039 | 1 |
| EPT1 | -0.008 | 0.001 | -0.004 | 0 |
| YER085C | 0.003 | 0.005 | 0.004 | 0 |
| YHR138C | -0.007 | 0.003 | -0.002 | 0 |
| PEX4 | 0.002 | -0.001 | 0.000 | 0 |
| LIN1 | -0.004 | -0.002 | -0.003 | 0 |
| RPL24B | 0.004 | 0.006 | 0.005 | 0 |
| OYE2 | -0.001 | 0.002 | 0.000 | 0 |
| YGR164W | 0.001 | -0.001 | 0.000 | 0 |
| SKN7 | -0.004 | 0.011 | 0.003 | 0 |
| ATF2 | 0.005 | 0.003 | 0.004 | 0 |
| GBP2 | -0.017 | -0.018 | -0.018 | 0 |
| FYV8 | -0.020 | -0.011 | -0.015 | 0 |
| HIS4 | -0.006 | -0.011 | -0.009 | 0 |
| RTA1 | 0.003 | 0.008 | 0.005 | 0 |
| YCL047C | 0.001 | -0.001 | 0.000 | 0 |
| CBP2 | -0.019 | 0.015 | -0.002 | 0 |

| | | | | |
|---|---|---|---|---|
| CHA1 | 0.000 | -0.001 | 0.000 | 0 |
| YHL026C | 0.002 | 0.005 | 0.003 | 0 |
| ADP1 | 0.003 | 0.008 | 0.005 | 0 |
| YAP3 | 0.010 | 0.006 | 0.008 | 0 |
| LEU3 | -0.008 | -0.005 | -0.006 | 0 |
| MIP6 | -0.006 | 0.002 | -0.002 | 0 |
| SLS1 | -0.017 | -0.012 | -0.015 | 0 |
| TMA20 | 0.004 | 0.009 | 0.006 | 0 |
| INM1 | -0.003 | -0.003 | -0.003 | 0 |
| FIR1 | 0.010 | 0.011 | 0.011 | 0 |
| OSH3 | -0.015 | -0.008 | -0.011 | 0 |
| SAP1 | -0.007 | -0.004 | -0.005 | 0 |
| HXT4 | 0.007 | 0.004 | 0.006 | 0 |
| PCL6 | -0.003 | 0.001 | -0.001 | 0 |
| CTM1 | -0.002 | 0.001 | -0.001 | 0 |
| MRI1 | -0.001 | -0.001 | -0.001 | 0 |
| NDT80 | -0.004 | 0.000 | -0.002 | 0 |
| ILV1 | -0.022 | -0.010 | -0.016 | 0 |
| SPS100 | 0.001 | -0.002 | -0.001 | 0 |
| PRE9 | 0.118 | 0.113 | 0.115 | 1 |
| REC104 | -0.005 | 0.004 | 0.000 | 0 |
| YGR149W | -0.001 | 0.004 | 0.001 | 0 |
| YHR182W | -0.001 | 0.002 | 0.001 | 0 |
| MRPS35 | 0.028 | 0.000 | 0.014 | 0 |
| SET5 | -0.013 | 0.004 | -0.004 | 0 |
| PBP1 | 0.009 | 0.012 | 0.011 | 0 |
| YCL012W | -0.002 | -0.005 | -0.003 | 0 |
| SNG1 | -0.001 | -0.002 | -0.002 | 0 |
| STE50 | 0.021 | 0.018 | 0.019 | 0 |
| RPS0A | 0.007 | 0.004 | 0.005 | 0 |
| SPS22 | -0.002 | -0.006 | -0.004 | 0 |
| YHL037C | -0.011 | -0.001 | -0.006 | 0 |
| VBA3 | -0.001 | 0.000 | -0.001 | 0 |
| RMD11 | -0.008 | -0.080 | -0.044 | 0 |
| POL4 | -0.005 | -0.004 | -0.004 | 0 |
| YHL008C | 0.010 | -0.007 | 0.001 | 0 |
| YLR126C | 0.000 | 0.006 | 0.003 | 0 |
| ARG4 | 0.010 | 0.018 | 0.014 | 0 |
| PUT1 | -0.010 | 0.001 | -0.005 | 0 |
| PAC2 | -0.005 | -0.008 | -0.006 | 0 |
| AAP1 | -0.010 | -0.012 | -0.011 | 0 |
| ZRG8 | -0.006 | 0.006 | 0.000 | 0 |
| PPE1 | -0.003 | -0.004 | -0.003 | 0 |
| CAJ1 | -0.016 | -0.007 | -0.012 | 0 |
| AHT1 | -0.031 | -0.002 | -0.017 | 0 |
| FCY21 | 0.004 | 0.004 | 0.004 | 0 |
| ERP5 | -0.004 | 0.004 | 0.000 | 0 |
| YER071C | 0.000 | 0.004 | 0.002 | 0 |
| YHR125W | 0.010 | 0.007 | 0.009 | 0 |
| YER087C-A | 0.021 | 0.027 | 0.024 | 1 |
| YHR139C-A | 0.001 | -0.009 | -0.004 | 0 |
| LSB1 | 0.011 | 0.001 | 0.006 | 0 |
| KEL1 | 0.007 | 0.016 | 0.011 | 0 |

| | | | | |
|---|---|---|---|---|
| YGR150C | 0.024 | -0.048 | -0.012 | 0 |
| GND1 | -0.001 | 0.013 | 0.006 | 0 |
| KRE11 | 0.004 | 0.006 | 0.005 | 0 |
| CRG1 | -0.005 | -0.008 | -0.007 | 0 |
| RNR4 | 0.073 | 0.055 | 0.064 | 1 |
| YCL013W | 0.002 | -0.003 | -0.001 | 0 |
| PMT6 | 0.000 | -0.005 | -0.003 | 0 |
| YCL033C | 0.001 | 0.000 | 0.001 | 0 |
| RSM27 | 0.087 | -0.032 | 0.028 | 0 |
| YCL049C | -0.004 | 0.002 | -0.001 | 0 |
| MUP3 | -0.010 | 0.000 | -0.005 | 0 |
| YCR001W | 0.006 | -0.003 | 0.002 | 0 |
| SPO11 | -0.018 | -0.002 | -0.010 | 0 |
| YCR015C | 0.008 | -0.002 | 0.003 | 0 |
| STE20 | -0.026 | -0.015 | -0.021 | 0 |
| DCN1 | 0.001 | 0.004 | 0.003 | 0 |
| YHR022C | 0.001 | -0.001 | 0.000 | 0 |
| YLR143W | -0.008 | -0.001 | -0.005 | 0 |
| YER010C | -0.003 | 0.002 | -0.001 | 0 |
| YHK8 | -0.018 | -0.008 | -0.013 | 0 |
| YER034W | 0.007 | 0.010 | 0.008 | 0 |
| PTC7 | 0.001 | 0.002 | 0.002 | 0 |
| TPA1 | -0.001 | 0.002 | 0.001 | 0 |
| HXT1 | -0.003 | -0.009 | -0.006 | 0 |
| FCY22 | 0.015 | 0.007 | 0.011 | 0 |
| UBA4 | 0.032 | 0.022 | 0.027 | 1 |
| VTC1 | -0.003 | -0.004 | -0.004 | 0 |
| YHR126C | 0.002 | 0.004 | 0.003 | 0 |
| PPT1 | -0.004 | -0.004 | -0.004 | 0 |
| CHS7 | 0.012 | 0.012 | 0.012 | 0 |
| YGR137W | -0.002 | 0.001 | 0.000 | 0 |
| YHR159W | -0.001 | 0.002 | 0.000 | 0 |
| YGR151C | 0.003 | 0.007 | 0.005 | 0 |
| SSP1 | 0.005 | 0.001 | 0.003 | 0 |
| YOL153C | -0.009 | 0.002 | -0.004 | 0 |
| YHR210C | -0.013 | -0.005 | -0.009 | 0 |
| TIM13 | -0.005 | 0.003 | -0.001 | 0 |
| BUD3 | 0.021 | 0.020 | 0.021 | 1 |
| ELP2 | 0.050 | 0.049 | 0.050 | 1 |
| LSB5 | -0.001 | 0.003 | 0.001 | 0 |
| CCH1 | -0.009 | -0.016 | -0.012 | 0 |
| APA1 | -0.003 | 0.009 | 0.003 | 0 |
| VMR1 | -0.006 | -0.003 | -0.004 | 0 |
| CDC10 | 0.033 | 0.037 | 0.035 | 1 |
| YHL021C | -0.008 | -0.006 | -0.007 | 0 |
| YCR016W | -0.017 | -0.038 | -0.028 | 0 |
| SHU1 | -0.003 | 0.005 | 0.001 | 0 |
| ZRT2 | 0.000 | 0.004 | 0.002 | 0 |
| YHI9 | 0.001 | -0.002 | 0.000 | 0 |
| ACF2 | -0.001 | 0.002 | 0.001 | 0 |
| TIR1 | -0.007 | -0.010 | -0.009 | 0 |
| FSH1 | -0.016 | 0.002 | -0.007 | 0 |
| EDC2 | -0.011 | -0.002 | -0.007 | 0 |

| | | | | |
|---|---|---|---|---|
| NMD2 | 0.005 | 0.001 | 0.003 | 0 |
| RSM18 | 0.018 | -0.006 | 0.006 | 0 |
| YHR095W | -0.010 | -0.010 | -0.010 | 0 |
| CEM1 | 0.015 | 0.004 | 0.010 | 0 |
| YHR112C | -0.013 | 0.002 | -0.006 | 0 |
| ALD5 | -0.004 | 0.001 | -0.002 | 0 |
| ARP1 | -0.014 | -0.010 | -0.012 | 0 |
| ASN2 | 0.002 | 0.007 | 0.004 | 0 |
| DSE2 | 0.011 | 0.008 | 0.010 | 0 |
| TPO2 | 0.002 | 0.001 | 0.002 | 0 |
| PEX18 | -0.002 | -0.002 | -0.002 | 0 |
| RSR1 | 0.014 | 0.007 | 0.010 | 0 |
| PTH1 | 0.006 | 0.017 | 0.012 | 0 |
| YGR168C | -0.004 | -0.009 | -0.006 | 0 |
| RER1 | 0.023 | 0.024 | 0.024 | 1 |
| YGR182C | -0.001 | 0.005 | 0.002 | 0 |
| DCC1 | 0.009 | 0.021 | 0.015 | 0 |
| PCT1 | 0.008 | 0.004 | 0.006 | 0 |
| GFD2 | -0.004 | -0.021 | -0.013 | 0 |
| ARN2 | -0.002 | -0.002 | -0.002 | 0 |
| LRE1 | -0.032 | -0.019 | -0.025 | 0 |
| SBP1 | 0.005 | 0.000 | 0.003 | 0 |
| MRPL32 | 0.019 | 0.015 | 0.017 | 0 |
| OPI1 | 0.018 | 0.015 | 0.017 | 0 |
| CWH43 | 0.003 | -0.001 | 0.001 | 0 |
| YHL005C | -0.001 | 0.004 | 0.002 | 0 |
| ACE2 | 0.020 | 0.023 | 0.021 | 0 |
| SLT2 | -0.011 | -0.001 | -0.006 | 0 |
| YMR074C | -0.007 | -0.008 | -0.008 | 0 |
| YLR149C | -0.003 | -0.009 | -0.006 | 0 |
| SPE1 | 0.027 | 0.032 | 0.029 | 1 |
| YLR173W | -0.003 | -0.002 | -0.003 | 0 |
| YGR039W | -0.008 | 0.005 | -0.001 | 0 |
| RPL37A | -0.005 | 0.006 | 0.001 | 0 |
| RSC1 | 0.037 | 0.046 | 0.042 | 1 |
| MSS51 | 0.015 | -0.048 | -0.017 | 0 |
| YGR071C | -0.013 | -0.005 | -0.009 | 0 |
| TPC1 | -0.004 | 0.003 | -0.001 | 0 |
| LAC1 | 0.008 | 0.003 | 0.006 | 0 |
| RPS23A | -0.003 | 0.003 | 0.000 | 0 |
| MAE1 | -0.003 | 0.006 | 0.002 | 0 |
| INP53 | 0.012 | 0.014 | 0.013 | 0 |
| YKL047W | -0.003 | 0.000 | -0.002 | 0 |
| IAH1 | -0.009 | -0.009 | -0.009 | 0 |
| YKL063C | -0.025 | -0.003 | -0.014 | 0 |
| RUP1 | -0.012 | -0.002 | -0.007 | 0 |
| YKL075C | 0.018 | 0.013 | 0.015 | 0 |
| YRR1 | -0.002 | 0.010 | 0.004 | 0 |
| YKL091C | -0.001 | -0.004 | -0.003 | 0 |
| MPC54 | 0.004 | -0.003 | 0.000 | 0 |
| AAT1 | -0.007 | -0.024 | -0.016 | 0 |
| THI72 | -0.002 | 0.016 | 0.007 | 0 |
| YKL123W | -0.006 | -0.022 | -0.014 | 0 |

| | | | | |
|---|---|---|---|---|
| NPT1 | -0.016 | -0.019 | -0.018 | 0 |
| YKL136W | 0.011 | 0.009 | 0.010 | 0 |
| YOR225W | 0.001 | -0.013 | -0.006 | 0 |
| SUE1 | -0.013 | -0.030 | -0.021 | 0 |
| ABP140 | -0.003 | -0.004 | -0.004 | 0 |
| KKQ8 | -0.008 | 0.011 | 0.001 | 0 |
| OSW1 | -0.001 | 0.004 | 0.001 | 0 |
| STM1 | 0.020 | 0.021 | 0.021 | 0 |
| ASH1 | -0.002 | 0.003 | 0.001 | 0 |
| IDP2 | 0.001 | 0.006 | 0.004 | 0 |
| BUD9 | -0.007 | -0.006 | -0.007 | 0 |
| SKG3 | 0.003 | 0.013 | 0.008 | 0 |
| LST7 | 0.002 | -0.003 | -0.001 | 0 |
| QRI5 | -0.004 | 0.006 | 0.001 | 0 |
| UPF3 | 0.009 | -0.001 | 0.004 | 0 |
| MSC3 | -0.004 | 0.001 | -0.001 | 0 |
| ASK10 | -0.020 | -0.001 | -0.010 | 0 |
| MRT4 | -0.025 | -0.008 | -0.016 | 0 |
| MEP1 | 0.002 | 0.004 | 0.003 | 0 |
| YKL031W | 0.002 | 0.011 | 0.007 | 0 |
| YOR111W | 0.008 | -0.001 | 0.004 | 0 |
| MNR2 | -0.002 | 0.003 | 0.001 | 0 |
| YOR139C | -0.003 | -0.004 | -0.003 | 0 |
| PSY1 | 0.018 | 0.022 | 0.020 | 0 |
| DDP1 | -0.010 | -0.010 | -0.010 | 0 |
| BUD2 | 0.009 | 0.008 | 0.008 | 0 |
| GAC1 | -0.001 | 0.009 | 0.004 | 0 |
| YKL107W | 0.002 | -0.006 | -0.002 | 0 |
| PEX27 | 0.001 | 0.001 | 0.001 | 0 |
| SSH4 | -0.010 | -0.001 | -0.005 | 0 |
| MGM1 | 0.030 | 0.016 | 0.023 | 0 |
| CMC1 | 0.006 | -0.008 | -0.001 | 0 |
| ISU2 | 0.007 | 0.007 | 0.007 | 0 |
| RPS27A | -0.011 | 0.008 | -0.002 | 0 |
| YOR240W | 0.009 | 0.001 | 0.005 | 0 |
| OSH7 | 0.005 | 0.002 | 0.003 | 0 |
| PCD1 | -0.020 | -0.004 | -0.012 | 0 |
| YKL187C | 0.011 | -0.002 | 0.004 | 0 |
| RFX1 | 0.011 | 0.004 | 0.008 | 0 |
| YGR042W | -0.002 | -0.008 | -0.005 | 0 |
| MDL1 | -0.002 | 0.002 | 0.000 | 0 |
| PEF1 | -0.048 | -0.033 | -0.040 | -1 |
| HMX1 | 0.001 | 0.000 | 0.000 | 0 |
| MRPL25 | -0.007 | -0.005 | -0.006 | 0 |
| CCC1 | -0.007 | 0.000 | -0.003 | 0 |
| MDR1 | -0.001 | 0.001 | 0.000 | 0 |
| UFD4 | 0.010 | 0.022 | 0.016 | 0 |
| YGR122W | -0.023 | -0.091 | -0.057 | -1 |
| IXR1 | 0.016 | 0.019 | 0.018 | 0 |
| CEX1 | 0.010 | 0.016 | 0.013 | 0 |
| ELM1 | 0.005 | 0.025 | 0.015 | 0 |
| RGA1 | -0.007 | 0.000 | -0.003 | 0 |
| YET1 | 0.000 | 0.008 | 0.004 | 0 |

| | | | | |
|---|---|---|---|---|
| SFL1 | 0.007 | 0.004 | 0.006 | 0 |
| YKL077W | -0.024 | -0.043 | -0.034 | -1 |
| YOR164C | 0.005 | 0.002 | 0.004 | 0 |
| MBR1 | 0.006 | -0.005 | 0.000 | 0 |
| RPS30B | -0.018 | -0.003 | -0.010 | 0 |
| HAP4 | -0.189 | 0.007 | -0.091 | 0 |
| SLK19 | 0.017 | 0.019 | 0.018 | 0 |
| YPK1 | -0.008 | -0.003 | -0.006 | 0 |
| STE4 | 0.007 | 0.001 | 0.004 | 0 |
| MRPL31 | 0.017 | 0.024 | 0.020 | 0 |
| HER1 | -0.004 | -0.005 | -0.005 | 0 |
| APE2 | 0.001 | 0.002 | 0.002 | 0 |
| MET7 | -0.016 | -0.039 | -0.028 | 0 |
| YKL169C | -0.029 | -0.035 | -0.032 | -1 |
| HNT3 | 0.007 | 0.015 | 0.011 | 0 |
| YLR152C | -0.006 | -0.005 | -0.006 | 0 |
| PXA2 | -0.003 | -0.010 | -0.007 | 0 |
| YLR177W | 0.002 | -0.003 | 0.000 | 0 |
| NQM1 | -0.006 | -0.011 | -0.009 | 0 |
| ATG26 | -0.007 | -0.007 | -0.007 | 0 |
| SPR3 | -0.004 | 0.000 | -0.002 | 0 |
| ENT2 | -0.005 | -0.002 | -0.003 | 0 |
| PEX8 | -0.006 | 0.004 | -0.001 | 0 |
| RSA3 | -0.007 | -0.016 | -0.011 | 0 |
| PCP1 | 0.050 | 0.058 | 0.054 | 1 |
| CCE1 | -0.023 | -0.007 | -0.015 | 0 |
| YOR097C | -0.006 | -0.007 | -0.006 | 0 |
| TUL1 | 0.012 | 0.011 | 0.011 | 0 |
| AZF1 | -0.003 | -0.027 | -0.015 | 0 |
| YKL050C | -0.004 | -0.003 | -0.003 | 0 |
| AFI1 | 0.001 | -0.002 | -0.001 | 0 |
| YKL066W | -0.007 | -0.001 | -0.004 | 0 |
| ARP8 | 0.014 | 0.016 | 0.015 | 0 |
| SMY1 | -0.017 | -0.041 | -0.029 | 0 |
| SEY1 | 0.023 | 0.007 | 0.015 | 0 |
| YJU3 | -0.006 | -0.001 | -0.004 | 0 |
| FYV12 | -0.004 | 0.002 | -0.001 | 0 |
| KTI12 | 0.041 | 0.043 | 0.042 | 1 |
| LIP5 | 0.007 | 0.013 | 0.010 | 0 |
| PGM1 | 0.017 | 0.017 | 0.017 | 0 |
| SAS5 | -0.001 | -0.004 | -0.002 | 0 |
| TGL1 | -0.007 | -0.009 | -0.008 | 0 |
| YOR228C | 0.009 | 0.003 | 0.006 | 0 |
| YKL158W | -0.006 | -0.006 | -0.006 | 0 |
| SSP2 | 0.006 | 0.012 | 0.009 | 0 |
| MRPL38 | -0.009 | -0.005 | -0.007 | 0 |
| YOR263C | 0.004 | 0.006 | 0.005 | 0 |
| RNH203 | -0.003 | 0.001 | -0.001 | 0 |
| CNB1 | -0.013 | -0.002 | -0.007 | 0 |
| MRP4 | -0.037 | 0.004 | -0.017 | 0 |
| RME1 | -0.006 | -0.018 | -0.012 | 0 |
| MMR1 | 0.042 | 0.032 | 0.037 | 1 |
| ADE6 | 0.002 | -0.013 | -0.005 | 0 |

| | | | | |
|---|---|---|---|---|
| HRD3 | -0.014 | -0.011 | -0.013 | 0 |
| PAC10 | 0.046 | 0.067 | 0.056 | 1 |
| YLR224W | 0.012 | 0.017 | 0.015 | 0 |
| YGR102C | -0.006 | -0.004 | -0.005 | 0 |
| PUT3 | -0.005 | 0.008 | 0.001 | 0 |
| KTR1 | -0.005 | 0.005 | 0.000 | 0 |
| YKL037W | 0.020 | 0.013 | 0.016 | 0 |
| YOR114W | -0.003 | 0.006 | 0.001 | 0 |
| SFK1 | 0.000 | 0.005 | 0.002 | 0 |
| ORT1 | -0.055 | -0.050 | -0.053 | -1 |
| YNK1 | -0.007 | -0.014 | -0.011 | 0 |
| LSC1 | -0.010 | 0.001 | -0.005 | 0 |
| VMA5 | -0.010 | 0.008 | -0.001 | 0 |
| SWT1 | 0.012 | 0.013 | 0.012 | 0 |
| CWP1 | -0.009 | 0.001 | -0.004 | 0 |
| SER1 | -0.006 | 0.006 | 0.000 | 0 |
| RAD27 | 0.011 | 0.029 | 0.020 | 0 |
| MCA1 | -0.011 | -0.010 | -0.010 | 0 |
| PMU1 | 0.001 | -0.004 | -0.002 | 0 |
| YOR214C | 0.007 | 0.004 | 0.006 | 0 |
| MRP8 | -0.001 | -0.004 | -0.002 | 0 |
| WTM2 | -0.012 | 0.003 | -0.004 | 0 |
| RCN1 | -0.002 | -0.015 | -0.009 | 0 |
| PUS7 | 0.023 | 0.029 | 0.026 | 1 |
| YKL171W | 0.007 | 0.006 | 0.006 | 0 |
| DSE3 | 0.005 | 0.005 | 0.005 | 0 |
| YLR164W | -0.002 | 0.000 | -0.001 | 0 |
| RPS25A | -0.010 | 0.006 | -0.002 | 0 |
| TFS1 | 0.008 | 0.001 | 0.005 | 0 |
| YGR045C | -0.002 | 0.004 | 0.001 | 0 |
| PEX13 | -0.007 | 0.002 | -0.002 | 0 |
| PNP1 | -0.003 | -0.004 | -0.004 | 0 |
| YGR079W | -0.005 | -0.006 | -0.005 | 0 |
| YLR225C | 0.004 | 0.006 | 0.005 | 0 |
| SRB5 | -0.041 | -0.021 | -0.031 | -1 |
| ATP7 | 0.025 | 0.027 | 0.026 | 1 |
| CRC1 | -0.007 | -0.004 | -0.006 | 0 |
| RGT1 | 0.000 | -0.009 | -0.005 | 0 |
| TRS33 | -0.010 | 0.004 | -0.003 | 0 |
| YKL053W | 0.025 | 0.037 | 0.031 | 1 |
| YOR131C | 0.010 | 0.006 | 0.008 | 0 |
| NUP100 | 0.013 | 0.008 | 0.010 | 0 |
| ELG1 | 0.019 | 0.026 | 0.023 | 0 |
| RPS28A | -0.005 | 0.004 | -0.001 | 0 |
| YKL097C | -0.038 | 0.005 | -0.016 | 0 |
| GSP2 | -0.001 | -0.003 | -0.002 | 0 |
| APN1 | -0.001 | 0.005 | 0.002 | 0 |
| BFR1 | 0.013 | 0.027 | 0.020 | 0 |
| MYO3 | -0.001 | 0.003 | 0.001 | 0 |
| YOR215C | 0.000 | 0.000 | 0.000 | 0 |
| LTV1 | -0.026 | -0.010 | -0.018 | 0 |
| WTM1 | -0.003 | -0.014 | -0.009 | 0 |
| ELF1 | 0.011 | 0.006 | 0.008 | 0 |

| | | | | |
|---|---|---|---|---|
| DGA1 | 0.005 | 0.009 | 0.007 | 0 |
| TPO5 | 0.004 | -0.006 | -0.001 | 0 |
| YOR277C | 0.017 | 0.016 | 0.016 | 0 |
| PUS5 | -0.005 | -0.004 | -0.004 | 0 |
| YGR031W | -0.004 | 0.000 | -0.002 | 0 |
| YLR179C | -0.015 | -0.005 | -0.010 | 0 |
| UPS1 | 0.053 | 0.047 | 0.050 | 1 |
| YGR064W | -0.014 | -0.009 | -0.011 | 0 |
| CLB4 | -0.008 | -0.005 | -0.006 | 0 |
| TWF1 | -0.030 | -0.004 | -0.017 | 0 |
| ADY4 | 0.002 | 0.008 | 0.005 | 0 |
| VMA21 | -0.055 | -0.003 | -0.029 | 0 |
| HCS1 | -0.001 | 0.005 | 0.002 | 0 |
| RAS1 | -0.004 | -0.007 | -0.006 | 0 |
| PTM1 | -0.028 | -0.057 | -0.042 | -1 |
| RTC5 | -0.010 | 0.005 | -0.003 | 0 |
| DEF1 | 0.006 | 0.015 | 0.010 | 0 |
| VPS17 | -0.015 | -0.017 | -0.016 | 0 |
| YKL069W | -0.012 | 0.005 | -0.004 | 0 |
| YOR152C | -0.002 | 0.002 | 0.000 | 0 |
| TEF4 | -0.005 | -0.028 | -0.017 | 0 |
| YOR170W | -0.027 | -0.037 | -0.032 | -1 |
| MTC2 | -0.004 | -0.037 | -0.021 | 0 |
| YOR186W | 0.010 | 0.002 | 0.006 | 0 |
| PRR1 | 0.002 | 0.005 | 0.004 | 0 |
| YOR199W | -0.022 | -0.038 | -0.030 | -1 |
| SHE2 | 0.007 | -0.012 | -0.003 | 0 |
| RUD3 | -0.011 | -0.003 | -0.007 | 0 |
| AVT3 | -0.006 | -0.008 | -0.007 | 0 |
| MKK1 | -0.010 | -0.008 | -0.009 | 0 |
| YKL161C | 0.003 | 0.011 | 0.007 | 0 |
| YOR246C | 0.013 | 0.022 | 0.017 | 0 |
| ZRT3 | -0.007 | -0.001 | -0.004 | 0 |
| RFM1 | -0.012 | -0.017 | -0.014 | 0 |
| YLR168C | 0.005 | 0.016 | 0.010 | 0 |
| TIM21 | 0.005 | -0.010 | -0.002 | 0 |
| SAM1 | 0.023 | -0.005 | 0.009 | 0 |
| SCM4 | -0.004 | -0.003 | -0.003 | 0 |
| YLR194C | -0.004 | -0.009 | -0.007 | 0 |
| YGR066C | 0.002 | 0.002 | 0.002 | 0 |
| YLR211C | 0.001 | 0.001 | 0.001 | 0 |
| SLX9 | 0.007 | 0.013 | 0.010 | 0 |
| MET14 | 0.003 | -0.005 | -0.001 | 0 |
| YGR107W | 0.001 | 0.005 | 0.003 | 0 |
| SPT23 | 0.011 | 0.000 | 0.006 | 0 |
| PIN2 | -0.003 | 0.006 | 0.001 | 0 |
| NFU1 | 0.002 | 0.012 | 0.007 | 0 |
| GCY1 | -0.002 | 0.008 | 0.003 | 0 |
| OAR1 | 0.027 | 0.042 | 0.035 | 1 |
| EFT1 | -0.013 | -0.006 | -0.010 | 0 |
| YKL070W | 0.002 | 0.006 | 0.004 | 0 |
| PDR5 | -0.001 | -0.001 | -0.001 | 0 |
| HOT13 | -0.008 | -0.048 | -0.028 | 0 |

| | | | | |
|---|---|---|---|---|
| YKL100C | 0.001 | 0.005 | 0.003 | 0 |
| TUF1 | -0.081 | -0.194 | -0.138 | -1 |
| SBA1 | 0.002 | -0.005 | -0.001 | 0 |
| YOR200W | -0.028 | 0.004 | -0.012 | 0 |
| YKL131W | 0.011 | 0.008 | 0.010 | 0 |
| STE13 | -0.009 | -0.006 | -0.007 | 0 |
| YKL147C | 0.005 | 0.000 | 0.002 | 0 |
| KIN4 | 0.010 | 0.010 | 0.010 | 0 |
| YKL162C | -0.018 | -0.002 | -0.010 | 0 |
| SRL1 | 0.005 | -0.001 | 0.002 | 0 |
| LST4 | 0.003 | -0.007 | -0.002 | 0 |
| FSH3 | -0.001 | 0.006 | 0.003 | 0 |
| YLR169W | 0.010 | 0.016 | 0.013 | 0 |
| RPL26B | -0.015 | -0.009 | -0.012 | 0 |
| VTA1 | -0.027 | -0.016 | -0.022 | 0 |
| YGR051C | -0.008 | 0.007 | -0.001 | 0 |
| PBA1 | 0.014 | 0.014 | 0.014 | 0 |
| YGR067C | -0.003 | -0.006 | -0.004 | 0 |
| CRR1 | 0.003 | 0.005 | 0.004 | 0 |
| MRP13 | -0.002 | 0.009 | 0.004 | 0 |
| DID4 | -0.023 | -0.042 | -0.032 | -1 |
| CLB1 | 0.002 | 0.006 | 0.004 | 0 |
| YKL023W | -0.005 | -0.003 | -0.004 | 0 |
| YOR105W | 0.006 | 0.003 | 0.004 | 0 |
| VPS24 | -0.035 | -0.025 | -0.030 | -1 |
| YOR121C | 0.000 | -0.008 | -0.004 | 0 |
| TMA19 | 0.009 | 0.050 | 0.030 | 0 |
| BAG7 | 0.005 | 0.004 | 0.004 | 0 |
| YKL071W | -0.012 | -0.011 | -0.011 | 0 |
| SLP1 | -0.233 | -0.220 | -0.227 | -1 |
| MDH1 | -0.009 | -0.008 | -0.009 | 0 |
| LCB4 | 0.008 | 0.007 | 0.008 | 0 |
| HSL1 | 0.003 | 0.023 | 0.013 | 0 |
| MSB1 | 0.002 | -0.013 | -0.005 | 0 |
| YKL118W | -0.029 | -0.067 | -0.048 | -1 |
| MRM1 | -0.013 | 0.002 | -0.006 | 0 |
| RMA1 | 0.001 | 0.005 | 0.003 | 0 |
| YOR220W | -0.008 | 0.000 | -0.004 | 0 |
| SDH1 | -0.001 | -0.011 | -0.006 | 0 |
| RPL33B | 0.004 | -0.007 | -0.001 | 0 |
| PIR3 | -0.001 | -0.007 | -0.004 | 0 |
| YOR248W | 0.010 | -0.005 | 0.003 | 0 |
| YKL177W | -0.001 | 0.004 | 0.001 | 0 |
| YOR283W | 0.002 | 0.003 | 0.002 | 0 |
| APS1 | 0.005 | 0.009 | 0.007 | 0 |
| YGR035C | -0.005 | -0.002 | -0.003 | 0 |
| SWI6 | 0.024 | 0.070 | 0.047 | 1 |
| FMP48 | 0.009 | -0.003 | 0.003 | 0 |
| YKE2 | 0.048 | 0.069 | 0.059 | 1 |
| ART5 | 0.040 | 0.041 | 0.041 | 1 |
| FRE1 | 0.001 | 0.001 | 0.001 | 0 |
| RPL11B | 0.027 | 0.019 | 0.023 | 0 |
| MRP17 | 0.010 | 0.020 | 0.015 | 0 |

| | | | | |
|---|---|---|---|---|
| CLB6 | -0.008 | 0.002 | -0.003 | 0 |
| PAN3 | 0.006 | 0.020 | 0.013 | 0 |
| VAM3 | -0.033 | 0.006 | -0.013 | 0 |
| PHD1 | -0.005 | -0.001 | -0.003 | 0 |
| LEO1 | 0.019 | 0.026 | 0.023 | 0 |
| NUP120 | 0.029 | 0.039 | 0.034 | 1 |
| IRC14 | -0.003 | 0.000 | -0.001 | 0 |
| STB6 | -0.015 | -0.022 | -0.019 | 0 |
| ISN1 | -0.007 | -0.007 | -0.007 | 0 |
| SRX1 | 0.006 | 0.004 | 0.005 | 0 |
| YRM1 | -0.002 | -0.001 | -0.001 | 0 |
| YKL102C | -0.018 | -0.006 | -0.012 | 0 |
| IES4 | 0.000 | 0.004 | 0.002 | 0 |
| VPH2 | -0.039 | 0.002 | -0.019 | 0 |
| HIS3 | 0.010 | -0.005 | 0.002 | 0 |
| YKL133C | -0.015 | 0.001 | -0.007 | 0 |
| MCT1 | 0.005 | 0.005 | 0.005 | 0 |
| DBR1 | -0.003 | 0.009 | 0.003 | 0 |
| IRC13 | 0.008 | -0.001 | 0.004 | 0 |
| PIR1 | -0.003 | -0.012 | -0.007 | 0 |
| YOR251C | -0.024 | 0.000 | -0.012 | 0 |
| STE3 | -0.003 | 0.003 | 0.000 | 0 |
| HUA2 | 0.005 | 0.014 | 0.010 | 0 |
| YLR171W | 0.007 | 0.010 | 0.009 | 0 |
| CAX4 | 0.040 | 0.081 | 0.060 | 1 |
| TOS4 | 0.017 | 0.008 | 0.013 | 0 |
| YGR054W | -0.048 | -0.060 | -0.054 | -1 |
| COQ9 | -0.033 | 0.003 | -0.015 | 0 |
| YGR069W | -0.007 | -0.009 | -0.008 | 0 |
| CPR6 | -0.001 | 0.011 | 0.005 | 0 |
| PDC6 | 0.000 | 0.002 | 0.001 | 0 |
| RPL14A | 0.001 | 0.007 | 0.004 | 0 |
| YGR111W | -0.010 | -0.007 | -0.008 | 0 |
| GPX1 | -0.005 | 0.008 | 0.002 | 0 |
| RGS2 | 0.008 | 0.004 | 0.006 | 0 |
| YKL044W | 0.000 | 0.002 | 0.001 | 0 |
| UBP2 | 0.004 | -0.002 | 0.001 | 0 |
| YKL061W | -0.011 | -0.001 | -0.006 | 0 |
| IDH2 | -0.033 | 0.000 | -0.017 | 0 |
| LHS1 | -0.034 | -0.028 | -0.031 | -1 |
| NFI1 | -0.020 | -0.003 | -0.012 | 0 |
| CYT2 | 0.015 | 0.017 | 0.016 | 0 |
| DCS2 | 0.004 | 0.016 | 0.010 | 0 |
| LAP4 | -0.012 | -0.029 | -0.020 | 0 |
| SPR1 | -0.011 | -0.009 | -0.010 | 0 |
| OAC1 | -0.010 | 0.001 | -0.005 | 0 |
| YOR205C | 0.039 | 0.019 | 0.029 | 0 |
| 1-Oct | 0.010 | 0.001 | 0.005 | 0 |
| ODC2 | 0.006 | -0.002 | 0.002 | 0 |
| MCR1 | -0.006 | 0.006 | 0.000 | 0 |
| HES1 | -0.004 | 0.000 | -0.002 | 0 |
| TPK3 | 0.003 | -0.008 | -0.002 | 0 |
| TMA16 | 0.000 | -0.001 | 0.000 | 0 |

| | | | | |
|---|---|---|---|---|
| COY1 | -0.007 | 0.001 | -0.003 | 0 |
| YOR285W | 0.005 | 0.010 | 0.007 | 0 |
| DPH5 | -0.009 | -0.002 | -0.006 | 0 |
| ACB1 | -0.017 | -0.013 | -0.015 | 0 |
| YLR184W | 0.054 | 0.028 | 0.041 | 1 |
| MUP1 | 0.008 | 0.015 | 0.011 | 0 |
| YLR202C | 0.021 | 0.003 | 0.012 | 0 |
| ROM1 | -0.015 | -0.010 | -0.013 | 0 |
| YLR217W | -0.005 | 0.004 | 0.000 | 0 |
| CTT1 | 0.007 | -0.010 | -0.002 | 0 |
| CAP1 | 0.001 | 0.018 | 0.009 | 0 |
| SHY1 | -0.019 | 0.001 | -0.009 | 0 |
| YKL027W | 0.000 | 0.001 | 0.000 | 0 |
| LEU9 | -0.004 | 0.000 | -0.002 | 0 |
| DCW1 | 0.007 | 0.010 | 0.008 | 0 |
| CAT5 | 0.002 | -0.001 | 0.001 | 0 |
| MSN4 | -0.011 | -0.001 | -0.006 | 0 |
| SIA1 | 0.004 | 0.007 | 0.006 | 0 |
| MUD2 | 0.035 | 0.033 | 0.034 | 1 |
| PNS1 | -0.009 | -0.005 | -0.007 | 0 |
| CUE2 | -0.001 | 0.002 | 0.000 | 0 |
| ALE1 | -0.006 | -0.005 | -0.006 | 0 |
| YKL105C | -0.003 | 0.001 | -0.001 | 0 |
| ULS1 | 0.006 | 0.001 | 0.004 | 0 |
| DGR2 | -0.004 | -0.002 | -0.003 | 0 |
| PTP2 | -0.004 | 0.001 | -0.001 | 0 |
| APL2 | 0.009 | 0.001 | 0.005 | 0 |
| YOR223W | 0.000 | 0.000 | 0.000 | 0 |
| YKL151C | -0.003 | -0.006 | -0.004 | 0 |
| YOR238W | 0.000 | 0.003 | 0.001 | 0 |
| MRP49 | 0.016 | 0.014 | 0.015 | 0 |
| NAT5 | 0.003 | 0.001 | 0.002 | 0 |
| LOT5 | -0.005 | -0.010 | -0.007 | 0 |
| YOR286W | 0.013 | 0.015 | 0.014 | 0 |
| MPD1 | -0.032 | -0.028 | -0.030 | -1 |
| SEC72 | 0.004 | 0.021 | 0.013 | 0 |
| YJL206C | 0.007 | 0.012 | 0.009 | 0 |
| IMH1 | -0.022 | -0.026 | -0.024 | -1 |
| RPS14B | -0.002 | 0.006 | 0.002 | 0 |
| KGD2 | -0.019 | 0.000 | -0.010 | 0 |
| ATG27 | -0.003 | -0.012 | -0.008 | 0 |
| YDR161W | 0.020 | 0.013 | 0.017 | 0 |
| FMP33 | -0.011 | -0.007 | -0.009 | 0 |
| SAS4 | 0.013 | 0.003 | 0.008 | 0 |
| YJL147C | 0.001 | -0.006 | -0.003 | 0 |
| YDR199W | 0.011 | 0.016 | 0.013 | 0 |
| YJL131C | 0.000 | 0.004 | 0.002 | 0 |
| RAD9 | 0.012 | 0.000 | 0.006 | 0 |
| RPS25B | 0.008 | 0.013 | 0.010 | 0 |
| COX20 | -0.016 | -0.007 | -0.012 | 0 |
| BUD8 | -0.023 | -0.011 | -0.017 | 0 |
| YGL199C | -0.009 | -0.002 | -0.006 | 0 |
| ROM2 | 0.022 | 0.014 | 0.018 | 0 |

| | | | | |
|---|---|---|---|---|
| CLG1 | 0.031 | 0.013 | 0.022 | 0 |
| VAC14 | -0.024 | -0.008 | -0.016 | 0 |
| SHE10 | -0.036 | -0.026 | -0.031 | -1 |
| DUS4 | 0.004 | 0.004 | 0.004 | 0 |
| RTF1 | 0.029 | 0.040 | 0.034 | 1 |
| ECM22 | -0.009 | -0.013 | -0.011 | 0 |
| VEL1 | 0.004 | 0.004 | 0.004 | 0 |
| RCK2 | -0.008 | -0.003 | -0.005 | 0 |
| NMA2 | -0.005 | 0.010 | 0.003 | 0 |
| RED1 | -0.003 | -0.005 | -0.004 | 0 |
| MTL1 | 0.019 | 0.011 | 0.015 | 0 |
| YLR280C | -0.002 | 0.017 | 0.008 | 0 |
| RPL21B | -0.030 | -0.042 | -0.036 | -1 |
| YJL218W | -0.007 | -0.007 | -0.007 | 0 |
| YLR294C | 0.002 | -0.004 | -0.001 | 0 |
| YJL206C-A | 0.000 | -0.006 | -0.003 | 0 |
| YLR311C | -0.001 | -0.007 | -0.004 | 0 |
| RPS22A | 0.009 | 0.023 | 0.016 | 0 |
| YDR149C | -0.015 | 0.007 | -0.004 | 0 |
| SWI3 | 0.017 | 0.010 | 0.013 | 0 |
| NBP2 | 0.027 | 0.017 | 0.022 | 0 |
| HSP150 | -0.009 | -0.020 | -0.015 | 0 |
| PLP1 | 0.002 | 0.000 | 0.001 | 0 |
| IDS2 | -0.007 | 0.009 | 0.001 | 0 |
| VPS64 | -0.023 | -0.009 | -0.016 | 0 |
| URA2 | -0.022 | 0.008 | -0.007 | 0 |
| SPR28 | 0.022 | 0.007 | 0.015 | 0 |
| TAL1 | -0.010 | -0.011 | -0.011 | 0 |
| EMP24 | -0.049 | -0.046 | -0.048 | -1 |
| SUR4 | 0.036 | 0.033 | 0.035 | 1 |
| KIP3 | -0.029 | -0.019 | -0.024 | 0 |
| REH1 | -0.014 | 0.000 | -0.007 | 0 |
| SAP4 | -0.001 | 0.001 | 0.000 | 0 |
| YLR407W | -0.001 | 0.003 | 0.001 | 0 |
| RAI1 | 0.041 | 0.037 | 0.039 | 1 |
| BNA5 | -0.006 | -0.003 | -0.004 | 0 |
| YPS5 | -0.005 | -0.003 | -0.004 | 0 |
| SSP120 | 0.003 | 0.000 | 0.001 | 0 |
| YGR011W | -0.003 | 0.002 | 0.000 | 0 |
| RPS28B | -0.002 | -0.019 | -0.011 | 0 |
| YGR025W | 0.006 | 0.000 | 0.003 | 0 |
| PCL5 | 0.004 | 0.003 | 0.004 | 0 |
| REE1 | -0.009 | 0.002 | -0.004 | 0 |
| ATP14 | 0.026 | 0.016 | 0.021 | 0 |
| RCY1 | 0.034 | 0.034 | 0.034 | 1 |
| YLR312C | -0.013 | -0.005 | -0.009 | 0 |
| RPL39 | 0.032 | 0.012 | 0.022 | 0 |
| NUM1 | -0.005 | -0.014 | -0.010 | 0 |
| CPS1 | 0.009 | 0.013 | 0.011 | 0 |
| CWC15 | 0.016 | 0.012 | 0.014 | 0 |
| CIS3 | 0.009 | 0.002 | 0.005 | 0 |
| ATC1 | 0.000 | 0.008 | 0.004 | 0 |
| SFH5 | -0.003 | 0.005 | 0.001 | 0 |

| | | | | |
|---|---|---|---|---|
| YDR203W | -0.064 | -0.112 | -0.088 | -1 |
| TRK1 | -0.012 | -0.003 | -0.007 | 0 |
| MFB1 | 0.024 | 0.007 | 0.016 | 0 |
| NUP2 | -0.008 | 0.005 | -0.002 | 0 |
| RTN1 | 0.013 | 0.017 | 0.015 | 0 |
| YLR356W | -0.010 | 0.002 | -0.004 | 0 |
| ARO8 | 0.014 | 0.012 | 0.013 | 0 |
| VID22 | 0.059 | 0.066 | 0.063 | 1 |
| YGL217C | -0.022 | -0.028 | -0.025 | -1 |
| RPS29A | 0.001 | 0.010 | 0.005 | 0 |
| YGL230C | -0.005 | 0.002 | -0.001 | 0 |
| YLR408C | -0.017 | -0.009 | -0.013 | 0 |
| PDE1 | -0.012 | 0.003 | -0.005 | 0 |
| YLR232W | 0.008 | 0.002 | 0.005 | 0 |
| YGL260W | 0.003 | 0.011 | 0.007 | 0 |
| SYM1 | 0.012 | 0.008 | 0.010 | 0 |
| YGR012W | -0.004 | -0.004 | -0.004 | 0 |
| NEJ1 | -0.005 | -0.005 | -0.005 | 0 |
| YGR026W | 0.008 | -0.001 | 0.003 | 0 |
| YLR281C | -0.003 | 0.007 | 0.002 | 0 |
| ATP4 | -0.010 | 0.012 | 0.001 | 0 |
| YJL216C | -0.043 | -0.005 | -0.024 | 0 |
| YLR296W | -0.005 | -0.015 | -0.010 | 0 |
| ECM25 | -0.007 | -0.010 | -0.008 | 0 |
| MRPL15 | -0.016 | -0.001 | -0.008 | 0 |
| BUD19 | 0.033 | 0.028 | 0.030 | 1 |
| CTH1 | 0.002 | -0.001 | 0.001 | 0 |
| YJL171C | -0.010 | -0.004 | -0.007 | 0 |
| TRM82 | -0.003 | 0.007 | 0.002 | 0 |
| FAR1 | 0.004 | 0.004 | 0.004 | 0 |
| YDR185C | -0.002 | -0.003 | -0.003 | 0 |
| YJL144W | 0.002 | -0.002 | 0.000 | 0 |
| COQ4 | -0.029 | -0.010 | -0.019 | 0 |
| PEX30 | -0.008 | 0.008 | 0.000 | 0 |
| YDR220C | -0.002 | 0.000 | -0.001 | 0 |
| SPO77 | -0.001 | 0.013 | 0.006 | 0 |
| LYS4 | -0.073 | -0.070 | -0.071 | -1 |
| RSC2 | 0.036 | 0.043 | 0.039 | 1 |
| KEX1 | 0.004 | -0.002 | 0.001 | 0 |
| YLR374C | -0.017 | -0.014 | -0.016 | 0 |
| YGL218W | 0.019 | 0.032 | 0.026 | 0 |
| ECM19 | -0.010 | 0.004 | -0.003 | 0 |
| YGL231C | 0.022 | 0.019 | 0.021 | 0 |
| VIP1 | 0.034 | 0.031 | 0.033 | 1 |
| ZIP2 | -0.011 | 0.001 | -0.005 | 0 |
| EST1 | 0.001 | 0.008 | 0.005 | 0 |
| YGL261C | -0.005 | -0.005 | -0.005 | 0 |
| YLR252W | 0.000 | 0.002 | 0.001 | 0 |
| MSB2 | 0.007 | 0.008 | 0.008 | 0 |
| PDR8 | 0.007 | 0.012 | 0.009 | 0 |
| GLR1 | -0.018 | -0.003 | -0.010 | 0 |
| YLR282C | 0.003 | 0.003 | 0.003 | 0 |
| YPL077C | -0.006 | -0.008 | -0.007 | 0 |

| | | | | |
|---|---|---|---|---|
| YJL215C | -0.001 | -0.005 | -0.003 | 0 |
| YLR297W | 0.000 | -0.006 | -0.003 | 0 |
| ACO2 | 0.006 | 0.006 | 0.006 | 0 |
| SPH1 | -0.009 | 0.000 | -0.005 | 0 |
| SWE1 | 0.004 | 0.005 | 0.005 | 0 |
| GIR2 | 0.004 | 0.003 | 0.003 | 0 |
| ASG7 | -0.006 | -0.029 | -0.017 | 0 |
| STB3 | -0.001 | 0.001 | 0.000 | 0 |
| FBP26 | -0.016 | -0.014 | -0.015 | 0 |
| YDR186C | 0.010 | 0.001 | 0.005 | 0 |
| IRC9 | 0.009 | 0.001 | 0.005 | 0 |
| EBS1 | 0.010 | 0.010 | 0.010 | 0 |
| RPL38 | -0.002 | -0.012 | -0.007 | 0 |
| GTB1 | 0.005 | 0.009 | 0.007 | 0 |
| FKS1 | -0.019 | -0.006 | -0.012 | 0 |
| MRPL7 | 0.036 | 0.027 | 0.032 | 1 |
| VPS38 | -0.035 | -0.037 | -0.036 | -1 |
| POX1 | -0.004 | -0.003 | -0.004 | 0 |
| STP3 | -0.019 | -0.007 | -0.013 | 0 |
| FRA2 | -0.015 | -0.015 | -0.015 | 0 |
| YLR392C | -0.004 | 0.000 | -0.002 | 0 |
| TAN1 | -0.035 | -0.044 | -0.039 | -1 |
| BER1 | -0.004 | -0.003 | -0.004 | 0 |
| RMR1 | 0.007 | 0.004 | 0.006 | 0 |
| TOP3 | 0.001 | 0.030 | 0.015 | 0 |
| YGL262W | -0.001 | -0.003 | -0.002 | 0 |
| YLR253W | 0.010 | 0.000 | 0.005 | 0 |
| YGR015C | 0.016 | 0.014 | 0.015 | 0 |
| BOP2 | 0.008 | 0.004 | 0.006 | 0 |
| RPS6A | -0.035 | -0.035 | -0.035 | -1 |
| YLR283W | 0.001 | 0.005 | 0.003 | 0 |
| YTA6 | 0.009 | -0.001 | 0.004 | 0 |
| HXT8 | -0.003 | -0.019 | -0.011 | 0 |
| ECM38 | -0.010 | -0.007 | -0.008 | 0 |
| MBB1 | -0.010 | -0.010 | -0.010 | 0 |
| NKP2 | -0.023 | 0.001 | -0.011 | 0 |
| MNN5 | -0.001 | 0.003 | 0.001 | 0 |
| ENT5 | -0.003 | 0.006 | 0.001 | 0 |
| YJL169W | -0.003 | 0.007 | 0.002 | 0 |
| HSP42 | 0.003 | 0.000 | 0.002 | 0 |
| VPS35 | -0.051 | -0.034 | -0.042 | -1 |
| HST4 | 0.004 | 0.000 | 0.002 | 0 |
| RPB4 | 0.018 | 0.016 | 0.017 | 0 |
| UME6 | -0.008 | -0.015 | -0.012 | 0 |
| YLR326W | -0.002 | -0.003 | -0.002 | 0 |
| YDR222W | -0.006 | 0.009 | 0.001 | 0 |
| RPL26A | -0.027 | -0.017 | -0.022 | 0 |
| YDR239C | -0.006 | 0.002 | -0.002 | 0 |
| STE11 | -0.001 | 0.008 | 0.004 | 0 |
| SIP2 | -0.005 | -0.007 | -0.006 | 0 |
| PSY3 | -0.018 | -0.006 | -0.012 | 0 |
| NIF3 | -0.001 | 0.010 | 0.005 | 0 |
| ATP10 | 0.012 | 0.023 | 0.017 | 0 |

| | | | | |
|---|---|---|---|---|
| ADE5,7 | -0.017 | 0.002 | -0.008 | 0 |
| YLR413W | -0.002 | -0.006 | -0.004 | 0 |
| HFM1 | 0.006 | -0.002 | 0.002 | 0 |
| YLR235C | 0.011 | 0.020 | 0.015 | 0 |
| COS12 | 0.005 | 0.003 | 0.004 | 0 |
| NDL1 | 0.000 | 0.011 | 0.006 | 0 |
| YGR016W | -0.009 | -0.003 | -0.006 | 0 |
| SEC22 | -0.006 | -0.011 | -0.009 | 0 |
| RLM1 | -0.002 | 0.009 | 0.003 | 0 |
| ECI1 | 0.007 | 0.003 | 0.005 | 0 |
| UBP16 | 0.007 | 0.002 | 0.004 | 0 |
| OPT1 | 0.001 | -0.003 | -0.001 | 0 |
| EXG1 | -0.012 | -0.001 | -0.006 | 0 |
| PHO90 | -0.012 | -0.009 | -0.011 | 0 |
| EST2 | 0.011 | 0.017 | 0.014 | 0 |
| YJL185C | -0.001 | 0.001 | 0.000 | 0 |
| YDR154C | 0.001 | 0.010 | 0.005 | 0 |
| SET2 | -0.008 | 0.012 | 0.002 | 0 |
| ARG82 | -0.004 | -0.005 | -0.004 | 0 |
| INO1 | 0.008 | 0.000 | 0.004 | 0 |
| NUP42 | 0.008 | 0.008 | 0.008 | 0 |
| YUR1 | 0.009 | 0.014 | 0.011 | 0 |
| YDR209C | -0.004 | 0.002 | -0.001 | 0 |
| TMA10 | 0.011 | 0.005 | 0.008 | 0 |
| CRF1 | -0.013 | 0.001 | -0.006 | 0 |
| YLR345W | 0.003 | -0.008 | -0.002 | 0 |
| BUD26 | 0.018 | 0.024 | 0.021 | 0 |
| NMD4 | 0.010 | 0.003 | 0.007 | 0 |
| MIG2 | -0.011 | -0.004 | -0.008 | 0 |
| FBP1 | -0.008 | -0.007 | -0.007 | 0 |
| EDC1 | -0.001 | 0.011 | 0.005 | 0 |
| COX8 | 0.004 | 0.008 | 0.006 | 0 |
| YGL235W | -0.001 | 0.004 | 0.001 | 0 |
| YLR414C | -0.009 | 0.004 | -0.003 | 0 |
| RTG2 | -0.069 | -0.010 | -0.039 | 0 |
| YLR236C | 0.000 | -0.010 | -0.005 | 0 |
| YGR001C | -0.003 | -0.005 | -0.004 | 0 |
| YLR255C | -0.001 | 0.008 | 0.003 | 0 |
| YGR017W | -0.018 | -0.001 | -0.010 | 0 |
| YLR269C | -0.006 | -0.010 | -0.008 | 0 |
| YPL088W | 0.002 | -0.001 | 0.000 | 0 |
| NNT1 | 0.008 | -0.002 | 0.003 | 0 |
| YPL073C | 0.014 | 0.010 | 0.012 | 0 |
| PEX2 | -0.001 | -0.005 | -0.003 | 0 |
| MET17 | -0.008 | -0.004 | -0.006 | 0 |
| UBP12 | -0.020 | -0.019 | -0.020 | 0 |
| BUD6 | 0.001 | 0.000 | 0.001 | 0 |
| MNN11 | -0.039 | -0.005 | -0.022 | 0 |
| CPR1 | 0.013 | 0.004 | 0.009 | 0 |
| QCR8 | 0.004 | 0.004 | 0.004 | 0 |
| RSM24 | 0.016 | 0.026 | 0.021 | 0 |
| YJL152W | 0.005 | 0.004 | 0.004 | 0 |
| YDR193W | 0.008 | 0.010 | 0.009 | 0 |

| | | | | |
|---|---|---|---|---|
| TIF2 | -0.010 | -0.003 | -0.006 | 0 |
| YDR210W | -0.011 | -0.004 | -0.008 | 0 |
| NMA1 | 0.012 | 0.002 | 0.007 | 0 |
| HTA1 | 0.013 | 0.015 | 0.014 | 0 |
| DIC1 | 0.002 | 0.008 | 0.005 | 0 |
| HOS2 | -0.012 | -0.011 | -0.011 | 0 |
| GRX8 | -0.005 | -0.011 | -0.008 | 0 |
| YPT32 | -0.018 | -0.003 | -0.010 | 0 |
| CSR1 | -0.012 | -0.007 | -0.009 | 0 |
| SDT1 | -0.006 | 0.002 | -0.002 | 0 |
| SKI2 | 0.000 | 0.005 | 0.003 | 0 |
| MTO1 | 0.004 | -0.003 | 0.001 | 0 |
| YLR415C | -0.002 | 0.002 | 0.000 | 0 |
| HXK2 | -0.058 | -0.041 | -0.050 | -1 |
| FAR10 | 0.008 | 0.008 | 0.008 | 0 |
| CUL3 | -0.008 | 0.002 | -0.003 | 0 |
| YLR257W | -0.011 | -0.001 | -0.006 | 0 |
| YGR018C | -0.007 | -0.005 | -0.006 | 0 |
| DCS1 | 0.000 | 0.007 | 0.004 | 0 |
| YDC1 | 0.005 | 0.006 | 0.005 | 0 |
| YLR287C | 0.009 | 0.010 | 0.009 | 0 |
| YPL071C | -0.003 | 0.003 | 0.000 | 0 |
| YJL211C | -0.010 | -0.003 | -0.007 | 0 |
| ACO1 | -0.017 | -0.009 | -0.013 | 0 |
| PAN5 | -0.003 | -0.010 | -0.006 | 0 |
| MMS22 | 0.030 | 0.070 | 0.050 | 1 |
| YJL181W | -0.006 | 0.008 | 0.001 | 0 |
| RPA14 | 0.022 | 0.006 | 0.014 | 0 |
| HAL5 | 0.008 | 0.007 | 0.007 | 0 |
| NGG1 | 0.042 | 0.087 | 0.065 | 1 |
| SNA3 | 0.003 | 0.001 | 0.002 | 0 |
| MSS116 | -0.034 | -0.009 | -0.022 | 0 |
| YJL135W | 0.007 | 0.006 | 0.006 | 0 |
| UPC2 | -0.009 | 0.006 | -0.001 | 0 |
| REC102 | 0.002 | 0.010 | 0.006 | 0 |
| ADK1 | -0.055 | -0.042 | -0.048 | -1 |
| YLR349W | 0.009 | 0.000 | 0.005 | 0 |
| GCN1 | 0.004 | -0.011 | -0.003 | 0 |
| YLR365W | -0.006 | 0.004 | -0.001 | 0 |
| NCS6 | 0.018 | 0.030 | 0.024 | 0 |
| CTF3 | -0.004 | -0.005 | -0.005 | 0 |
| OST5 | 0.000 | 0.014 | 0.007 | 0 |
| YLR400W | 0.003 | 0.003 | 0.003 | 0 |
| HAP2 | -0.001 | -0.004 | -0.002 | 0 |
| YLR416C | -0.004 | -0.012 | -0.008 | 0 |
| FZF1 | 0.006 | 0.007 | 0.006 | 0 |
| LIP2 | -0.009 | -0.022 | -0.015 | 0 |
| PEX31 | -0.001 | 0.004 | 0.001 | 0 |
| GSY2 | 0.007 | -0.013 | -0.003 | 0 |
| UGA1 | -0.003 | 0.004 | 0.000 | 0 |
| YLR271W | -0.004 | -0.005 | -0.005 | 0 |
| ELP3 | 0.050 | 0.052 | 0.051 | 1 |
| YLR287-A | 0.002 | -0.023 | -0.010 | 0 |

| | | | | |
|---|---|---|---|---|
| MUK1 | 0.003 | -0.006 | -0.001 | 0 |
| CBP1 | -0.031 | -0.020 | -0.025 | 0 |
| UBC12 | 0.008 | 0.002 | 0.005 | 0 |
| ELO1 | 0.006 | -0.004 | 0.001 | 0 |
| VPS65 | -0.025 | -0.003 | -0.014 | 0 |
| YJL182C | -0.006 | 0.005 | 0.000 | 0 |
| YDR157W | 0.002 | 0.005 | 0.003 | 0 |
| TPK1 | 0.000 | 0.004 | 0.002 | 0 |
| SDH4 | 0.005 | -0.001 | 0.002 | 0 |
| YJL150W | -0.004 | 0.009 | 0.003 | 0 |
| REF2 | -0.028 | -0.009 | -0.018 | 0 |
| LCB3 | 0.019 | 0.019 | 0.019 | 0 |
| AHA1 | -0.006 | -0.001 | -0.004 | 0 |
| CHS5 | 0.009 | 0.020 | 0.014 | 0 |
| SIR4 | -0.033 | -0.011 | -0.022 | 0 |
| ORM2 | -0.012 | -0.004 | -0.008 | 0 |
| DSD1 | -0.002 | 0.001 | -0.001 | 0 |
| YLR366W | -0.003 | 0.004 | 0.000 | 0 |
| VAM7 | -0.012 | 0.003 | -0.005 | 0 |
| MTC3 | 0.015 | 0.013 | 0.014 | 0 |
| DUS3 | 0.001 | 0.000 | 0.000 | 0 |
| KAP114 | 0.007 | 0.008 | 0.007 | 0 |
| VPS36 | -0.031 | -0.027 | -0.029 | -1 |
| ZRT1 | 0.008 | 0.000 | 0.004 | 0 |
| YLR241W | -0.004 | -0.004 | -0.004 | 0 |
| PRP18 | -0.030 | -0.009 | -0.019 | 0 |
| LCB5 | -0.010 | -0.005 | -0.008 | 0 |
| VMA7 | 0.017 | 0.020 | 0.018 | 0 |
| PIG1 | -0.001 | 0.011 | 0.005 | 0 |
| BRO1 | -0.022 | -0.173 | -0.097 | -1 |
| MEC3 | 0.002 | 0.005 | 0.004 | 0 |
| BTS1 | -0.003 | 0.008 | 0.002 | 0 |
| NUC1 | -0.009 | -0.011 | -0.010 | 0 |
| CDA1 | -0.006 | -0.007 | -0.006 | 0 |
| YJL193W | 0.001 | -0.007 | -0.003 | 0 |
| ATP12 | -0.013 | -0.004 | -0.009 | 0 |
| HOM2 | -0.003 | 0.008 | 0.003 | 0 |
| YJL163C | -0.014 | 0.003 | -0.005 | 0 |
| CSN9 | 0.002 | -0.001 | 0.001 | 0 |
| YJL149W | -0.001 | 0.004 | 0.002 | 0 |
| CBS2 | 0.005 | 0.005 | 0.005 | 0 |
| MRS3 | 0.003 | 0.000 | 0.002 | 0 |
| YDR215C | -0.001 | -0.010 | -0.005 | 0 |
| JIP3 | -0.007 | -0.016 | -0.011 | 0 |
| IVY1 | 0.013 | -0.001 | 0.006 | 0 |
| NIT3 | -0.013 | 0.003 | -0.005 | 0 |
| MDS3 | -0.017 | -0.022 | -0.019 | 0 |
| RPS22B | -0.033 | 0.005 | -0.014 | 0 |
| SKI8 | -0.024 | -0.023 | -0.023 | -1 |
| IKI3 | 0.057 | 0.052 | 0.055 | 1 |
| VID30 | 0.007 | 0.015 | 0.011 | 0 |
| YLR402W | 0.035 | 0.026 | 0.031 | 1 |
| YGL242C | -0.003 | 0.005 | 0.001 | 0 |

| | | | | |
|---|---|---|---|---|
| CDC73 | -0.022 | -0.005 | -0.014 | 0 |
| ADH4 | -0.006 | -0.001 | -0.003 | 0 |
| ARV1 | 0.036 | 0.049 | 0.042 | 1 |
| MUQ1 | -0.001 | -0.022 | -0.011 | 0 |
| VPS63 | -0.016 | -0.016 | -0.016 | 0 |
| YGR021W | 0.006 | 0.010 | 0.008 | 0 |
| YLR278C | 0.018 | 0.011 | 0.015 | 0 |
| RPS9A | 0.009 | 0.000 | 0.004 | 0 |
| GUF1 | 0.000 | 0.004 | 0.002 | 0 |
| YPL068C | 0.001 | 0.003 | 0.002 | 0 |
| LAA1 | -0.005 | -0.002 | -0.003 | 0 |
| CDA2 | 0.002 | 0.002 | 0.002 | 0 |
| SOP4 | -0.006 | -0.002 | -0.004 | 0 |
| EKI1 | -0.012 | 0.006 | -0.003 | 0 |
| PFD1 | -0.008 | 0.000 | -0.004 | 0 |
| SAC3 | 0.048 | 0.050 | 0.049 | 1 |
| JJJ2 | -0.008 | 0.000 | -0.004 | 0 |
| YDR179W-A | -0.025 | -0.008 | -0.016 | 0 |
| RPA34 | 0.007 | 0.007 | 0.007 | 0 |
| RKM2 | -0.003 | 0.003 | 0.000 | 0 |
| YJL132W | 0.002 | 0.003 | 0.002 | 0 |
| ADR1 | 0.004 | -0.015 | -0.005 | 0 |
| MID2 | -0.028 | -0.020 | -0.024 | -1 |
| YDR230W | 0.015 | 0.017 | 0.016 | 0 |
| YLR352W | -0.006 | 0.000 | -0.003 | 0 |
| YIP4 | -0.012 | 0.001 | -0.006 | 0 |
| MDM30 | -0.038 | -0.031 | -0.034 | -1 |
| YGL214W | 0.007 | 0.003 | 0.005 | 0 |
| SWC7 | 0.001 | -0.012 | -0.005 | 0 |
| FLD1 | -0.018 | -0.011 | -0.014 | 0 |
| TAD1 | 0.004 | -0.001 | 0.001 | 0 |
| RPN13 | 0.028 | 0.024 | 0.026 | 1 |
| MNT2 | -0.016 | -0.001 | -0.008 | 0 |
| IRC20 | 0.004 | 0.000 | 0.002 | 0 |
| STF2 | -0.002 | 0.005 | 0.001 | 0 |
| YPT6 | -0.049 | -0.085 | -0.067 | -1 |
| YGR022C | 0.007 | 0.002 | 0.005 | 0 |
| YLR279W | 0.019 | 0.009 | 0.014 | 0 |
| YPL080C | 0.014 | 0.009 | 0.012 | 0 |
| YLR290C | 0.006 | 0.004 | 0.005 | 0 |
| YPL067C | -0.007 | -0.001 | -0.004 | 0 |
| YPL066W | 0.006 | -0.010 | -0.002 | 0 |
| FRE7 | 0.003 | 0.007 | 0.005 | 0 |
| OAZ1 | -0.010 | -0.033 | -0.022 | 0 |
| FUS3 | 0.004 | 0.037 | 0.020 | 0 |
| APL3 | -0.002 | 0.000 | -0.001 | 0 |
| CTF19 | 0.005 | 0.021 | 0.013 | 0 |
| PIN4 | -0.152 | -0.153 | -0.152 | -1 |
| PDH1 | 0.004 | 0.003 | 0.003 | 0 |
| KIP1 | 0.000 | -0.013 | -0.007 | 0 |
| CLB5 | 0.038 | 0.011 | 0.025 | 0 |
| NUP170 | 0.007 | 0.010 | 0.009 | 0 |
| RPS23B | 0.016 | 0.026 | 0.021 | 0 |

| | | | | |
|---|---|---|---|---|
| YPL158C | 0.003 | -0.007 | -0.002 | 0 |
| YPR150W | 0.002 | 0.004 | 0.003 | 0 |
| YGL015C | -0.003 | -0.010 | -0.006 | 0 |
| MMS1 | 0.035 | 0.000 | 0.017 | 0 |
| SKI3 | 0.011 | -0.014 | -0.002 | 0 |
| RIM8 | -0.037 | -0.068 | -0.052 | -1 |
| YCR090C | -0.007 | -0.001 | -0.004 | 0 |
| PYC1 | -0.004 | -0.018 | -0.011 | 0 |
| STF1 | 0.010 | -0.003 | 0.004 | 0 |
| FMP37 | 0.008 | -0.028 | -0.010 | 0 |
| YER188W | 0.003 | -0.011 | -0.004 | 0 |
| ATG2 | 0.000 | 0.009 | 0.004 | 0 |
| DUG1 | 0.000 | -0.004 | -0.002 | 0 |
| YNL228W | 0.003 | -0.010 | -0.003 | 0 |
| TOS2 | -0.002 | -0.006 | -0.004 | 0 |
| YNL211C | -0.011 | 0.018 | 0.004 | 0 |
| YGR235C | -0.006 | 0.004 | -0.001 | 0 |
| SLZ1 | -0.005 | -0.006 | -0.005 | 0 |
| VPS28 | -0.017 | -0.035 | -0.026 | 0 |
| HTA2 | -0.002 | 0.010 | 0.004 | 0 |
| ARL3 | -0.077 | -0.026 | -0.051 | -1 |
| PEP1 | -0.023 | -0.033 | -0.028 | -1 |
| YPL035C | -0.009 | 0.014 | 0.003 | 0 |
| MRPL16 | 0.001 | 0.021 | 0.011 | 0 |
| HST2 | 0.004 | 0.003 | 0.003 | 0 |
| SAS3 | -0.009 | 0.008 | -0.001 | 0 |
| YPR003C | -0.002 | -0.006 | -0.004 | 0 |
| PRX1 | 0.000 | -0.008 | -0.004 | 0 |
| THI22 | 0.002 | -0.007 | -0.003 | 0 |
| PET112 | -0.009 | 0.003 | -0.003 | 0 |
| MSS18 | 0.027 | 0.038 | 0.033 | 1 |
| YBL094C | 0.006 | 0.008 | 0.007 | 0 |
| MRP2 | -0.012 | -0.030 | -0.021 | 0 |
| RPL24A | 0.019 | 0.021 | 0.020 | 0 |
| QCR2 | 0.027 | 0.013 | 0.020 | 0 |
| YGL046W | -0.038 | -0.036 | -0.037 | -1 |
| YNG2 | -0.005 | 0.008 | 0.002 | 0 |
| PUS2 | 0.001 | -0.014 | -0.007 | 0 |
| SEM1 | 0.075 | 0.071 | 0.073 | 1 |
| YGL081W | -0.009 | -0.011 | -0.010 | 0 |
| RPL22B | 0.006 | 0.009 | 0.007 | 0 |
| ZWF1 | 0.023 | 0.017 | 0.020 | 0 |
| YFR045W | 0.002 | 0.000 | 0.001 | 0 |
| YNL226W | 0.022 | 0.025 | 0.023 | 1 |
| HSV2 | -0.003 | -0.005 | -0.004 | 0 |
| YNL208W | 0.011 | 0.017 | 0.014 | 0 |
| SFB3 | 0.017 | 0.012 | 0.015 | 0 |
| CWC27 | -0.010 | -0.010 | -0.010 | 0 |
| PDR3 | -0.012 | -0.011 | -0.012 | 0 |
| DIG1 | 0.006 | -0.011 | -0.002 | 0 |
| APN2 | -0.005 | -0.006 | -0.005 | 0 |
| SRL4 | -0.016 | 0.006 | -0.005 | 0 |
| YPL014W | 0.001 | 0.017 | 0.009 | 0 |

| | | | | |
|---|---|---|---|---|
| YBL053W | -0.006 | -0.008 | -0.007 | 0 |
| YPR004C | -0.006 | 0.007 | 0.000 | 0 |
| YBL065W | -0.055 | -0.047 | -0.051 | -1 |
| AXL1 | 0.009 | -0.012 | -0.001 | 0 |
| YBL081W | 0.025 | 0.033 | 0.029 | 1 |
| CTF4 | 0.046 | 0.016 | 0.031 | 0 |
| ERP6 | -0.001 | 0.001 | 0.000 | 0 |
| URN1 | 0.011 | 0.011 | 0.011 | 0 |
| KAP122 | 0.032 | 0.044 | 0.038 | 1 |
| MET16 | 0.013 | 0.003 | 0.008 | 0 |
| AGA2 | -0.003 | -0.016 | -0.009 | 0 |
| AQY1 | -0.004 | -0.025 | -0.014 | 0 |
| TIF4632 | -0.009 | -0.009 | -0.009 | 0 |
| KIN82 | -0.010 | -0.002 | -0.006 | 0 |
| MRH4 | 0.007 | 0.017 | 0.012 | 0 |
| SNA2 | 0.013 | 0.015 | 0.014 | 0 |
| YGL082W | -0.006 | -0.006 | -0.006 | 0 |
| YFR032C | 0.005 | -0.002 | 0.001 | 0 |
| LAP3 | 0.009 | -0.004 | 0.003 | 0 |
| CNN1 | 0.010 | -0.016 | -0.003 | 0 |
| JJJ1 | -0.007 | -0.033 | -0.020 | 0 |
| AZR1 | -0.035 | 0.003 | -0.016 | 0 |
| RTT106 | -0.017 | -0.020 | -0.018 | 0 |
| SPG1 | 0.009 | 0.023 | 0.016 | 0 |
| YNL195C | -0.008 | -0.021 | -0.014 | 0 |
| YPL062W | 0.049 | 0.035 | 0.042 | 1 |
| LDB7 | 0.330 | 0.044 | 0.187 | 1 |
| CAM1 | 0.003 | -0.008 | -0.003 | 0 |
| HAP3 | -0.002 | -0.004 | -0.003 | 0 |
| SVL3 | -0.016 | -0.007 | -0.012 | 0 |
| URA7 | 0.024 | 0.014 | 0.019 | 0 |
| MRPS16 | -0.001 | 0.014 | 0.006 | 0 |
| TOD6 | -0.003 | 0.003 | 0.000 | 0 |
| HAL1 | -0.014 | -0.006 | -0.010 | 0 |
| SEF1 | -0.070 | -0.047 | -0.058 | -1 |
| YPR123C | -0.025 | -0.043 | -0.034 | -1 |
| ALG3 | -0.017 | -0.006 | -0.011 | 0 |
| MEP3 | 0.005 | 0.012 | 0.008 | 0 |
| CDH1 | -0.005 | -0.047 | -0.026 | 0 |
| YPR153W | -0.015 | -0.041 | -0.028 | 0 |
| ATE1 | -0.008 | -0.002 | -0.005 | 0 |
| YPR170C | -0.003 | 0.006 | 0.002 | 0 |
| HOP2 | -0.018 | -0.031 | -0.025 | 0 |
| HPA2 | -0.004 | -0.016 | -0.010 | 0 |
| TYW3 | 0.003 | -0.014 | -0.006 | 0 |
| MSH3 | -0.007 | 0.001 | -0.003 | 0 |
| YDR535C | 0.008 | -0.002 | 0.003 | 0 |
| SCY1 | 0.001 | 0.007 | 0.004 | 0 |
| RPL29 | -0.008 | -0.007 | -0.007 | 0 |
| KEX2 | 0.021 | 0.016 | 0.019 | 0 |
| BNA6 | -0.001 | -0.006 | -0.004 | 0 |
| SQS1 | -0.009 | -0.015 | -0.012 | 0 |
| AMA1 | 0.001 | -0.003 | -0.001 | 0 |

| | | | | |
|---|---|---|---|---|
| YNL205C | 0.012 | 0.008 | 0.010 | 0 |
| YGR237C | -0.004 | 0.000 | -0.002 | 0 |
| YNL194C | 0.000 | 0.003 | 0.001 | 0 |
| ALD6 | 0.039 | -0.012 | 0.013 | 0 |
| SLA1 | -0.018 | -0.019 | -0.019 | 0 |
| SGF11 | 0.017 | 0.004 | 0.011 | 0 |
| PIM1 | 0.000 | 0.029 | 0.015 | 0 |
| TRM44 | -0.001 | 0.005 | 0.002 | 0 |
| FUI1 | -0.011 | 0.006 | -0.002 | 0 |
| YPL009C | 0.020 | 0.005 | 0.013 | 0 |
| YBL055C | -0.010 | -0.008 | -0.009 | 0 |
| ISR1 | 0.002 | -0.004 | -0.001 | 0 |
| UBP13 | -0.039 | -0.061 | -0.050 | -1 |
| CTR1 | -0.002 | 0.005 | 0.001 | 0 |
| YBL083C | -0.019 | 0.002 | -0.009 | 0 |
| VPS66 | -0.003 | 0.000 | -0.001 | 0 |
| RPN14 | 0.010 | 0.005 | 0.008 | 0 |
| PIN3 | 0.011 | 0.006 | 0.009 | 0 |
| CKB1 | 0.006 | 0.016 | 0.011 | 0 |
| BSP1 | 0.002 | 0.003 | 0.002 | 0 |
| YGL034C | -0.019 | -0.037 | -0.028 | 0 |
| OPT2 | -0.009 | -0.030 | -0.019 | 0 |
| MST27 | 0.011 | -0.014 | -0.001 | 0 |
| CDC50 | 0.038 | 0.039 | 0.039 | 1 |
| SGF73 | 0.022 | 0.024 | 0.023 | 1 |
| STL1 | 0.009 | -0.017 | -0.004 | 0 |
| GUP1 | 0.026 | 0.046 | 0.036 | 1 |
| QCR6 | 0.006 | 0.031 | 0.019 | 0 |
| YTP1 | 0.016 | 0.000 | 0.008 | 0 |
| RMD8 | -0.001 | -0.010 | -0.005 | 0 |
| ATG4 | -0.004 | 0.004 | 0.000 | 0 |
| YGR226C | 0.010 | 0.015 | 0.013 | 0 |
| SPS18 | 0.010 | 0.007 | 0.009 | 0 |
| PFK1 | -0.028 | -0.002 | -0.015 | 0 |
| YNL193W | -0.021 | -0.010 | -0.015 | 0 |
| LPE10 | 0.008 | 0.006 | 0.007 | 0 |
| HIR1 | -0.012 | -0.005 | -0.009 | 0 |
| ELC1 | -0.001 | -0.031 | -0.016 | 0 |
| NCL1 | 0.007 | 0.016 | 0.011 | 0 |
| SUV3 | -0.074 | 0.022 | -0.026 | 0 |
| ECM13 | -0.006 | 0.001 | -0.003 | 0 |
| CHL1 | 0.009 | 0.000 | 0.005 | 0 |
| PTC3 | -0.011 | -0.008 | -0.009 | 0 |
| YPR109W | 0.000 | 0.007 | 0.004 | 0 |
| PRS4 | 0.020 | 0.031 | 0.025 | 0 |
| YLH47 | -0.004 | 0.000 | -0.002 | 0 |
| BOI1 | -0.002 | 0.007 | 0.002 | 0 |
| TAZ1 | -0.005 | -0.019 | -0.012 | 0 |
| COG7 | -0.004 | -0.026 | -0.015 | 0 |
| NCA2 | 0.013 | 0.006 | 0.009 | 0 |
| ALK1 | -0.004 | 0.003 | -0.001 | 0 |
| YPR172W | -0.015 | -0.034 | -0.024 | 0 |
| MIG1 | -0.019 | -0.037 | -0.028 | 0 |

| | | | | |
|---|---|---|---|---|
| YPR195C | 0.000 | -0.025 | -0.012 | 0 |
| PRM8 | 0.006 | 0.008 | 0.007 | 0 |
| GIT1 | -0.006 | -0.003 | -0.004 | 0 |
| NPY1 | -0.008 | 0.005 | -0.001 | 0 |
| PAD1 | 0.000 | 0.000 | 0.000 | 0 |
| YGL085W | -0.010 | -0.015 | -0.013 | 0 |
| PHO4 | -0.006 | -0.011 | -0.009 | 0 |
| SIN4 | 0.011 | 0.006 | 0.008 | 0 |
| YMR31 | 0.007 | 0.003 | 0.005 | 0 |
| ALG9 | 0.004 | -0.002 | 0.001 | 0 |
| DIE2 | 0.006 | 0.008 | 0.007 | 0 |
| SPS19 | 0.008 | 0.003 | 0.005 | 0 |
| YAP1802 | 0.011 | 0.012 | 0.011 | 0 |
| CHS1 | -0.033 | -0.062 | -0.048 | -1 |
| PDR12 | -0.008 | -0.011 | -0.009 | 0 |
| ALK2 | -0.002 | 0.019 | 0.009 | 0 |
| SSN3 | 0.017 | 0.038 | 0.028 | 0 |
| RPL19B | -0.007 | -0.019 | -0.013 | 0 |
| SKS1 | -0.001 | 0.012 | 0.005 | 0 |
| YBL044W | -0.008 | -0.015 | -0.011 | 0 |
| NCR1 | -0.003 | 0.014 | 0.005 | 0 |
| PTH2 | 0.001 | -0.002 | -0.001 | 0 |
| DBF20 | -0.004 | -0.009 | -0.007 | 0 |
| AST1 | 0.001 | -0.010 | -0.005 | 0 |
| YPR126C | -0.001 | -0.002 | -0.001 | 0 |
| YBL086C | -0.008 | 0.004 | -0.002 | 0 |
| KAR3 | 0.005 | 0.000 | 0.003 | 0 |
| PMC1 | 0.005 | -0.002 | 0.001 | 0 |
| TPO3 | 0.005 | 0.003 | 0.004 | 0 |
| PIB2 | -0.041 | -0.007 | -0.024 | 0 |
| VPS4 | 0.008 | -0.019 | -0.005 | 0 |
| YGL036W | -0.007 | -0.007 | -0.007 | 0 |
| YPR196W | -0.015 | -0.011 | -0.013 | 0 |
| ERV14 | 0.022 | 0.029 | 0.025 | 1 |
| YCR099C | 0.000 | -0.002 | -0.001 | 0 |
| AFT1 | -0.002 | 0.018 | 0.008 | 0 |
| YDR539W | 0.003 | -0.012 | -0.004 | 0 |
| MAD1 | -0.002 | -0.003 | -0.003 | 0 |
| YFR035C | -0.001 | 0.009 | 0.004 | 0 |
| YNL235C | -0.013 | -0.006 | -0.009 | 0 |
| HXK1 | 0.009 | 0.004 | 0.007 | 0 |
| MGS1 | 0.001 | 0.010 | 0.005 | 0 |
| YGR228W | -0.004 | 0.001 | -0.002 | 0 |
| YNL203C | 0.009 | 0.010 | 0.010 | 0 |
| YGR242W | 0.006 | -0.004 | 0.001 | 0 |
| DUG3 | -0.008 | -0.009 | -0.009 | 0 |
| SUR1 | 0.019 | 0.001 | 0.010 | 0 |
| YBL010C | 0.003 | 0.036 | 0.020 | 0 |
| YPL041C | -0.006 | -0.003 | -0.005 | 0 |
| YBL028C | 0.007 | -0.005 | 0.001 | 0 |
| YPL025C | -0.006 | -0.010 | -0.008 | 0 |
| COR1 | -0.002 | 0.018 | 0.008 | 0 |
| AEP3 | 0.031 | 0.015 | 0.023 | 0 |

| | | | | |
|---|---|---|---|---|
| SHP1 | 0.071 | 0.021 | 0.046 | 1 |
| YPR114W | -0.014 | -0.031 | -0.023 | 0 |
| YBL070C | -0.002 | -0.002 | -0.002 | 0 |
| YPR127W | 0.000 | -0.004 | -0.002 | 0 |
| RPL23A | 0.012 | 0.015 | 0.014 | 0 |
| ASN1 | 0.003 | 0.007 | 0.005 | 0 |
| BRP1 | 0.019 | 0.006 | 0.013 | 0 |
| YPR157W | 0.015 | 0.007 | 0.011 | 0 |
| YGL024W | -0.032 | -0.017 | -0.025 | 0 |
| YPR174C | -0.004 | -0.011 | -0.008 | 0 |
| PNC1 | -0.002 | -0.008 | -0.005 | 0 |
| YPR197C | -0.023 | -0.011 | -0.017 | 0 |
| SDS23 | 0.012 | 0.013 | 0.012 | 0 |
| YCR100C | -0.001 | 0.011 | 0.005 | 0 |
| YGL072C | 0.021 | 0.022 | 0.022 | 1 |
| IRC4 | 0.002 | -0.007 | -0.003 | 0 |
| MMS2 | 0.002 | -0.034 | -0.016 | 0 |
| CDC26 | -0.006 | -0.014 | -0.010 | 0 |
| YNL234W | 0.000 | -0.005 | -0.003 | 0 |
| YFR054C | 0.007 | 0.005 | 0.006 | 0 |
| YNL217W | -0.004 | -0.002 | -0.003 | 0 |
| BNS1 | -0.014 | 0.008 | -0.003 | 0 |
| PSY2 | -0.001 | 0.002 | 0.001 | 0 |
| FMP43 | 0.006 | 0.004 | 0.005 | 0 |
| YNL190W | -0.003 | -0.005 | -0.004 | 0 |
| YPL056C | 0.013 | 0.008 | 0.010 | 0 |
| SCT1 | -0.001 | -0.037 | -0.019 | 0 |
| ISM1 | -0.016 | -0.004 | -0.010 | 0 |
| YBL029W | -0.018 | 0.002 | -0.008 | 0 |
| MET12 | 0.006 | 0.011 | 0.009 | 0 |
| PSY4 | -0.008 | 0.003 | -0.002 | 0 |
| ULA1 | 0.012 | 0.006 | 0.009 | 0 |
| YBL059W | 0.012 | -0.003 | 0.004 | 0 |
| YPR115W | -0.005 | -0.015 | -0.010 | 0 |
| YBL071C | 0.023 | 0.011 | 0.017 | 0 |
| ANT1 | -0.004 | 0.001 | -0.001 | 0 |
| TEL1 | 0.000 | -0.001 | 0.000 | 0 |
| YPR146C | -0.013 | 0.002 | -0.005 | 0 |
| YGL010W | -0.004 | -0.021 | -0.013 | 0 |
| CUR1 | 0.002 | 0.010 | 0.006 | 0 |
| PGD1 | -0.013 | 0.006 | -0.003 | 0 |
| HDA3 | -0.016 | -0.024 | -0.020 | 0 |
| YGL039W | 0.006 | -0.023 | -0.008 | 0 |
| SGE1 | -0.017 | -0.006 | -0.011 | 0 |
| YGL057C | 0.006 | 0.003 | 0.005 | 0 |
| YCR101C | -0.002 | 0.007 | 0.002 | 0 |
| RPL7A | 0.020 | 0.029 | 0.025 | 0 |
| YDR541C | -0.002 | -0.007 | -0.005 | 0 |
| MF(ALPHA)2 | 0.004 | -0.005 | 0.000 | 0 |
| IRC5 | -0.009 | -0.007 | -0.008 | 0 |
| BNI4 | 0.003 | -0.032 | -0.015 | 0 |
| IRC7 | 0.006 | 0.004 | 0.005 | 0 |
| IES2 | -0.013 | 0.017 | 0.002 | 0 |

| | | | | |
|---|---|---|---|---|
| PHB2 | 0.011 | 0.020 | 0.015 | 0 |
| YNL200C | -0.001 | -0.005 | -0.003 | 0 |
| LSC2 | 0.004 | -0.014 | -0.005 | 0 |
| SWT21 | 0.003 | 0.014 | 0.009 | 0 |
| LGE1 | -0.070 | -0.111 | -0.090 | -1 |
| YBL012C | 0.007 | -0.018 | -0.006 | 0 |
| YPL039W | -0.025 | -0.008 | -0.017 | 0 |
| SHE1 | -0.005 | -0.014 | -0.009 | 0 |
| RAD1 | 0.003 | 0.010 | 0.006 | 0 |
| EDE1 | -0.014 | -0.009 | -0.012 | 0 |
| SNF8 | -0.038 | 0.003 | -0.018 | 0 |
| YEL1 | 0.000 | 0.004 | 0.002 | 0 |
| YPR116W | 0.032 | 0.006 | 0.019 | 0 |
| RPS8A | 0.008 | -0.020 | -0.006 | 0 |
| SCD6 | 0.023 | 0.015 | 0.019 | 0 |
| AVT5 | 0.011 | 0.015 | 0.013 | 0 |
| YPR147C | -0.009 | 0.010 | 0.000 | 0 |
| ERG4 | -0.004 | -0.010 | -0.007 | 0 |
| KRE6 | -0.054 | -0.010 | -0.032 | 0 |
| TRP5 | -0.006 | 0.008 | 0.001 | 0 |
| GDB1 | 0.008 | 0.017 | 0.012 | 0 |
| YGL041C | 0.005 | -0.017 | -0.006 | 0 |
| ARR1 | -0.007 | -0.006 | -0.007 | 0 |
| RAD6 | 0.015 | -0.045 | -0.015 | 0 |
| YCR102C | 0.002 | 0.011 | 0.007 | 0 |
| HNM1 | -0.006 | 0.001 | -0.002 | 0 |
| YER039C-A | -0.005 | 0.003 | -0.001 | 0 |
| LIF1 | -0.005 | -0.010 | -0.007 | 0 |
| SAP155 | -0.011 | -0.013 | -0.012 | 0 |
| PDR16 | -0.022 | -0.047 | -0.035 | -1 |
| YFR056C | 0.001 | 0.007 | 0.004 | 0 |
| PEX17 | -0.003 | 0.007 | 0.002 | 0 |
| NAS6 | -0.010 | 0.011 | 0.000 | 0 |
| GCR2 | 0.010 | 0.016 | 0.013 | 0 |
| CPD1 | 0.000 | 0.009 | 0.004 | 0 |
| YNL184C | 0.016 | -0.010 | 0.003 | 0 |
| LEE1 | -0.005 | -0.018 | -0.012 | 0 |
| FMT1 | 0.007 | -0.007 | 0.000 | 0 |
| MET31 | -0.009 | -0.023 | -0.016 | 0 |
| HEK2 | -0.006 | 0.002 | -0.002 | 0 |
| ECM23 | 0.000 | 0.006 | 0.003 | 0 |
| YBL048W | -0.008 | -0.009 | -0.008 | 0 |
| HAT1 | 0.003 | 0.012 | 0.008 | 0 |
| SKT5 | 0.012 | 0.022 | 0.017 | 0 |
| YPR117W | -0.011 | -0.004 | -0.007 | 0 |
| SSA3 | 0.011 | -0.009 | 0.001 | 0 |
| YPR130C | 0.018 | 0.022 | 0.020 | 0 |
| MRP21 | -0.004 | -0.006 | -0.005 | 0 |
| YPR148C | -0.014 | 0.002 | -0.006 | 0 |
| PDR1 | -0.013 | -0.016 | -0.015 | 0 |
| GPH1 | 0.001 | -0.011 | -0.005 | 0 |
| CWH41 | 0.011 | 0.029 | 0.020 | 0 |
| ATG13 | -0.015 | 0.003 | -0.006 | 0 |

| | | | | |
|---|---|---|---|---|
| YGL042C | 0.034 | 0.022 | 0.028 | 1 |
| ARR2 | -0.008 | -0.026 | -0.017 | 0 |
| PKP2 | -0.013 | 0.006 | -0.003 | 0 |
| ADH7 | -0.004 | 0.016 | 0.006 | 0 |
| DBP3 | 0.009 | 0.005 | 0.007 | 0 |
| YER091C-A | 0.012 | 0.026 | 0.019 | 0 |
| PAN2 | 0.018 | 0.017 | 0.017 | 0 |
| ERJ5 | 0.043 | 0.039 | 0.041 | 1 |
| ELA1 | -0.002 | -0.014 | -0.008 | 0 |
| YFR057W | -0.004 | -0.012 | -0.008 | 0 |
| YNL213C | -0.012 | -0.006 | -0.009 | 0 |
| PHO81 | -0.002 | 0.016 | 0.007 | 0 |
| YNL198C | 0.013 | 0.022 | 0.018 | 0 |
| MGA1 | 0.001 | -0.001 | 0.000 | 0 |
| NPR1 | 0.001 | -0.006 | -0.002 | 0 |
| KTR6 | -0.002 | 0.005 | 0.001 | 0 |
| ACH1 | 0.008 | -0.016 | -0.004 | 0 |
| EGD1 | 0.003 | -0.026 | -0.011 | 0 |
| YBL036C | -0.024 | -0.010 | -0.017 | 0 |
| VTC3 | 0.002 | -0.005 | -0.002 | 0 |
| MOH1 | -0.002 | 0.001 | -0.001 | 0 |
| CIT3 | -0.012 | 0.003 | -0.004 | 0 |
| YBL062W | 0.014 | 0.012 | 0.013 | 0 |
| CLB2 | -0.002 | -0.003 | -0.003 | 0 |
| ATG8 | -0.008 | -0.010 | -0.009 | 0 |
| GRE2 | -0.010 | 0.006 | -0.002 | 0 |
| MAP2 | 0.001 | 0.000 | 0.001 | 0 |
| NCE102 | 0.007 | 0.018 | 0.012 | 0 |
| PUF4 | 0.007 | 0.002 | 0.004 | 0 |
| TIF3 | -0.001 | -0.005 | -0.003 | 0 |
| SCW11 | 0.017 | 0.029 | 0.023 | 0 |
| MLC2 | -0.013 | -0.023 | -0.018 | 0 |
| DST1 | 0.037 | 0.038 | 0.037 | 1 |
| ARR3 | -0.003 | -0.016 | -0.010 | 0 |
| YBP2 | 0.001 | 0.014 | 0.008 | 0 |
| RDS1 | 0.007 | -0.001 | 0.003 | 0 |
| YGL079W | -0.009 | -0.002 | -0.006 | 0 |
| UBP5 | -0.009 | 0.000 | -0.004 | 0 |
| TOS8 | 0.003 | -0.018 | -0.008 | 0 |
| IRC6 | 0.016 | 0.009 | 0.013 | 0 |
| URE2 | -0.056 | -0.063 | -0.059 | -1 |
| MRPL9 | 0.030 | -0.002 | 0.014 | 0 |
| VID27 | 0.012 | 0.013 | 0.013 | 0 |
| YHB1 | 0.004 | 0.003 | 0.003 | 0 |
| WHI3 | -0.032 | -0.006 | -0.019 | 0 |
| ECM15 | -0.010 | -0.001 | -0.005 | 0 |
| YNL179C | 0.000 | 0.019 | 0.009 | 0 |
| MRPL22 | -0.006 | -0.006 | -0.006 | 0 |
| SEC28 | 0.019 | 0.019 | 0.019 | 0 |
| RPL42A | 0.013 | 0.020 | 0.017 | 0 |
| YIA6 | -0.002 | 0.002 | 0.000 | 0 |
| YKL200C | -0.010 | -0.013 | -0.011 | 0 |
| YIL059C | -0.002 | 0.006 | 0.002 | 0 |

| | | | | |
|---|---|---|---|---|
| SRY1 | 0.009 | 0.002 | 0.006 | 0 |
| YFL015C | -0.003 | 0.001 | -0.001 | 0 |
| YKR015C | 0.001 | -0.004 | -0.002 | 0 |
| YFL035C-B | -0.004 | -0.001 | -0.003 | 0 |
| OPI8 | -0.017 | 0.019 | 0.001 | 0 |
| YFL052W | -0.003 | 0.002 | -0.001 | 0 |
| RHO4 | 0.007 | -0.002 | 0.003 | 0 |
| CMK1 | -0.009 | 0.002 | -0.003 | 0 |
| YDR249C | 0.001 | 0.005 | 0.003 | 0 |
| GND2 | 0.000 | 0.011 | 0.005 | 0 |
| SWM1 | 0.015 | 0.017 | 0.016 | 0 |
| YOR1 | -0.008 | -0.020 | -0.014 | 0 |
| YDR274C | -0.012 | -0.009 | -0.011 | 0 |
| EST3 | 0.009 | 0.010 | 0.010 | 0 |
| RTT103 | -0.017 | 0.011 | -0.003 | 0 |
| DAL2 | 0.003 | 0.004 | 0.004 | 0 |
| YDR307W | 0.005 | 0.013 | 0.009 | 0 |
| YKL162C-A | 0.000 | 0.008 | 0.004 | 0 |
| ASP1 | 0.000 | 0.000 | 0.000 | 0 |
| YKR078W | -0.001 | 0.002 | 0.001 | 0 |
| INP51 | 0.014 | 0.004 | 0.009 | 0 |
| ABF2 | 0.031 | -0.017 | 0.007 | 0 |
| YIL025C | 0.006 | 0.022 | 0.014 | 0 |
| YMR086C-A | -0.010 | 0.020 | 0.005 | 0 |
| CBR1 | 0.011 | 0.023 | 0.017 | 0 |
| AIF1 | 0.003 | -0.004 | 0.000 | 0 |
| YNL176C | -0.002 | -0.006 | -0.004 | 0 |
| YIL077C | -0.014 | 0.000 | -0.007 | 0 |
| YGP1 | 0.004 | 0.000 | 0.002 | 0 |
| NAS2 | -0.005 | 0.000 | -0.003 | 0 |
| LOS1 | -0.001 | 0.007 | 0.003 | 0 |
| YIL060W | -0.010 | -0.005 | -0.007 | 0 |
| MCH2 | 0.001 | 0.007 | 0.004 | 0 |
| PAU5 | 0.003 | -0.008 | -0.003 | 0 |
| YKR016W | 0.019 | 0.020 | 0.019 | 0 |
| RPO41 | -0.003 | -0.014 | -0.009 | 0 |
| UTH1 | -0.001 | -0.012 | -0.006 | 0 |
| DAK2 | -0.005 | -0.005 | -0.005 | 0 |
| TRM2 | 0.007 | -0.004 | 0.001 | 0 |
| GSY1 | -0.001 | 0.004 | 0.002 | 0 |
| EXG2 | -0.005 | 0.004 | 0.000 | 0 |
| BGL2 | -0.001 | 0.003 | 0.001 | 0 |
| BSC2 | 0.005 | 0.008 | 0.006 | 0 |
| MET28 | 0.000 | 0.000 | 0.000 | 0 |
| YDR290W | -0.014 | 0.001 | -0.006 | 0 |
| DAL7 | -0.001 | 0.006 | 0.002 | 0 |
| GIC2 | 0.006 | 0.013 | 0.010 | 0 |
| DID2 | -0.022 | -0.023 | -0.023 | -1 |
| MRPL35 | -0.002 | -0.013 | -0.008 | 0 |
| MTD1 | 0.008 | 0.006 | 0.007 | 0 |
| EPS1 | 0.006 | -0.001 | 0.002 | 0 |
| IRC21 | -0.013 | -0.008 | -0.010 | 0 |
| KRE27 | 0.018 | 0.019 | 0.019 | 0 |

| | | | | |
|---|---|---|---|---|
| YMR086W | -0.002 | -0.001 | -0.001 | 0 |
| SKG6 | 0.012 | 0.008 | 0.010 | 0 |
| NOP13 | -0.001 | -0.003 | -0.002 | 0 |
| AIR1 | -0.018 | -0.021 | -0.020 | 0 |
| ASI2 | 0.012 | 0.005 | 0.008 | 0 |
| URM1 | 0.021 | 0.010 | 0.015 | 0 |
| ADD66 | 0.012 | 0.015 | 0.014 | 0 |
| YIL067C | 0.000 | 0.004 | 0.002 | 0 |
| YKL222C | 0.000 | 0.010 | 0.005 | 0 |
| GAT1 | 0.004 | 0.013 | 0.009 | 0 |
| YKR017C | -0.011 | -0.003 | -0.007 | 0 |
| YFL040W | -0.001 | 0.000 | -0.001 | 0 |
| YKR043C | 0.015 | -0.002 | 0.007 | 0 |
| YFL054C | -0.005 | 0.000 | -0.002 | 0 |
| RPS21A | 0.003 | 0.011 | 0.007 | 0 |
| YFR016C | -0.008 | -0.003 | -0.005 | 0 |
| YDR250C | 0.001 | 0.004 | 0.003 | 0 |
| YGR259C | -0.001 | -0.011 | -0.006 | 0 |
| YDR262W | -0.003 | -0.013 | -0.008 | 0 |
| YGR283C | -0.017 | -0.013 | -0.015 | 0 |
| PMP3 | 0.000 | -0.017 | -0.008 | 0 |
| YAP5 | -0.005 | 0.006 | 0.000 | 0 |
| HRQ1 | -0.008 | 0.007 | 0.000 | 0 |
| MGA2 | -0.031 | -0.008 | -0.019 | 0 |
| SUM1 | -0.077 | -0.059 | -0.068 | -1 |
| CCP1 | -0.001 | -0.004 | -0.003 | 0 |
| PEP7 | 0.040 | 0.041 | 0.040 | 1 |
| NUP133 | 0.032 | 0.022 | 0.027 | 1 |
| TIR3 | -0.003 | -0.015 | -0.009 | 0 |
| YMR075C-A | 0.002 | 0.013 | 0.008 | 0 |
| YIL028W | 0.012 | 0.005 | 0.008 | 0 |
| YMR087W | -0.007 | 0.001 | -0.003 | 0 |
| AGE2 | 0.015 | 0.012 | 0.013 | 0 |
| COS10 | 0.051 | 0.059 | 0.055 | 1 |
| MDG1 | -0.002 | 0.001 | 0.000 | 0 |
| SDS3 | 0.011 | -0.009 | 0.001 | 0 |
| YNL157W | -0.001 | 0.003 | 0.001 | 0 |
| FAA3 | -0.008 | -0.006 | -0.007 | 0 |
| YKL207W | 0.008 | 0.027 | 0.018 | 0 |
| RPS24B | 0.005 | -0.010 | -0.003 | 0 |
| VPS1 | -0.014 | -0.012 | -0.013 | 0 |
| BUD27 | 0.006 | -0.002 | 0.002 | 0 |
| YKR018C | -0.002 | 0.005 | 0.002 | 0 |
| FET5 | -0.005 | -0.001 | -0.003 | 0 |
| UIP5 | 0.007 | 0.001 | 0.004 | 0 |
| AGP3 | -0.004 | 0.002 | -0.001 | 0 |
| GLG1 | 0.012 | -0.002 | 0.005 | 0 |
| YFR017C | -0.004 | 0.019 | 0.007 | 0 |
| PAM1 | -0.021 | 0.002 | -0.009 | 0 |
| TNA1 | 0.000 | -0.015 | -0.008 | 0 |
| DIN7 | -0.006 | -0.010 | -0.008 | 0 |
| ERV29 | 0.037 | 0.038 | 0.038 | 1 |
| MTH1 | 0.000 | 0.006 | 0.003 | 0 |

| | | | | |
|---|---|---|---|---|
| MUC1 | -0.006 | 0.005 | 0.000 | 0 |
| SSD1 | -0.050 | -0.058 | -0.054 | -1 |
| LYS1 | -0.003 | -0.011 | -0.007 | 0 |
| SSF2 | -0.008 | 0.011 | 0.001 | 0 |
| GPT2 | 0.005 | -0.005 | 0.000 | 0 |
| PEX3 | -0.006 | -0.002 | -0.004 | 0 |
| HBS1 | 0.001 | -0.010 | -0.004 | 0 |
| YIL012W | 0.005 | -0.004 | 0.000 | 0 |
| RCO1 | 0.000 | 0.002 | 0.001 | 0 |
| YIL029C | 0.010 | 0.001 | 0.005 | 0 |
| VBA1 | -0.006 | 0.013 | 0.003 | 0 |
| PIG2 | -0.005 | 0.006 | 0.001 | 0 |
| YOL013W-A | 0.015 | 0.006 | 0.011 | 0 |
| YNL170W | 0.058 | 0.041 | 0.050 | 1 |
| YIL086C | -0.009 | -0.004 | -0.007 | 0 |
| NSG2 | 0.002 | 0.003 | 0.002 | 0 |
| DOT5 | -0.015 | -0.001 | -0.008 | 0 |
| RPL42B | 0.026 | 0.021 | 0.024 | 1 |
| MAM33 | -0.003 | -0.005 | -0.004 | 0 |
| OSH6 | -0.004 | 0.019 | 0.008 | 0 |
| BST1 | -0.016 | -0.020 | -0.018 | 0 |
| VPS51 | 0.002 | 0.002 | 0.002 | 0 |
| YFL043C | 0.003 | -0.002 | 0.001 | 0 |
| YKR045C | 0.005 | -0.012 | -0.004 | 0 |
| AAD6 | -0.003 | 0.001 | -0.001 | 0 |
| TIF1 | 0.014 | -0.009 | 0.003 | 0 |
| YFR018C | -0.007 | -0.002 | -0.005 | 0 |
| BTT1 | -0.001 | 0.015 | 0.007 | 0 |
| APL6 | -0.029 | -0.145 | -0.087 | -1 |
| AKR1 | 0.001 | 0.003 | 0.002 | 0 |
| ZUO1 | 0.034 | 0.021 | 0.027 | 1 |
| YDR278C | 0.005 | 0.006 | 0.005 | 0 |
| YIR020C | -0.013 | 0.004 | -0.005 | 0 |
| DPL1 | 0.010 | 0.012 | 0.011 | 0 |
| YIR035C | 0.001 | 0.001 | 0.001 | 0 |
| PIB1 | -0.002 | 0.008 | 0.003 | 0 |
| MET1 | 0.009 | 0.011 | 0.010 | 0 |
| UBX5 | 0.010 | 0.023 | 0.017 | 0 |
| ECM40 | -0.087 | -0.120 | -0.103 | -1 |
| PDR11 | 0.008 | -0.007 | 0.001 | 0 |
| VPS20 | -0.037 | -0.018 | -0.027 | 0 |
| YIL032C | 0.007 | -0.005 | 0.001 | 0 |
| YTA12 | 0.001 | -0.023 | -0.011 | 0 |
| DFG10 | 0.001 | 0.013 | 0.007 | 0 |
| ADH1 | -0.030 | -0.045 | -0.037 | -1 |
| YNL171C | -0.003 | -0.010 | -0.006 | 0 |
| YIL087C | -0.003 | 0.005 | 0.001 | 0 |
| YNL155W | -0.002 | -0.003 | -0.003 | 0 |
| YIL015C-A | -0.005 | 0.004 | 0.000 | 0 |
| CBT1 | -0.005 | 0.004 | 0.000 | 0 |
| PCI8 | 0.006 | -0.003 | 0.002 | 0 |
| YKR005C | -0.006 | 0.001 | -0.002 | 0 |
| STE2 | 0.000 | -0.013 | -0.006 | 0 |

| | | | | |
|---|---|---|---|---|
| ALY1 | -0.007 | 0.003 | -0.002 | 0 |
| OTU1 | 0.006 | 0.000 | 0.003 | 0 |
| YKR047W | 0.008 | -0.003 | 0.002 | 0 |
| LOC1 | 0.018 | 0.056 | 0.037 | 0 |
| UTP30 | 0.017 | 0.009 | 0.013 | 0 |
| YFR020W | -0.003 | 0.006 | 0.001 | 0 |
| MET32 | -0.002 | -0.002 | -0.002 | 0 |
| SAY1 | 0.013 | -0.001 | 0.006 | 0 |
| PEX10 | -0.003 | -0.016 | -0.010 | 0 |
| BIO2 | -0.004 | -0.003 | -0.004 | 0 |
| RNH202 | 0.007 | 0.005 | 0.006 | 0 |
| YIR020W-B | -0.002 | 0.001 | -0.001 | 0 |
| IRC24 | 0.006 | 0.004 | 0.005 | 0 |
| RAD34 | -0.003 | 0.008 | 0.002 | 0 |
| YKR070W | 0.007 | -0.003 | 0.002 | 0 |
| IRC3 | -0.005 | 0.001 | -0.002 | 0 |
| RIM9 | -0.023 | -0.028 | -0.025 | -1 |
| MNT3 | 0.000 | -0.005 | -0.002 | 0 |
| CTF18 | -0.004 | -0.006 | -0.005 | 0 |
| CAP2 | 0.008 | -0.002 | 0.003 | 0 |
| YMR090W | -0.011 | 0.006 | -0.003 | 0 |
| PCL7 | -0.004 | 0.009 | 0.003 | 0 |
| YOL087C | -0.078 | -0.156 | -0.117 | -1 |
| PSD1 | 0.066 | 0.039 | 0.052 | 1 |
| AVT7 | -0.013 | -0.002 | -0.007 | 0 |
| YCK2 | -0.003 | -0.008 | -0.005 | 0 |
| NOT3 | -0.007 | 0.009 | 0.001 | 0 |
| TRP3 | -0.009 | 0.003 | -0.003 | 0 |
| MEH1 | -0.011 | -0.006 | -0.008 | 0 |
| GYP8 | 0.000 | -0.013 | -0.006 | 0 |
| DBP7 | -0.030 | -0.017 | -0.024 | 0 |
| FMP32 | 0.005 | -0.005 | 0.000 | 0 |
| NAP1 | 0.015 | -0.002 | 0.007 | 0 |
| YFR006W | -0.008 | 0.010 | 0.001 | 0 |
| KTR2 | 0.008 | 0.002 | 0.005 | 0 |
| ATG18 | -0.012 | -0.002 | -0.007 | 0 |
| CHL4 | -0.013 | 0.006 | -0.004 | 0 |
| YGR266W | 0.009 | 0.007 | 0.008 | 0 |
| YDR266C | -0.008 | -0.001 | -0.004 | 0 |
| YGR287C | 0.002 | 0.002 | 0.002 | 0 |
| PHM6 | 0.000 | 0.005 | 0.002 | 0 |
| MRS1 | -0.002 | 0.011 | 0.004 | 0 |
| HDA2 | -0.015 | -0.011 | -0.013 | 0 |
| HYR1 | 0.004 | 0.003 | 0.003 | 0 |
| IPK1 | 0.031 | 0.047 | 0.039 | 1 |
| SIS2 | 0.005 | 0.009 | 0.007 | 0 |
| YDR333C | -0.008 | 0.005 | -0.001 | 0 |
| KAR5 | -0.003 | -0.014 | -0.008 | 0 |
| BAR1 | 0.005 | -0.002 | 0.002 | 0 |
| NAM7 | 0.012 | 0.002 | 0.007 | 0 |
| CKA1 | 0.009 | -0.002 | 0.003 | 0 |
| NPL6 | -0.003 | -0.026 | -0.015 | 0 |
| RHR2 | 0.017 | 0.036 | 0.027 | 0 |

| | | | | |
|---|---|---|---|---|
| MPD2 | -0.011 | 0.009 | -0.001 | 0 |
| FMP41 | -0.008 | -0.007 | -0.007 | 0 |
| ICE2 | 0.005 | 0.001 | 0.003 | 0 |
| ALF1 | 0.078 | 0.051 | 0.065 | 1 |
| PKP1 | -0.005 | -0.001 | -0.003 | 0 |
| SAC1 | 0.007 | 0.011 | 0.009 | 0 |
| SER33 | 0.005 | -0.001 | 0.002 | 0 |
| FOX2 | -0.006 | 0.013 | 0.003 | 0 |
| CAF16 | 0.014 | 0.009 | 0.011 | 0 |
| GCN3 | -0.008 | 0.005 | -0.001 | 0 |
| RGD2 | -0.004 | -0.014 | -0.009 | 0 |
| FMP46 | 0.008 | -0.005 | 0.001 | 0 |
| YFH7 | -0.014 | 0.001 | -0.007 | 0 |
| OAF3 | 0.002 | 0.004 | 0.003 | 0 |
| ROG3 | -0.001 | -0.001 | -0.001 | 0 |
| RMD5 | 0.002 | 0.005 | 0.004 | 0 |
| HUA1 | 0.000 | -0.002 | -0.001 | 0 |
| YDR269C | -0.008 | -0.016 | -0.012 | 0 |
| MAL13 | 0.011 | 0.018 | 0.014 | 0 |
| YDR282C | 0.001 | 0.000 | 0.001 | 0 |
| YIR024C | -0.007 | 0.005 | -0.001 | 0 |
| SUR2 | -0.028 | -0.015 | -0.022 | 0 |
| OMS1 | -0.005 | -0.006 | -0.005 | 0 |
| YKR073C | 0.007 | -0.006 | 0.000 | 0 |
| SWR1 | -0.083 | -0.074 | -0.079 | -1 |
| UBX4 | 0.027 | 0.009 | 0.018 | 0 |
| SNL1 | 0.041 | 0.026 | 0.033 | 1 |
| ISF1 | -0.007 | -0.007 | -0.007 | 0 |
| CST6 | -0.001 | 0.000 | -0.001 | 0 |
| AIP1 | -0.008 | 0.000 | -0.004 | 0 |
| YIL057C | -0.013 | 0.001 | -0.006 | 0 |
| HAL9 | 0.010 | -0.026 | -0.008 | 0 |
| SKO1 | -0.002 | -0.004 | -0.003 | 0 |
| RSM25 | 0.018 | 0.019 | 0.018 | 0 |
| DPH2 | 0.004 | 0.007 | 0.006 | 0 |
| SYG1 | -0.011 | -0.004 | -0.007 | 0 |
| DOA1 | -0.015 | -0.031 | -0.023 | 0 |
| YIL089W | 0.001 | 0.007 | 0.004 | 0 |
| YKR011C | -0.015 | -0.008 | -0.011 | 0 |
| AGX1 | 0.008 | -0.006 | 0.001 | 0 |
| GMH1 | -0.010 | -0.001 | -0.005 | 0 |
| EMP47 | 0.003 | -0.002 | 0.001 | 0 |
| TRK2 | -0.003 | 0.002 | 0.000 | 0 |
| FAR7 | 0.005 | 0.006 | 0.006 | 0 |
| PAM17 | 0.017 | 0.021 | 0.019 | 0 |
| PES4 | 0.008 | -0.001 | 0.003 | 0 |
| CTA1 | -0.012 | 0.010 | -0.001 | 0 |
| YGR269W | -0.009 | -0.011 | -0.010 | 0 |
| CCC2 | -0.004 | -0.006 | -0.005 | 0 |
| YGR290W | -0.003 | 0.004 | 0.000 | 0 |
| DPP1 | 0.005 | 0.000 | 0.002 | 0 |
| MND2 | -0.007 | 0.001 | -0.003 | 0 |
| ATP5 | 0.006 | -0.013 | -0.004 | 0 |

| | | | | |
|---|---|---|---|---|
| GTT1 | 0.012 | 0.003 | 0.008 | 0 |
| HIM1 | -0.012 | 0.008 | -0.002 | 0 |
| YKR074W | 0.020 | -0.015 | 0.003 | 0 |
| MSN5 | -0.005 | 0.007 | 0.001 | 0 |
| AVO2 | 0.002 | -0.015 | -0.006 | 0 |
| VID28 | 0.008 | 0.000 | 0.004 | 0 |
| YMR082C | -0.013 | 0.010 | -0.002 | 0 |
| PRM2 | 0.002 | 0.014 | 0.008 | 0 |
| YNR070W | 0.007 | 0.005 | 0.006 | 0 |
| YIL064W | 0.000 | 0.006 | 0.003 | 0 |
| MSH2 | 0.001 | 0.006 | 0.004 | 0 |
| BNI5 | -0.013 | -0.011 | -0.012 | 0 |
| PRK1 | -0.004 | 0.009 | 0.002 | 0 |
| PEX1 | 0.002 | -0.007 | -0.002 | 0 |
| RPL34B | 0.016 | 0.021 | 0.018 | 0 |
| YRA2 | -0.005 | -0.003 | -0.004 | 0 |
| YIL092W | -0.006 | -0.001 | -0.004 | 0 |
| YKR012C | -0.014 | 0.007 | -0.004 | 0 |
| HAC1 | 0.037 | 0.040 | 0.039 | 1 |
| SPO14 | -0.048 | -0.071 | -0.060 | -1 |
| SWP82 | 0.007 | 0.005 | 0.006 | 0 |
| YKR051W | 0.006 | -0.016 | -0.005 | 0 |
| GCN20 | 0.009 | 0.001 | 0.005 | 0 |
| PEX5 | -0.009 | 0.011 | 0.001 | 0 |
| LSB3 | 0.007 | -0.001 | 0.003 | 0 |
| RKM4 | -0.004 | -0.007 | -0.005 | 0 |
| YTA7 | 0.030 | 0.041 | 0.035 | 1 |
| YDR271C | -0.006 | -0.005 | -0.006 | 0 |
| ECM12 | 0.006 | 0.017 | 0.011 | 0 |
| ZIP1 | -0.001 | 0.004 | 0.002 | 0 |
| YVH1 | 0.004 | 0.018 | 0.011 | 0 |
| CPR5 | 0.007 | 0.013 | 0.010 | 0 |
| YPS6 | 0.007 | 0.004 | 0.006 | 0 |
| MCM21 | -0.006 | -0.006 | -0.006 | 0 |
| YKR075C | 0.011 | -0.007 | 0.002 | 0 |
| YDR336W | -0.007 | 0.002 | -0.003 | 0 |
| NAT4 | 0.007 | 0.004 | 0.006 | 0 |
| HIS6 | -0.020 | -0.026 | -0.023 | 0 |
| ADH3 | -0.001 | 0.010 | 0.004 | 0 |
| TED1 | -0.024 | -0.025 | -0.025 | -1 |
| YNR071C | -0.002 | 0.005 | 0.002 | 0 |
| FIS1 | 0.012 | 0.002 | 0.007 | 0 |
| SPO21 | -0.013 | -0.004 | -0.008 | 0 |
| YNL165W | -0.003 | -0.007 | -0.005 | 0 |
| YIL096C | 0.007 | 0.003 | 0.005 | 0 |
| PTK1 | 0.001 | 0.002 | 0.002 | 0 |
| YIL054W | -0.001 | 0.015 | 0.007 | 0 |
| URA1 | 0.005 | -0.049 | -0.022 | 0 |
| YFL006W | -0.009 | 0.000 | -0.005 | 0 |
| PRY2 | -0.011 | 0.010 | 0.000 | 0 |
| YFL032W | 0.034 | 0.041 | 0.037 | 1 |
| YKR032W | -0.014 | 0.001 | -0.006 | 0 |
| ALR2 | 0.006 | -0.003 | 0.002 | 0 |

| | | | | |
|---|---|---|---|---|
| MRS4 | -0.010 | -0.005 | -0.008 | 0 |
| UBP6 | 0.006 | 0.000 | 0.003 | 0 |
| VHS1 | -0.003 | -0.005 | -0.004 | 0 |
| ULI1 | 0.008 | 0.008 | 0.008 | 0 |
| HSP78 | 0.002 | -0.005 | -0.001 | 0 |
| RTT102 | -0.002 | 0.006 | 0.002 | 0 |
| GLO2 | -0.001 | -0.008 | -0.005 | 0 |
| YHR039C-B | 0.011 | -0.010 | 0.001 | 0 |
| YDR286C | -0.017 | 0.008 | -0.004 | 0 |
| DAL1 | -0.011 | -0.005 | -0.008 | 0 |
| HNT2 | 0.002 | 0.021 | 0.011 | 0 |
| YIR042C | 0.011 | 0.005 | 0.008 | 0 |
| YDR319C | -0.013 | -0.004 | -0.009 | 0 |
| ECM4 | 0.001 | -0.009 | -0.004 | 0 |
| MRPS28 | 0.002 | 0.011 | 0.006 | 0 |
| MOT3 | -0.004 | -0.011 | -0.008 | 0 |
| YKE4 | -0.005 | -0.011 | -0.008 | 0 |
| YMR084W | -0.002 | 0.009 | 0.003 | 0 |
| APQ12 | 0.011 | 0.019 | 0.015 | 0 |
| HXT17 | 0.009 | 0.007 | 0.008 | 0 |
| HOP1 | 0.006 | -0.002 | 0.002 | 0 |
| YOL092W | -0.012 | -0.013 | -0.012 | 0 |
| IBD2 | 0.004 | 0.009 | 0.007 | 0 |
| FYV10 | -0.002 | -0.006 | -0.004 | 0 |
| YKL199C | -0.005 | 0.015 | 0.005 | 0 |
| YIL055C | -0.014 | 0.000 | -0.007 | 0 |
| JEN1 | -0.004 | 0.000 | -0.002 | 0 |
| HXT10 | 0.002 | -0.006 | -0.002 | 0 |
| YPT52 | -0.023 | -0.004 | -0.013 | 0 |
| YFL034W | 0.001 | -0.001 | 0.000 | 0 |
| YKR033C | -0.020 | -0.024 | -0.022 | -1 |
| YFL051C | 0.000 | -0.003 | -0.001 | 0 |
| DYN1 | -0.003 | -0.008 | -0.005 | 0 |
| YFR012W | -0.019 | -0.008 | -0.014 | 0 |
| YDR248C | -0.001 | -0.003 | -0.002 | 0 |
| RPL2A | 0.013 | 0.013 | 0.013 | 0 |
| YAP6 | 0.009 | 0.005 | 0.007 | 0 |
| SCW4 | -0.020 | -0.028 | -0.024 | -1 |
| DON1 | -0.006 | -0.017 | -0.011 | 0 |
| YHR079C-B | 0.008 | 0.003 | 0.005 | 0 |
| INM2 | -0.004 | 0.002 | -0.001 | 0 |
| DAL4 | -0.014 | 0.000 | -0.007 | 0 |
| YDR306C | 0.005 | 0.012 | 0.009 | 0 |
| YKL033W-A | -0.003 | -0.001 | -0.002 | 0 |
| SWA2 | 0.035 | 0.041 | 0.038 | 1 |
| MSA2 | 0.007 | 0.005 | 0.006 | 0 |
| YIL001W | -0.003 | 0.004 | 0.001 | 0 |
| TVP18 | 0.010 | 0.002 | 0.006 | 0 |
| YIL024C | -0.007 | 0.004 | -0.002 | 0 |
| YMR085W | 0.007 | -0.010 | -0.001 | 0 |
| GVP36 | -0.002 | -0.001 | -0.002 | 0 |
| YNR073C | 0.004 | 0.008 | 0.006 | 0 |
| SPO22 | 0.012 | -0.006 | 0.003 | 0 |

| | | | | |
|---|---|---|---|---|
| TRM10 | -0.016 | 0.007 | -0.005 | 0 |
| HMI1 | 0.011 | 0.001 | 0.006 | 0 |
| CSR2 | 0.010 | -0.008 | 0.001 | 0 |
| ZEO1 | 0.000 | -0.012 | -0.006 | 0 |
| MAK3 | -0.066 | -0.056 | -0.061 | -1 |
| SMF1 | 0.000 | -0.020 | -0.010 | 0 |
| UBA3 | 0.014 | 0.002 | 0.008 | 0 |
| DYN2 | 0.006 | 0.006 | 0.006 | 0 |
| YPR084W | 0.005 | -0.002 | 0.001 | 0 |
| YER097W | 0.002 | -0.004 | -0.001 | 0 |
| SNT309 | 0.076 | 0.063 | 0.069 | 1 |
| YHR162W | 0.013 | -0.005 | 0.004 | 0 |
| ASF1 | 0.056 | 0.037 | 0.046 | 1 |
| RAD5 | 0.035 | 0.024 | 0.029 | 1 |
| TOK1 | -0.001 | -0.003 | -0.002 | 0 |
| MRPS8 | -0.001 | 0.002 | 0.000 | 0 |
| YJL068C | 0.003 | 0.002 | 0.002 | 0 |
| YMR316C-A | 0.001 | -0.011 | -0.005 | 0 |
| ZAP1 | -0.021 | 0.002 | -0.010 | 0 |
| CMK2 | -0.003 | 0.003 | 0.000 | 0 |
| LOH1 | -0.002 | 0.004 | 0.001 | 0 |
| YML095C-A | 0.049 | 0.055 | 0.052 | 1 |
| HSE1 | -0.009 | -0.017 | -0.013 | 0 |
| URA5 | -0.020 | -0.010 | -0.015 | 0 |
| FYV4 | 0.017 | -0.011 | 0.003 | 0 |
| YML122C | -0.005 | -0.002 | -0.004 | 0 |
| ENB1 | -0.009 | 0.001 | -0.004 | 0 |
| YMR103C | -0.009 | -0.014 | -0.011 | 0 |
| BUD4 | 0.018 | 0.015 | 0.017 | 0 |
| RPL15B | -0.010 | 0.003 | -0.003 | 0 |
| MCM22 | 0.000 | 0.003 | 0.002 | 0 |
| ICL2 | -0.004 | -0.006 | -0.005 | 0 |
| SRP40 | -0.008 | -0.003 | -0.006 | 0 |
| COQ3 | -0.021 | 0.047 | 0.013 | 0 |
| SRO7 | -0.004 | -0.005 | -0.005 | 0 |
| SHR5 | -0.013 | -0.009 | -0.011 | 0 |
| NHP6A | 0.006 | -0.001 | 0.002 | 0 |
| TRM11 | 0.008 | 0.004 | 0.006 | 0 |
| HOS1 | -0.004 | 0.008 | 0.002 | 0 |
| GLC3 | -0.004 | 0.004 | 0.000 | 0 |
| VPS69 | -0.017 | 0.011 | -0.003 | 0 |
| UBP9 | 0.016 | 0.008 | 0.012 | 0 |
| SPT10 | 0.057 | 0.027 | 0.042 | 1 |
| MTG2 | -0.009 | 0.005 | -0.002 | 0 |
| MDV1 | -0.005 | 0.002 | -0.001 | 0 |
| SMF3 | -0.009 | -0.003 | -0.006 | 0 |
| SRS2 | 0.018 | 0.015 | 0.016 | 0 |
| YMR316C-B | 0.006 | -0.003 | 0.001 | 0 |
| YJL055W | 0.001 | -0.001 | 0.000 | 0 |
| RPS7A | -0.022 | 0.014 | -0.004 | 0 |
| IRC18 | 0.007 | 0.003 | 0.005 | 0 |
| YML096W | 0.006 | -0.017 | -0.005 | 0 |
| PRS3 | 0.013 | 0.045 | 0.029 | 0 |

| | | | | |
|---|---|---|---|---|
| PML39 | 0.017 | 0.009 | 0.013 | 0 |
| HTD2 | -0.017 | 0.018 | 0.001 | 0 |
| PHO84 | 0.008 | -0.015 | -0.003 | 0 |
| YOL159C | -0.018 | -0.001 | -0.009 | 0 |
| PGM2 | -0.002 | -0.004 | -0.003 | 0 |
| VPS25 | -0.005 | -0.020 | -0.012 | 0 |
| YMR122C | -0.001 | 0.005 | 0.002 | 0 |
| ECM17 | 0.017 | 0.013 | 0.015 | 0 |
| YHR192W | -0.042 | -0.040 | -0.041 | -1 |
| YOL098C | 0.005 | -0.004 | 0.000 | 0 |
| VMA13 | -0.022 | -0.004 | -0.013 | 0 |
| MDY2 | -0.021 | -0.013 | -0.017 | 0 |
| YPR053C | 0.012 | 0.003 | 0.008 | 0 |
| MDH2 | -0.009 | -0.015 | -0.012 | 0 |
| SPE3 | -0.020 | 0.041 | 0.011 | 0 |
| YER064C | -0.152 | 0.006 | -0.073 | 0 |
| YPR089W | 0.008 | 0.002 | 0.005 | 0 |
| CAF130 | 0.026 | 0.006 | 0.016 | 0 |
| NIT2 | 0.000 | -0.001 | 0.000 | 0 |
| SVP26 | -0.007 | -0.005 | -0.006 | 0 |
| GZF3 | -0.004 | 0.001 | -0.001 | 0 |
| MLH2 | 0.001 | 0.005 | 0.003 | 0 |
| SIP4 | 0.003 | 0.011 | 0.007 | 0 |
| EAR1 | -0.031 | -0.027 | -0.029 | -1 |
| YJL067W | -0.003 | -0.007 | -0.005 | 0 |
| DIA1 | 0.003 | -0.008 | -0.003 | 0 |
| PEP8 | -0.031 | -0.041 | -0.036 | -1 |
| MCH5 | 0.001 | 0.001 | 0.001 | 0 |
| SNX4 | 0.008 | 0.002 | 0.005 | 0 |
| VPS9 | -0.021 | -0.025 | -0.023 | -1 |
| YHL039W | -0.002 | -0.001 | -0.002 | 0 |
| YML108W | -0.014 | -0.012 | -0.013 | 0 |
| YHR127W | 0.008 | 0.012 | 0.010 | 0 |
| TUB3 | -0.013 | -0.008 | -0.011 | 0 |
| YOL160W | 0.020 | -0.003 | 0.008 | 0 |
| YKU80 | 0.006 | 0.013 | 0.009 | 0 |
| URA8 | 0.010 | -0.003 | 0.003 | 0 |
| PKR1 | 0.011 | 0.008 | 0.010 | 0 |
| YJR146W | 0.004 | -0.009 | -0.002 | 0 |
| SUT2 | 0.010 | -0.002 | 0.004 | 0 |
| PTR2 | -0.031 | 0.009 | -0.011 | 0 |
| YOL099C | 0.004 | -0.005 | -0.001 | 0 |
| IRC16 | 0.003 | -0.006 | -0.002 | 0 |
| MSB4 | 0.011 | -0.003 | 0.004 | 0 |
| SMK1 | 0.001 | -0.006 | -0.003 | 0 |
| YGK3 | 0.007 | -0.005 | 0.001 | 0 |
| MED1 | 0.050 | 0.056 | 0.053 | 1 |
| YER077C | 0.006 | 0.007 | 0.007 | 0 |
| YPR090W | 0.006 | 0.007 | 0.006 | 0 |
| YGR210C | 0.012 | 0.002 | 0.007 | 0 |
| LSM1 | 0.027 | 0.036 | 0.031 | 1 |
| CTF8 | 0.003 | -0.007 | -0.002 | 0 |
| PRM10 | 0.001 | 0.001 | 0.001 | 0 |

| | | | | |
|---|---|---|---|---|
| YLR036C | 0.010 | -0.002 | 0.004 | 0 |
| ARG3 | 0.042 | 0.041 | 0.041 | 1 |
| YMR181C | -0.002 | -0.020 | -0.011 | 0 |
| MPM1 | -0.004 | 0.005 | 0.001 | 0 |
| YMR317W | -0.006 | -0.007 | -0.006 | 0 |
| TDH1 | 0.002 | -0.003 | 0.000 | 0 |
| FAA1 | -0.015 | -0.007 | -0.011 | 0 |
| MAD2 | 0.014 | 0.000 | 0.007 | 0 |
| ARG81 | 0.000 | -0.008 | -0.004 | 0 |
| YHR003C | -0.008 | 0.002 | -0.003 | 0 |
| ZDS2 | 0.009 | -0.001 | 0.004 | 0 |
| YHR131C | 0.008 | 0.005 | 0.006 | 0 |
| MSC1 | 0.000 | -0.001 | 0.000 | 0 |
| YOL162W | -0.017 | -0.008 | -0.012 | 0 |
| SPG4 | -0.005 | 0.005 | 0.000 | 0 |
| ADO1 | -0.012 | -0.020 | -0.016 | 0 |
| YMR124W | -0.006 | -0.006 | -0.006 | 0 |
| HMS2 | 0.002 | 0.001 | 0.001 | 0 |
| YPR012W | 0.004 | 0.005 | 0.005 | 0 |
| PCK1 | 0.025 | 0.011 | 0.018 | 0 |
| PKH2 | 0.008 | -0.003 | 0.002 | 0 |
| YPR039W | -0.003 | -0.008 | -0.005 | 0 |
| SKM1 | 0.009 | -0.001 | 0.004 | 0 |
| BRR1 | 0.015 | 0.006 | 0.010 | 0 |
| VPS68 | -0.007 | -0.014 | -0.010 | 0 |
| YPR071W | 0.002 | 0.006 | 0.004 | 0 |
| YER078C | 0.001 | 0.007 | 0.004 | 0 |
| YPR092W | 0.011 | 0.002 | 0.006 | 0 |
| RIM4 | -0.002 | -0.006 | -0.004 | 0 |
| MTC1 | -0.011 | -0.020 | -0.016 | 0 |
| EGD2 | -0.002 | 0.007 | 0.002 | 0 |
| YJL107C | -0.007 | 0.000 | -0.003 | 0 |
| DAN2 | 0.006 | 0.003 | 0.005 | 0 |
| ALY2 | 0.008 | 0.004 | 0.006 | 0 |
| YMR209C | 0.001 | -0.001 | 0.000 | 0 |
| YJL064W | 0.007 | 0.015 | 0.011 | 0 |
| ADH6 | -0.007 | -0.009 | -0.008 | 0 |
| IRC8 | 0.029 | 0.002 | 0.015 | 0 |
| COX11 | -0.023 | 0.036 | 0.006 | 0 |
| UBC8 | 0.002 | -0.001 | 0.001 | 0 |
| TSL1 | -0.010 | -0.005 | -0.007 | 0 |
| NEM1 | -0.016 | -0.003 | -0.009 | 0 |
| DAT1 | -0.068 | -0.050 | -0.059 | -1 |
| YHR180W | 0.010 | 0.007 | 0.009 | 0 |
| YML131W | 0.011 | -0.001 | 0.005 | 0 |
| YOL163W | -0.004 | -0.004 | -0.004 | 0 |
| MYO5 | 0.005 | 0.010 | 0.007 | 0 |
| ABM1 | -0.010 | 0.000 | -0.005 | 0 |
| STO1 | -0.004 | 0.007 | 0.001 | 0 |
| YJR149W | -0.101 | 0.001 | -0.050 | 0 |
| YPR014C | -0.010 | -0.006 | -0.008 | 0 |
| UBP11 | 0.003 | 0.004 | 0.004 | 0 |
| IZH4 | -0.003 | -0.012 | -0.008 | 0 |

| | | | | |
|---|---|---|---|---|
| TIP41 | -0.012 | -0.025 | -0.019 | 0 |
| YOL114C | -0.002 | -0.009 | -0.005 | 0 |
| YMC1 | 0.011 | 0.025 | 0.018 | 0 |
| YOL131W | 0.012 | 0.006 | 0.009 | 0 |
| LTP1 | 0.010 | 0.002 | 0.006 | 0 |
| DOT6 | 0.003 | 0.002 | 0.003 | 0 |
| ASR1 | 0.006 | 0.012 | 0.009 | 0 |
| SNF6 | -0.008 | -0.033 | -0.020 | 0 |
| ALB1 | -0.028 | -0.006 | -0.017 | 0 |
| YLL030C | 0.014 | 0.008 | 0.011 | 0 |
| IME2 | -0.007 | 0.000 | -0.003 | 0 |
| COX12 | 0.002 | 0.003 | 0.002 | 0 |
| TAX4 | -0.002 | 0.003 | 0.001 | 0 |
| URA10 | -0.004 | -0.004 | -0.004 | 0 |
| DLS1 | 0.008 | 0.022 | 0.015 | 0 |
| FET4 | 0.011 | 0.000 | 0.005 | 0 |
| YJL049W | 0.008 | -0.011 | -0.002 | 0 |
| ODC1 | -0.002 | -0.013 | -0.008 | 0 |
| YPT31 | 0.014 | 0.005 | 0.010 | 0 |
| [illegible]ML100W-A | -0.008 | 0.003 | -0.002 | 0 |
| STP2 | 0.000 | 0.008 | 0.004 | 0 |
| ATR1 | 0.018 | -0.008 | 0.005 | 0 |
| PFS1 | 0.003 | 0.009 | 0.006 | 0 |
| MVP1 | -0.003 | -0.012 | -0.007 | 0 |
| OPI3 | -0.005 | 0.008 | 0.002 | 0 |
| HFD1 | -0.004 | -0.003 | -0.003 | 0 |
| YMR1 | 0.001 | -0.001 | 0.000 | 0 |
| DLT1 | 0.004 | 0.001 | 0.002 | 0 |
| DAL5 | -0.008 | 0.006 | -0.001 | 0 |
| YPR015C | 0.001 | 0.004 | 0.002 | 0 |
| BAS1 | -0.021 | 0.002 | -0.010 | 0 |
| ITR2 | 0.002 | -0.011 | -0.005 | 0 |
| PUF2 | 0.002 | -0.009 | -0.004 | 0 |
| PAP2 | 0.003 | -0.007 | -0.002 | 0 |
| YPR059C | 0.014 | 0.019 | 0.016 | 0 |
| GAS4 | 0.003 | 0.010 | 0.007 | 0 |
| TKL1 | 0.031 | 0.033 | 0.032 | 1 |
| TRP2 | -0.007 | -0.004 | -0.005 | 0 |
| SYT1 | -0.008 | -0.013 | -0.010 | 0 |
| YSC84 | 0.017 | 0.000 | 0.009 | 0 |
| YJL120W | 0.037 | 0.028 | 0.032 | 1 |
| YLL044W | -0.004 | 0.003 | -0.001 | 0 |
| MEF2 | 0.003 | -0.028 | -0.012 | 0 |
| RIC1 | -0.022 | -0.014 | -0.018 | 0 |
| IML2 | -0.010 | -0.004 | -0.007 | 0 |
| YMR279C | -0.004 | -0.001 | -0.003 | 0 |
| MRPL8 | 0.001 | 0.000 | 0.001 | 0 |
| YMR320W | -0.015 | -0.011 | -0.013 | 0 |
| UBX6 | -0.001 | -0.006 | -0.004 | 0 |
| DAK1 | 0.011 | -0.024 | -0.007 | 0 |
| SPO73 | -0.006 | -0.020 | -0.013 | 0 |
| CUE4 | 0.013 | 0.010 | 0.011 | 0 |
| SOD2 | -0.007 | 0.004 | -0.001 | 0 |

| | | | | |
|---|---|---|---|---|
| NAB6 | -0.030 | -0.032 | -0.031 | -1 |
| MDM31 | 0.007 | 0.010 | 0.008 | 0 |
| SNO1 | 0.021 | 0.004 | 0.012 | 0 |
| HOC1 | -0.005 | -0.013 | -0.009 | 0 |
| YMR111C | -0.010 | -0.002 | -0.006 | 0 |
| YJR111C | -0.005 | -0.002 | -0.004 | 0 |
| SAS2 | -0.013 | 0.005 | -0.004 | 0 |
| YJR154W | -0.002 | -0.004 | -0.003 | 0 |
| DSS4 | -0.001 | -0.012 | -0.007 | 0 |
| SKG1 | 0.005 | 0.004 | 0.004 | 0 |
| NDJ1 | -0.001 | -0.010 | -0.005 | 0 |
| OPI11 | 0.041 | 0.039 | 0.040 | 1 |
| MSN1 | -0.018 | -0.021 | -0.020 | 0 |
| ARO7 | -0.013 | 0.028 | 0.007 | 0 |
| PFK27 | -0.001 | 0.005 | 0.002 | 0 |
| OPY2 | 0.006 | 0.006 | 0.006 | 0 |
| MET6 | 0.081 | 0.070 | 0.075 | 1 |
| YPR096C | -0.010 | -0.007 | -0.009 | 0 |
| YSC83 | 0.007 | 0.004 | 0.006 | 0 |
| RPE1 | 0.021 | 0.024 | 0.022 | 1 |
| YBT1 | 0.000 | 0.007 | 0.003 | 0 |
| LSB6 | 0.005 | -0.002 | 0.002 | 0 |
| YLR040C | -0.002 | 0.009 | 0.003 | 0 |
| SCP160 | 0.000 | 0.000 | 0.000 | 0 |
| FKS3 | -0.006 | 0.009 | 0.001 | 0 |
| LAS21 | 0.033 | -0.001 | 0.016 | 0 |
| RAD50 | 0.079 | 0.021 | 0.050 | 1 |
| RTT101 | 0.058 | 0.005 | 0.032 | 0 |
| COG8 | 0.006 | -0.017 | -0.006 | 0 |
| THO1 | -0.016 | -0.014 | -0.015 | 0 |
| ML102C-A | -0.008 | -0.015 | -0.011 | 0 |
| YHR009C | 0.003 | -0.006 | -0.002 | 0 |
| ML117W-A | -0.016 | -0.023 | -0.019 | 0 |
| YLL007C | 0.012 | 0.012 | 0.012 | 0 |
| SNZ1 | 0.004 | 0.005 | 0.005 | 0 |
| BNA2 | -0.003 | 0.001 | -0.001 | 0 |
| YMR114C | -0.011 | 0.002 | -0.005 | 0 |
| YJR115W | -0.003 | 0.005 | 0.001 | 0 |
| POM152 | 0.002 | -0.002 | 0.000 | 0 |
| OMA1 | 0.000 | -0.001 | 0.000 | 0 |
| RLF2 | -0.005 | -0.013 | -0.009 | 0 |
| SIR1 | 0.019 | 0.021 | 0.020 | 0 |
| WSC3 | -0.003 | -0.023 | -0.013 | 0 |
| YPR045C | 0.031 | 0.038 | 0.035 | 1 |
| RRI2 | 0.008 | -0.003 | 0.003 | 0 |
| JID1 | -0.008 | -0.009 | -0.008 | 0 |
| IKI1 | 0.061 | 0.060 | 0.060 | 1 |
| YPR076W | 0.001 | 0.002 | 0.002 | 0 |
| IES5 | 0.010 | 0.004 | 0.007 | 0 |
| YPR097W | 0.002 | 0.001 | 0.002 | 0 |
| YHR032W | 0.021 | -0.004 | 0.008 | 0 |
| YJL118W | -0.001 | -0.009 | -0.005 | 0 |
| LDB18 | 0.016 | -0.004 | 0.006 | 0 |

| | | | | |
|---|---|---|---|---|
| CHS6 | 0.017 | 0.016 | 0.016 | 0 |
| YLR041W | -0.003 | 0.000 | -0.001 | 0 |
| PRY1 | 0.001 | 0.008 | 0.005 | 0 |
| GLC8 | 0.015 | 0.010 | 0.012 | 0 |
| BNA3 | -0.181 | 0.001 | -0.090 | 0 |
| MRPL17 | -0.039 | 0.000 | -0.020 | 0 |
| YJL046W | -0.021 | 0.002 | -0.009 | 0 |
| FPR3 | -0.003 | -0.019 | -0.011 | 0 |
| YER066W | -0.007 | -0.005 | -0.006 | 0 |
| CAC2 | -0.006 | -0.012 | -0.009 | 0 |
| THR1 | 0.006 | 0.002 | 0.004 | 0 |
| NGL3 | 0.005 | -0.005 | 0.000 | 0 |
| RIM13 | -0.037 | -0.009 | -0.023 | 0 |
| YMR099C | 0.022 | 0.009 | 0.015 | 0 |
| YJR079W | -0.011 | 0.010 | -0.001 | 0 |
| MGR3 | -0.004 | -0.003 | -0.003 | 0 |
| RSF2 | -0.001 | -0.007 | -0.004 | 0 |
| YMR130W | -0.012 | -0.002 | -0.007 | 0 |
| TVP38 | 0.001 | 0.002 | 0.001 | 0 |
| ATP20 | 0.023 | 0.027 | 0.025 | 1 |
| NFT1 | -0.007 | -0.006 | -0.006 | 0 |
| YOL106W | 0.003 | -0.014 | -0.006 | 0 |
| MCM16 | 0.011 | -0.019 | -0.004 | 0 |
| YOL118C | 0.005 | -0.003 | 0.001 | 0 |
| FCY1 | -0.012 | -0.006 | -0.009 | 0 |
| BSC6 | 0.005 | 0.005 | 0.005 | 0 |
| YPR077C | -0.012 | 0.004 | -0.004 | 0 |
| YER093C-A | 0.025 | 0.004 | 0.015 | 0 |
| YPR098C | -0.006 | 0.000 | -0.003 | 0 |
| YHR045W | -0.008 | -0.005 | -0.007 | 0 |
| YJL119C | -0.007 | -0.001 | -0.004 | 0 |
| YLL059C | -0.003 | -0.001 | -0.002 | 0 |
| SAP185 | 0.007 | 0.006 | 0.007 | 0 |
| YLR050C | 0.007 | 0.006 | 0.006 | 0 |
| ICS3 | -0.008 | -0.003 | -0.006 | 0 |
| ELP6 | 0.046 | 0.041 | 0.044 | 1 |
| YHC3 | -0.008 | 0.007 | 0.000 | 0 |
| PRM1 | 0.001 | -0.011 | -0.005 | 0 |
| YJL045W | -0.009 | -0.006 | -0.008 | 0 |
| YML090W | 0.011 | 0.004 | 0.007 | 0 |
| BUB1 | -0.006 | -0.005 | -0.006 | 0 |
| PPA1 | 0.028 | 0.016 | 0.022 | 0 |
| YML119W | -0.016 | -0.012 | -0.014 | 0 |
| GOR1 | 0.015 | -0.001 | 0.007 | 0 |
| MUB1 | 0.039 | 0.038 | 0.038 | 1 |
| EAF6 | 0.008 | 0.010 | 0.009 | 0 |
| ASC1 | 0.036 | 0.026 | 0.031 | 1 |
| YJR128W | -0.002 | -0.007 | -0.005 | 0 |
| JLP2 | -0.034 | 0.001 | -0.016 | 0 |
| TGL4 | 0.002 | -0.010 | -0.004 | 0 |
| YPR027C | -0.004 | -0.006 | -0.005 | 0 |
| YKR104W | 0.006 | 0.010 | 0.008 | 0 |
| YOL107W | 0.003 | -0.006 | -0.001 | 0 |

| | | | | |
|---|---|---|---|---|
| MSF1 | -0.022 | 0.035 | 0.006 | 0 |
| MCH4 | -0.001 | -0.006 | -0.004 | 0 |
| YPR063C | -0.002 | -0.011 | -0.006 | 0 |
| RTC1 | 0.005 | -0.006 | -0.001 | 0 |
| RAD51 | 0.008 | 0.002 | 0.005 | 0 |
| YPR099C | 0.023 | -0.011 | 0.006 | 0 |
| SSZ1 | 0.054 | 0.016 | 0.035 | 0 |
| PHO86 | -0.013 | -0.004 | -0.008 | 0 |
| YLR030W | -0.011 | -0.001 | -0.006 | 0 |
| MRPL49 | -0.002 | -0.002 | -0.002 | 0 |
| IES3 | 0.001 | 0.013 | 0.007 | 0 |
| JEM1 | -0.018 | -0.014 | -0.016 | 0 |
| TGL3 | 0.009 | 0.004 | 0.007 | 0 |
| BIT61 | 0.001 | 0.013 | 0.007 | 0 |
| TOS6 | 0.018 | -0.002 | 0.008 | 0 |
| GYP6 | -0.002 | -0.008 | -0.005 | 0 |
| GIM5 | 0.044 | 0.038 | 0.041 | 1 |
| YGR201C | -0.019 | -0.012 | -0.015 | 0 |
| NUP188 | 0.007 | 0.002 | 0.004 | 0 |
| SRB2 | 0.014 | 0.032 | 0.023 | 0 |
| NDI1 | 0.014 | 0.001 | 0.007 | 0 |
| PPM2 | -0.004 | -0.006 | -0.005 | 0 |
| SRT1 | 0.002 | 0.007 | 0.004 | 0 |
| ACF4 | -0.006 | -0.019 | -0.013 | 0 |
| (MR119W-A | -0.016 | 0.004 | -0.006 | 0 |
| YJR129C | 0.001 | -0.006 | -0.003 | 0 |
| REC114 | -0.009 | 0.007 | -0.001 | 0 |
| PXL1 | -0.014 | -0.013 | -0.014 | 0 |
| YOP1 | -0.006 | 0.000 | -0.003 | 0 |
| VBA5 | -0.001 | 0.005 | 0.002 | 0 |
| INO4 | 0.069 | 0.075 | 0.072 | 1 |
| ATG11 | -0.019 | -0.019 | -0.019 | 0 |
| RPS19A | 0.027 | 0.011 | 0.019 | 0 |
| ROX1 | -0.009 | 0.003 | -0.003 | 0 |
| YBR232C | 0.000 | -0.002 | -0.001 | 0 |
| MRL1 | 0.000 | 0.007 | 0.003 | 0 |
| SHC1 | 0.001 | -0.004 | -0.001 | 0 |
| MRPL51 | -0.013 | 0.008 | -0.002 | 0 |
| YHR140W | 0.007 | -0.005 | 0.001 | 0 |
| NCA3 | -0.006 | 0.002 | -0.002 | 0 |
| YLR031W | -0.007 | 0.000 | -0.003 | 0 |
| BCK1 | -0.003 | 0.001 | -0.001 | 0 |
| RPL13B | -0.003 | 0.000 | -0.001 | 0 |
| ARG2 | 0.055 | -0.001 | 0.027 | 0 |
| YMR315W | -0.002 | 0.001 | 0.000 | 0 |
| IKS1 | -0.002 | 0.004 | 0.001 | 0 |
| PHA2 | 0.005 | 0.001 | 0.003 | 0 |
| YJL043W | -0.002 | -0.006 | -0.004 | 0 |
| RAD10 | 0.017 | 0.004 | 0.010 | 0 |
| ADE3 | -0.094 | -0.098 | -0.096 | -1 |
| MDM1 | -0.006 | -0.010 | -0.008 | 0 |
| GTR1 | 0.000 | 0.000 | 0.000 | 0 |
| YOL150C | 0.011 | -0.005 | 0.003 | 0 |

| | | | | |
|---|---|---|---|---|
| YMR102C | 0.021 | 0.009 | 0.015 | 0 |
| YJR088C | 0.008 | 0.010 | 0.009 | 0 |
| ADE17 | -0.021 | 0.010 | -0.006 | 0 |
| STR2 | 0.007 | -0.001 | 0.003 | 0 |
| GID8 | 0.008 | 0.008 | 0.008 | 0 |
| SRL3 | 0.004 | 0.002 | 0.003 | 0 |
| APL4 | 0.010 | 0.019 | 0.014 | 0 |
| COX19 | -0.002 | -0.002 | -0.002 | 0 |
| TMA7 | -0.002 | -0.013 | -0.008 | 0 |
| YDR026C | -0.008 | 0.000 | -0.004 | 0 |
| TSR2 | 0.000 | 0.006 | 0.003 | 0 |
| NRG1 | -0.001 | 0.013 | 0.006 | 0 |
| HMG2 | 0.003 | 0.007 | 0.005 | 0 |
| SSH1 | 0.010 | 0.030 | 0.020 | 0 |
| YBR300C | -0.001 | 0.003 | 0.001 | 0 |
| BIO3 | -0.001 | -0.004 | -0.002 | 0 |
| RBK1 | 0.004 | 0.002 | 0.003 | 0 |
| SPC72 | -0.016 | 0.016 | 0.000 | 0 |
| IMG2 | 0.011 | 0.000 | 0.006 | 0 |
| YJL016W | 0.000 | -0.003 | -0.001 | 0 |
| YJR080C | -0.004 | -0.001 | -0.002 | 0 |
| SPC1 | 0.002 | 0.001 | 0.002 | 0 |
| YJR100C | -0.002 | -0.011 | -0.007 | 0 |
| YJR026W | 0.006 | 0.011 | 0.008 | 0 |
| IBA57 | 0.002 | -0.008 | -0.003 | 0 |
| BFA1 | -0.003 | -0.017 | -0.010 | 0 |
| BAT2 | 0.002 | -0.004 | -0.001 | 0 |
| GET3 | -0.009 | -0.006 | -0.008 | 0 |
| YOR300W | -0.009 | -0.015 | -0.012 | 0 |
| YDL118W | 0.038 | 0.042 | 0.040 | 1 |
| YDL206W | -0.003 | 0.004 | 0.000 | 0 |
| LYS21 | -0.006 | 0.000 | -0.003 | 0 |
| SHS1 | -0.031 | -0.007 | -0.019 | 0 |
| BUD30 | 0.015 | 0.011 | 0.013 | 0 |
| ADY3 | 0.004 | 0.002 | 0.003 | 0 |
| YDL172C | -0.001 | -0.006 | -0.003 | 0 |
| YDR010C | -0.018 | -0.011 | -0.014 | 0 |
| RPL41A | -0.003 | 0.006 | 0.002 | 0 |
| YLR422W | -0.006 | -0.027 | -0.016 | 0 |
| VPS54 | 0.004 | 0.016 | 0.010 | 0 |
| ECM30 | -0.061 | -0.101 | -0.081 | -1 |
| BAP3 | -0.015 | -0.008 | -0.011 | 0 |
| SST2 | -0.010 | -0.020 | -0.015 | 0 |
| YBR284W | -0.005 | -0.002 | -0.004 | 0 |
| ΥMR158C-B | 0.006 | 0.003 | 0.005 | 0 |
| ΥCL001W-A | -0.009 | -0.006 | -0.008 | 0 |
| MNT4 | -0.005 | 0.003 | -0.001 | 0 |
| PHO87 | -0.003 | -0.012 | -0.008 | 0 |
| ACS1 | 0.000 | 0.004 | 0.002 | 0 |
| SOL2 | -0.005 | -0.004 | -0.005 | 0 |
| YJL017W | -0.005 | 0.006 | 0.001 | 0 |
| SOD1 | 0.003 | -0.025 | -0.011 | 0 |
| YJR030C | 0.002 | -0.002 | 0.000 | 0 |

| | | | | |
|---|---|---|---|---|
| YJR124C | 0.018 | 0.022 | 0.020 | 0 |
| YJR054W | 0.001 | -0.012 | -0.006 | 0 |
| DAN1 | -0.009 | 0.002 | -0.004 | 0 |
| DUN1 | 0.017 | 0.026 | 0.022 | 0 |
| YDL119C | 0.054 | 0.054 | 0.054 | 1 |
| UGA4 | -0.003 | -0.008 | -0.006 | 0 |
| YDL133W | -0.036 | -0.039 | -0.038 | -1 |
| GCS1 | -0.001 | 0.002 | 0.000 | 0 |
| MSH5 | -0.006 | -0.002 | -0.004 | 0 |
| LRG1 | 0.006 | 0.006 | 0.006 | 0 |
| YDL173W | 0.000 | -0.012 | -0.006 | 0 |
| SCH9 | 0.008 | -0.003 | 0.003 | 0 |
| ATG17 | 0.021 | 0.010 | 0.016 | 0 |
| REG1 | 0.042 | 0.034 | 0.038 | 1 |
| DIF1 | -0.006 | -0.002 | -0.004 | 0 |
| YAL064C-A | 0.003 | -0.005 | -0.001 | 0 |
| RIF2 | 0.000 | 0.003 | 0.002 | 0 |
| YBR285W | -0.007 | -0.011 | -0.009 | 0 |
| FRE4 | -0.004 | -0.002 | -0.003 | 0 |
| YCR043C | 0.007 | 0.001 | 0.004 | 0 |
| YAL058C-A | 0.010 | 0.027 | 0.019 | 0 |
| YCR076C | -0.006 | -0.009 | -0.007 | 0 |
| BBC1 | 0.000 | -0.001 | 0.000 | 0 |
| CSN12 | -0.009 | -0.007 | -0.008 | 0 |
| MET3 | 0.013 | 0.000 | 0.006 | 0 |
| ECM27 | -0.019 | -0.025 | -0.022 | 0 |
| GEA1 | 0.006 | 0.003 | 0.005 | 0 |
| ENT3 | 0.003 | 0.013 | 0.008 | 0 |
| HIT1 | 0.005 | 0.002 | 0.004 | 0 |
| PGU1 | -0.009 | 0.004 | -0.002 | 0 |
| QRI7 | 0.019 | 0.023 | 0.021 | 0 |
| YOR309C | 0.001 | 0.006 | 0.003 | 0 |
| YDL121C | -0.004 | -0.014 | -0.009 | 0 |
| YDL211C | 0.007 | 0.001 | 0.004 | 0 |
| PPH21 | -0.005 | -0.002 | -0.004 | 0 |
| HO | -0.003 | -0.010 | -0.006 | 0 |
| CLB3 | -0.002 | 0.001 | 0.000 | 0 |
| YDL241W | -0.005 | -0.009 | -0.007 | 0 |
| DLD1 | -0.003 | -0.005 | -0.004 | 0 |
| SNQ2 | -0.039 | -0.025 | -0.032 | -1 |
| YDL186W | 0.007 | 0.003 | 0.005 | 0 |
| TUS1 | 0.005 | -0.008 | -0.002 | 0 |
| YDR029W | -0.009 | 0.003 | -0.003 | 0 |
| CAR2 | 0.001 | 0.006 | 0.004 | 0 |
| SCS22 | -0.004 | -0.002 | -0.003 | 0 |
| FMP27 | -0.003 | 0.001 | -0.001 | 0 |
| APE3 | 0.001 | 0.000 | 0.001 | 0 |
| HTL1 | 0.013 | 0.013 | 0.013 | 0 |
| YNR061C | 0.006 | -0.005 | 0.000 | 0 |
| YCR045C | -0.011 | -0.006 | -0.009 | 0 |
| FLO1 | 0.001 | 0.003 | 0.002 | 0 |
| PAT1 | 0.003 | 0.001 | 0.002 | 0 |
| YJL021C | 0.004 | 0.007 | 0.005 | 0 |

| | | | | |
|---|---|---|---|---|
| YJR087W | 0.029 | 0.038 | 0.033 | 1 |
| YJR011C | 0.017 | 0.015 | 0.016 | 0 |
| YJR107W | -0.001 | -0.018 | -0.010 | 0 |
| RAV1 | -0.070 | -0.165 | -0.117 | -1 |
| VPS70 | -0.006 | 0.009 | 0.001 | 0 |
| YJR056C | -0.009 | -0.004 | -0.006 | 0 |
| BYE1 | 0.037 | -0.012 | 0.013 | 0 |
| PHO2 | -0.008 | -0.017 | -0.012 | 0 |
| RPL35A | 0.014 | 0.026 | 0.020 | 0 |
| NOP6 | -0.006 | 0.003 | -0.002 | 0 |
| YDL134C-A | -0.002 | 0.000 | -0.001 | 0 |
| SSB1 | -0.008 | -0.008 | -0.008 | 0 |
| YDL156W | 0.001 | 0.005 | 0.003 | 0 |
| YDL242W | -0.002 | -0.008 | -0.005 | 0 |
| AIR2 | 0.009 | 0.006 | 0.008 | 0 |
| RAD61 | -0.001 | 0.007 | 0.003 | 0 |
| YDL187C | -0.001 | 0.005 | 0.002 | 0 |
| YLR426W | -0.018 | -0.042 | -0.030 | 0 |
| RAD28 | -0.001 | -0.001 | -0.001 | 0 |
| RPS1A | 0.004 | 0.018 | 0.011 | 0 |
| FMP21 | 0.001 | 0.004 | 0.003 | 0 |
| YLR456W | -0.007 | -0.005 | -0.006 | 0 |
| BSD2 | -0.011 | -0.017 | -0.014 | 0 |
| SLM5 | -0.001 | 0.008 | 0.004 | 0 |
| YNR062C | -0.002 | 0.002 | 0.000 | 0 |
| YCR049C | -0.007 | 0.000 | -0.004 | 0 |
| YCL006C | -0.011 | -0.006 | -0.009 | 0 |
| PTC6 | 0.005 | 0.006 | 0.006 | 0 |
| YJL022W | -0.011 | -0.001 | -0.006 | 0 |
| JSN1 | -0.006 | -0.002 | -0.004 | 0 |
| TMA22 | -0.007 | 0.005 | -0.001 | 0 |
| CPA2 | 0.016 | 0.024 | 0.020 | 0 |
| RAD26 | -0.005 | -0.018 | -0.012 | 0 |
| MNS1 | -0.002 | 0.007 | 0.002 | 0 |
| APS2 | -0.009 | 0.001 | -0.004 | 0 |
| YKL030W | 0.003 | 0.002 | 0.002 | 0 |
| MSS2 | 0.008 | -0.060 | -0.026 | 0 |
| ARF1 | 0.002 | 0.004 | 0.003 | 0 |
| UBP1 | 0.001 | 0.014 | 0.007 | 0 |
| PRR2 | -0.347 | -0.002 | -0.174 | 0 |
| RDI1 | -0.020 | -0.027 | -0.024 | 0 |
| PTP1 | 0.005 | 0.001 | 0.003 | 0 |
| YDL157C | 0.012 | 0.010 | 0.011 | 0 |
| AAD4 | -0.007 | 0.002 | -0.002 | 0 |
| YDL176W | -0.006 | -0.017 | -0.011 | 0 |
| YDR015C | 0.002 | 0.004 | 0.003 | 0 |
| PPH22 | 0.002 | 0.004 | 0.003 | 0 |
| MAG2 | 0.005 | -0.001 | 0.002 | 0 |
| MIC14 | 0.003 | 0.010 | 0.006 | 0 |
| ECM7 | -0.004 | 0.006 | 0.001 | 0 |
| YBR271W | 0.002 | 0.000 | 0.001 | 0 |
| YLR460C | -0.003 | -0.002 | -0.002 | 0 |
| CTP1 | -0.009 | 0.002 | -0.003 | 0 |

| | | | | |
|---|---|---|---|---|
| YMR194C-A | -0.002 | -0.003 | -0.002 | 0 |
| PMP1 | 0.002 | -0.004 | -0.001 | 0 |
| YNR063W | 0.001 | 0.004 | 0.003 | 0 |
| YCR050C | 0.001 | 0.003 | 0.002 | 0 |
| YCL022C | 0.000 | -0.002 | -0.001 | 0 |
| SRB8 | 0.108 | 0.080 | 0.094 | 1 |
| PET130 | 0.011 | 0.000 | 0.006 | 0 |
| IME1 | -0.012 | -0.004 | -0.008 | 0 |
| YJR015W | 0.003 | 0.002 | 0.003 | 0 |
| RSM7 | 0.004 | -0.005 | 0.000 | 0 |
| HUL4 | 0.003 | 0.012 | 0.007 | 0 |
| XPT1 | 0.001 | -0.004 | -0.001 | 0 |
| PTK2 | 0.002 | 0.012 | 0.007 | 0 |
| SPE4 | -0.006 | -0.001 | -0.004 | 0 |
| YDL109C | 0.005 | 0.005 | 0.005 | 0 |
| ASF2 | -0.006 | -0.007 | -0.007 | 0 |
| SNA4 | 0.003 | 0.000 | 0.001 | 0 |
| GDH2 | 0.002 | 0.009 | 0.006 | 0 |
| RPL35B | 0.011 | 0.022 | 0.017 | 0 |
| BRE4 | 0.003 | 0.004 | 0.003 | 0 |
| STE7 | 0.010 | 0.004 | 0.007 | 0 |
| NTH1 | -0.001 | 0.005 | 0.002 | 0 |
| YDL177C | 0.000 | -0.002 | -0.001 | 0 |
| KCS1 | -0.066 | -0.060 | -0.063 | -1 |
| RBS1 | -0.007 | 0.002 | -0.002 | 0 |
| YLR428C | -0.009 | 0.003 | -0.003 | 0 |
| PST2 | -0.002 | -0.005 | -0.004 | 0 |
| YLR444C | -0.001 | -0.006 | -0.004 | 0 |
| UBX7 | 0.000 | 0.006 | 0.003 | 0 |
| PAU4 | -0.018 | -0.001 | -0.009 | 0 |
| YBR292C | -0.021 | -0.006 | -0.013 | 0 |
| YMR326C | -0.001 | 0.000 | 0.000 | 0 |
| YCR025C | -0.004 | -0.002 | -0.003 | 0 |
| YNR064C | -0.007 | -0.015 | -0.011 | 0 |
| YCR051W | -0.003 | -0.002 | -0.002 | 0 |
| AHC2 | -0.010 | -0.016 | -0.013 | 0 |
| APS3 | -0.158 | -0.070 | -0.114 | -1 |
| RPL43B | -0.010 | 0.001 | -0.005 | 0 |
| YJR018W | 0.060 | 0.050 | 0.055 | 1 |
| YJR116W | 0.000 | 0.014 | 0.007 | 0 |
| POL32 | 0.041 | 0.032 | 0.037 | 1 |
| SGM1 | -0.010 | -0.016 | -0.013 | 0 |
| CBF1 | -0.003 | -0.006 | -0.004 | 0 |
| GAS2 | 0.002 | -0.004 | -0.001 | 0 |
| TMA17 | 0.001 | 0.009 | 0.005 | 0 |
| GGC1 | 0.029 | -0.013 | 0.008 | 0 |
| YDL124W | -0.005 | -0.001 | -0.003 | 0 |
| RRI1 | -0.002 | 0.002 | 0.000 | 0 |
| ARF2 | -0.004 | -0.011 | -0.007 | 0 |
| OST4 | 0.001 | 0.003 | 0.002 | 0 |
| ENT1 | 0.000 | 0.005 | 0.003 | 0 |
| RCR2 | -0.011 | 0.004 | -0.004 | 0 |
| DLD2 | -0.004 | -0.001 | -0.003 | 0 |

| | | | | |
|---|---|---|---|---|
| YDR018C | -0.007 | 0.001 | -0.003 | 0 |
| UFD2 | 0.187 | 0.190 | 0.188 | 1 |
| CRN1 | -0.008 | -0.001 | -0.005 | 0 |
| MRH1 | 0.001 | -0.001 | 0.000 | 0 |
| YLR445W | -0.006 | -0.001 | -0.004 | 0 |
| CHK1 | 0.003 | 0.007 | 0.005 | 0 |
| VBA2 | 0.001 | -0.010 | -0.005 | 0 |
| HUB1 | -0.008 | -0.004 | -0.006 | 0 |
| NPP1 | 0.000 | 0.001 | 0.000 | 0 |
| YNR065C | -0.002 | 0.003 | 0.001 | 0 |
| YIH1 | 0.003 | 0.000 | 0.002 | 0 |
| ATG22 | -0.007 | -0.005 | -0.006 | 0 |
| YCR085W | 0.001 | -0.003 | -0.001 | 0 |
| VPS53 | 0.010 | 0.027 | 0.019 | 0 |
| SFC1 | -0.009 | 0.005 | -0.002 | 0 |
| TES1 | 0.012 | 0.008 | 0.010 | 0 |
| STE24 | -0.027 | -0.015 | -0.021 | 0 |
| CYC1 | 0.014 | 0.024 | 0.019 | 0 |
| HOM6 | -0.005 | -0.020 | -0.012 | 0 |
| YJR061W | -0.007 | -0.008 | -0.008 | 0 |
| ERV41 | -0.015 | -0.020 | -0.017 | 0 |
| TRM3 | 0.005 | -0.001 | 0.002 | 0 |
| YDL199C | -0.005 | 0.002 | -0.002 | 0 |
| HNT1 | 0.000 | 0.007 | 0.004 | 0 |
| YDL218W | 0.004 | 0.001 | 0.003 | 0 |
| RGT2 | -0.005 | -0.006 | -0.006 | 0 |
| YDL233W | -0.270 | -0.011 | -0.140 | 0 |
| YDL162C | 0.005 | -0.002 | 0.001 | 0 |
| RAD57 | 0.022 | 0.015 | 0.019 | 0 |
| PCL9 | -0.001 | -0.001 | -0.001 | 0 |
| GCV1 | 0.000 | 0.016 | 0.008 | 0 |
| ATG23 | -0.002 | 0.001 | -0.001 | 0 |
| LYS14 | -0.026 | -0.030 | -0.028 | -1 |
| YLR446W | 0.001 | 0.001 | 0.001 | 0 |
| YBR277C | 0.027 | 0.018 | 0.022 | 0 |
| YML010C-B | 0.023 | 0.018 | 0.020 | 0 |
| PCA1 | -0.010 | 0.000 | -0.005 | 0 |
| LYS9 | 0.024 | 0.016 | 0.020 | 0 |
| RHB1 | 0.007 | 0.015 | 0.011 | 0 |
| YNR066C | -0.003 | 0.000 | -0.001 | 0 |
| YCR061W | -0.002 | -0.005 | -0.004 | 0 |
| FYV5 | 0.007 | 0.006 | 0.007 | 0 |
| CSM1 | -0.004 | -0.015 | -0.010 | 0 |
| HAM1 | 0.003 | -0.008 | -0.002 | 0 |
| AVT1 | -0.001 | 0.008 | 0.004 | 0 |
| YJR096W | -0.012 | -0.001 | -0.007 | 0 |
| YJR020W | 0.006 | 0.001 | 0.004 | 0 |
| ILM1 | 0.017 | 0.023 | 0.020 | 0 |
| UTR1 | -0.010 | -0.006 | -0.008 | 0 |
| HIR3 | -0.004 | 0.002 | -0.001 | 0 |
| NTA1 | 0.000 | -0.012 | -0.006 | 0 |
| ITT1 | 0.009 | 0.001 | 0.005 | 0 |
| ATG20 | -0.001 | 0.006 | 0.003 | 0 |

| | | | | |
|---|---|---|---|---|
| MGT1 | -0.012 | -0.011 | -0.011 | 0 |
| PCL2 | -0.004 | -0.005 | -0.005 | 0 |
| DTD1 | -0.006 | 0.006 | 0.000 | 0 |
| CRD1 | 0.026 | 0.040 | 0.033 | 1 |
| GYP7 | -0.001 | 0.001 | 0.000 | 0 |
| SFA1 | 0.002 | 0.000 | 0.001 | 0 |
| MAF1 | 0.002 | 0.004 | 0.003 | 0 |
| YDL180W | -0.002 | -0.005 | -0.003 | 0 |
| YDR020C | -0.001 | 0.004 | 0.002 | 0 |
| IMD3 | -0.011 | 0.009 | -0.001 | 0 |
| ARO3 | 0.003 | 0.006 | 0.004 | 0 |
| VMA6 | -0.023 | -0.039 | -0.031 | -1 |
| DPB3 | 0.027 | 0.025 | 0.026 | 1 |
| UNG1 | 0.000 | -0.007 | -0.004 | 0 |
| PHO89 | -0.011 | -0.004 | -0.007 | 0 |
| BRE5 | 0.030 | 0.034 | 0.032 | 1 |
| FEN2 | 0.077 | 0.010 | 0.044 | 0 |
| DSE4 | 0.004 | 0.011 | 0.008 | 0 |
| HCM1 | -0.001 | -0.004 | -0.002 | 0 |
| YCL074W | -0.003 | 0.002 | -0.001 | 0 |
| YCR087C-A | -0.006 | -0.005 | -0.006 | 0 |
| LIA1 | -0.035 | -0.010 | -0.023 | 0 |
| APL1 | -0.006 | -0.003 | -0.005 | 0 |
| JJJ3 | 0.013 | 0.007 | 0.010 | 0 |
| REC107 | -0.010 | -0.009 | -0.009 | 0 |
| JHD2 | -0.002 | -0.004 | -0.003 | 0 |
| ISY1 | 0.005 | -0.010 | -0.002 | 0 |
| YJR142W | -0.018 | -0.017 | -0.017 | 0 |
| RPA12 | -0.004 | -0.012 | -0.008 | 0 |
| TCB3 | -0.015 | -0.022 | -0.019 | 0 |
| YDL114W | -0.010 | -0.008 | -0.009 | 0 |
| TRM8 | 0.006 | 0.011 | 0.008 | 0 |
| VCX1 | 0.007 | 0.001 | 0.004 | 0 |
| FMP45 | -0.004 | -0.002 | -0.003 | 0 |
| YDL144C | 0.000 | 0.006 | 0.003 | 0 |
| PHO13 | 0.009 | 0.014 | 0.011 | 0 |
| UGX2 | 0.000 | -0.004 | -0.002 | 0 |
| SOK1 | 0.001 | -0.002 | -0.001 | 0 |
| INH1 | 0.006 | 0.002 | 0.004 | 0 |
| CIS1 | 0.005 | 0.013 | 0.009 | 0 |
| CNA1 | -0.004 | -0.007 | -0.005 | 0 |
| EHD3 | 0.004 | 0.000 | 0.002 | 0 |
| RPL6B | 0.027 | 0.046 | 0.037 | 1 |
| DUG2 | -0.015 | -0.008 | -0.011 | 0 |
| ATP18 | 0.004 | 0.004 | 0.004 | 0 |
| MAL33 | -0.011 | -0.008 | -0.010 | 0 |
| BIO5 | -0.001 | 0.001 | 0.000 | 0 |
| RPS14A | 0.001 | 0.005 | 0.003 | 0 |
| YNR068C | 0.010 | -0.003 | 0.004 | 0 |
| RAD18 | 0.008 | -0.003 | 0.003 | 0 |
| YCL075W | 0.001 | 0.006 | 0.004 | 0 |
| YCR087W | -0.008 | 0.001 | -0.004 | 0 |
| MOG1 | 0.007 | 0.002 | 0.005 | 0 |

| | | | | |
|---|---|---|---|---|
| YJR008W | 0.002 | 0.001 | 0.001 | 0 |
| YJR098C | -0.002 | 0.001 | 0.000 | 0 |
| MDE1 | -0.004 | 0.003 | 0.000 | 0 |
| YJR120W | 0.009 | 0.014 | 0.011 | 0 |
| OSM1 | -0.013 | -0.006 | -0.010 | 0 |
| MGM101 | 0.025 | 0.033 | 0.029 | 1 |
| OPI6 | -0.010 | -0.001 | -0.006 | 0 |
| GAT2 | 0.002 | 0.002 | 0.002 | 0 |
| NUP84 | 0.052 | 0.029 | 0.041 | 1 |
| ACK1 | -0.002 | -0.002 | -0.002 | 0 |
| YDL129W | 0.000 | -0.002 | -0.001 | 0 |
| HBT1 | -0.007 | 0.006 | -0.001 | 0 |
| LDB17 | -0.010 | -0.012 | -0.011 | 0 |
| YDL237W | -0.003 | -0.006 | -0.004 | 0 |
| UGA3 | -0.012 | 0.004 | -0.004 | 0 |
| YDR008C | 0.002 | 0.005 | 0.004 | 0 |
| LYS20 | 0.011 | 0.015 | 0.013 | 0 |
| FYV1 | -0.009 | -0.010 | -0.009 | 0 |
| YOR325W | -0.030 | -0.008 | -0.019 | 0 |
| YLR434C | -0.009 | -0.006 | -0.007 | 0 |
| YDR042C | -0.009 | -0.013 | -0.011 | 0 |
| FPR4 | -0.002 | 0.007 | 0.003 | 0 |
| MRPL27 | 0.009 | 0.019 | 0.014 | 0 |
| SAM37 | 0.064 | 0.047 | 0.056 | 1 |
| MAL31 | -0.005 | -0.003 | -0.004 | 0 |
| BIO4 | 0.004 | -0.001 | 0.002 | 0 |
| FEN1 | 0.048 | 0.040 | 0.044 | 1 |
| CYS3 | -0.008 | -0.012 | -0.010 | 0 |
| ATG15 | -0.007 | -0.003 | -0.005 | 0 |
| YCL076W | 0.007 | 0.009 | 0.008 | 0 |
| YJL007C | -0.002 | -0.014 | -0.008 | 0 |
| MIR1 | 0.013 | 0.015 | 0.014 | 0 |
| TDH2 | 0.012 | 0.012 | 0.012 | 0 |
| YUH1 | 0.002 | 0.014 | 0.008 | 0 |
| BNA1 | -0.001 | -0.002 | -0.001 | 0 |
| ATP2 | -0.001 | 0.001 | 0.000 | 0 |
| RAD7 | -0.005 | -0.014 | -0.010 | 0 |
| RPS4A | 0.007 | 0.020 | 0.013 | 0 |
| BUG1 | 0.005 | 0.003 | 0.004 | 0 |
| HOT1 | -0.006 | -0.021 | -0.013 | 0 |
| CYK3 | 0.008 | 0.001 | 0.004 | 0 |
| RTN2 | -0.013 | 0.000 | -0.007 | 0 |
| RPP1B | 0.017 | 0.027 | 0.022 | 0 |
| WHI4 | -0.004 | 0.000 | -0.002 | 0 |
| ATG9 | -0.004 | 0.001 | -0.001 | 0 |
| GUD1 | 0.004 | 0.006 | 0.005 | 0 |
| GLT1 | -0.011 | -0.003 | -0.007 | 0 |
| GAL3 | 0.002 | 0.004 | 0.003 | 0 |
| YDL183C | -0.010 | 0.002 | -0.004 | 0 |
| RPS11A | -0.001 | -0.006 | -0.004 | 0 |
| YOR333C | 0.004 | 0.001 | 0.003 | 0 |
| YOR345C | 0.007 | -0.039 | -0.016 | 0 |
| HOS4 | -0.008 | -0.004 | -0.006 | 0 |

| | | | | |
|---|---|---|---|---|
| PUB1 | 0.007 | 0.023 | 0.015 | 0 |
| ASG1 | -0.005 | -0.009 | -0.007 | 0 |
| SIW14 | 0.025 | 0.037 | 0.031 | 1 |
| RPL40A | 0.006 | 0.001 | 0.003 | 0 |
| CIT1 | -0.012 | -0.014 | -0.013 | 0 |
| YIL163C | 0.008 | -0.009 | 0.000 | 0 |
| YNR014W | -0.011 | -0.007 | -0.009 | 0 |
| IST3 | 0.020 | 0.032 | 0.026 | 0 |
| ALG12 | -0.005 | 0.004 | -0.001 | 0 |
| COX5A | 0.005 | -0.018 | -0.007 | 0 |
| YNR048W | 0.000 | 0.002 | 0.001 | 0 |
| NIS1 | -0.019 | -0.018 | -0.019 | 0 |
| YBL107C | 0.004 | 0.003 | 0.004 | 0 |
| OCA1 | 0.016 | 0.030 | 0.023 | 0 |
| GRX7 | 0.015 | -0.001 | 0.007 | 0 |
| CPT1 | -0.006 | -0.010 | -0.008 | 0 |
| YBR027C | -0.004 | -0.012 | -0.008 | 0 |
| MDM34 | 0.024 | 0.013 | 0.019 | 0 |
| QDR3 | 0.003 | 0.000 | 0.001 | 0 |
| VOA1 | -0.004 | 0.001 | -0.001 | 0 |
| MUM2 | -0.009 | -0.005 | -0.007 | 0 |
| TRM13 | 0.004 | 0.006 | 0.005 | 0 |
| TAT1 | 0.001 | -0.011 | -0.005 | 0 |
| AAH1 | -0.006 | -0.009 | -0.007 | 0 |
| YKL215C | -0.004 | -0.003 | -0.003 | 0 |
| DMA2 | 0.009 | 0.003 | 0.006 | 0 |
| YKR041W | -0.004 | -0.008 | -0.006 | 0 |
| YNL057W | -0.006 | 0.015 | 0.005 | 0 |
| TBS1 | -0.015 | -0.029 | -0.022 | 0 |
| YOR366W | 0.010 | -0.004 | 0.003 | 0 |
| SDP1 | -0.009 | 0.005 | -0.002 | 0 |
| ARK1 | -0.010 | -0.001 | -0.006 | 0 |
| CSM2 | -0.003 | -0.009 | -0.006 | 0 |
| YNL034W | 0.002 | -0.008 | -0.003 | 0 |
| MLP2 | 0.008 | -0.001 | 0.004 | 0 |
| ATO2 | 0.004 | 0.002 | 0.003 | 0 |
| NIT1 | 0.017 | 0.003 | 0.010 | 0 |
| SMM1 | -0.024 | -0.014 | -0.019 | 0 |
| YIR007W | 0.003 | 0.007 | 0.005 | 0 |
| SSK2 | -0.001 | -0.008 | -0.004 | 0 |
| OCA2 | 0.015 | 0.017 | 0.016 | 0 |
| MSO1 | -0.005 | 0.001 | -0.002 | 0 |
| TPM1 | 0.038 | 0.015 | 0.026 | 0 |
| MNN2 | -0.001 | -0.010 | -0.006 | 0 |
| RPS9B | -0.001 | 0.010 | 0.005 | 0 |
| YBR028C | 0.000 | 0.005 | 0.002 | 0 |
| MSP1 | -0.009 | -0.002 | -0.005 | 0 |
| TCM62 | -0.008 | 0.008 | 0.000 | 0 |
| YGR110W | 0.001 | 0.000 | 0.000 | 0 |
| UBP14 | 0.050 | 0.043 | 0.046 | 1 |
| YAK1 | 0.002 | 0.000 | 0.001 | 0 |
| YBR071W | -0.011 | -0.012 | -0.011 | 0 |
| THO2 | 0.018 | 0.006 | 0.012 | 0 |

| | | | | |
|---|---|---|---|---|
| FRE2 | -0.002 | 0.002 | 0.000 | 0 |
| YNL108C | -0.005 | -0.017 | -0.011 | 0 |
| PET10 | 0.013 | 0.008 | 0.011 | 0 |
| YIL127C | -0.003 | -0.003 | -0.003 | 0 |
| YOR379C | 0.020 | -0.021 | -0.001 | 0 |
| POR2 | 0.001 | -0.003 | -0.001 | 0 |
| HDA1 | -0.029 | -0.042 | -0.036 | -1 |
| RPL16A | 0.006 | 0.010 | 0.008 | 0 |
| YNL035C | -0.003 | 0.004 | 0.000 | 0 |
| YIL152W | -0.008 | 0.001 | -0.003 | 0 |
| YNR004W | -0.004 | 0.021 | 0.008 | 0 |
| YIL165C | 0.002 | -0.005 | -0.001 | 0 |
| YNR018W | -0.003 | 0.000 | -0.002 | 0 |
| MSL1 | -0.007 | 0.010 | 0.001 | 0 |
| PPG1 | 0.020 | 0.001 | 0.010 | 0 |
| AQR1 | 0.001 | 0.001 | 0.001 | 0 |
| YBL095W | -0.012 | 0.000 | -0.006 | 0 |
| EOS1 | -0.008 | -0.003 | -0.006 | 0 |
| NTH2 | -0.005 | -0.044 | -0.024 | 0 |
| YNL100W | 0.008 | 0.021 | 0.015 | 0 |
| YBR016W | 0.009 | -0.039 | -0.015 | 0 |
| OCA4 | 0.015 | 0.027 | 0.021 | 0 |
| RKM3 | -0.012 | -0.006 | -0.009 | 0 |
| GSC2 | -0.005 | -0.004 | -0.004 | 0 |
| GIP1 | -0.004 | 0.000 | -0.002 | 0 |
| YGR117C | 0.009 | -0.001 | 0.004 | 0 |
| AKL1 | 0.027 | 0.006 | 0.017 | 0 |
| HSP26 | -0.009 | -0.008 | -0.009 | 0 |
| EAF7 | -0.005 | -0.011 | -0.008 | 0 |
| TOF2 | 0.001 | -0.022 | -0.011 | 0 |
| AVT4 | -0.005 | 0.007 | 0.001 | 0 |
| YSR3 | -0.003 | 0.001 | -0.001 | 0 |
| YNL058C | 0.000 | 0.016 | 0.008 | 0 |
| PEX32 | 0.001 | 0.012 | 0.007 | 0 |
| DOM34 | -0.015 | -0.030 | -0.022 | 0 |
| HIS5 | -0.015 | 0.014 | -0.001 | 0 |
| YNL022C | -0.002 | 0.000 | -0.001 | 0 |
| FLX1 | -0.005 | -0.003 | -0.004 | 0 |
| IDH1 | 0.001 | 0.032 | 0.017 | 0 |
| RRD1 | -0.009 | -0.027 | -0.018 | 0 |
| YNR005C | -0.018 | -0.042 | -0.030 | 0 |
| YIL166C | -0.004 | -0.008 | -0.006 | 0 |
| ARE2 | 0.005 | -0.005 | 0.000 | 0 |
| GAT4 | -0.001 | 0.004 | 0.001 | 0 |
| SOL1 | 0.012 | -0.004 | 0.004 | 0 |
| SUN4 | 0.017 | 0.002 | 0.009 | 0 |
| YBL096C | 0.006 | -0.004 | 0.001 | 0 |
| PMS1 | 0.010 | 0.000 | 0.005 | 0 |
| LEU4 | 0.009 | 0.016 | 0.012 | 0 |
| GAL7 | 0.001 | -0.003 | -0.001 | 0 |
| YCR102W-A | -0.009 | -0.023 | -0.016 | 0 |
| RPL4A | 0.002 | 0.004 | 0.003 | 0 |
| ORM1 | -0.008 | -0.013 | -0.011 | 0 |

| | | | | |
|---|---|---|---|---|
| ZTA1 | -0.004 | 0.006 | 0.001 | 0 |
| KEL2 | 0.003 | 0.000 | 0.001 | 0 |
| TRM7 | -0.009 | 0.001 | -0.004 | 0 |
| YJL160C | -0.005 | 0.000 | -0.003 | 0 |
| RDH54 | 0.002 | -0.005 | -0.001 | 0 |
| FPR1 | -0.007 | 0.000 | -0.004 | 0 |
| IRS4 | -0.001 | -0.005 | -0.003 | 0 |
| RAS2 | -0.012 | -0.009 | -0.010 | 0 |
| VAC7 | 0.004 | 0.015 | 0.009 | 0 |
| SSE2 | 0.007 | -0.010 | -0.001 | 0 |
| HRB1 | 0.002 | 0.012 | 0.007 | 0 |
| PRM5 | 0.010 | -0.004 | 0.003 | 0 |
| FAP1 | -0.011 | -0.015 | -0.013 | 0 |
| VHS2 | -0.004 | -0.020 | -0.012 | 0 |
| YNL040W | 0.001 | -0.003 | -0.001 | 0 |
| IMP2' | 0.020 | 0.006 | 0.013 | 0 |
| SDL1 | 0.003 | -0.001 | 0.001 | 0 |
| ATP23 | 0.014 | 0.019 | 0.016 | 0 |
| YIR014W | 0.008 | -0.001 | 0.004 | 0 |
| MRPS12 | -0.019 | 0.004 | -0.008 | 0 |
| RPL9B | -0.001 | 0.000 | -0.001 | 0 |
| BNA4 | 0.007 | -0.013 | -0.003 | 0 |
| SAL1 | -0.009 | 0.011 | 0.001 | 0 |
| RCR1 | 0.000 | -0.001 | -0.001 | 0 |
| YNL105W | 0.005 | -0.003 | 0.001 | 0 |
| GAL10 | -0.008 | -0.007 | -0.008 | 0 |
| RPL41B | 0.004 | -0.005 | 0.000 | 0 |
| YBR032W | -0.012 | 0.007 | -0.003 | 0 |
| KSS1 | 0.002 | -0.017 | -0.008 | 0 |
| FMP23 | -0.002 | 0.008 | 0.003 | 0 |
| PEX21 | 0.002 | 0.008 | 0.005 | 0 |
| YBR062C | 0.019 | 0.008 | 0.013 | 0 |
| YBR074W | 0.001 | 0.002 | 0.001 | 0 |
| YJL213W | -0.011 | -0.006 | -0.009 | 0 |
| YNL134C | 0.000 | 0.005 | 0.003 | 0 |
| YKR023W | 0.005 | 0.007 | 0.006 | 0 |
| APP1 | 0.015 | 0.008 | 0.012 | 0 |
| ERP1 | -0.084 | -0.049 | -0.066 | -1 |
| SGA1 | 0.001 | 0.010 | 0.005 | 0 |
| BIT2 | 0.007 | 0.008 | 0.007 | 0 |
| MRP7 | 0.021 | 0.020 | 0.021 | 1 |
| RPI1 | -0.029 | -0.039 | -0.034 | -1 |
| YNL024C | 0.002 | 0.010 | 0.006 | 0 |
| TMA108 | -0.005 | -0.012 | -0.008 | 0 |
| COG6 | -0.008 | -0.009 | -0.009 | 0 |
| GUT2 | -0.003 | 0.001 | -0.001 | 0 |
| VPS27 | -0.047 | -0.040 | -0.043 | -1 |
| YIL168W | 0.000 | 0.001 | 0.001 | 0 |
| YNR021W | 0.002 | -0.012 | -0.005 | 0 |
| YIR016W | -0.010 | 0.005 | -0.003 | 0 |
| RSM19 | 0.003 | -0.009 | -0.003 | 0 |
| FKH2 | -0.037 | -0.087 | -0.062 | -1 |
| ATP1 | -0.010 | -0.079 | -0.045 | 0 |

| | | | | |
|---|---|---|---|---|
| MKT1 | -0.002 | -0.005 | -0.004 | 0 |
| UGA2 | 0.009 | 0.006 | 0.007 | 0 |
| INP52 | -0.003 | 0.018 | 0.008 | 0 |
| GAL1 | 0.001 | 0.000 | 0.000 | 0 |
| TGL2 | 0.000 | -0.007 | -0.004 | 0 |
| EDS1 | -0.008 | -0.002 | -0.005 | 0 |
| YGR050C | 0.008 | -0.004 | 0.002 | 0 |
| RPS11B | -0.003 | 0.015 | 0.006 | 0 |
| SOL4 | -0.007 | -0.011 | -0.009 | 0 |
| YBR075W | 0.003 | -0.002 | 0.001 | 0 |
| CWP2 | -0.002 | 0.002 | 0.000 | 0 |
| FYV6 | 0.079 | 0.050 | 0.065 | 1 |
| BCH2 | 0.000 | -0.001 | 0.000 | 0 |
| YNL092W | -0.007 | 0.007 | 0.000 | 0 |
| TEC1 | 0.002 | -0.020 | -0.009 | 0 |
| YIL100W | 0.007 | 0.010 | 0.009 | 0 |
| HSM3 | 0.030 | 0.029 | 0.030 | 1 |
| ASI3 | 0.005 | -0.015 | -0.005 | 0 |
| QDR1 | -0.020 | -0.010 | -0.015 | 0 |
| SSN8 | 0.047 | 0.068 | 0.058 | 1 |
| TPM2 | 0.008 | -0.008 | 0.000 | 0 |
| YNL043C | -0.014 | -0.012 | -0.013 | 0 |
| UBP7 | -0.001 | -0.018 | -0.010 | 0 |
| ATG3 | -0.004 | 0.005 | 0.000 | 0 |
| MRPL50 | 0.010 | -0.005 | 0.003 | 0 |
| SNF3 | -0.017 | -0.004 | -0.011 | 0 |
| ZRG17 | 0.009 | 0.010 | 0.010 | 0 |
| TOM7 | 0.023 | 0.024 | 0.023 | 1 |
| YBL100C | -0.006 | 0.005 | 0.000 | 0 |
| TCB2 | 0.006 | 0.001 | 0.003 | 0 |
| DSF2 | -0.012 | 0.011 | -0.001 | 0 |
| YAF9 | -0.054 | -0.039 | -0.046 | -1 |
| FUR4 | -0.004 | -0.008 | -0.006 | 0 |
| HMO1 | 0.005 | 0.005 | 0.005 | 0 |
| HMT1 | -0.005 | 0.005 | 0.000 | 0 |
| YGR053C | -0.003 | -0.008 | -0.005 | 0 |
| REG2 | -0.007 | -0.003 | -0.005 | 0 |
| YGR250C | -0.007 | -0.003 | -0.005 | 0 |
| YBR063C | 0.001 | -0.004 | -0.001 | 0 |
| ECM8 | 0.008 | -0.006 | 0.001 | 0 |
| YKL115C | -0.011 | -0.003 | -0.007 | 0 |
| NRK1 | 0.004 | -0.002 | 0.001 | 0 |
| SAP190 | -0.021 | -0.009 | -0.015 | 0 |
| RPL19A | 0.020 | 0.009 | 0.014 | 0 |
| XBP1 | -0.013 | -0.010 | -0.011 | 0 |
| RIF1 | -0.002 | -0.007 | -0.005 | 0 |
| IDP3 | -0.006 | -0.008 | -0.007 | 0 |
| QDR2 | 0.000 | 0.044 | 0.022 | 0 |
| CRZ1 | 0.011 | 0.011 | 0.011 | 0 |
| REV7 | 0.018 | 0.002 | 0.010 | 0 |
| YIP3 | -0.004 | -0.017 | -0.010 | 0 |
| COA1 | -0.003 | -0.005 | -0.004 | 0 |
| LRO1 | -0.009 | -0.009 | -0.009 | 0 |

| | | | | |
|---|---|---|---|---|
| HXT12 | 0.003 | 0.000 | 0.002 | 0 |
| MPP6 | 0.011 | -0.006 | 0.003 | 0 |
| TRP1 | 0.003 | 0.011 | 0.007 | 0 |
| YNR040W | 0.007 | -0.025 | -0.009 | 0 |
| LAT1 | 0.006 | 0.033 | 0.019 | 0 |
| ECM21 | -0.058 | -0.004 | -0.031 | 0 |
| RHO2 | -0.004 | -0.009 | -0.006 | 0 |
| FLR1 | -0.008 | -0.010 | -0.009 | 0 |
| YNL115C | -0.008 | -0.007 | -0.007 | 0 |
| POA1 | -0.001 | 0.002 | 0.000 | 0 |
| RAV2 | -0.046 | -0.120 | -0.083 | -1 |
| CSG2 | 0.016 | 0.027 | 0.022 | 0 |
| SPT4 | -0.021 | -0.005 | -0.013 | 0 |
| YBR051W | -0.004 | -0.005 | -0.005 | 0 |
| YBR064W | -0.012 | 0.008 | -0.002 | 0 |
| SLM4 | -0.008 | -0.006 | -0.007 | 0 |
| CTK1 | -0.010 | 0.022 | 0.006 | 0 |
| TEP1 | 0.000 | 0.001 | 0.000 | 0 |
| SET3 | -0.001 | -0.007 | -0.004 | 0 |
| SWS2 | -0.009 | 0.021 | 0.006 | 0 |
| YBR090C | -0.001 | 0.004 | 0.002 | 0 |
| DPH1 | -0.003 | -0.014 | -0.008 | 0 |
| PPS1 | 0.003 | 0.015 | 0.009 | 0 |
| YNL010W | -0.061 | -0.157 | -0.109 | -1 |
| SIM1 | 0.008 | -0.034 | -0.013 | 0 |
| YNL028W | -0.001 | -0.002 | -0.001 | 0 |
| AXL2 | 0.029 | 0.020 | 0.024 | 1 |
| YNL045W | -0.010 | 0.002 | -0.004 | 0 |
| BNR1 | -0.012 | -0.010 | -0.011 | 0 |
| NRM1 | 0.006 | 0.017 | 0.012 | 0 |
| VTH1 | 0.004 | -0.005 | 0.000 | 0 |
| YNR025C | 0.013 | -0.003 | 0.005 | 0 |
| YDR048C | -0.003 | 0.000 | -0.001 | 0 |
| COQ2 | 0.019 | 0.036 | 0.027 | 0 |
| RNH201 | 0.008 | 0.009 | 0.008 | 0 |
| SFT2 | -0.002 | -0.004 | -0.003 | 0 |
| NST1 | 0.001 | 0.000 | 0.000 | 0 |
| HHF1 | 0.013 | 0.016 | 0.014 | 0 |
| NCS2 | 0.020 | 0.020 | 0.020 | 1 |
| CHS3 | 0.008 | 0.003 | 0.005 | 0 |
| MSC2 | -0.006 | -0.001 | -0.004 | 0 |
| SCO1 | 0.028 | 0.041 | 0.034 | 1 |
| PIL1 | 0.002 | -0.001 | 0.001 | 0 |
| RFS1 | -0.002 | 0.011 | 0.004 | 0 |
| ECM2 | -0.001 | 0.007 | 0.003 | 0 |
| YNL146W | 0.003 | 0.004 | 0.004 | 0 |
| MST1 | -0.008 | -0.015 | -0.012 | 0 |
| FAR11 | 0.008 | 0.020 | 0.014 | 0 |
| DAL80 | -0.023 | -0.028 | -0.026 | -1 |
| APJ1 | 0.010 | 0.017 | 0.014 | 0 |
| YBR100W | -0.014 | -0.006 | -0.010 | 0 |
| SLM1 | 0.005 | 0.002 | 0.003 | 0 |
| SAF1 | -0.001 | 0.015 | 0.007 | 0 |

| | | | | |
|---|---|---|---|---|
| SPO1 | -0.003 | -0.018 | -0.011 | 0 |
| AYR1 | -0.005 | -0.033 | -0.019 | 0 |
| KTR5 | -0.002 | 0.002 | 0.000 | 0 |
| YIL141W | 0.028 | 0.020 | 0.024 | 1 |
| YNL046W | -0.009 | -0.017 | -0.013 | 0 |
| POT1 | -0.012 | -0.013 | -0.012 | 0 |
| CSE2 | 0.037 | 0.034 | 0.036 | 1 |
| SGN1 | 0.008 | -0.002 | 0.003 | 0 |
| BUD17 | -0.029 | -0.023 | -0.026 | -1 |
| YFR011C | 0.030 | 0.012 | 0.021 | 0 |
| YNR042W | -0.018 | -0.017 | -0.018 | 0 |
| RTG3 | -0.011 | -0.002 | -0.007 | 0 |
| YPT53 | -0.005 | -0.016 | -0.011 | 0 |
| HHT1 | 0.019 | 0.026 | 0.023 | 0 |
| YNL120C | 0.004 | 0.018 | 0.011 | 0 |
| SCO2 | 0.006 | 0.000 | 0.003 | 0 |
| YDR445C | -0.006 | -0.007 | -0.007 | 0 |
| FIG1 | 0.004 | -0.003 | 0.001 | 0 |
| NNF2 | 0.042 | 0.038 | 0.040 | 1 |
| YBR053C | -0.006 | -0.005 | -0.005 | 0 |
| RPS21B | -0.024 | -0.009 | -0.016 | 0 |
| NRG2 | -0.001 | -0.001 | -0.001 | 0 |
| YJL175W | -0.010 | -0.017 | -0.013 | 0 |
| MFA2 | -0.007 | 0.005 | -0.001 | 0 |
| MNN4 | 0.002 | 0.001 | 0.001 | 0 |
| NMA111 | 0.007 | -0.008 | 0.000 | 0 |
| CAF4 | -0.021 | -0.004 | -0.013 | 0 |
| RPL16B | 0.017 | -0.006 | 0.006 | 0 |
| ASI1 | -0.001 | 0.012 | 0.005 | 0 |
| PFK26 | 0.000 | -0.002 | -0.001 | 0 |
| YBR287W | 0.000 | -0.003 | -0.001 | 0 |
| YNL013C | 0.003 | -0.008 | -0.003 | 0 |
| KGD1 | 0.007 | 0.007 | 0.007 | 0 |
| HHF2 | 0.002 | -0.017 | -0.008 | 0 |
| PAN6 | -0.001 | -0.020 | -0.011 | 0 |
| SFB2 | -0.007 | 0.020 | 0.007 | 0 |
| YIL161W | 0.003 | -0.003 | 0.000 | 0 |
| URK1 | 0.001 | 0.018 | 0.009 | 0 |
| MPH1 | 0.003 | -0.041 | -0.019 | 0 |
| CPR8 | -0.005 | 0.004 | 0.000 | 0 |
| IOC3 | 0.002 | 0.000 | 0.001 | 0 |
| PET494 | -0.013 | 0.012 | -0.001 | 0 |
| MLF3 | 0.002 | -0.002 | 0.000 | 0 |
| YBL104C | -0.002 | -0.003 | -0.002 | 0 |
| YNL095C | 0.004 | 0.003 | 0.003 | 0 |
| YBR012C | 0.001 | -0.015 | -0.007 | 0 |
| TOM70 | 0.021 | 0.020 | 0.021 | 0 |
| OLA1 | -0.003 | 0.000 | -0.001 | 0 |
| YDR537C | -0.024 | -0.014 | -0.019 | 0 |
| FAT1 | -0.025 | -0.040 | -0.032 | -1 |
| DBF2 | 0.011 | -0.018 | -0.004 | 0 |
| YRO2 | -0.017 | -0.009 | -0.013 | 0 |
| GLG2 | -0.011 | 0.006 | -0.002 | 0 |

| | | | | |
|---|---|---|---|---|
| TIP1 | -0.005 | 0.009 | 0.002 | 0 |
| RPL17B | 0.005 | -0.004 | 0.000 | 0 |
| YNL144C | 0.003 | -0.003 | 0.000 | 0 |
| YKL202W | 0.003 | 0.005 | 0.004 | 0 |
| YNL122C | 0.008 | 0.003 | 0.006 | 0 |
| GAP1 | -0.005 | -0.013 | -0.009 | 0 |
| YDJ1 | 0.048 | 0.050 | 0.049 | 1 |
| PTC4 | -0.006 | -0.019 | -0.013 | 0 |
| YIL108W | 0.004 | -0.018 | -0.007 | 0 |
| APM3 | -0.046 | -0.195 | -0.121 | -1 |
| PBI2 | -0.011 | -0.032 | -0.021 | 0 |
| MET18 | 0.020 | 0.019 | 0.019 | 0 |
| HHT2 | 0.018 | 0.015 | 0.017 | 0 |
| ECM37 | -0.006 | -0.033 | -0.020 | 0 |
| YNL050C | 0.004 | -0.011 | -0.003 | 0 |
| SUC2 | -0.012 | 0.007 | -0.002 | 0 |
| PHO91 | 0.012 | 0.003 | 0.007 | 0 |
| YIR003W | 0.001 | -0.045 | -0.022 | 0 |
| YNR029C | -0.018 | -0.018 | -0.018 | 0 |
| COG5 | -0.026 | -0.025 | -0.025 | -1 |
| FPK1 | -0.042 | -0.032 | -0.037 | -1 |
| MKS1 | -0.008 | -0.039 | -0.024 | 0 |
| SRO77 | -0.006 | -0.049 | -0.027 | 0 |
| PHO23 | 0.030 | 0.016 | 0.023 | 0 |
| YBR013C | -0.016 | 0.006 | -0.005 | 0 |
| ESBP6 | 0.010 | 0.018 | 0.014 | 0 |
| ETR1 | 0.000 | 0.022 | 0.011 | 0 |
| YFR039C | -0.014 | 0.001 | -0.006 | 0 |
| CST26 | 0.004 | 0.009 | 0.006 | 0 |
| YGR093W | -0.003 | 0.002 | -0.001 | 0 |
| YBR056W | 0.007 | -0.001 | 0.003 | 0 |
| BAP2 | 0.004 | 0.002 | 0.003 | 0 |
| YNL143C | -0.016 | -0.001 | -0.008 | 0 |
| EAP1 | 0.012 | 0.009 | 0.010 | 0 |
| MLS1 | 0.005 | 0.000 | 0.003 | 0 |
| YKR040C | -0.013 | 0.003 | -0.005 | 0 |
| MTQ1 | -0.008 | 0.005 | -0.002 | 0 |
| CCZ1 | 0.013 | 0.017 | 0.015 | 0 |
| MNI1 | -0.001 | 0.029 | 0.014 | 0 |
| SNF5 | 0.003 | -0.011 | -0.004 | 0 |
| SUL1 | -0.004 | -0.010 | -0.007 | 0 |
| TPS1 | -0.043 | -0.030 | -0.036 | -1 |
| YCR062W | 0.003 | -0.003 | 0.000 | 0 |
| YBR144C | -0.001 | 0.019 | 0.009 | 0 |
| TPS2 | -0.020 | -0.047 | -0.034 | 0 |
| TOS1 | 0.006 | 0.001 | 0.003 | 0 |
| NAM2 | 0.019 | -0.011 | 0.004 | 0 |
| YDL009C | -0.022 | 0.002 | -0.010 | 0 |
| WAR1 | -0.002 | -0.005 | -0.003 | 0 |
| DIA3 | 0.005 | -0.010 | -0.002 | 0 |
| RTT10 | -0.012 | -0.001 | -0.006 | 0 |
| YDL041W | -0.002 | -0.038 | -0.020 | 0 |
| YIR043C | -0.006 | -0.001 | -0.004 | 0 |

| | | | | |
|---|---|---|---|---|
| MBP1 | 0.027 | 0.007 | 0.017 | 0 |
| YDL073W | -0.018 | -0.007 | -0.012 | 0 |
| MOD5 | 0.002 | 0.005 | 0.004 | 0 |
| ASM4 | -0.018 | -0.005 | -0.012 | 0 |
| GIP3 | -0.011 | -0.012 | -0.012 | 0 |
| SSN2 | 0.011 | 0.037 | 0.024 | 0 |
| HIS2 | -0.009 | 0.007 | -0.001 | 0 |
| PFA5 | -0.014 | -0.010 | -0.012 | 0 |
| RPL40B | 0.006 | 0.028 | 0.017 | 0 |
| PEX29 | -0.026 | -0.038 | -0.032 | -1 |
| MSN2 | 0.002 | 0.010 | 0.006 | 0 |
| RSM28 | 0.003 | -0.004 | 0.000 | 0 |
| EAF3 | -0.009 | 0.013 | 0.002 | 0 |
| YDR509W | 0.015 | 0.022 | 0.018 | 0 |
| AAC3 | -0.006 | 0.006 | 0.000 | 0 |
| API2 | 0.037 | 0.018 | 0.028 | 0 |
| PHO88 | 0.127 | 0.149 | 0.138 | 1 |
| YGL109W | 0.000 | 0.013 | 0.007 | 0 |
| DAN3 | -0.005 | 0.010 | 0.002 | 0 |
| VMA2 | -0.048 | -0.071 | -0.060 | -1 |
| SED4 | 0.036 | 0.035 | 0.035 | 1 |
| ADH5 | -0.012 | 0.003 | -0.005 | 0 |
| GAL83 | -0.001 | -0.006 | -0.004 | 0 |
| YSY6 | 0.015 | 0.013 | 0.014 | 0 |
| CST9 | -0.002 | 0.016 | 0.007 | 0 |
| GRX6 | -0.009 | -0.001 | -0.005 | 0 |
| YDL025C | 0.019 | 0.024 | 0.022 | 0 |
| RTC6 | 0.005 | 0.023 | 0.014 | 0 |
| SIR2 | 0.016 | -0.001 | 0.007 | 0 |
| YIR044C | -0.006 | -0.004 | -0.005 | 0 |
| YDL057W | -0.001 | 0.004 | 0.001 | 0 |
| RIM20 | -0.033 | -0.049 | -0.041 | -1 |
| NUR1 | -0.007 | -0.001 | -0.004 | 0 |
| GPA1 | 0.005 | -0.001 | 0.002 | 0 |
| ECM11 | -0.003 | 0.000 | -0.001 | 0 |
| MET10 | 0.002 | 0.013 | 0.007 | 0 |
| MRPL28 | 0.004 | -0.014 | -0.005 | 0 |
| MLP1 | -0.011 | 0.001 | -0.005 | 0 |
| DIG2 | -0.007 | -0.014 | -0.011 | 0 |
| YPK2 | 0.006 | -0.009 | -0.002 | 0 |
| VPS3 | -0.019 | -0.019 | -0.019 | 0 |
| YME1 | 0.018 | 0.008 | 0.013 | 0 |
| ACN9 | -0.006 | 0.001 | -0.003 | 0 |
| YBR090C-A | 0.003 | -0.008 | -0.003 | 0 |
| HLR1 | -0.020 | -0.015 | -0.018 | 0 |
| YOL073C | 0.013 | 0.018 | 0.015 | 0 |
| FRM2 | -0.019 | -0.004 | -0.012 | 0 |
| ATG14 | -0.012 | -0.016 | -0.014 | 0 |
| CPR4 | 0.009 | 0.012 | 0.010 | 0 |
| MRPS9 | -0.014 | -0.010 | -0.012 | 0 |
| PHM8 | -0.004 | -0.011 | -0.007 | 0 |
| DEM1 | 0.003 | -0.015 | -0.006 | 0 |
| RPL31B | -0.002 | 0.008 | 0.003 | 0 |

| | | | | |
|---|---|---|---|---|
| YDL011C | -0.006 | 0.008 | 0.001 | 0 |
| CSM3 | 0.000 | -0.002 | -0.001 | 0 |
| YDL026W | -0.004 | -0.007 | -0.005 | 0 |
| MTF2 | 0.019 | -0.140 | -0.061 | 0 |
| YJR003C | -0.002 | -0.028 | -0.015 | 0 |
| RAD59 | 0.015 | 0.022 | 0.018 | 0 |
| RXT3 | 0.019 | 0.027 | 0.023 | 0 |
| CAF20 | -0.008 | -0.002 | -0.005 | 0 |
| RAM1 | 0.016 | -0.004 | 0.006 | 0 |
| RPS17B | -0.002 | -0.001 | -0.001 | 0 |
| CRP1 | 0.001 | -0.009 | -0.004 | 0 |
| STP1 | -0.043 | -0.059 | -0.051 | -1 |
| YKR096W | 0.017 | 0.022 | 0.019 | 0 |
| PHO8 | 0.000 | 0.012 | 0.006 | 0 |
| YNL109W | -0.002 | -0.012 | -0.007 | 0 |
| PUF6 | 0.003 | 0.026 | 0.015 | 0 |
| ATH1 | -0.005 | -0.005 | -0.005 | 0 |
| EMI1 | 0.011 | 0.008 | 0.010 | 0 |
| PHO3 | 0.017 | -0.014 | 0.001 | 0 |
| IML3 | -0.006 | 0.007 | 0.000 | 0 |
| CUE3 | -0.001 | 0.012 | 0.006 | 0 |
| RIM1 | -0.116 | 0.022 | -0.047 | 0 |
| OPY1 | -0.001 | 0.016 | 0.008 | 0 |
| SSK22 | 0.006 | -0.013 | -0.004 | 0 |
| RTC2 | -0.010 | -0.001 | -0.005 | 0 |
| CYS4 | 0.008 | -0.004 | 0.002 | 0 |
| ARL1 | -0.077 | -0.224 | -0.150 | -1 |
| APT1 | 0.017 | -0.002 | 0.007 | 0 |
| YDL012C | 0.004 | 0.009 | 0.007 | 0 |
| (MR135W-A | 0.000 | -0.006 | -0.003 | 0 |
| YDL027C | -0.004 | -0.008 | -0.006 | 0 |
| MMT2 | 0.003 | -0.004 | -0.001 | 0 |
| MRP10 | 0.000 | 0.027 | 0.013 | 0 |
| RPS29B | -0.008 | -0.001 | -0.005 | 0 |
| HPF1 | -0.004 | 0.008 | 0.002 | 0 |
| VAM6 | 0.015 | 0.020 | 0.017 | 0 |
| MBF1 | 0.008 | 0.007 | 0.008 | 0 |
| UBX3 | -0.002 | 0.007 | 0.003 | 0 |
| SYC1 | 0.002 | -0.020 | -0.009 | 0 |
| ADA2 | 0.093 | 0.115 | 0.104 | 1 |
| ATG7 | 0.004 | -0.007 | -0.001 | 0 |
| RMT2 | 0.000 | 0.005 | 0.003 | 0 |
| FLO10 | -0.002 | -0.002 | -0.002 | 0 |
| CWC21 | -0.047 | -0.046 | -0.047 | -1 |
| POP2 | -0.028 | -0.016 | -0.022 | 0 |
| ITR1 | -0.011 | 0.006 | -0.002 | 0 |
| NTO1 | 0.004 | 0.002 | 0.003 | 0 |
| GRX2 | -0.003 | -0.006 | -0.005 | 0 |
| PHO5 | 0.008 | -0.009 | 0.000 | 0 |
| APA2 | -0.005 | 0.006 | 0.000 | 0 |
| YBR108W | -0.008 | -0.025 | -0.017 | 0 |
| YGL114W | 0.005 | -0.006 | -0.001 | 0 |
| SYP1 | -0.010 | -0.011 | -0.010 | 0 |

| | | | | |
|---|---|---|---|---|
| SHE3 | -0.016 | 0.003 | -0.006 | 0 |
| ERS1 | -0.026 | -0.034 | -0.030 | -1 |
| YSW1 | -0.026 | 0.002 | -0.012 | 0 |
| HCR1 | 0.029 | 0.035 | 0.032 | 1 |
| UBS1 | 0.011 | 0.002 | 0.007 | 0 |
| YOX1 | 0.014 | 0.000 | 0.007 | 0 |
| SLX5 | 0.046 | 0.035 | 0.040 | 1 |
| PSO2 | -0.009 | 0.002 | -0.004 | 0 |
| YDL032W | 0.010 | 0.004 | 0.007 | 0 |
| COS6 | 0.002 | -0.020 | -0.009 | 0 |
| NPC2 | -0.009 | -0.003 | -0.006 | 0 |
| MDM35 | 0.029 | 0.019 | 0.024 | 0 |
| YDL062W | 0.000 | 0.019 | 0.009 | 0 |
| RBL2 | -0.004 | 0.014 | 0.005 | 0 |
| MDH3 | 0.007 | -0.007 | 0.000 | 0 |
| PMT5 | -0.009 | -0.009 | -0.009 | 0 |
| DCI1 | 0.000 | 0.009 | 0.005 | 0 |
| RPS18A | -0.003 | 0.018 | 0.008 | 0 |
| MHP1 | -0.006 | -0.003 | -0.005 | 0 |
| PKH3 | 0.013 | 0.018 | 0.015 | 0 |
| CCW12 | 0.009 | 0.008 | 0.008 | 0 |
| VPS52 | 0.016 | 0.009 | 0.012 | 0 |
| HOL1 | -0.006 | 0.002 | -0.002 | 0 |
| RPL37B | 0.005 | 0.005 | 0.005 | 0 |
| ERV2 | 0.003 | -0.009 | -0.003 | 0 |
| YDR514C | -0.009 | -0.003 | -0.006 | 0 |
| PBY1 | 0.008 | -0.009 | -0.001 | 0 |
| YDR532C | -0.003 | -0.008 | -0.006 | 0 |
| YSA1 | -0.002 | 0.021 | 0.010 | 0 |
| SNF4 | -0.004 | -0.027 | -0.015 | 0 |
| BPH1 | -0.013 | -0.002 | -0.007 | 0 |
| AGP2 | -0.014 | -0.007 | -0.011 | 0 |
| TRX3 | -0.006 | 0.008 | 0.001 | 0 |
| ARA1 | -0.019 | 0.014 | -0.003 | 0 |
| THI7 | -0.001 | 0.006 | 0.003 | 0 |
| TYR1 | 0.008 | 0.011 | 0.010 | 0 |
| CGI121 | 0.009 | 0.011 | 0.010 | 0 |
| ERP3 | 0.009 | -0.011 | -0.001 | 0 |
| CIN4 | -0.014 | -0.009 | -0.011 | 0 |
| SLM3 | 0.002 | -0.008 | -0.003 | 0 |
| YHR132W-A | 0.005 | -0.005 | 0.000 | 0 |
| STP4 | -0.010 | 0.003 | -0.004 | 0 |
| YKR106W | 0.008 | -0.013 | -0.003 | 0 |
| YDL063C | 0.008 | 0.011 | 0.009 | 0 |
| PNT1 | -0.001 | 0.014 | 0.007 | 0 |
| MRK1 | 0.003 | 0.003 | 0.003 | 0 |
| YDL094C | 0.002 | 0.001 | 0.001 | 0 |
| IWR1 | 0.049 | 0.044 | 0.047 | 1 |
| YHP1 | -0.013 | 0.001 | -0.006 | 0 |
| YJL070C | 0.014 | -0.005 | 0.004 | 0 |
| YDR467C | 0.005 | -0.009 | -0.002 | 0 |
| CCW14 | 0.007 | 0.010 | 0.008 | 0 |
| VPS72 | -0.067 | -0.059 | -0.063 | -1 |

| | | | | |
|---|---|---|---|---|
| BSC5 | -0.005 | -0.009 | -0.007 | 0 |
| PLM2 | -0.015 | -0.013 | -0.014 | 0 |
| RPL43A | 0.053 | 0.032 | 0.043 | 1 |
| EMI2 | -0.006 | 0.008 | 0.001 | 0 |
| RXT2 | 0.022 | 0.023 | 0.022 | 1 |
| HSP31 | -0.001 | 0.009 | 0.004 | 0 |
| YBR113W | 0.005 | -0.003 | 0.001 | 0 |
| YGL117W | -0.007 | -0.002 | -0.005 | 0 |
| SNT1 | -0.029 | -0.022 | -0.025 | -1 |
| HSL7 | 0.005 | -0.003 | 0.001 | 0 |
| TUP1 | 0.054 | 0.041 | 0.048 | 1 |
| APD1 | -0.013 | -0.006 | -0.010 | 0 |
| ERF2 | 0.001 | -0.013 | -0.006 | 0 |
| NPL4 | 0.010 | -0.022 | -0.006 | 0 |
| YMD8 | 0.001 | 0.010 | 0.005 | 0 |
| OSH2 | -0.002 | 0.002 | 0.000 | 0 |
| RIM11 | -0.005 | -0.011 | -0.008 | 0 |
| YDL034W | -0.017 | -0.015 | -0.016 | 0 |
| SSM4 | 0.021 | 0.031 | 0.026 | 1 |
| KNH1 | 0.004 | 0.012 | 0.008 | 0 |
| PEX19 | -0.010 | 0.006 | -0.002 | 0 |
| HRK1 | 0.001 | -0.003 | -0.001 | 0 |
| THI3 | -0.008 | 0.003 | -0.002 | 0 |
| LSP1 | -0.013 | -0.015 | -0.014 | 0 |
| PMT1 | 0.002 | 0.005 | 0.004 | 0 |
| YFL013W-A | 0.032 | 0.040 | 0.036 | 1 |
| PPN1 | -0.011 | -0.005 | -0.008 | 0 |
| PRY3 | 0.011 | 0.002 | 0.007 | 0 |
| SDC1 | -0.003 | -0.007 | -0.005 | 0 |
| MRPL4 | -0.007 | 0.012 | 0.002 | 0 |
| VPS60 | -0.021 | -0.020 | -0.021 | -1 |
| CYB5 | -0.014 | -0.008 | -0.011 | 0 |
| LPP1 | -0.002 | 0.008 | 0.003 | 0 |
| YPR050C | -0.043 | -0.082 | -0.062 | -1 |
| GRH1 | -0.003 | 0.011 | 0.004 | 0 |
| MMS4 | -0.001 | 0.001 | 0.000 | 0 |
| FIT1 | -0.004 | 0.019 | 0.008 | 0 |
| RAD16 | 0.010 | 0.001 | 0.005 | 0 |
| YGL118C | -0.006 | 0.007 | 0.001 | 0 |
| IMG1 | -0.015 | -0.032 | -0.024 | 0 |
| YBR134W | 0.019 | 0.014 | 0.017 | 0 |
| ABP1 | 0.000 | -0.005 | -0.003 | 0 |
| SLI15 | -0.012 | 0.001 | -0.006 | 0 |
| YLR334C | 0.008 | -0.009 | -0.001 | 0 |
| SEC66 | 0.031 | 0.011 | 0.021 | 0 |
| VPS71 | -0.066 | -0.061 | -0.063 | -1 |
| RPN4 | 0.072 | 0.091 | 0.082 | 1 |
| YMR160W | -0.006 | -0.022 | -0.014 | 0 |
| GPR1 | 0.004 | 0.012 | 0.008 | 0 |
| YIL058W | -0.006 | 0.004 | -0.001 | 0 |
| YDL050C | 0.010 | 0.002 | 0.006 | 0 |
| SNO4 | -0.005 | -0.010 | -0.008 | 0 |
| IDP1 | 0.016 | 0.031 | 0.023 | 0 |

| | | | | |
|---|---|---|---|---|
| YOR268C | 0.013 | 0.007 | 0.010 | 0 |
| RPP1A | 0.019 | -0.012 | 0.004 | 0 |
| IRC15 | -0.009 | -0.003 | -0.006 | 0 |
| THI74 | 0.001 | 0.002 | 0.001 | 0 |
| HSP12 | -0.031 | -0.010 | -0.021 | 0 |
| TSA2 | -0.013 | 0.004 | -0.004 | 0 |
| KHA1 | 0.007 | 0.005 | 0.006 | 0 |
| UGO1 | 0.035 | 0.007 | 0.021 | 0 |
| SIR3 | 0.005 | -0.020 | -0.007 | 0 |
| PAC11 | -0.007 | 0.012 | 0.003 | 0 |
| PEX11 | -0.006 | -0.009 | -0.008 | 0 |
| SPG3 | -0.005 | -0.004 | -0.004 | 0 |
| YPR064W | -0.009 | -0.012 | -0.011 | 0 |
| EUG1 | 0.002 | -0.002 | 0.000 | 0 |
| YBR099C | -0.010 | -0.010 | -0.010 | 0 |
| YGL101W | 0.010 | 0.021 | 0.015 | 0 |
| LYS2 | -0.007 | -0.042 | -0.024 | 0 |
| GPG1 | -0.007 | 0.003 | -0.002 | 0 |
| BUD23 | -0.007 | 0.007 | 0.000 | 0 |
| YBR137W | 0.011 | -0.004 | 0.004 | 0 |
| FIG2 | 0.002 | -0.009 | -0.004 | 0 |
| ICS2 | -0.015 | 0.001 | -0.007 | 0 |
| YLR346C | 0.006 | 0.004 | 0.005 | 0 |
| SMY2 | 0.007 | 0.002 | 0.005 | 0 |
| GPM2 | 0.020 | 0.030 | 0.025 | 0 |
| DDR48 | -0.005 | -0.001 | -0.003 | 0 |
| PUS9 | 0.015 | 0.011 | 0.013 | 0 |
| LHP1 | 0.010 | -0.003 | 0.003 | 0 |
| SRV2 | -0.005 | -0.037 | -0.021 | 0 |
| YDL068W | -0.011 | 0.023 | 0.006 | 0 |
| PAC1 | 0.000 | -0.002 | -0.001 | 0 |
| RPL13A | 0.007 | 0.008 | 0.007 | 0 |
| SMA1 | -0.008 | 0.008 | 0.000 | 0 |
| LRS4 | -0.003 | -0.015 | -0.009 | 0 |
| YFL019C | -0.006 | -0.001 | -0.003 | 0 |
| YDR455C | -0.009 | 0.009 | 0.000 | 0 |
| GSH1 | 0.010 | -0.017 | -0.004 | 0 |
| RPL27B | 0.026 | 0.025 | 0.025 | 1 |
| PKH1 | -0.001 | 0.009 | 0.004 | 0 |
| REC8 | -0.009 | 0.016 | 0.004 | 0 |
| PSP1 | -0.009 | -0.015 | -0.012 | 0 |
| YPR078C | -0.003 | -0.002 | -0.003 | 0 |
| FPR2 | 0.008 | 0.000 | 0.004 | 0 |
| VPS73 | 0.001 | 0.002 | 0.001 | 0 |
| YBR116C | 0.005 | 0.002 | 0.003 | 0 |
| MON1 | 0.019 | 0.020 | 0.020 | 0 |
| ARE1 | -0.009 | -0.029 | -0.019 | 0 |
| YBR138C | 0.009 | -0.013 | -0.002 | 0 |
| RPL21A | 0.054 | 0.058 | 0.056 | 1 |
| AMN1 | -0.016 | -0.008 | -0.012 | 0 |
| YLR358C | 0.036 | 0.027 | 0.031 | 1 |
| RMD1 | -0.006 | -0.012 | -0.009 | 0 |
| CAT2 | -0.002 | 0.002 | 0.000 | 0 |

| | | | | |
|---|---|---|---|---|
| GPD1 | -0.006 | -0.007 | -0.007 | 0 |
| BSC1 | 0.012 | 0.000 | 0.006 | 0 |
| DAL81 | -0.046 | -0.064 | -0.055 | -1 |
| SLC1 | 0.012 | -0.007 | 0.002 | 0 |
| YNL140C | -0.006 | 0.002 | -0.002 | 0 |
| CBS1 | 0.032 | 0.036 | 0.034 | 1 |
| VPH1 | -0.013 | -0.105 | -0.059 | 0 |
| RPS16B | -0.023 | -0.020 | -0.022 | -1 |
| YPL034W | -0.004 | -0.017 | -0.010 | 0 |
| DOT1 | 0.017 | 0.010 | 0.013 | 0 |
| YFL042C | -0.002 | 0.006 | 0.002 | 0 |
| NHX1 | -0.010 | -0.007 | -0.008 | 0 |
| SET4 | -0.004 | 0.000 | -0.002 | 0 |
| YDR474C | -0.010 | 0.005 | -0.003 | 0 |
| YDR491C | -0.014 | -0.020 | -0.017 | 0 |
| HAA1 | -0.004 | 0.008 | 0.002 | 0 |
| YDR506C | 0.002 | -0.004 | -0.001 | 0 |
| SPT7 | -0.004 | -0.017 | -0.011 | 0 |
| URC2 | -0.002 | -0.016 | -0.009 | 0 |
| SIF2 | 0.000 | 0.009 | 0.004 | 0 |
| ARC1 | 0.016 | 0.022 | 0.019 | 0 |
| MUD1 | 0.010 | 0.004 | 0.007 | 0 |
| MET13 | -0.021 | -0.017 | -0.019 | 0 |
| THR4 | 0.003 | -0.012 | -0.005 | 0 |
| YBR139W | 0.006 | 0.001 | 0.003 | 0 |
| GRX1 | -0.002 | 0.004 | 0.001 | 0 |
| YBR159W | -0.009 | 0.001 | -0.004 | 0 |
| DCR2 | -0.016 | -0.024 | -0.020 | 0 |
| NHP10 | 0.011 | 0.019 | 0.015 | 0 |
| PRM6 | -0.006 | 0.002 | -0.002 | 0 |
| YDL038C | 0.003 | -0.018 | -0.007 | 0 |
| DCG1 | 0.000 | -0.005 | -0.002 | 0 |
| PBP4 | 0.000 | -0.012 | -0.006 | 0 |
| MEP2 | -0.014 | -0.002 | -0.008 | 0 |
| BDF2 | 0.015 | 0.026 | 0.020 | 0 |
| FSF1 | -0.014 | -0.012 | -0.013 | 0 |
| NDE2 | 0.010 | 0.012 | 0.011 | 0 |
| PMA2 | -0.014 | -0.007 | -0.011 | 0 |
| APT2 | 0.015 | 0.009 | 0.012 | 0 |
| FAB1 | -0.009 | -0.010 | -0.009 | 0 |
| TOM1 | 0.002 | 0.005 | 0.004 | 0 |
| PBS2 | -0.036 | -0.030 | -0.033 | -1 |
| JIP4 | -0.007 | 0.001 | -0.003 | 0 |
| SMA2 | -0.005 | 0.005 | 0.000 | 0 |
| IZH1 | -0.005 | -0.003 | -0.004 | 0 |
| YPR013C | 0.012 | 0.012 | 0.012 | 0 |
| GIN4 | 0.009 | -0.014 | -0.002 | 0 |
| UBC4 | 0.050 | 0.046 | 0.048 | 1 |
| SPS2 | -0.011 | -0.007 | -0.009 | 0 |
| YMC2 | 0.021 | 0.011 | 0.016 | 0 |
| RMD9 | -0.011 | 0.011 | 0.000 | 0 |
| CBP6 | -0.012 | 0.003 | -0.004 | 0 |
| SCS3 | 0.014 | 0.016 | 0.015 | 0 |

| | | | | |
|---|---|---|---|---|
| TAH1 | -0.005 | 0.001 | -0.002 | 0 |
| YBR141C | 0.008 | -0.006 | 0.001 | 0 |
| PAA1 | -0.016 | -0.029 | -0.022 | 0 |
| CSH1 | -0.001 | 0.006 | 0.003 | 0 |
| ARC18 | 0.038 | 0.014 | 0.026 | 0 |
| PTC1 | 0.039 | 0.047 | 0.043 | 1 |
| HMG1 | 0.001 | 0.014 | 0.008 | 0 |
| YDL023C | 0.005 | 0.006 | 0.006 | 0 |
| YOR364W | -0.020 | -0.004 | -0.012 | 0 |
| PRM7 | -0.003 | -0.013 | -0.008 | 0 |
| DAL3 | -0.015 | 0.001 | -0.007 | 0 |
| MCH1 | -0.010 | 0.003 | -0.004 | 0 |
| ATP11 | 0.005 | 0.009 | 0.007 | 0 |
| YDL071C | 0.011 | 0.016 | 0.014 | 0 |
| TPO4 | -0.005 | 0.012 | 0.003 | 0 |
| YDL086W | 0.005 | -0.004 | 0.000 | 0 |
| YDR442W | 0.039 | 0.028 | 0.034 | 1 |
| YFR024C | -0.003 | -0.011 | -0.007 | 0 |
| HEH2 | -0.004 | -0.016 | -0.010 | 0 |
| YLR455W | 0.002 | 0.008 | 0.005 | 0 |
| YDR476C | -0.012 | -0.009 | -0.011 | 0 |
| VAN1 | -0.003 | -0.013 | -0.008 | 0 |
| YPR022C | 0.003 | -0.004 | -0.001 | 0 |
| GNP1 | 0.001 | 0.017 | 0.009 | 0 |
| MIS1 | 0.006 | 0.012 | 0.009 | 0 |
| AGE1 | 0.001 | 0.017 | 0.009 | 0 |
| VID24 | 0.006 | -0.002 | 0.002 | 0 |
| YGL108C | 0.000 | -0.005 | -0.002 | 0 |
| GRS1 | -0.015 | -0.006 | -0.011 | 0 |
| SOH1 | 0.023 | 0.006 | 0.015 | 0 |
| RSM23 | 0.002 | 0.000 | 0.001 | 0 |
| DHR2 | -0.002 | 0.010 | 0.004 | 0 |
| YGL146C | -0.013 | -0.011 | -0.012 | 0 |
| MTR2 | 0.005 | 0.004 | 0.004 | 0 |
| YGL160W | -0.001 | 0.022 | 0.010 | 0 |
| LAS1 | 0.024 | 0.009 | 0.016 | 0 |
| SAE2 | 0.000 | -0.003 | -0.002 | 0 |
| SED5 | -0.017 | 0.013 | -0.002 | 0 |
| YFL063W | -0.004 | -0.017 | -0.011 | 0 |
| RRN5 | -0.001 | 0.007 | 0.003 | 0 |
| YCK3 | -0.072 | 0.010 | -0.031 | 0 |
| YEF3 | 0.005 | 0.000 | 0.002 | 0 |
| COX15 | 0.009 | -0.015 | -0.003 | 0 |
| SEC61 | 0.443 | 0.283 | 0.363 | 1 |
| YER158C | -0.014 | 0.001 | -0.006 | 0 |
| noname | 0.001 | -0.002 | 0.000 | 0 |
| TMT1 | -0.017 | 0.001 | -0.008 | 0 |
| MRPL23 | 0.034 | 0.000 | 0.017 | 0 |
| YER187W | -0.015 | 0.002 | -0.007 | 0 |
| SFP1 | 0.014 | -0.003 | 0.005 | 0 |
| RIB1 | 0.008 | -0.007 | 0.001 | 0 |
| SIT4 | -0.058 | -0.009 | -0.033 | 0 |
| MHR1 | 0.011 | -0.012 | 0.000 | 0 |

| | | | | |
|---|---|---|---|---|
| UMP1 | 0.134 | 0.129 | 0.131 | 1 |
| MTG1 | 0.003 | -0.017 | -0.007 | 0 |
| MAL32 | -0.014 | -0.003 | -0.008 | 0 |
| SYS1 | -0.236 | -0.020 | -0.128 | 0 |
| DEG1 | 0.050 | 0.039 | 0.045 | 1 |
| YJR038C | -0.019 | -0.009 | -0.014 | 0 |
| RAD2 | -0.007 | 0.007 | 0.000 | 0 |
| ISA2 | -0.005 | -0.022 | -0.013 | 0 |
| SNT2 | -0.009 | -0.009 | -0.009 | 0 |
| RRP14 | 0.001 | -0.001 | 0.000 | 0 |
| RPL9A | 0.012 | 0.019 | 0.015 | 0 |
| ACP1 | -0.020 | -0.014 | -0.017 | 0 |
| YIP5 | -0.003 | 0.002 | -0.001 | 0 |
| BET3 | -0.018 | -0.013 | -0.016 | 0 |
| YGL176C | -0.017 | -0.009 | -0.013 | 0 |
| RSC58 | 0.007 | 0.023 | 0.015 | 0 |
| SWI4 | 0.051 | 0.018 | 0.034 | 0 |
| RMP1 | 0.002 | 0.005 | 0.003 | 0 |
| DSE1 | 0.005 | 0.011 | 0.008 | 0 |
| MCM5 | 0.003 | 0.014 | 0.008 | 0 |
| MAG1 | 0.003 | -0.003 | 0.000 | 0 |
| AFG2 | 0.017 | 0.016 | 0.017 | 0 |
| YNL011C | -0.010 | 0.004 | -0.003 | 0 |
| ORC1 | 0.002 | 0.014 | 0.008 | 0 |
| ECM32 | -0.021 | -0.005 | -0.013 | 0 |
| PET123 | 0.023 | 0.000 | 0.011 | 0 |
| YMR052C-A | -0.004 | -0.006 | -0.005 | 0 |
| VPS33 | -0.005 | -0.004 | -0.004 | 0 |
| TOR1 | -0.008 | 0.003 | -0.002 | 0 |
| GET1 | -0.006 | 0.009 | 0.001 | 0 |
| COX9 | 0.007 | -0.009 | -0.001 | 0 |
| PRO1 | 0.003 | -0.011 | -0.004 | 0 |
| GSM1 | -0.012 | 0.007 | -0.003 | 0 |
| ATP25 | 0.001 | -0.016 | -0.008 | 0 |
| AAD3 | -0.003 | 0.008 | 0.002 | 0 |
| CTK2 | -0.001 | -0.020 | -0.010 | 0 |
| MSH4 | -0.019 | -0.006 | -0.013 | 0 |
| DHH1 | -0.030 | -0.022 | -0.026 | -1 |
| HEF3 | -0.011 | -0.004 | -0.007 | 0 |
| YGL132W | -0.010 | -0.012 | -0.011 | 0 |
| YKL088W | 0.006 | 0.007 | 0.006 | 0 |
| ARO2 | -0.029 | -0.016 | -0.022 | 0 |
| SDS22 | -0.003 | 0.011 | 0.004 | 0 |
| SUT1 | -0.016 | -0.001 | -0.008 | 0 |
| DRE2 | 0.008 | 0.008 | 0.008 | 0 |
| YGL177W | -0.025 | 0.004 | -0.011 | 0 |
| SPC3 | 0.020 | -0.011 | 0.004 | 0 |
| TMN3 | -0.006 | 0.004 | -0.001 | 0 |
| SEC10 | 0.020 | 0.020 | 0.020 | 0 |
| YER128W | -0.018 | -0.001 | -0.009 | 0 |
| GCD7 | 0.017 | 0.026 | 0.021 | 0 |
| DDI1 | -0.001 | 0.004 | 0.001 | 0 |
| UTP21 | 0.017 | 0.019 | 0.018 | 0 |

| | | | | |
|---|---|---|---|---|
| SPT2 | 0.025 | 0.020 | 0.022 | 0 |
| POB3 | 0.028 | 0.030 | 0.029 | 1 |
| BMH1 | 0.008 | 0.014 | 0.011 | 0 |
| VRP1 | 0.023 | 0.007 | 0.015 | 0 |
| FAR3 | 0.003 | 0.000 | 0.002 | 0 |
| DOC1 | 0.012 | -0.011 | 0.001 | 0 |
| YIL102C | -0.012 | 0.003 | -0.005 | 0 |
| OCH1 | 0.027 | 0.029 | 0.028 | 1 |
| BRE1 | -0.077 | -0.039 | -0.058 | -1 |
| RPL2B | 0.012 | 0.011 | 0.012 | 0 |
| HEM14 | 0.004 | -0.013 | -0.004 | 0 |
| RIB4 | 0.003 | -0.015 | -0.006 | 0 |
| AMD2 | -0.001 | 0.006 | 0.003 | 0 |
| VTC4 | -0.004 | -0.001 | -0.002 | 0 |
| VTC2 | -0.015 | -0.013 | -0.014 | 0 |
| NRP1 | 0.029 | 0.029 | 0.029 | 1 |
| SLH1 | -0.011 | 0.006 | -0.002 | 0 |
| VPS15 | -0.002 | -0.022 | -0.012 | 0 |
| ITC1 | -0.001 | -0.001 | -0.001 | 0 |
| YJU2 | 0.010 | 0.009 | 0.010 | 0 |
| YGL149W | -0.030 | -0.027 | -0.028 | -1 |
| MIA40 | 0.008 | 0.007 | 0.007 | 0 |
| RAD54 | 0.025 | 0.017 | 0.021 | 0 |
| RPF2 | -0.007 | -0.002 | -0.005 | 0 |
| TOS3 | -0.014 | 0.004 | -0.005 | 0 |
| RGR1 | 0.016 | 0.021 | 0.019 | 0 |
| BOI2 | 0.001 | -0.008 | -0.003 | 0 |
| RPS31 | 0.000 | -0.006 | -0.003 | 0 |
| SAK1 | -0.041 | 0.002 | -0.019 | 0 |
| STT4 | 0.008 | 0.012 | 0.010 | 0 |
| FTR1 | -0.001 | 0.005 | 0.002 | 0 |
| SPP382 | 0.021 | 0.018 | 0.020 | 0 |
| RAD4 | -0.011 | 0.001 | -0.005 | 0 |
| BET5 | -0.012 | -0.003 | -0.007 | 0 |
| PDA1 | -0.013 | -0.012 | -0.012 | 0 |
| OPI9 | -0.011 | -0.014 | -0.013 | 0 |
| STB2 | -0.028 | -0.023 | -0.025 | -1 |
| GRX5 | -0.003 | -0.013 | -0.008 | 0 |
| COX5B | -0.009 | 0.001 | -0.004 | 0 |
| RPB9 | 0.015 | 0.006 | 0.011 | 0 |
| RPL31A | 0.014 | 0.044 | 0.029 | 0 |
| RNR3 | -0.009 | -0.013 | -0.011 | 0 |
| BIM1 | 0.017 | 0.013 | 0.015 | 0 |
| TPD3 | 0.094 | -0.016 | 0.039 | 0 |
| YSP2 | -0.001 | 0.001 | 0.000 | 0 |
| MAD3 | -0.012 | -0.012 | -0.012 | 0 |
| BLM10 | -0.018 | -0.025 | -0.022 | 0 |
| TFP1 | 0.006 | -0.020 | -0.007 | 0 |
| ABZ1 | -0.005 | -0.001 | -0.003 | 0 |
| FES1 | -0.017 | -0.021 | -0.019 | 0 |
| RPL1B | 0.024 | 0.003 | 0.013 | 0 |
| SLD2 | 0.001 | 0.005 | 0.003 | 0 |
| NUT1 | 0.024 | 0.032 | 0.028 | 1 |

| | | | | |
|---|---|---|---|---|
| TOR2 | 0.012 | 0.016 | 0.014 | 0 |
| YRB30 | -0.008 | 0.002 | -0.003 | 0 |
| ORC3 | 0.012 | 0.006 | 0.009 | 0 |
| ATG1 | -0.013 | -0.004 | -0.008 | 0 |
| BOS1 | -0.010 | -0.018 | -0.014 | 0 |
| SPR6 | -0.003 | -0.002 | -0.002 | 0 |
| NMT1 | -0.020 | -0.005 | -0.012 | 0 |
| YER130C | 0.001 | 0.008 | 0.004 | 0 |
| CDC25 | 0.012 | 0.012 | 0.012 | 0 |
| PEA2 | -0.045 | -0.030 | -0.038 | -1 |
| SEC39 | 0.010 | 0.016 | 0.013 | 0 |
| YER163C | -0.005 | -0.002 | -0.004 | 0 |
| PGA3 | 0.017 | 0.013 | 0.015 | 0 |
| DMC1 | -0.003 | -0.008 | -0.005 | 0 |
| SSQ1 | -0.009 | -0.023 | -0.016 | 0 |
| STV1 | -0.071 | -0.066 | -0.069 | -1 |
| MNN9 | -0.005 | -0.001 | -0.003 | 0 |
| POG1 | -0.021 | 0.000 | -0.010 | 0 |
| VPS45 | 0.015 | -0.010 | 0.002 | 0 |
| SNF1 | -0.006 | -0.004 | -0.005 | 0 |
| LPD1 | -0.012 | 0.006 | -0.003 | 0 |
| MIG3 | -0.003 | 0.002 | -0.001 | 0 |
| YDR417C | 0.001 | 0.009 | 0.005 | 0 |
| YJL027C | -0.005 | -0.001 | -0.003 | 0 |
| WWM1 | -0.002 | -0.001 | -0.001 | 0 |
| YOR331C | 0.002 | -0.021 | -0.010 | 0 |
| YGR273C | -0.003 | -0.005 | -0.004 | 0 |
| MRPL36 | 0.000 | -0.022 | -0.011 | 0 |
| MRM2 | 0.030 | 0.012 | 0.021 | 0 |
| ABF1 | 0.011 | 0.008 | 0.010 | 0 |
| YGL152C | -0.001 | 0.000 | 0.000 | 0 |
| UBA1 | 0.004 | 0.001 | 0.003 | 0 |
| YGL165C | -0.013 | 0.002 | -0.006 | 0 |
| GPI13 | 0.022 | 0.012 | 0.017 | 0 |
| GTS1 | -0.018 | 0.001 | -0.009 | 0 |
| SMC4 | 0.014 | 0.021 | 0.017 | 0 |
| SLX8 | 0.052 | 0.034 | 0.043 | 1 |
| PWP1 | 0.011 | 0.010 | 0.011 | 0 |
| RPS26B | -0.014 | 0.000 | -0.007 | 0 |
| CDC3 | 0.003 | 0.007 | 0.005 | 0 |
| SPI1 | -0.023 | -0.006 | -0.015 | 0 |
| NBP1 | 0.002 | 0.013 | 0.007 | 0 |
| CHD1 | -0.019 | -0.031 | -0.025 | 0 |
| ERG13 | 0.019 | 0.018 | 0.019 | 0 |
| ISC10 | -0.010 | -0.003 | -0.007 | 0 |
| STE23 | -0.008 | -0.004 | -0.006 | 0 |
| BUB2 | -0.006 | -0.011 | -0.008 | 0 |
| VPS16 | 0.085 | -0.006 | 0.040 | 0 |
| FKH1 | -0.012 | -0.009 | -0.010 | 0 |
| YGL088W | -0.007 | 0.006 | -0.001 | 0 |
| KRE2 | 0.001 | 0.010 | 0.005 | 0 |
| BUD32 | 0.008 | -0.015 | -0.004 | 0 |
| ERG28 | -0.007 | -0.002 | -0.005 | 0 |

| | | | | |
|---|---|---|---|---|
| GRR1 | 0.036 | 0.000 | 0.018 | 0 |
| YDR444W | -0.004 | 0.005 | 0.001 | 0 |
| YJL028W | -0.013 | 0.000 | -0.006 | 0 |
| AUA1 | -0.009 | 0.008 | -0.001 | 0 |
| PET8 | -0.003 | 0.002 | 0.000 | 0 |
| RNH70 | -0.008 | -0.012 | -0.010 | 0 |
| NAT1 | -0.001 | 0.002 | 0.001 | 0 |
| YGL138C | -0.020 | -0.010 | -0.015 | 0 |
| SRP21 | 0.023 | 0.000 | 0.011 | 0 |
| PEX14 | -0.007 | -0.005 | -0.006 | 0 |
| PAP1 | 0.007 | 0.020 | 0.013 | 0 |
| CUP2 | -0.018 | -0.003 | -0.011 | 0 |
| COF1 | 0.011 | -0.012 | 0.000 | 0 |
| AST2 | -0.002 | 0.002 | 0.000 | 0 |
| GAA1 | 0.080 | -0.015 | 0.033 | 0 |
| RPL23B | -0.010 | -0.005 | -0.007 | 0 |
| NOP56 | -0.011 | 0.002 | -0.004 | 0 |
| PMD1 | -0.009 | -0.002 | -0.006 | 0 |
| TAD3 | 0.000 | 0.006 | 0.003 | 0 |
| UBP3 | 0.051 | 0.015 | 0.033 | 0 |
| SPT5 | -0.003 | 0.009 | 0.003 | 0 |
| DNF1 | -0.007 | -0.023 | -0.015 | 0 |
| SEC59 | 0.011 | 0.015 | 0.013 | 0 |
| YER181C | -0.012 | -0.001 | -0.006 | 0 |
| YLR391W | -0.001 | -0.004 | -0.003 | 0 |
| AAC1 | -0.008 | -0.004 | -0.006 | 0 |
| PHO85 | -0.013 | 0.053 | 0.020 | 0 |
| OM45 | -0.020 | 0.003 | -0.008 | 0 |
| GIM3 | 0.042 | 0.046 | 0.044 | 1 |
| YDR521W | 0.014 | 0.019 | 0.017 | 0 |
| AEP1 | 0.013 | 0.011 | 0.012 | 0 |
| HIS1 | -0.023 | -0.018 | -0.021 | 0 |
| CTS1 | 0.015 | 0.014 | 0.015 | 0 |
| MFA1 | 0.001 | -0.021 | -0.010 | 0 |
| SAG1 | -0.004 | -0.006 | -0.005 | 0 |
| YFL012W | -0.008 | -0.009 | -0.009 | 0 |
| ECM33 | 0.014 | 0.010 | 0.012 | 0 |
| MAL11 | -0.004 | -0.006 | -0.005 | 0 |
| YET3 | -0.004 | -0.002 | -0.003 | 0 |
| FLC3 | -0.020 | -0.007 | -0.014 | 0 |
| SDH3 | -0.001 | 0.003 | 0.001 | 0 |
| LYS5 | -0.079 | -0.120 | -0.100 | -1 |
| ECM9 | 0.011 | 0.023 | 0.017 | 0 |
| PMR1 | -0.005 | -0.011 | -0.008 | 0 |
| SSL1 | 0.001 | 0.008 | 0.004 | 0 |
| SSA4 | -0.001 | 0.004 | 0.001 | 0 |
| SEN2 | 0.001 | 0.011 | 0.006 | 0 |
| SHO1 | -0.019 | -0.008 | -0.013 | 0 |
| TUB4 | 0.014 | 0.019 | 0.016 | 0 |
| YER134C | -0.029 | -0.010 | -0.020 | 0 |
| SFH1 | 0.000 | 0.006 | 0.003 | 0 |
| YER152C | -0.013 | -0.003 | -0.008 | 0 |
| NSE5 | 0.010 | 0.011 | 0.011 | 0 |

| | | | | |
|---|---|---|---|---|
| BCK2 | 0.004 | 0.004 | 0.004 | 0 |
| TAP42 | 0.015 | 0.011 | 0.013 | 0 |
| FMP10 | -0.012 | -0.015 | -0.014 | 0 |
| VPS34 | 0.024 | 0.000 | 0.012 | 0 |
| YMR057C | -0.003 | 0.000 | -0.001 | 0 |
| RMI1 | 0.014 | 0.010 | 0.012 | 0 |
| YIL158W | 0.000 | -0.001 | -0.001 | 0 |
| CNM67 | -0.001 | -0.059 | -0.030 | 0 |
| SPS1 | 0.000 | 0.000 | 0.000 | 0 |
| SOV1 | -0.039 | -0.013 | -0.026 | 0 |
| DAP2 | -0.013 | -0.003 | -0.008 | 0 |
| MRPL11 | 0.009 | -0.018 | -0.004 | 0 |
| YDR493W | -0.001 | 0.005 | 0.002 | 0 |
| CPR7 | 0.011 | 0.012 | 0.011 | 0 |
| IES1 | 0.021 | 0.025 | 0.023 | 1 |
| END3 | 0.000 | 0.006 | 0.003 | 0 |
| YGR291C | 0.000 | 0.003 | 0.001 | 0 |
| GLO3 | 0.026 | 0.024 | 0.025 | 1 |
| YGL140C | -0.012 | -0.008 | -0.010 | 0 |
| RPC25 | 0.030 | 0.044 | 0.037 | 1 |
| AMS1 | -0.013 | -0.009 | -0.011 | 0 |
| NTR2 | 0.014 | 0.013 | 0.014 | 0 |
| HUR1 | 0.007 | 0.006 | 0.007 | 0 |
| NSE1 | 0.009 | 0.015 | 0.012 | 0 |
| MAM1 | 0.000 | 0.003 | 0.002 | 0 |
| CFT2 | 0.006 | 0.012 | 0.009 | 0 |
| AVT6 | -0.013 | -0.002 | -0.007 | 0 |
| UTP13 | 0.017 | 0.026 | 0.021 | 0 |
| YER135C | -0.010 | -0.003 | -0.006 | 0 |
| CWC24 | 0.010 | 0.012 | 0.011 | 0 |
| PET122 | 0.005 | -0.023 | -0.009 | 0 |
| YML6 | 0.018 | 0.013 | 0.015 | 0 |
| RPH1 | 0.002 | 0.003 | 0.002 | 0 |
| NUP116 | -0.023 | -0.035 | -0.029 | -1 |
| FAU1 | -0.033 | -0.016 | -0.025 | 0 |
| MAP1 | 0.001 | -0.001 | 0.000 | 0 |
| FET3 | -0.008 | 0.005 | -0.002 | 0 |
| YGR219W | -0.033 | -0.015 | -0.024 | 0 |
| DJP1 | 0.013 | 0.013 | 0.013 | 0 |
| MRPL13 | 0.008 | -0.010 | -0.001 | 0 |
| SSK1 | -0.024 | -0.026 | -0.025 | -1 |
| COQ5 | 0.006 | -0.011 | -0.003 | 0 |
| YHR033W | -0.007 | -0.002 | -0.004 | 0 |
| PAF1 | 0.002 | -0.016 | -0.007 | 0 |
| SAM2 | -0.019 | 0.012 | -0.004 | 0 |
| YJR037W | -0.008 | -0.002 | -0.005 | 0 |
| RIM15 | -0.021 | -0.018 | -0.019 | 0 |
| FMC1 | 0.013 | 0.014 | 0.013 | 0 |
| MAL12 | 0.001 | -0.004 | -0.001 | 0 |
| APQ13 | 0.000 | -0.007 | -0.003 | 0 |
| HUL5 | -0.008 | -0.005 | -0.006 | 0 |
| RPT1 | 0.059 | 0.069 | 0.064 | 1 |
| YGL157W | -0.005 | -0.012 | -0.009 | 0 |

| | | | | |
|---|---|---|---|---|
| RPC37 | 0.014 | 0.010 | 0.012 | 0 |
| SPO74 | -0.003 | -0.016 | -0.010 | 0 |
| RLP24 | -0.003 | 0.021 | 0.009 | 0 |
| YER108C | -0.009 | 0.000 | -0.005 | 0 |
| CLF1 | 0.000 | 0.014 | 0.007 | 0 |
| YER119C-A | -0.022 | -0.017 | -0.019 | 0 |
| IFH1 | 0.006 | 0.016 | 0.011 | 0 |
| YER137C | 0.001 | 0.002 | 0.001 | 0 |
| KAP95 | 0.023 | 0.016 | 0.020 | 0 |
| OXA1 | -0.003 | -0.020 | -0.012 | 0 |
| NDC1 | 0.064 | 0.031 | 0.048 | 1 |
| ADK2 | -0.006 | -0.001 | -0.004 | 0 |
| ERB1 | 0.018 | 0.023 | 0.020 | 0 |
| YER184C | -0.018 | 0.001 | -0.009 | 0 |
| CHC1 | -0.048 | -0.046 | -0.047 | -1 |
| PET191 | -0.014 | -0.018 | -0.016 | 0 |
| PET54 | 0.024 | -0.006 | 0.009 | 0 |
| VHR1 | -0.015 | 0.005 | -0.005 | 0 |
| MNN10 | 0.042 | 0.033 | 0.037 | 1 |
| YML010W-A | 0.009 | 0.025 | 0.017 | 0 |
| BUL2 | -0.060 | -0.061 | -0.060 | -1 |
| PER1 | 0.007 | -0.003 | 0.002 | 0 |
| RPL6A | -0.008 | 0.002 | -0.003 | 0 |
| YGR122C-A | -0.006 | 0.003 | -0.002 | 0 |
| GON7 | -0.003 | -0.020 | -0.012 | 0 |
| PDX3 | -0.070 | -0.021 | -0.046 | -1 |
| MRF1 | 0.017 | 0.001 | 0.009 | 0 |
| SRP102 | 0.000 | 0.006 | 0.003 | 0 |
| RCK1 | -0.003 | 0.000 | -0.001 | 0 |
| KAE1 | 0.024 | 0.023 | 0.023 | 1 |
| KEM1 | 0.012 | 0.023 | 0.017 | 0 |
| TEN1 | 0.043 | 0.036 | 0.040 | 1 |
| FLO8 | -0.017 | 0.002 | -0.007 | 0 |
| APC2 | 0.000 | -0.008 | -0.004 | 0 |
| SCS2 | -0.017 | -0.022 | -0.019 | 0 |
| CDC42 | 0.054 | 0.039 | 0.046 | 1 |
| RTR1 | -0.013 | -0.004 | -0.009 | 0 |
| ILV5 | 0.017 | 0.010 | 0.013 | 0 |
| BEM2 | -0.005 | -0.018 | -0.012 | 0 |
| RRN11 | 0.015 | 0.006 | 0.011 | 0 |
| RAD24 | 0.002 | -0.004 | -0.001 | 0 |
| SEC14 | 0.011 | 0.019 | 0.015 | 0 |
| PUG1 | -0.022 | -0.007 | -0.014 | 0 |
| COG1 | 0.037 | 0.042 | 0.040 | 1 |
| YJR039W | -0.009 | -0.004 | -0.006 | 0 |
| SMI1 | 0.001 | 0.002 | 0.002 | 0 |
| KTR7 | -0.012 | -0.008 | -0.010 | 0 |
| MSW1 | 0.009 | -0.006 | 0.002 | 0 |
| CCS1 | 0.008 | -0.006 | 0.001 | 0 |
| CTK3 | 0.006 | -0.006 | 0.000 | 0 |
| BUD31 | 0.017 | 0.039 | 0.028 | 0 |
| SLF1 | -0.014 | 0.002 | -0.006 | 0 |
| ANB1 | 0.009 | -0.002 | 0.003 | 0 |

| | | | | |
|---|---|---|---|---|
| SPT20 | -0.002 | -0.018 | -0.010 | 0 |
| PPT2 | -0.002 | -0.002 | -0.002 | 0 |
| NOT5 | -0.003 | 0.011 | 0.004 | 0 |
| ROG1 | -0.019 | -0.018 | -0.018 | 0 |
| FAS1 | 0.000 | 0.006 | 0.003 | 0 |
| YGL159W | -0.018 | -0.008 | -0.013 | 0 |
| TFA2 | 0.004 | 0.026 | 0.015 | 0 |
| BUD13 | -0.029 | -0.019 | -0.024 | 0 |
| SDO1 | 0.004 | 0.018 | 0.011 | 0 |
| KAP123 | -0.029 | -0.021 | -0.025 | -1 |
| YLR132C | 0.019 | 0.027 | 0.023 | 0 |
| YER121W | 0.001 | 0.003 | 0.002 | 0 |
| YLR243W | 0.010 | 0.016 | 0.013 | 0 |
| YER140W | -0.160 | -0.213 | -0.186 | -1 |
| ADE13 | 0.008 | 0.010 | 0.009 | 0 |
| YER156C | 0.000 | 0.009 | 0.004 | 0 |
| TEM1 | 0.015 | 0.010 | 0.012 | 0 |
| GRX4 | 0.000 | -0.003 | -0.002 | 0 |
| UTP15 | 0.011 | 0.023 | 0.017 | 0 |
| YER186C | -0.010 | -0.004 | -0.007 | 0 |
| BDF1 | 0.021 | 0.000 | 0.011 | 0 |
| VPS55 | -0.010 | -0.016 | -0.013 | 0 |
| RRN10 | 0.014 | 0.009 | 0.012 | 0 |
| LYS12 | 0.003 | 0.028 | 0.016 | 0 |
| GCN2 | 0.001 | -0.001 | 0.000 | 0 |
| COX14 | -0.014 | 0.002 | -0.006 | 0 |
| LTE1 | -0.001 | -0.010 | -0.005 | 0 |
| COX16 | -0.002 | 0.016 | 0.007 | 0 |
| PTC2 | -0.047 | -0.010 | -0.029 | 0 |
| GEF1 | -0.001 | -0.011 | -0.006 | 0 |
| ENO1 | 0.007 | 0.002 | 0.005 | 0 |
| PLC1 | 0.005 | 0.009 | 0.007 | 0 |
| YOR008C-A | -0.007 | 0.001 | -0.003 | 0 |
| CTF13 | -0.003 | 0.013 | 0.005 | 0 |
| ERG8 | 0.020 | 0.018 | 0.019 | 0 |
| ABC1 | 0.006 | 0.001 | 0.004 | 0 |
| NIP1 | -0.007 | -0.023 | -0.015 | 0 |
| YKR099C-A | -0.006 | -0.003 | -0.004 | 0 |
| NAF1 | 0.003 | 0.007 | 0.005 | 0 |
| YML007C-A | -0.124 | -0.078 | -0.101 | -1 |
| NAR1 | 0.010 | 0.004 | 0.007 | 0 |
| YMR272W-B | -0.001 | 0.002 | 0.001 | 0 |
| KRI1 | -0.008 | 0.002 | -0.003 | 0 |
| DDR2 | -0.007 | -0.003 | -0.005 | 0 |
| YOL022C | 0.015 | 0.011 | 0.013 | 0 |
| YPL038W-A | -0.020 | -0.005 | -0.012 | 0 |
| MED7 | -0.003 | -0.003 | -0.003 | 0 |
| LSM7 | 0.029 | 0.024 | 0.026 | 1 |
| YAL016C-B | -0.011 | -0.004 | -0.007 | 0 |
| YBR072C-A | 0.002 | 0.000 | 0.001 | 0 |
| BUD16 | 0.001 | -0.004 | -0.002 | 0 |
| TPN1 | -0.024 | 0.008 | -0.008 | 0 |
| YDR246W-A | -0.021 | -0.007 | -0.014 | 0 |

| | | | | |
|---|---|---|---|---|
| YER175W-A | -0.006 | -0.006 | -0.006 | 0 |
| YGR121W-A | -0.009 | -0.007 | -0.008 | 0 |
| YHR086W-A | -0.005 | 0.003 | -0.001 | 0 |
| GCN5 | 0.066 | 0.040 | 0.053 | 1 |
| SPC24 | -0.004 | -0.003 | -0.004 | 0 |
| YAL044W-A | -0.011 | -0.007 | -0.009 | 0 |
| RRP5 | 0.006 | 0.008 | 0.007 | 0 |
| PCL10 | 0.002 | -0.003 | -0.001 | 0 |
| PRE5 | 0.104 | 0.105 | 0.105 | 1 |
| YLL006W-A | 0.000 | -0.003 | -0.002 | 0 |
| TOM22 | 0.036 | 0.040 | 0.038 | 1 |
| YML054C-A | -0.010 | -0.010 | -0.010 | 0 |
| SUI1 | -0.057 | 0.001 | -0.028 | 0 |
| YMR315W-A | -0.003 | -0.004 | -0.004 | 0 |
| ZIM17 | 0.020 | 0.017 | 0.018 | 0 |
| ATP19 | 0.007 | 0.005 | 0.006 | 0 |
| SMC5 | -0.004 | 0.006 | 0.001 | 0 |
| ERI1 | 0.002 | -0.002 | 0.000 | 0 |
| CDC33 | -0.004 | 0.000 | -0.002 | 0 |
| ADE12 | -0.010 | 0.003 | -0.003 | 0 |
| YAL037C-A | -0.006 | -0.011 | -0.008 | 0 |
| YBR085C-A | -0.010 | -0.046 | -0.028 | 0 |
| YCR075W-A | 0.017 | 0.012 | 0.015 | 0 |
| RPS27B | -0.005 | -0.011 | -0.008 | 0 |
| SLO1 | 0.017 | 0.011 | 0.014 | 0 |
| ANP1 | 0.039 | 0.027 | 0.033 | 1 |
| YGR146C-A | -0.006 | -0.003 | -0.004 | 0 |
| ROX3 | -0.011 | -0.014 | -0.013 | 0 |
| YHR175W-A | -0.012 | 0.002 | -0.005 | 0 |
| SSB2 | -0.023 | -0.013 | -0.018 | 0 |
| YJL136W-A | -0.006 | 0.000 | -0.003 | 0 |
| ECM16 | 0.005 | 0.012 | 0.009 | 0 |
| RNA1 | 0.061 | 0.045 | 0.053 | 1 |
| MPT5 | -0.013 | -0.009 | -0.011 | 0 |
| RLP7 | 0.019 | 0.015 | 0.017 | 0 |
| YLR264C-A | -0.030 | -0.025 | -0.027 | -1 |
| RPC31 | -0.001 | 0.011 | 0.005 | 0 |
| HUG1 | -0.011 | -0.007 | -0.009 | 0 |
| CWC25 | 0.027 | 0.026 | 0.027 | 1 |
| YNL042W-B | -0.009 | 0.000 | -0.005 | 0 |
| RFA2 | 0.015 | 0.011 | 0.013 | 0 |
| YOL086W-A | 0.002 | 0.000 | 0.001 | 0 |
| PRE6 | 0.086 | 0.101 | 0.094 | 1 |
| YPL119C-A | 0.002 | -0.013 | -0.005 | 0 |
| NOP8 | 0.022 | 0.017 | 0.019 | 0 |
| MRPL10 | 0.016 | 0.014 | 0.015 | 0 |
| YAL067W-A | -0.010 | -0.004 | -0.007 | 0 |
| SUS1 | 0.012 | 0.004 | 0.008 | 0 |
| YDL085C-A | 0.000 | 0.001 | 0.000 | 0 |
| TIM11 | 0.014 | 0.013 | 0.013 | 0 |
| BUD25 | 0.010 | 0.004 | 0.007 | 0 |
| YGR169C-A | -0.004 | -0.002 | -0.003 | 0 |
| YLR419W | 0.000 | 0.000 | 0.000 | 0 |

| | | | | |
|---|---|---|---|---|
| YIL002W-A | -0.002 | 0.010 | 0.004 | 0 |
| COQ6 | -0.293 | -0.029 | -0.161 | -1 |
| RRB1 | 0.021 | 0.017 | 0.019 | 0 |
| YCL021W-A | -0.016 | -0.017 | -0.016 | 0 |
| TAF9 | -0.013 | 0.001 | -0.006 | 0 |
| STR3 | -0.009 | -0.006 | -0.007 | 0 |
| LST8 | 0.020 | 0.017 | 0.019 | 0 |
| ISA1 | 0.000 | -0.013 | -0.007 | 0 |
| PGA1 | -0.004 | 0.000 | -0.002 | 0 |
| YMR001C-A | -0.012 | -0.005 | -0.008 | 0 |
| YNL247W | 0.002 | 0.013 | 0.007 | 0 |
| YNL067W-B | -0.006 | -0.001 | -0.003 | 0 |
| YNL313C | 0.007 | 0.011 | 0.009 | 0 |
| YOL097W-A | -0.005 | -0.009 | -0.007 | 0 |
| RIB2 | -0.041 | -0.028 | -0.035 | -1 |
| YPL152W-A | -0.003 | 0.001 | -0.001 | 0 |
| PSF3 | 0.013 | 0.016 | 0.015 | 0 |
| YAR035C-A | -0.009 | -0.009 | -0.009 | 0 |
| YBR182C-A | -0.002 | 0.000 | -0.001 | 0 |
| YDL159W-A | 0.005 | 0.004 | 0.004 | 0 |
| BUR2 | -0.001 | -0.004 | -0.002 | 0 |
| YDR379C-A | -0.013 | -0.004 | -0.008 | 0 |
| YJR085C | -0.013 | -0.010 | -0.011 | 0 |
| YFL041W-A | 0.000 | 0.001 | 0.001 | 0 |
| YGR174W-A | 0.008 | 0.003 | 0.005 | 0 |
| YIL046W-A | 0.005 | 0.002 | 0.004 | 0 |
| YJR005C-A | -0.017 | -0.017 | -0.017 | 0 |
| YPL183W | 0.008 | 0.002 | 0.005 | 0 |
| YMR134W | 0.015 | 0.012 | 0.013 | 0 |
| HBN1 | -0.006 | 0.003 | -0.002 | 0 |
| CUS1 | 0.016 | 0.018 | 0.017 | 0 |
| YGL185C | -0.009 | -0.006 | -0.008 | 0 |
| GPI15 | 0.028 | 0.017 | 0.023 | 0 |
| YLR285C-A | 0.011 | 0.003 | 0.007 | 0 |
| YMR013W-A | -0.011 | -0.004 | -0.007 | 0 |
| FOL1 | 0.015 | 0.009 | 0.012 | 0 |
| YNL097C-A | -0.002 | -0.001 | -0.002 | 0 |
| PFS2 | 0.031 | 0.029 | 0.030 | 1 |
| YOL159C-A | -0.006 | -0.001 | -0.004 | 0 |
| NUF2 | -0.061 | 0.002 | -0.029 | 0 |
| COA2 | 0.004 | 0.005 | 0.005 | 0 |
| UTP23 | -0.002 | -0.001 | -0.001 | 0 |
| MDM32 | -0.018 | -0.017 | -0.017 | 0 |
| YBL008W-A | 0.006 | -0.002 | 0.002 | 0 |
| YBR196C-A | -0.020 | -0.027 | -0.024 | -1 |
| YDL160C-A | -0.001 | -0.006 | -0.003 | 0 |
| RPS12 | 0.001 | 0.001 | 0.001 | 0 |
| YDR524C-B | 0.021 | 0.022 | 0.021 | 1 |
| STE18 | -0.007 | -0.011 | -0.009 | 0 |
| YFR012W-A | -0.010 | -0.005 | -0.008 | 0 |
| YGR204C-A | 0.000 | 0.000 | 0.000 | 0 |
| YIL134C-A | 0.007 | 0.009 | 0.008 | 0 |
| TIM8 | -0.010 | 0.004 | -0.003 | 0 |

| | | | | |
|---|---|---|---|---|
| SWP1 | 0.011 | -0.009 | 0.001 | 0 |
| TIF11 | -0.010 | 0.015 | 0.002 | 0 |
| CDC55 | -0.013 | 0.001 | -0.006 | 0 |
| NOP2 | 0.009 | 0.022 | 0.015 | 0 |
| YLR307C-A | 0.003 | -0.004 | 0.000 | 0 |
| APC1 | 0.005 | 0.018 | 0.012 | 0 |
| YMR105W-A | -0.012 | -0.007 | -0.010 | 0 |
| DSL1 | 0.001 | 0.005 | 0.003 | 0 |
| DGR1 | -0.001 | -0.002 | -0.001 | 0 |
| RPC34 | 0.020 | 0.019 | 0.020 | 0 |
| YOL164W-A | 0.005 | 0.001 | 0.003 | 0 |
| BRX1 | -0.019 | 0.003 | -0.008 | 0 |
| YPR108W-A | -0.017 | -0.006 | -0.011 | 0 |
| RAT1 | 0.019 | 0.007 | 0.013 | 0 |
| PUS4 | 0.013 | 0.010 | 0.011 | 0 |
| YBL029C-A | -0.002 | 0.002 | 0.000 | 0 |
| YBR196C-B | -0.018 | -0.002 | -0.010 | 0 |
| YDR003W-A | -0.009 | -0.002 | -0.006 | 0 |
| MDJ1 | -0.010 | -0.079 | -0.045 | 0 |
| YDR524W-A | 0.020 | 0.012 | 0.016 | 0 |
| RSM22 | 0.024 | 0.001 | 0.012 | 0 |
| YFR032C-B | -0.004 | 0.007 | 0.001 | 0 |
| YNL296W | 0.016 | 0.003 | 0.010 | 0 |
| TOM5 | 0.011 | 0.007 | 0.009 | 0 |
| YIR018C-A | -0.009 | 0.002 | -0.003 | 0 |
| YJR151W-A | 0.004 | -0.003 | 0.000 | 0 |
| GCR1 | -0.049 | -0.003 | -0.026 | 0 |
| CEP3 | 0.013 | 0.021 | 0.017 | 0 |
| PRP24 | 0.004 | 0.013 | 0.008 | 0 |
| COX13 | -0.009 | -0.003 | -0.006 | 0 |
| IMP4 | -0.001 | 0.011 | 0.005 | 0 |
| YLR312C-B | 0.001 | -0.003 | -0.001 | 0 |
| YNL181W | 0.028 | 0.019 | 0.024 | 0 |
| YMR175W-A | -0.008 | -0.006 | -0.007 | 0 |
| YNL260C | 0.010 | 0.018 | 0.014 | 0 |
| YNL146C-A | -0.004 | -0.001 | -0.002 | 0 |
| PRP2 | 0.002 | 0.006 | 0.004 | 0 |
| YOR020W-A | 0.000 | 0.000 | 0.000 | 0 |
| AVO1 | 0.001 | 0.011 | 0.006 | 0 |
| YPR159C-A | -0.001 | 0.001 | 0.000 | 0 |
| YOR060C | 0.015 | 0.004 | 0.009 | 0 |
| SAP30 | 0.033 | 0.031 | 0.032 | 1 |
| YBL039W-A | -0.065 | -0.061 | -0.063 | -1 |
| YBR200W-A | -0.012 | -0.003 | -0.008 | 0 |
| YER087W | 0.042 | 0.044 | 0.043 | 1 |
| RPL27A | 0.003 | 0.009 | 0.006 | 0 |
| MTM1 | -0.006 | -0.012 | -0.009 | 0 |
| CLC1 | -0.007 | -0.010 | -0.009 | 0 |
| RSM26 | 0.004 | 0.019 | 0.011 | 0 |
| YGL006W-A | 0.000 | -0.004 | -0.002 | 0 |
| EFG1 | 0.013 | -0.008 | 0.002 | 0 |
| YIR021W-A | -0.008 | 0.001 | -0.003 | 0 |
| ARG8 | -0.015 | -0.024 | -0.020 | 0 |

| | | | | |
|---|---|---|---|---|
| YKL018C-A | 0.008 | 0.000 | 0.004 | 0 |
| YMR185W | 0.006 | -0.002 | 0.002 | 0 |
| PRS2 | 0.000 | -0.006 | -0.003 | 0 |
| HSH155 | 0.002 | 0.002 | 0.002 | 0 |
| IME4 | -0.009 | 0.000 | -0.004 | 0 |
| POL1 | 0.005 | 0.009 | 0.007 | 0 |
| YLR342W-A | 0.002 | -0.002 | 0.000 | 0 |
| IPI3 | 0.013 | 0.007 | 0.010 | 0 |
| YMR182W-A | -0.021 | -0.015 | -0.018 | 0 |
| POL2 | 0.000 | 0.004 | 0.002 | 0 |
| YNL162W-A | -0.002 | 0.007 | 0.003 | 0 |
| TIM23 | 0.020 | 0.014 | 0.017 | 0 |
| YOR034C-A | -0.012 | -0.001 | -0.006 | 0 |
| RFC4 | 0.010 | 0.010 | 0.010 | 0 |
| CYC8 | -0.015 | -0.014 | -0.014 | 0 |
| RTS2 | 0.012 | 0.003 | 0.007 | 0 |
| MCK1 | -0.061 | -0.032 | -0.046 | -1 |
| YBL071C-B | -0.033 | -0.041 | -0.037 | -1 |
| YBR221W-A | 0.002 | 0.001 | 0.002 | 0 |
| YDR034W-B | -0.005 | -0.010 | -0.007 | 0 |
| SOM1 | 0.017 | 0.007 | 0.012 | 0 |
| YJR112W-A | -0.003 | 0.005 | 0.001 | 0 |
| YGL007C-A | -0.008 | 0.005 | -0.001 | 0 |
| YHL015W-A | -0.002 | -0.007 | -0.005 | 0 |
| YJL012C-A | 0.009 | 0.000 | 0.004 | 0 |
| MRPL20 | 0.040 | 0.031 | 0.036 | 1 |
| YKL068W-A | -0.008 | -0.010 | -0.009 | 0 |
| RPL36B | 0.012 | 0.000 | 0.006 | 0 |
| ROT1 | 0.011 | 0.009 | 0.010 | 0 |
| NUP157 | -0.001 | -0.008 | -0.004 | 0 |
| HAS1 | 0.008 | 0.007 | 0.007 | 0 |
| MET4 | -0.006 | 0.008 | 0.001 | 0 |
| YLR361C-A | 0.005 | -0.006 | -0.001 | 0 |
| RIO2 | 0.019 | 0.016 | 0.018 | 0 |
| YMR194C-B | -0.009 | -0.008 | -0.009 | 0 |
| YIF1 | 0.020 | 0.026 | 0.023 | 1 |
| YNL277W-A | -0.004 | 0.000 | -0.002 | 0 |
| SEC12 | 0.115 | 0.100 | 0.107 | 1 |
| YOR161C-C | -0.009 | 0.005 | -0.002 | 0 |
| WRS1 | 0.053 | 0.045 | 0.049 | 1 |
| TIF4631 | 0.006 | 0.006 | 0.006 | 0 |
| RKI1 | 0.019 | 0.021 | 0.020 | 0 |
| MET2 | -0.023 | -0.009 | -0.016 | 0 |
| KTI11 | 0.070 | 0.068 | 0.069 | 1 |
| YBR230W-A | -0.004 | -0.004 | -0.004 | 0 |
| IES6 | 0.030 | 0.025 | 0.027 | 1 |
| TFB5 | 0.013 | 0.004 | 0.008 | 0 |
| YER053C-A | 0.005 | -0.005 | 0.000 | 0 |
| YJR114W | -0.072 | -0.117 | -0.095 | -1 |
| YGL041C-B | -0.006 | 0.001 | -0.003 | 0 |
| YHR007C-A | -0.002 | -0.002 | -0.002 | 0 |
| HTB2 | -0.008 | 0.002 | -0.003 | 0 |
| YJL047C-A | 0.012 | -0.003 | 0.005 | 0 |

| | | | | |
|---|---|---|---|---|
| CTR9 | -0.007 | -0.046 | -0.027 | 0 |
| YKL096C-B | -0.004 | -0.003 | -0.004 | 0 |
| YPL277C | -0.012 | -0.001 | -0.007 | 0 |
| ERG12 | 0.000 | 0.001 | 0.001 | 0 |
| LIP1 | 0.030 | 0.031 | 0.031 | 1 |
| NOP15 | 0.001 | 0.015 | 0.008 | 0 |
| YLR363W-A | -0.006 | -0.007 | -0.006 | 0 |
| RAP1 | 0.016 | 0.009 | 0.012 | 0 |
| YMR230W-A | -0.002 | -0.005 | -0.003 | 0 |
| POP3 | 0.008 | 0.010 | 0.009 | 0 |
| YOL013W-B | -0.004 | 0.002 | -0.001 | 0 |
| TRM112 | -0.003 | 0.001 | -0.001 | 0 |
| YOR293C-A | -0.020 | 0.002 | -0.009 | 0 |
| TPT1 | 0.270 | 0.291 | 0.281 | 1 |
| YGR272C | 0.013 | 0.016 | 0.014 | 0 |
| OST2 | -0.010 | -0.014 | -0.012 | 0 |
| YBL101W-C | -0.005 | -0.007 | -0.006 | 0 |
| YBR296C-A | -0.007 | 0.003 | -0.002 | 0 |
| YDR169C-A | -0.007 | -0.006 | -0.006 | 0 |
| POR1 | 0.036 | 0.024 | 0.030 | 1 |
| YCL007C | -0.004 | -0.009 | -0.006 | 0 |
| PMT4 | 0.003 | 0.004 | 0.003 | 0 |
| YGL188C-A | -0.019 | -0.008 | -0.013 | 0 |
| YHR022C-A | -0.001 | -0.005 | -0.003 | 0 |
| YJL062W-A | -0.010 | 0.002 | -0.004 | 0 |
| ZPS1 | -0.002 | 0.000 | -0.001 | 0 |
| YKL106C-A | -0.002 | 0.003 | 0.000 | 0 |
| DML1 | -0.007 | -0.002 | -0.005 | 0 |
| YER186W-A | 0.002 | -0.010 | -0.004 | 0 |
| ATM1 | 0.009 | 0.012 | 0.010 | 0 |
| DBP2 | 0.011 | 0.009 | 0.010 | 0 |
| YLR406C-A | -0.003 | 0.004 | 0.000 | 0 |
| SSU72 | -0.011 | -0.004 | -0.007 | 0 |
| YMR242W-A | 0.024 | 0.026 | 0.025 | 1 |
| SEC21 | -0.005 | -0.002 | -0.004 | 0 |
| YOL019W-A | -0.011 | -0.010 | -0.011 | 0 |
| RPB11 | 0.032 | 0.030 | 0.031 | 1 |
| YOR316C-A | -0.008 | 0.003 | -0.002 | 0 |
| ALR1 | 0.015 | 0.015 | 0.015 | 0 |
| TFC7 | 0.008 | 0.001 | 0.005 | 0 |
| YBR056W-A | -0.014 | -0.008 | -0.011 | 0 |
| ADE2 | -0.002 | -0.004 | -0.003 | 0 |
| YCL001W-B | 0.000 | 0.000 | 0.000 | 0 |
| YDR182W-A | -0.011 | -0.002 | -0.007 | 0 |
| PEP3 | 0.005 | 0.002 | 0.004 | 0 |
| DAN4 | 0.004 | -0.001 | 0.002 | 0 |
| RNR1 | 0.016 | 0.007 | 0.011 | 0 |
| YHR050W-A | 0.010 | 0.004 | 0.007 | 0 |
| NAT3 | 0.032 | 0.016 | 0.024 | 0 |
| YJL077W-B | 0.001 | 0.002 | 0.001 | 0 |
| BDS1 | -0.005 | 0.000 | -0.002 | 0 |
| ARP5 | -0.015 | -0.019 | -0.017 | 0 |
| CEF1 | 0.006 | 0.011 | 0.009 | 0 |

| | | | | |
|---|---|---|---|---|
| SEH1 | -0.010 | -0.020 | -0.015 | 0 |
| PSE1 | 0.010 | 0.012 | 0.011 | 0 |
| YKR004C-A | -0.018 | -0.009 | -0.013 | 0 |
| RPC19 | 0.000 | 0.006 | 0.003 | 0 |
| YLR412C-A | -0.004 | 0.000 | -0.002 | 0 |
| CSL4 | 0.012 | 0.006 | 0.009 | 0 |
| YMR247W-A | 0.001 | 0.000 | 0.000 | 0 |
| RFC3 | -0.005 | 0.006 | 0.001 | 0 |
| YOL038C-A | -0.014 | -0.006 | -0.010 | 0 |
| DIS3 | -0.004 | 0.003 | 0.000 | 0 |
| YOR376W-A | 0.002 | -0.002 | 0.000 | 0 |
| HRT1 | 0.023 | 0.034 | 0.029 | 1 |
| RPT5 | 0.083 | 0.070 | 0.076 | 1 |
| TSC3 | 0.036 | 0.019 | 0.027 | 0 |
| YCL012C | -0.002 | -0.004 | -0.003 | 0 |
| YCL057C-A | 0.034 | 0.026 | 0.030 | 1 |
| YDR194W-A | -0.009 | -0.004 | -0.007 | 0 |
| SBH1 | 0.017 | 0.018 | 0.018 | 0 |
| YGR035W-A | -0.011 | -0.010 | -0.011 | 0 |
| CGR1 | -0.009 | 0.008 | 0.000 | 0 |
| HPR1 | 0.044 | 0.028 | 0.036 | 1 |
| YJL127C-B | -0.014 | -0.001 | -0.007 | 0 |
| APC4 | 0.003 | -0.010 | -0.003 | 0 |
| YPT1 | -0.029 | -0.040 | -0.035 | -1 |
| YDR196C | -0.008 | -0.004 | -0.006 | 0 |
| MTR3 | 0.001 | -0.004 | -0.002 | 0 |
| BFR2 | 0.002 | 0.017 | 0.010 | 0 |
| MCD4 | -0.015 | -0.008 | -0.011 | 0 |
| NCB2 | 0.004 | -0.004 | 0.000 | 0 |
| TAF11 | -0.021 | -0.002 | -0.011 | 0 |
| SEC20 | -0.010 | -0.006 | -0.008 | 0 |
| KAR1 | -0.004 | 0.004 | 0.000 | 0 |
| PRO3 | 0.004 | 0.015 | 0.009 | 0 |
| SGT1 | -0.013 | -0.016 | -0.014 | 0 |
| SMC1 | 0.001 | 0.002 | 0.001 | 0 |
| MEX67 | -0.010 | -0.002 | -0.006 | 0 |
| PMA1 | 0.010 | 0.001 | 0.006 | 0 |
| CDC24 | -0.022 | -0.029 | -0.025 | -1 |
| CEG1 | 0.039 | 0.061 | 0.050 | 1 |
| PRE7 | 0.008 | -0.003 | 0.002 | 0 |
| SMD1 | 0.038 | 0.055 | 0.046 | 1 |
| RPG1 | 0.006 | 0.001 | 0.003 | 0 |
| MES1 | 0.000 | -0.002 | -0.001 | 0 |
| RPB5 | -0.005 | -0.003 | -0.004 | 0 |
| UTP9 | 0.004 | -0.013 | -0.005 | 0 |
| TSC10 | -0.004 | -0.003 | -0.004 | 0 |
| NUP159 | -0.005 | -0.007 | -0.006 | 0 |
| NOP1 | -0.021 | -0.029 | -0.025 | -1 |
| RPC17 | -0.010 | -0.011 | -0.011 | 0 |
| CDC48 | -0.004 | 0.001 | -0.002 | 0 |
| DPB11 | 0.011 | 0.018 | 0.015 | 0 |
| YPD1 | 0.009 | 0.011 | 0.010 | 0 |
| RFC2 | -0.003 | 0.000 | -0.002 | 0 |

| | | | | |
|---|---|---|---|---|
| BRN1 | -0.001 | -0.007 | -0.004 | 0 |
| DOP1 | -0.003 | -0.004 | -0.004 | 0 |
| NIC96 | -0.008 | -0.004 | -0.006 | 0 |
| SPC19 | -0.007 | 0.007 | 0.000 | 0 |
| BRF1 | -0.001 | -0.004 | -0.003 | 0 |
| CFT1 | 0.004 | 0.000 | 0.002 | 0 |
| SNU114 | -0.011 | -0.002 | -0.007 | 0 |
| UTP5 | 0.000 | -0.002 | -0.001 | 0 |
| PRE8 | 0.074 | 0.089 | 0.081 | 1 |
| LCD1 | 0.013 | 0.001 | 0.007 | 0 |
| SRP1 | 0.003 | -0.013 | -0.005 | 0 |
| GCD11 | 0.006 | -0.004 | 0.001 | 0 |
| PNO1 | -0.001 | 0.000 | 0.000 | 0 |
| CDC4 | -0.020 | 0.001 | -0.009 | 0 |
| SPT14 | 0.013 | 0.012 | 0.013 | 0 |
| SCL1 | 0.082 | 0.092 | 0.087 | 1 |
| PTA1 | -0.008 | -0.021 | -0.015 | 0 |
| SEC27 | -0.004 | -0.004 | -0.004 | 0 |
| SEC17 | -0.029 | -0.019 | -0.024 | 0 |
| GCD2 | -0.017 | -0.004 | -0.011 | 0 |
| SEC18 | -0.012 | 0.004 | -0.004 | 0 |
| FOL2 | -0.015 | -0.008 | -0.011 | 0 |
| CDC28 | -0.002 | 0.005 | 0.002 | 0 |
| RIX1 | -0.003 | -0.005 | -0.004 | 0 |
| NFS1 | -0.001 | 0.005 | 0.002 | 0 |
| TAO3 | 0.008 | -0.001 | 0.003 | 0 |
| TSC13 | 0.011 | -0.007 | 0.002 | 0 |
| RRN7 | -0.005 | -0.002 | -0.004 | 0 |
| CCT4 | -0.007 | 0.014 | 0.003 | 0 |
| PHS1 | -0.005 | 0.010 | 0.002 | 0 |
| YRB1 | -0.005 | 0.004 | -0.001 | 0 |
| NPA3 | 0.006 | 0.005 | 0.006 | 0 |
| PGS1 | -0.001 | -0.015 | -0.008 | 0 |
| SSY1 | -0.011 | 0.000 | -0.006 | 0 |
| RPN11 | 0.001 | -0.001 | 0.000 | 0 |
| MSS4 | 0.006 | 0.010 | 0.008 | 0 |
| SAM35 | 0.009 | 0.007 | 0.008 | 0 |
| GPI11 | -0.033 | -0.015 | -0.024 | 0 |
| SPC34 | 0.023 | 0.018 | 0.021 | 0 |
| TRS120 | 0.004 | 0.008 | 0.006 | 0 |
| TAF13 | 0.048 | 0.066 | 0.057 | 1 |
| SMT3 | 0.010 | 0.004 | 0.007 | 0 |
| POP1 | -0.005 | -0.013 | -0.009 | 0 |
| ARB1 | -0.013 | -0.011 | -0.012 | 0 |
| RPB2 | 0.004 | 0.000 | 0.002 | 0 |
| GNA1 | 0.006 | -0.001 | 0.002 | 0 |
| RVB2 | -0.021 | -0.023 | -0.022 | -1 |
| JAC1 | -0.036 | -0.023 | -0.029 | -1 |
| RFA1 | 0.001 | -0.008 | -0.003 | 0 |
| ROK1 | 0.004 | 0.011 | 0.008 | 0 |
| ILS1 | -0.004 | 0.003 | 0.000 | 0 |
| UTP22 | -0.023 | -0.010 | -0.017 | 0 |
| RFC5 | -0.002 | -0.005 | -0.004 | 0 |

| | | | | |
|---|---|---|---|---|
| RPS20 | -0.009 | -0.005 | -0.007 | 0 |
| POP7 | -0.016 | -0.006 | -0.011 | 0 |
| FAF1 | 0.008 | 0.005 | 0.007 | 0 |
| RRP7 | -0.008 | 0.003 | -0.002 | 0 |
| SLN1 | -0.077 | -0.091 | -0.084 | -1 |
| CDC7 | -0.014 | -0.007 | -0.011 | 0 |
| BET4 | 0.002 | 0.009 | 0.006 | 0 |
| NOP14 | -0.002 | -0.006 | -0.004 | 0 |
| TIM17 | 0.013 | 0.016 | 0.014 | 0 |
| KRS1 | 0.018 | 0.007 | 0.012 | 0 |
| CDC11 | -0.020 | -0.018 | -0.019 | 0 |
| CDC53 | -0.007 | -0.010 | -0.008 | 0 |
| SEC1 | -0.021 | -0.019 | -0.020 | 0 |
| RET2 | 0.005 | 0.001 | 0.003 | 0 |
| GCD6 | 0.004 | 0.009 | 0.007 | 0 |
| NMD3 | 0.004 | -0.001 | 0.001 | 0 |
| SRB7 | 0.026 | 0.020 | 0.023 | 0 |
| DAD2 | 0.011 | 0.010 | 0.010 | 0 |
| TIF35 | 0.044 | 0.006 | 0.025 | 0 |
| TAF4 | 0.014 | 0.014 | 0.014 | 0 |
| RBA50 | -0.002 | -0.003 | -0.003 | 0 |
| NRD1 | 0.000 | -0.009 | -0.004 | 0 |
| KRE29 | -0.008 | 0.001 | -0.004 | 0 |
| TOA1 | 0.009 | 0.024 | 0.016 | 0 |
| FRS2 | -0.046 | 0.004 | -0.021 | 0 |
| IQG1 | 0.012 | 0.005 | 0.008 | 0 |
| STT3 | 0.012 | 0.009 | 0.011 | 0 |
| SEN34 | -0.001 | -0.006 | -0.004 | 0 |
| MCM6 | -0.013 | -0.014 | -0.014 | 0 |
| CDC27 | -0.004 | 0.012 | 0.004 | 0 |
| PRP31 | 0.011 | -0.004 | 0.003 | 0 |
| POL30 | 0.009 | -0.005 | 0.002 | 0 |
| MED8 | -0.045 | -0.059 | -0.052 | -1 |
| TIM44 | -0.012 | -0.005 | -0.009 | 0 |
| PDI1 | -0.014 | -0.017 | -0.016 | 0 |
| MCM10 | 0.011 | 0.011 | 0.011 | 0 |
| MPS1 | -0.001 | -0.002 | -0.002 | 0 |
| HCA4 | -0.017 | -0.019 | -0.018 | 0 |
| SAS10 | -0.019 | -0.014 | -0.017 | 0 |
| ERG20 | 0.000 | 0.007 | 0.003 | 0 |
| HEM13 | 0.017 | 0.011 | 0.014 | 0 |
| FIP1 | -0.004 | 0.000 | -0.002 | 0 |
| RPO21 | -0.009 | -0.006 | -0.008 | 0 |
| TAF10 | 0.046 | 0.049 | 0.048 | 1 |
| RNA15 | 0.001 | -0.001 | 0.000 | 0 |
| HTB1 | 0.008 | 0.001 | 0.004 | 0 |
| RPB3 | 0.022 | 0.019 | 0.020 | 0 |
| GPI8 | -0.016 | -0.014 | -0.015 | 0 |
| SFI1 | 0.008 | 0.006 | 0.007 | 0 |
| GPI17 | 0.008 | 0.015 | 0.012 | 0 |
| MCM1 | 0.000 | -0.010 | -0.005 | 0 |
| YDR531W | -0.009 | -0.004 | -0.007 | 0 |
| ORC5 | -0.004 | 0.002 | -0.001 | 0 |

| | | | | |
|---|---|---|---|---|
| SAH1 | -0.012 | -0.010 | -0.011 | 0 |
| MGE1 | -0.003 | 0.008 | 0.003 | 0 |
| EPL1 | 0.004 | 0.007 | 0.006 | 0 |
| YAH1 | 0.016 | 0.015 | 0.015 | 0 |
| HEM2 | 0.004 | 0.009 | 0.006 | 0 |
| CDC15 | -0.020 | -0.029 | -0.024 | 0 |
| VRG4 | -0.032 | -0.031 | -0.032 | -1 |
| RPL32 | -0.032 | -0.032 | -0.032 | -1 |
| RRP46 | 0.000 | -0.006 | -0.003 | 0 |
| MRS5 | -0.039 | -0.048 | -0.043 | -1 |
| BCD1 | -0.005 | -0.014 | -0.010 | 0 |
| PGI1 | 0.024 | 0.013 | 0.018 | 0 |
| IRR1 | -0.006 | -0.011 | -0.008 | 0 |
| SPB1 | 0.003 | -0.001 | 0.001 | 0 |
| YIL171W | -0.013 | -0.004 | -0.009 | 0 |
| ARP2 | 0.005 | 0.018 | 0.011 | 0 |
| KAR2 | -0.085 | -0.090 | -0.088 | -1 |
| CDC36 | 0.001 | 0.001 | 0.001 | 0 |
| KRE9 | 0.005 | 0.015 | 0.010 | 0 |
| RPC11 | 0.021 | 0.023 | 0.022 | 1 |
| YJR141W | -0.003 | 0.000 | -0.002 | 0 |
| CDC37 | -0.001 | -0.002 | -0.002 | 0 |
| OLE1 | -0.020 | -0.015 | -0.017 | 0 |
| HEM1 | 0.007 | 0.002 | 0.005 | 0 |
| STH1 | 0.001 | 0.001 | 0.001 | 0 |
| FCF1 | -0.013 | -0.006 | -0.010 | 0 |
| DRS1 | 0.002 | -0.010 | -0.004 | 0 |
| GUK1 | -0.001 | -0.001 | -0.001 | 0 |
| RNA14 | 0.017 | 0.008 | 0.013 | 0 |
| WBP1 | -0.012 | -0.010 | -0.011 | 0 |
| ACC1 | -0.011 | -0.011 | -0.011 | 0 |
| LSM4 | -0.014 | -0.009 | -0.012 | 0 |
| GCD1 | -0.007 | 0.006 | 0.000 | 0 |
| YPI1 | 0.004 | 0.011 | 0.008 | 0 |
| ARP7 | 0.021 | 0.013 | 0.017 | 0 |
| RPT6 | 0.103 | 0.102 | 0.102 | 1 |
| UTP20 | -0.011 | -0.006 | -0.009 | 0 |
| SEC15 | -0.011 | -0.005 | -0.008 | 0 |
| PKC1 | -0.012 | -0.016 | -0.014 | 0 |
| TEL2 | 0.009 | 0.000 | 0.004 | 0 |
| EXO84 | -0.002 | -0.004 | -0.003 | 0 |
| NCP1 | -0.005 | 0.008 | 0.001 | 0 |
| MCM7 | 0.002 | 0.000 | 0.001 | 0 |
| ULP2 | -0.014 | -0.021 | -0.017 | 0 |
| KRR1 | -0.005 | -0.028 | -0.016 | 0 |
| PRI1 | 0.040 | 0.041 | 0.040 | 1 |
| FAD1 | 0.001 | 0.006 | 0.003 | 0 |
| TIM54 | -0.003 | -0.007 | -0.005 | 0 |
| NUS1 | -0.002 | 0.002 | 0.000 | 0 |
| PRP21 | 0.004 | -0.003 | 0.001 | 0 |
| DBF4 | -0.015 | -0.002 | -0.008 | 0 |
| PRP40 | -0.001 | 0.005 | 0.002 | 0 |
| SLU7 | -0.017 | -0.015 | -0.016 | 0 |

| | | | | |
|---|---|---|---|---|
| SEC7 | -0.038 | -0.037 | -0.037 | -1 |
| HSF1 | -0.002 | -0.009 | -0.005 | 0 |
| PRP42 | 0.005 | -0.004 | 0.000 | 0 |
| CCT2 | 0.033 | 0.031 | 0.032 | 1 |
| YDR341C | 0.023 | 0.020 | 0.021 | 0 |
| PAM18 | -0.002 | 0.001 | -0.001 | 0 |
| TFB3 | -0.006 | -0.001 | -0.004 | 0 |
| TIF34 | 0.023 | 0.031 | 0.027 | 1 |
| MCM3 | -0.010 | 0.000 | -0.005 | 0 |
| ARC35 | 0.024 | 0.018 | 0.021 | 0 |
| RSP5 | -0.021 | -0.040 | -0.031 | -1 |
| RPA190 | 0.009 | -0.003 | 0.003 | 0 |
| SAD1 | 0.012 | 0.014 | 0.013 | 0 |
| TIF5 | 0.008 | 0.014 | 0.011 | 0 |
| DUO1 | -0.006 | -0.006 | -0.006 | 0 |
| RRN6 | -0.007 | 0.001 | -0.003 | 0 |
| CSE1 | -0.015 | -0.022 | -0.018 | 0 |
| GPI18 | 0.006 | 0.000 | 0.003 | 0 |
| COG2 | -0.002 | -0.009 | -0.005 | 0 |
| CMD1 | -0.066 | -0.075 | -0.071 | -1 |
| MED6 | 0.000 | 0.004 | 0.002 | 0 |
| ARC40 | -0.037 | 0.040 | 0.002 | 0 |
| MMF1 | -0.006 | -0.014 | -0.010 | 0 |
| RSC6 | -0.001 | -0.004 | -0.002 | 0 |
| SEC11 | -0.036 | -0.039 | -0.038 | -1 |
| TSR1 | -0.001 | -0.002 | -0.001 | 0 |
| NUP82 | -0.005 | -0.009 | -0.007 | 0 |
| SEC31 | 0.003 | -0.007 | -0.002 | 0 |
| SUI2 | 0.009 | -0.003 | 0.003 | 0 |
| CDC34 | -0.004 | 0.010 | 0.003 | 0 |
| RAM2 | -0.007 | 0.006 | 0.000 | 0 |
| TAF12 | 0.002 | 0.005 | 0.004 | 0 |
| SUP35 | -0.001 | -0.002 | -0.002 | 0 |
| LSG1 | -0.001 | 0.000 | 0.000 | 0 |
| SEC26 | 0.004 | -0.002 | 0.001 | 0 |
| DSN1 | -0.010 | -0.003 | -0.006 | 0 |
| TFC6 | 0.004 | 0.003 | 0.004 | 0 |
| MSL5 | 0.001 | 0.003 | 0.002 | 0 |
| SPP41 | -0.006 | -0.007 | -0.006 | 0 |
| TOM40 | 0.002 | 0.003 | 0.002 | 0 |
| PCM1 | -0.019 | -0.016 | -0.018 | 0 |
| MVD1 | -0.009 | 0.004 | -0.003 | 0 |
| NSA2 | 0.002 | -0.003 | 0.000 | 0 |
| NUD1 | -0.001 | -0.008 | -0.005 | 0 |
| PTR3 | 0.017 | 0.003 | 0.010 | 0 |
| SGV1 | 0.005 | 0.010 | 0.007 | 0 |
| MNP1 | -0.013 | 0.000 | -0.007 | 0 |
| POP8 | 0.000 | 0.005 | 0.002 | 0 |
| SNU71 | -0.004 | 0.005 | 0.001 | 0 |
| IPP1 | 0.061 | 0.053 | 0.057 | 1 |
| ERG1 | -0.006 | 0.001 | -0.003 | 0 |
| ALG1 | -0.010 | 0.002 | -0.004 | 0 |
| TRM5 | 0.008 | 0.001 | 0.005 | 0 |

| | | | | |
|---|---|---|---|---|
| ALG7 | 0.002 | 0.009 | 0.006 | 0 |
| SNP1 | 0.003 | 0.006 | 0.005 | 0 |
| CTR86 | -0.007 | 0.000 | -0.004 | 0 |
| PRE3 | -0.015 | -0.010 | -0.013 | 0 |
| RPN6 | 0.073 | 0.074 | 0.073 | 1 |
| PSF2 | 0.011 | 0.008 | 0.009 | 0 |
| HEM3 | 0.011 | 0.002 | 0.007 | 0 |
| ILV3 | 0.000 | -0.006 | -0.003 | 0 |
| LCB2 | -0.002 | 0.001 | 0.000 | 0 |
| MAK11 | -0.020 | -0.003 | -0.011 | 0 |
| YCG1 | -0.004 | 0.004 | 0.000 | 0 |
| UBC1 | -0.013 | -0.005 | -0.009 | 0 |
| PRP43 | 0.009 | 0.007 | 0.008 | 0 |
| SNU56 | 0.006 | 0.000 | 0.003 | 0 |
| SQT1 | 0.001 | 0.004 | 0.002 | 0 |
| YDR367W | -0.001 | 0.002 | 0.001 | 0 |
| MAS1 | -0.011 | -0.006 | -0.009 | 0 |
| TLG1 | -0.011 | -0.016 | -0.014 | 0 |
| RNT1 | -0.019 | -0.014 | -0.016 | 0 |
| PMI40 | 0.006 | -0.004 | 0.001 | 0 |
| NOG2 | -0.005 | -0.007 | -0.006 | 0 |
| GLC7 | 0.005 | 0.004 | 0.005 | 0 |
| SWI1 | 0.004 | 0.001 | 0.002 | 0 |
| RSC8 | -0.005 | 0.008 | 0.002 | 0 |
| PRP4 | 0.004 | -0.006 | -0.001 | 0 |
| NBP35 | 0.001 | 0.012 | 0.006 | 0 |
| RFT1 | -0.010 | -0.006 | -0.008 | 0 |
| POP6 | -0.003 | 0.000 | -0.002 | 0 |
| CDS1 | -0.012 | -0.009 | -0.011 | 0 |
| TFG1 | -0.007 | -0.005 | -0.006 | 0 |
| CKS1 | -0.004 | 0.001 | -0.001 | 0 |
| IPI1 | 0.003 | -0.002 | 0.000 | 0 |
| ENP1 | -0.001 | 0.011 | 0.005 | 0 |
| ARC15 | 0.005 | 0.007 | 0.006 | 0 |
| CDC39 | -0.004 | -0.007 | -0.006 | 0 |
| OST1 | 0.006 | 0.002 | 0.004 | 0 |
| SNU23 | -0.003 | 0.003 | 0.000 | 0 |
| SMC3 | -0.008 | -0.011 | -0.010 | 0 |
| GLE1 | 0.006 | 0.011 | 0.008 | 0 |
| LSM8 | -0.004 | -0.004 | -0.004 | 0 |
| STN1 | 0.001 | 0.008 | 0.005 | 0 |
| CDC16 | -0.013 | -0.015 | -0.014 | 0 |
| RPT3 | 0.003 | -0.002 | 0.000 | 0 |
| SCC2 | -0.004 | -0.006 | -0.005 | 0 |
| PRP38 | 0.040 | 0.022 | 0.031 | 1 |
| PRP28 | -0.012 | -0.010 | -0.011 | 0 |
| CCT3 | 0.019 | 0.015 | 0.017 | 0 |
| FRQ1 | -0.005 | -0.004 | -0.004 | 0 |
| CBF5 | -0.016 | -0.006 | -0.011 | 0 |
| TRS31 | -0.005 | -0.006 | -0.006 | 0 |
| RRN9 | -0.009 | 0.004 | -0.003 | 0 |
| SEC3 | 0.013 | 0.020 | 0.016 | 0 |
| HRP1 | -0.005 | -0.017 | -0.011 | 0 |

| | | | | |
|---|---|---|---|---|
| SCC4 | 0.005 | 0.000 | 0.003 | 0 |
| ERG10 | 0.016 | 0.009 | 0.012 | 0 |
| PRE4 | 0.088 | 0.101 | 0.095 | 1 |
| EFB1 | -0.020 | -0.020 | -0.020 | 0 |
| NUP145 | 0.003 | 0.002 | 0.002 | 0 |
| PET9 | 0.001 | -0.005 | -0.002 | 0 |
| TAM41 | -0.005 | -0.007 | -0.006 | 0 |
| CHS2 | 0.015 | 0.006 | 0.010 | 0 |
| HIP1 | -0.038 | 0.060 | 0.011 | 0 |
| IRA1 | -0.017 | -0.021 | -0.019 | 0 |
| BIG1 | 0.003 | 0.016 | 0.010 | 0 |
| DUT1 | 0.000 | 0.001 | 0.000 | 0 |
| RPN2 | -0.003 | -0.002 | -0.002 | 0 |
| MCD1 | 0.002 | 0.002 | 0.002 | 0 |
| CYR1 | 0.013 | -0.005 | 0.004 | 0 |
| NSE4 | -0.006 | -0.005 | -0.006 | 0 |
| ARP4 | -0.022 | -0.028 | -0.025 | -1 |
| NHP2 | -0.020 | 0.001 | -0.010 | 0 |
| SSC1 | -0.010 | 0.002 | -0.004 | 0 |
| RRP1 | -0.014 | 0.001 | -0.006 | 0 |
| TTI1 | 0.020 | 0.019 | 0.019 | 0 |
| NUG1 | -0.007 | -0.011 | -0.009 | 0 |
| CDC1 | -0.002 | -0.013 | -0.008 | 0 |
| ESP1 | -0.006 | -0.022 | -0.014 | 0 |
| CIA1 | 0.001 | -0.001 | 0.000 | 0 |
| MTR4 | 0.004 | -0.005 | -0.001 | 0 |
| ARH1 | -0.024 | -0.011 | -0.017 | 0 |
| CDC123 | -0.015 | -0.009 | -0.012 | 0 |
| SNM1 | -0.010 | -0.003 | -0.007 | 0 |
| SIS1 | -0.007 | -0.003 | -0.005 | 0 |
| PRE1 | 0.070 | 0.061 | 0.066 | 1 |
| DCP1 | 0.006 | 0.010 | 0.008 | 0 |
| SPB4 | 0.012 | 0.024 | 0.018 | 0 |
| NOP4 | 0.005 | 0.008 | 0.007 | 0 |
| RPN12 | 0.065 | 0.070 | 0.068 | 1 |
| POP5 | -0.019 | -0.028 | -0.024 | 0 |
| SRM1 | 0.007 | -0.003 | 0.002 | 0 |
| POL12 | -0.009 | -0.015 | -0.012 | 0 |
| ERG25 | -0.026 | -0.024 | -0.025 | -1 |
| PRP6 | -0.019 | -0.005 | -0.012 | 0 |
| YPP1 | -0.002 | 0.002 | 0.000 | 0 |
| SUP45 | -0.006 | -0.019 | -0.012 | 0 |
| ORC6 | -0.024 | -0.024 | -0.024 | -1 |
| TRS20 | -0.033 | -0.040 | -0.037 | -1 |
| YIL083C | 0.000 | -0.001 | 0.000 | 0 |
| RPT2 | 0.048 | 0.055 | 0.052 | 1 |
| CCT8 | 0.017 | 0.016 | 0.016 | 0 |
| RRP42 | -0.007 | -0.014 | -0.010 | 0 |
| EXO70 | 0.010 | 0.006 | 0.008 | 0 |
| SHR3 | -0.016 | 0.005 | -0.005 | 0 |
| ARP3 | 0.008 | 0.024 | 0.016 | 0 |
| RLI1 | 0.019 | 0.010 | 0.014 | 0 |
| ASK1 | -0.002 | 0.001 | 0.000 | 0 |

| | | | | |
|---|---|---|---|---|
| SRB4 | 0.005 | -0.007 | -0.001 | 0 |
| RVB1 | 0.001 | -0.007 | -0.003 | 0 |
| NAT2 | -0.008 | -0.004 | -0.006 | 0 |
| NSE3 | -0.006 | 0.007 | 0.000 | 0 |
| RRN3 | 0.013 | 0.003 | 0.008 | 0 |
| YRA1 | 0.003 | 0.010 | 0.006 | 0 |
| YSH1 | 0.006 | -0.006 | 0.000 | 0 |
| RIB3 | -0.001 | 0.001 | 0.000 | 0 |
| GCD10 | 0.003 | 0.001 | 0.002 | 0 |
| PRP22 | 0.009 | 0.002 | 0.005 | 0 |
| DBP5 | -0.017 | -0.009 | -0.013 | 0 |
| SEC4 | -0.001 | 0.005 | 0.002 | 0 |
| SPC29 | 0.005 | -0.008 | -0.001 | 0 |
| ERG26 | 0.003 | 0.017 | 0.010 | 0 |
| CDC19 | 0.025 | 0.019 | 0.022 | 0 |
| RPS2 | 0.000 | 0.000 | 0.000 | 0 |
| VHT1 | 0.007 | 0.005 | 0.006 | 0 |
| ALG14 | -0.006 | -0.008 | -0.007 | 0 |
| ZPR1 | 0.002 | 0.004 | 0.003 | 0 |
| RIB7 | 0.004 | 0.003 | 0.004 | 0 |
| DNA2 | -0.012 | -0.006 | -0.009 | 0 |
| RIB5 | -0.328 | -0.337 | -0.332 | -1 |
| SHQ1 | -0.001 | -0.001 | -0.001 | 0 |
| APC11 | -0.011 | -0.013 | -0.012 | 0 |
| NOP9 | 0.003 | 0.002 | 0.002 | 0 |
| YFH1 | -0.005 | -0.006 | -0.006 | 0 |
| TRL1 | -0.016 | -0.005 | -0.010 | 0 |
| CDC13 | -0.005 | -0.013 | -0.009 | 0 |
| YAE1 | -0.026 | 0.000 | -0.013 | 0 |
| PDS1 | 0.007 | 0.006 | 0.006 | 0 |
| FBA1 | -0.010 | 0.002 | -0.004 | 0 |
| RIO1 | -0.008 | -0.005 | -0.006 | 0 |
| MAK5 | -0.012 | -0.013 | -0.012 | 0 |
| APC5 | 0.000 | 0.002 | 0.001 | 0 |
| PWP2 | -0.015 | -0.005 | -0.010 | 0 |
| ALA1 | -0.005 | -0.009 | -0.007 | 0 |
| RPC53 | 0.012 | 0.008 | 0.010 | 0 |
| IDI1 | -0.006 | -0.011 | -0.009 | 0 |
| SEC5 | -0.007 | -0.010 | -0.008 | 0 |
| TIF6 | 0.017 | 0.030 | 0.024 | 0 |
| UBA2 | -0.003 | -0.007 | -0.005 | 0 |
| PIS1 | -0.001 | 0.004 | 0.001 | 0 |
| YGL069C | -0.031 | -0.046 | -0.039 | -1 |
| MRS11 | 0.016 | 0.019 | 0.017 | 0 |
| YGR114C | -0.010 | 0.003 | -0.004 | 0 |
| RET1 | -0.005 | 0.000 | -0.002 | 0 |
| CDC23 | 0.012 | -0.001 | 0.005 | 0 |
| MYO2 | -0.008 | -0.010 | -0.009 | 0 |
| RNR2 | -0.006 | -0.014 | -0.010 | 0 |
| NOG1 | 0.003 | 0.007 | 0.005 | 0 |
| CDC8 | -0.005 | 0.008 | 0.002 | 0 |
| BBP1 | -0.008 | -0.003 | -0.006 | 0 |
| NOC3 | 0.002 | 0.003 | 0.003 | 0 |

| | | | | |
|---|---|---|---|---|
| YTH1 | 0.010 | 0.012 | 0.011 | 0 |
| SGD1 | -0.007 | -0.009 | -0.008 | 0 |
| AOS1 | -0.006 | -0.003 | -0.004 | 0 |
| GPI12 | 0.005 | 0.000 | 0.002 | 0 |
| ALG2 | -0.016 | -0.002 | -0.009 | 0 |
| MRPS18 | 0.000 | 0.015 | 0.008 | 0 |
| RIX7 | 0.012 | -0.002 | 0.005 | 0 |
| HEM15 | -0.003 | 0.000 | -0.001 | 0 |
| TAF3 | 0.001 | -0.008 | -0.003 | 0 |
| PRP46 | 0.003 | -0.005 | -0.001 | 0 |
| PFY1 | 0.001 | 0.000 | 0.000 | 0 |
| SPP381 | 0.003 | 0.000 | 0.001 | 0 |
| SEC63 | 0.013 | 0.016 | 0.015 | 0 |
| RSA4 | -0.006 | -0.002 | -0.004 | 0 |
| RPA43 | -0.010 | -0.004 | -0.007 | 0 |
| YDL152W | 0.000 | 0.007 | 0.004 | 0 |
| TBF1 | -0.001 | 0.011 | 0.005 | 0 |
| YDR187C | 0.000 | -0.005 | -0.003 | 0 |
| HTS1 | 0.008 | 0.005 | 0.006 | 0 |
| RPB7 | 0.017 | 0.003 | 0.010 | 0 |
| SPN1 | 0.022 | 0.041 | 0.031 | 1 |
| YGL074C | -0.008 | 0.010 | 0.001 | 0 |
| LSM3 | -0.014 | -0.010 | -0.012 | 0 |
| CBF2 | -0.006 | 0.006 | 0.000 | 0 |
| RFC1 | -0.001 | -0.005 | -0.003 | 0 |
| KOG1 | 0.009 | 0.009 | 0.009 | 0 |
| SCD5 | -0.043 | -0.011 | -0.027 | 0 |
| YJL032W | 0.038 | -0.007 | 0.015 | 0 |
| SEC62 | 0.100 | 0.093 | 0.096 | 1 |
| URB1 | -0.005 | -0.007 | -0.006 | 0 |
| DIM1 | -0.006 | -0.005 | -0.005 | 0 |
| YLR076C | -0.020 | -0.027 | -0.024 | -1 |
| RPN7 | 0.074 | 0.076 | 0.075 | 1 |
| YLR339C | 0.006 | 0.010 | 0.008 | 0 |
| RPC82 | 0.006 | 0.004 | 0.005 | 0 |
| BDP1 | -0.008 | 0.004 | -0.002 | 0 |
| MPS2 | 0.004 | -0.008 | -0.002 | 0 |
| DBP6 | -0.003 | 0.005 | 0.001 | 0 |
| YCS4 | -0.009 | -0.004 | -0.007 | 0 |
| NOC2 | 0.000 | 0.003 | 0.002 | 0 |
| GPI2 | 0.011 | 0.011 | 0.011 | 0 |
| CET1 | 0.018 | 0.010 | 0.014 | 0 |
| SPP2 | 0.002 | -0.003 | -0.001 | 0 |
| CNS1 | -0.001 | -0.003 | -0.002 | 0 |
| TRE2 | -0.003 | -0.003 | -0.003 | 0 |
| ATP16 | -0.014 | -0.017 | -0.016 | 0 |
| SOG2 | -0.014 | -0.011 | -0.012 | 0 |
| YDL163W | 0.038 | 0.029 | 0.034 | 1 |
| RPL33A | 0.012 | 0.021 | 0.016 | 0 |
| SLY1 | -0.024 | -0.027 | -0.026 | -1 |
| TAH18 | 0.004 | -0.005 | -0.001 | 0 |
| RRP17 | -0.004 | -0.013 | -0.009 | 0 |
| RRP9 | 0.010 | 0.003 | 0.006 | 0 |

| | | | | |
|---|---|---|---|---|
| CDC20 | -0.016 | 0.004 | -0.006 | 0 |
| ENP2 | -0.001 | 0.002 | 0.000 | 0 |
| CFD1 | -0.004 | 0.003 | 0.000 | 0 |
| TAD2 | 0.001 | -0.005 | -0.002 | 0 |
| TFA1 | -0.011 | -0.006 | -0.009 | 0 |
| CDC45 | -0.013 | -0.002 | -0.007 | 0 |
| RPC40 | -0.013 | -0.007 | -0.010 | 0 |
| YLR379W | 0.002 | -0.004 | -0.001 | 0 |
| ISD11 | -0.003 | -0.005 | -0.004 | 0 |
| TOP2 | 0.003 | 0.002 | 0.002 | 0 |
| USE1 | 0.014 | 0.006 | 0.010 | 0 |
| ESF2 | -0.008 | 0.008 | 0.000 | 0 |
| YHC1 | -0.011 | -0.021 | -0.016 | 0 |
| YOR218C | 0.020 | 0.013 | 0.016 | 0 |
| NSL1 | -0.006 | 0.003 | -0.001 | 0 |
| YPL238C | 0.046 | 0.056 | 0.051 | 1 |
| SMP3 | -0.006 | -0.010 | -0.008 | 0 |
| YBR190W | -0.009 | -0.007 | -0.008 | 0 |
| RPN8 | 0.005 | 0.008 | 0.007 | 0 |
| PRP9 | 0.002 | 0.000 | 0.001 | 0 |
| MRS6 | -0.010 | -0.012 | -0.011 | 0 |
| CDC9 | -0.023 | -0.007 | -0.015 | 0 |
| NOP53 | -0.008 | -0.002 | -0.005 | 0 |
| TCP1 | 0.030 | 0.024 | 0.027 | 1 |
| TFB4 | -0.006 | -0.007 | -0.006 | 0 |
| GPI19 | -0.005 | -0.001 | -0.003 | 0 |
| RRP15 | 0.016 | 0.015 | 0.016 | 0 |
| GPI10 | 0.000 | 0.011 | 0.005 | 0 |
| TYS1 | 0.000 | -0.004 | -0.002 | 0 |
| YRB2 | -0.001 | -0.004 | -0.003 | 0 |
| GWT1 | -0.010 | -0.013 | -0.011 | 0 |
| UGP1 | 0.006 | 0.003 | 0.004 | 0 |
| MDN1 | -0.006 | 0.004 | -0.001 | 0 |
| SEN1 | 0.007 | 0.000 | 0.003 | 0 |
| SPT6 | 0.006 | -0.012 | -0.003 | 0 |
| RCL1 | -0.001 | 0.006 | 0.002 | 0 |
| PRP39 | -0.010 | -0.007 | -0.008 | 0 |
| RPB8 | -0.067 | -0.073 | -0.070 | -1 |
| SUA7 | -0.019 | -0.011 | -0.015 | 0 |
| CCL1 | 0.004 | 0.012 | 0.008 | 0 |
| PUP1 | 0.065 | 0.067 | 0.066 | 1 |
| RIM2 | 0.000 | 0.006 | 0.003 | 0 |
| YTM1 | -0.014 | -0.004 | -0.009 | 0 |
| DBP10 | 0.002 | -0.002 | 0.000 | 0 |
| RRP12 | 0.002 | -0.008 | -0.003 | 0 |
| CWC2 | -0.007 | 0.002 | -0.002 | 0 |
| CDC60 | 0.005 | 0.003 | 0.004 | 0 |
| PCF11 | 0.010 | 0.005 | 0.007 | 0 |
| DIB1 | 0.016 | 0.025 | 0.021 | 0 |
| PRP3 | -0.001 | 0.012 | 0.005 | 0 |
| NOC4 | 0.027 | 0.024 | 0.026 | 1 |
| TIP20 | -0.009 | -0.004 | -0.006 | 0 |
| YGR190C | -0.003 | -0.012 | -0.007 | 0 |

| | | | | |
|---|---|---|---|---|
| SEC6 | -0.033 | -0.048 | -0.041 | -1 |
| UTP10 | -0.014 | -0.003 | -0.009 | 0 |
| PRI2 | 0.001 | -0.005 | -0.002 | 0 |
| DIP2 | -0.009 | -0.003 | -0.006 | 0 |
| YLR458W | -0.005 | -0.001 | -0.003 | 0 |
| YNL114C | -0.058 | -0.035 | -0.046 | -1 |
| SDA1 | -0.002 | -0.005 | -0.003 | 0 |
| MIM1 | 0.006 | 0.015 | 0.011 | 0 |
| RSE1 | 0.001 | 0.009 | 0.005 | 0 |
| DFR1 | -0.009 | -0.005 | -0.007 | 0 |
| PRP45 | -0.005 | 0.000 | -0.002 | 0 |
| GLN1 | 0.006 | 0.007 | 0.007 | 0 |
| SME1 | 0.036 | 0.038 | 0.037 | 1 |
| ABD1 | -0.006 | 0.003 | -0.002 | 0 |
| HEM4 | -0.006 | 0.001 | -0.002 | 0 |
| PRP11 | -0.005 | -0.012 | -0.009 | 0 |
| ULP1 | 0.006 | -0.008 | -0.001 | 0 |
| TIM22 | -0.005 | -0.004 | -0.004 | 0 |
| IPL1 | 0.000 | -0.006 | -0.003 | 0 |
| TRS23 | 0.000 | -0.001 | 0.000 | 0 |
| ASA1 | -0.008 | -0.004 | -0.006 | 0 |
| YDR526C | 0.036 | -0.038 | -0.001 | 0 |
| NUT2 | 0.011 | 0.003 | 0.007 | 0 |
| SUA5 | 0.018 | 0.000 | 0.009 | 0 |
| YGR251W | -0.005 | 0.007 | 0.001 | 0 |
| RHO3 | 0.004 | 0.008 | 0.006 | 0 |
| GCD14 | -0.005 | 0.004 | -0.001 | 0 |
| CSE4 | -0.015 | -0.009 | -0.012 | 0 |
| YLR140W | -0.028 | 0.006 | -0.011 | 0 |
| RPM2 | 0.005 | -0.001 | 0.002 | 0 |
| MAK16 | -0.020 | -0.019 | -0.020 | 0 |
| KRE33 | -0.004 | 0.002 | -0.001 | 0 |
| MET30 | 0.004 | -0.002 | 0.001 | 0 |
| YOL134C | 0.025 | -0.044 | -0.009 | 0 |
| TAF8 | 0.003 | 0.011 | 0.007 | 0 |
| ESA1 | 0.010 | 0.010 | 0.010 | 0 |
| MCM2 | -0.010 | -0.008 | -0.009 | 0 |
| FHL1 | 0.001 | 0.006 | 0.003 | 0 |
| MTR10 | -0.006 | -0.009 | -0.008 | 0 |
| PRP5 | -0.007 | 0.002 | -0.002 | 0 |
| YOR287C | 0.001 | 0.001 | 0.001 | 0 |
| PSA1 | 0.008 | 0.026 | 0.017 | 0 |
| TIM50 | -0.017 | 0.018 | 0.000 | 0 |
| PSF1 | -0.005 | -0.016 | -0.010 | 0 |
| SRP72 | 0.005 | 0.010 | 0.007 | 0 |
| SRP101 | 0.003 | 0.006 | 0.004 | 0 |
| SRP54 | 0.000 | 0.008 | 0.004 | 0 |
| UTR5 | -0.003 | 0.007 | 0.002 | 0 |
| JIP5 | -0.005 | -0.006 | -0.005 | 0 |
| YGL239C | 0.001 | 0.002 | 0.002 | 0 |
| YGR265W | -0.010 | -0.001 | -0.006 | 0 |
| TID3 | 0.014 | 0.014 | 0.014 | 0 |
| CDC6 | 0.002 | 0.000 | 0.001 | 0 |

| | | | | |
|---|---|---|---|---|
| YKL083W | -0.028 | -0.002 | -0.015 | 0 |
| ACS2 | 0.005 | -0.006 | -0.001 | 0 |
| UTP14 | -0.009 | -0.008 | -0.008 | 0 |
| USO1 | -0.004 | -0.012 | -0.008 | 0 |
| NAM9 | -0.002 | 0.011 | 0.005 | 0 |
| THS1 | -0.003 | -0.006 | -0.004 | 0 |
| HSP10 | 0.009 | 0.004 | 0.006 | 0 |
| VTI1 | -0.004 | 0.003 | 0.000 | 0 |
| CDC31 | -0.007 | -0.004 | -0.005 | 0 |
| YBL073W | 0.000 | -0.002 | -0.001 | 0 |
| MRD1 | 0.008 | 0.005 | 0.007 | 0 |
| MED4 | 0.000 | 0.004 | 0.002 | 0 |
| POP4 | -0.004 | -0.004 | -0.004 | 0 |
| RRS1 | 0.016 | 0.017 | 0.017 | 0 |
| LUC7 | -0.011 | -0.006 | -0.009 | 0 |
| FAL1 | -0.001 | -0.018 | -0.009 | 0 |
| SAR1 | -0.018 | -0.025 | -0.022 | 0 |
| TFB1 | -0.016 | -0.001 | -0.008 | 0 |
| PRE2 | 0.091 | 0.095 | 0.093 | 1 |
| BUR6 | -0.005 | 0.011 | 0.003 | 0 |
| DPB2 | -0.008 | -0.013 | -0.010 | 0 |
| GUS1 | 0.007 | -0.008 | 0.000 | 0 |
| CWC22 | -0.004 | -0.003 | -0.003 | 0 |
| PAN1 | 0.000 | 0.020 | 0.010 | 0 |
| YJL195C | 0.000 | 0.001 | 0.001 | 0 |
| HYM1 | -0.009 | -0.005 | -0.007 | 0 |
| YLR198C | -0.023 | 0.034 | 0.006 | 0 |
| CDC5 | -0.015 | 0.002 | -0.007 | 0 |
| RSC3 | -0.001 | -0.001 | -0.001 | 0 |
| NET1 | -0.003 | -0.012 | -0.007 | 0 |
| CDC21 | 0.017 | -0.004 | 0.006 | 0 |
| TAF7 | 0.005 | 0.020 | 0.012 | 0 |
| YOR262W | 0.016 | 0.004 | 0.010 | 0 |
| AAR2 | -0.003 | -0.004 | -0.003 | 0 |
| YPR142C | 0.002 | 0.004 | 0.003 | 0 |
| LAS17 | -0.003 | 0.001 | -0.001 | 0 |
| YCL041C | -0.002 | -0.009 | -0.006 | 0 |
| NOP58 | -0.006 | -0.010 | -0.008 | 0 |
| POL3 | 0.002 | -0.011 | -0.004 | 0 |
| SEN54 | 0.000 | 0.006 | 0.003 | 0 |
| RSM10 | 0.005 | -0.004 | 0.001 | 0 |
| FAS2 | 0.014 | 0.010 | 0.012 | 0 |
| UTP4 | -0.011 | -0.001 | -0.006 | 0 |
| COG4 | 0.009 | 0.000 | 0.005 | 0 |
| TUB2 | 0.013 | 0.023 | 0.018 | 0 |
| BET2 | 0.017 | 0.008 | 0.012 | 0 |
| UFD1 | -0.003 | -0.005 | -0.004 | 0 |
| ERG11 | -0.008 | -0.001 | -0.004 | 0 |
| RPR2 | -0.018 | 0.002 | -0.008 | 0 |
| YJR012C | 0.009 | -0.001 | 0.004 | 0 |
| TRZ1 | 0.009 | -0.001 | 0.004 | 0 |
| YLR230W | 0.006 | 0.001 | 0.004 | 0 |
| SEN15 | -0.012 | -0.006 | -0.009 | 0 |

| | | | | |
|---|---|---|---|---|
| POL5 | -0.009 | -0.014 | -0.011 | 0 |
| CBK1 | -0.011 | -0.006 | -0.008 | 0 |
| NNF1 | -0.008 | 0.005 | -0.002 | 0 |
| UFE1 | 0.013 | 0.002 | 0.007 | 0 |
| FCP1 | 0.002 | 0.000 | 0.001 | 0 |
| KRE5 | 0.007 | 0.007 | 0.007 | 0 |
| ORC2 | -0.002 | -0.008 | -0.005 | 0 |
| DPM1 | -0.003 | 0.004 | 0.000 | 0 |
| DED1 | 0.004 | 0.001 | 0.002 | 0 |
| PBN1 | -0.002 | -0.015 | -0.009 | 0 |
| HSH49 | 0.005 | -0.002 | 0.002 | 0 |
| KIN28 | 0.003 | 0.000 | 0.001 | 0 |
| SEC16 | 0.003 | 0.009 | 0.006 | 0 |
| HEM12 | 0.031 | 0.009 | 0.020 | 0 |
| SRP68 | -0.003 | 0.003 | 0.000 | 0 |
| YDR355C | -0.024 | -0.037 | -0.031 | -1 |
| ECO1 | -0.002 | 0.013 | 0.006 | 0 |
| YGR073C | 0.014 | 0.005 | 0.010 | 0 |
| BRL1 | 0.004 | 0.002 | 0.003 | 0 |
| YJL009W | -0.041 | -0.050 | -0.045 | -1 |
| GPI14 | -0.001 | -0.021 | -0.011 | 0 |
| SOF1 | 0.018 | 0.009 | 0.014 | 0 |
| HSP60 | -0.007 | -0.029 | -0.018 | 0 |
| PDS5 | -0.003 | -0.001 | -0.002 | 0 |
| RPN3 | 0.037 | 0.025 | 0.031 | 1 |
| RIA1 | -0.011 | -0.005 | -0.008 | 0 |
| SPC42 | -0.006 | -0.001 | -0.003 | 0 |
| NUP1 | 0.003 | -0.001 | 0.001 | 0 |
| SAM50 | -0.016 | -0.004 | -0.010 | 0 |
| PRT1 | 0.003 | 0.008 | 0.005 | 0 |
| YBR089W | -0.040 | -0.049 | -0.045 | -1 |
| PZF1 | 0.006 | 0.001 | 0.004 | 0 |
| PGK1 | -0.008 | -0.016 | -0.012 | 0 |
| SCM3 | -0.001 | -0.004 | -0.003 | 0 |
| YDR053W | -0.020 | 0.025 | 0.002 | 0 |
| SPC110 | 0.000 | -0.005 | -0.003 | 0 |
| CDC14 | -0.008 | 0.002 | -0.003 | 0 |
| VAS1 | 0.022 | 0.010 | 0.016 | 0 |
| RPP1 | 0.019 | 0.015 | 0.017 | 0 |
| YJL015C | 0.009 | 0.016 | 0.012 | 0 |
| ESS1 | -0.001 | 0.001 | 0.000 | 0 |
| GRC3 | -0.007 | -0.010 | -0.009 | 0 |
| DBP9 | -0.002 | 0.000 | -0.001 | 0 |
| FOL3 | 0.005 | 0.009 | 0.007 | 0 |
| CCA1 | -0.002 | -0.019 | -0.011 | 0 |
| PIK1 | -0.023 | -0.013 | -0.018 | 0 |
| MPE1 | -0.012 | 0.010 | -0.001 | 0 |
| RPO31 | -0.008 | -0.013 | -0.011 | 0 |
| NOB1 | 0.007 | 0.009 | 0.008 | 0 |
| RET3 | -0.018 | -0.046 | -0.032 | 0 |
| MTW1 | 0.002 | 0.014 | 0.008 | 0 |
| RRP43 | -0.014 | -0.009 | -0.012 | 0 |
| RPN5 | 0.052 | 0.054 | 0.053 | 1 |

| | | | | |
|---|---|---|---|---|
| MAK21 | 0.001 | -0.002 | -0.001 | 0 |
| BCP1 | -0.002 | -0.003 | -0.002 | 0 |
| ALG13 | 0.001 | 0.002 | 0.001 | 0 |
| DAM1 | -0.006 | -0.009 | -0.008 | 0 |
| RRP3 | 0.012 | 0.012 | 0.012 | 0 |
| YJL018W | 0.017 | 0.017 | 0.017 | 0 |
| TAH11 | -0.012 | -0.012 | -0.012 | 0 |
| YLL037W | -0.014 | -0.026 | -0.020 | 0 |
| YLR317W | -0.013 | 0.017 | 0.002 | 0 |
| TRS130 | 0.006 | 0.010 | 0.008 | 0 |
| SMC2 | 0.005 | -0.002 | 0.001 | 0 |
| SEC2 | -0.008 | 0.003 | -0.002 | 0 |
| RSC4 | -0.013 | 0.004 | -0.005 | 0 |
| GLN4 | 0.014 | 0.000 | 0.007 | 0 |
| PLP2 | 0.000 | 0.016 | 0.008 | 0 |
| TFB2 | -0.008 | 0.009 | 0.000 | 0 |
| YBR124W | -0.005 | -0.041 | -0.023 | 0 |
