## Supplemental File 1 for "Chaperone-Mediated Reflux of Secretory Proteins to the Cytosol During Endoplasmic Reticulum Stress"

### SUPPLEMENTARY MATERIALS AND METHODS

#### Immunofluorescence:

Cells were grown to mid-log phase, collected by centrifugation at 3000rpm, then resuspended in 1mL of 4% paraformaldehyde solution for 30 minutes at room temperature. Cells were then centrifuged again and the pellet washed twice with 5mL of KS buffer (0.1 M KPO4/1.2 M sorbitol) before resuspending again in KS buffer. After fixation, spheroplasts were generated using 20T zymolyase and resuspended again in KS buffer. Slides were washed with cold acetone and then treated with 0.1%-polylysine before adding spheroplasts; slides were then washed and blocked with PBS-BSA for 30 minutes. Slides were incubated with the primary antibody overnight, then washed three times with PBS-BSA. Secondary antibodies were prepared in PBS-BSA and slides were incubated with the secondary antibodies for 2hrs before washing again with PBS-BSA. Slides were then mounted and sealed with nail polish.

#### Quantitative Real-Time PCR

RNA was isolated from whole cells using Qiagen RNeasy kit. 1μg total RNA was reverse transcribed using the QuantiTect Reverse Transcription Kit (Qiagen) and amplified using the StepOnePlus Real-Time PCR System (Applied Biosystems) with SYBR green. Thermal cycles were: 5 min at 95 °C, 40 cycles of 15 s at 95 °C, 30 s at 60 °C. Gene expression levels were normalized to PGK1. Primers used for Q-PCR were as follows: GFP: 5'-acaagcagaagaacggcatc-3' and 5'-gcaggtgctcaggtagtggt-3'; PGK1: 5'-ctcactcttctatggtcggtttc-3' and 5'-gagaacccaaccagaccatt-3'.

#### Subcellular Fractionation

Yeast organelles were fractionated using ultracentrifugation according to the protocol described in [1].

**Estimating the fraction of newly synthesized eroGFP-Glyc**

The time evolution of the eroGFP translation rate (m) and the concentration of eroGFP protein (P) were estimated from a two-step transcription-translation model:

$$\frac{dm}{dt} = -\frac{m}{T_M}$$

$$\frac{dP}{dt} = -\frac{\ln(2)}{T_P}(m - P)$$

m and P are normalized to their steady state levels in galactose conditions and are thus unitless variables. $\tau_M$, defined as the half-life of the eroGFP translation rate, was estimated from the pulse-label data. $\ln(2)/\tau_P$ is the half-life of eroGFP proteins that decay due to dilution from cell division. $\tau_P$ is set to 120 min, which corresponds to the doubling time of yeast in SD media.

**Mass Spectrometry Analysis**

Immunoprecipitated FLAG-eroGFP-Glyc was purified by gel electrophoresis and subjected to in-gel trypsin digestion. Gel bands were diced and washed three times with 25 mM $NH_4HCO_3$/50% ACN, and evaporated to dryness. Cysteines were reduced by incubation with 10 mM DTT in 25 mM $NH_4HCO_3$ at 56°C for 1 hour, and free sulfhydryl groups were alkylated by incubation with 55 mM iodoacetamide for 45 minutes at room temperature. Gel pieces were washed twice with 25 mM $NH_4HCO_3$/50% ACN and evaporated to dryness. Gel pieces were rehydrated with 12.5 ng/µL trypsin in 25 mM

$NH_4HCO_3$ and incubated at 37°C overnight. The supernatant was removed and combined with two extractions with 50% ACN/5% formic acid. The combined supernatant was subjected to concentration on C18 ZipTips (Millipore) according to the manufacturer's specifications. Following evaporation, samples were resuspended in 0.1% formic acid for liquid chromatography and mass spectrometry (LC-MS/MS) analysis.

Digested peptide mixtures were analyzed in technical duplicate on a Thermo Scientific LTQ Orbitrap Elite mass spectrometry system equipped with a Proxeon Easy nLC 1000 ultra high-pressure liquid chromatography and autosampler system. Samples were injected onto a C18 column (25 cm x 75 um I.D. packed with ReproSil Pur C18 AQ 1.9 um particles) in 0.1% formic acid and then subjected to a 2-hours gradient from 0.1% formic acid to 30% ACN/0.1% formic acid. The mass spectrometer collected data in a data-dependent fashion, collecting one full scan in the Orbitrap at 120,000 resolution followed by 20 collision-induced dissociation MS/MS scans in the dual linear ion trap for the 20 most intense peaks from the full scan. Dynamic exclusion was enabled for 30 seconds with a repeat count of 1. Charge state screening was employed to reject analysis of singly charged species or species for which a charge could not be assigned. Extracted ion chromatograms for peptides of interest were generated using the Thermo Scientific XCalibur QualBrowser software package.

#### SUPPLEMENTARY FIGURE LEGENDS

**Figure S1.** ER-targeted eroGFP re-localizes to the cytosol during ER stress

(A) Time course of eroGFP ratio changes in WT cells upon exposure to DTT (blue) or Tm (red).

(B) eroGFP redox state in WT cells treated with Tm (6µg/mL) or DTT (1mM) for the indicated time points. Extracts were treated with NEM, resolved on non-reducing SDS-PAGE, and immunoblotted against GFP.

(C) Quantification of reduced eroGFP(red) percentage in WT cells treated with either Tm or DTT, ratios calculated from a non-reducing SDS-PAGE after alkylating the protein lysates with NEM for 30 minutes.

(D) Confocal images of wild-type cells expressing eroGFP and cytosolic tdTomato treated with Tm (6µg/mL) for the indicated time points.

**Figure S2.** Pre-existing eroGFP-Glyc is refluxed from the ER during ER stress.

(A) Schematic of eroGFP-Glyc. An *N*-linked protein glycosylation recognition sequence was inserted into the 9 amino acid linker between eroGFP and the HDEL retrieval sequence.

(B) Immunoblot (anti-GFP) of protein extracts from wild-type cells expressing eroGFP and eroGFP-Glyc.

(C) Schematic of FLAG-tagged GAL1-eroGFP-Glyc (top) and qPCR to determine relative eroGFP-Glyc mRNA levels. Cells were grown on Galactose for 4hrs, then shifted to glucose; time 0 indicates glucose addition.

(D) Pulse-label for wild-type cells expressing FLAG-tagged GAL1-eroGFPGlyc treated according to (B).

(E) Half-life of the FLAG-tagged eroGFP-Glyc translation rate (ꞇM = 82min) is calculated by fitting an exponential decay curve, e(-t/ꞇM), to the pulse-label data. The solid line represents the curve fit and the dashed lines represent 99% confidence bounds.

(F) The fraction of newly synthesized FLAG-tagged eroGFP-Glyc is estimated from a two-step transcription/translation model (see Methods).

(G) Confocal microscopy for eroGFP-Glyc. Cells were treated with galactose for 4hrs and then shifted to glucose containing media. After 2hrs on glucose, Tm was added for an additional 2 hrs.

(H) Two-dimensional gel electrophoresis followed by western blot of GAL1-eroGFP-Glyc for the indicated condition. FLAG-tagged eroGFP-ND indicates the GAL1-eroGFP-Glyc N to D point mutant. Duplicated blots are shown in the vertical and horizontal directions to aid visualization/alignment in both dimensions.

(I) Ratios of aspartate to asparagine at the position marked by * for cytosolic roGFP-Glyc (croGFP) and eroGFP-Glyc treated with Tm or PNGase where indicated.

**Figure S3.** Experimental Design and Analysis for the eroGFP screen.

(A) Schematic of mating strategy used to generate the eroGFP expressing gene libraries using the Synthetic Genetic Array strategy (Tong et al, 2001).

(B and C) Histograms of the differences between eroGFP replicate measurements for untreated and tunicamycin treated samples respectively

(D and E) Fit of the histograms from B and C modeled as the sum of two Gaussian distributions according to [2]. The fitted lines are in orange overlaid over the histograms from B and C.

(F and G) Histograms of the mean data for untreated and tunicamycin treated samples respectively in green overlaid over the histograms described in D-E.

(H) Enrichment of Gene Ontology terms (P<0.001) for the indicated ER functions.

**Figure S4.** ER protein reflux is not reliant on canonical ERAD machinery.

(A) Immunoprecipitation (IP) using anti-GFP antibody of extracts from wild-type cells treated with tunicamycin (or untreated) in the (presence/absence) of MG132, followed by immunoblot analysis with anti-ubiquitin (Ub) and anti-GFP antibodies.
(B) Immunoprecipitation (IP) using anti-HA antibody of extracts from wild-type cells expressing HA-tagged-CPY* treated with tunicamycin (or untreated) in the (presence/absence) of MG132, followed by immunoblot analysis with anti-ubiquitin (Ub) and anti-HA antibodies.

*Heavy chain

**Figure S5.** Reflux of ER proteins requires HLJ1

(A) ER targeted yemEos3.2 was first converted by UV in *hlj1*Δ cells and then cells were treated with Tm (6μg/mL). Images (550nm) were taken exactly after Tm addition and 120 minutes after treatment.

(B) Quantification of *hlj1*Δ images with ER- targeted mEos3.2.

(C) Confocal images for *glr1*Δ treated with Tm (6μg/mL) for 2hrs.

(D) Cyto-roGFP redox state in WT, *hlj1*Δ and *glr1*Δ cells treated with (6μg/mL) Tm. Protein extracts were treated with NEM, resolved on non-reducing SDS-PAGE, and immunoblotted against GFP.

(E) Immunoblot (anti-FLAG) of protein extracts from *hlj1*Δ cells overexpressing FLAG tagged HLJ1 after induction with galactose for the indicated time points.

(F) Confocal images for *hlj1*Δ cells overexpressing WT-HLJ1 and expressing the ER-targeted mEos3.2 after shifting to Galactose containing media for the indicated time points.

(G) Subcellular protein fractionation of Myc-CPR5, PDI1 and eroGFP in WT, and *hlj1*Δ cells treated with Tm.

(H) Subcellular protein fractionation of CPR5, PDI1, and eroGFP in the ERAD double mutant (*hrd1*Δ *doa10*Δ) expressing either CPY or CPY* under the conditional promoter CUP1 (-/+) 200µM copper sulfate (CuSO4).

(I) Viability assay after Tm challenge for 4 hours in WT, *hlj1*Δ and *sse1*Δ cells.

(J) Plate sensitivity assay of WT, *hlj1*Δ and *sse1*Δ cells on Tm containing agar plates.

**Movie S1:** ER-targeted mEos3.2 in untreated WT cells followed for 120 minutes after photoconversion by a UV pulse.

**Movie S2:** ER-targeted mEos3.2 in Tm-treated WT cells followed for 120 minutes after photoconversion by a UV pulse.

**Movie S3:** ER-targeted mEos3.2 in Tm-treated WT cells followed for 8hours after photoconversion by a UV pulse.

**Movie S4:** ER-targeted mEos3.2 in Tm-treated WT cells followed for 120 minutes after photoconversion by a UV pulse.

**Movie S5:** ER-targeted mEos3.2 in Tm-treated *hrd1*Δ*doa10*Δ cells followed for 120 minutes after photoconversion by a UV pulse.

**Movie S6:** ER-targeted mEos3.2 in *hrd1*Δ*doa10*Δ cells expressing CPY* after addition of copper.

1. Rieder, S.E. and S.D. Emr, *Isolation of subcellular fractions from the yeast Saccharomyces cerevisiae.* Curr Protoc Cell Biol, 2001. **Chapter 3**: p. Unit 3 8.
2. Breslow, D.K., et al., *A comprehensive strategy enabling high-resolution functional analysis of the yeast genome.* Nat Methods, 2008. **5**(8): p. 711-8.
