## Supplementary figures and images for "Chaperone-Mediated Reflux of Secretory Proteins to the Cytosol During Endoplasmic Reticulum Stress"

### Supplemental Figure 1

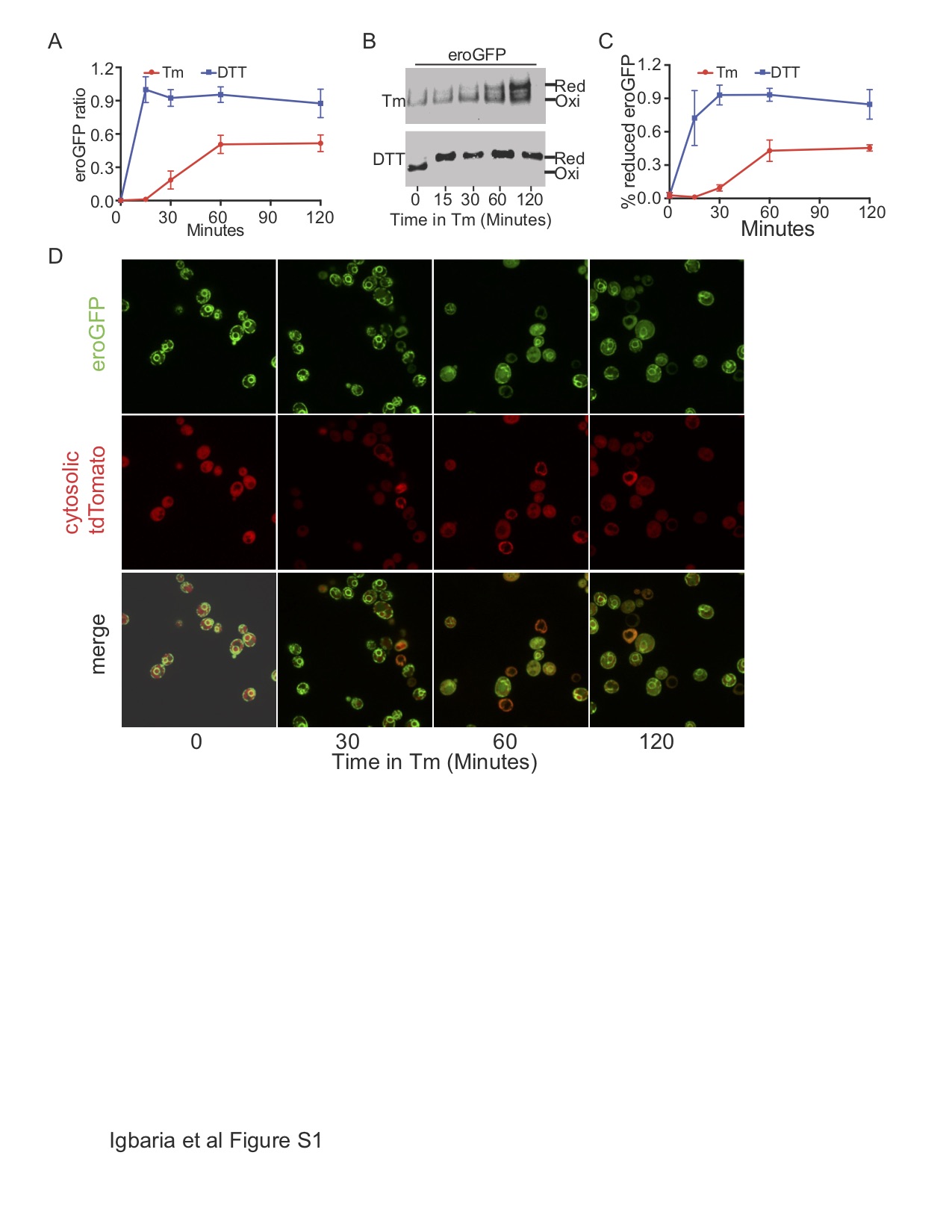

### Supplemental Figure 2

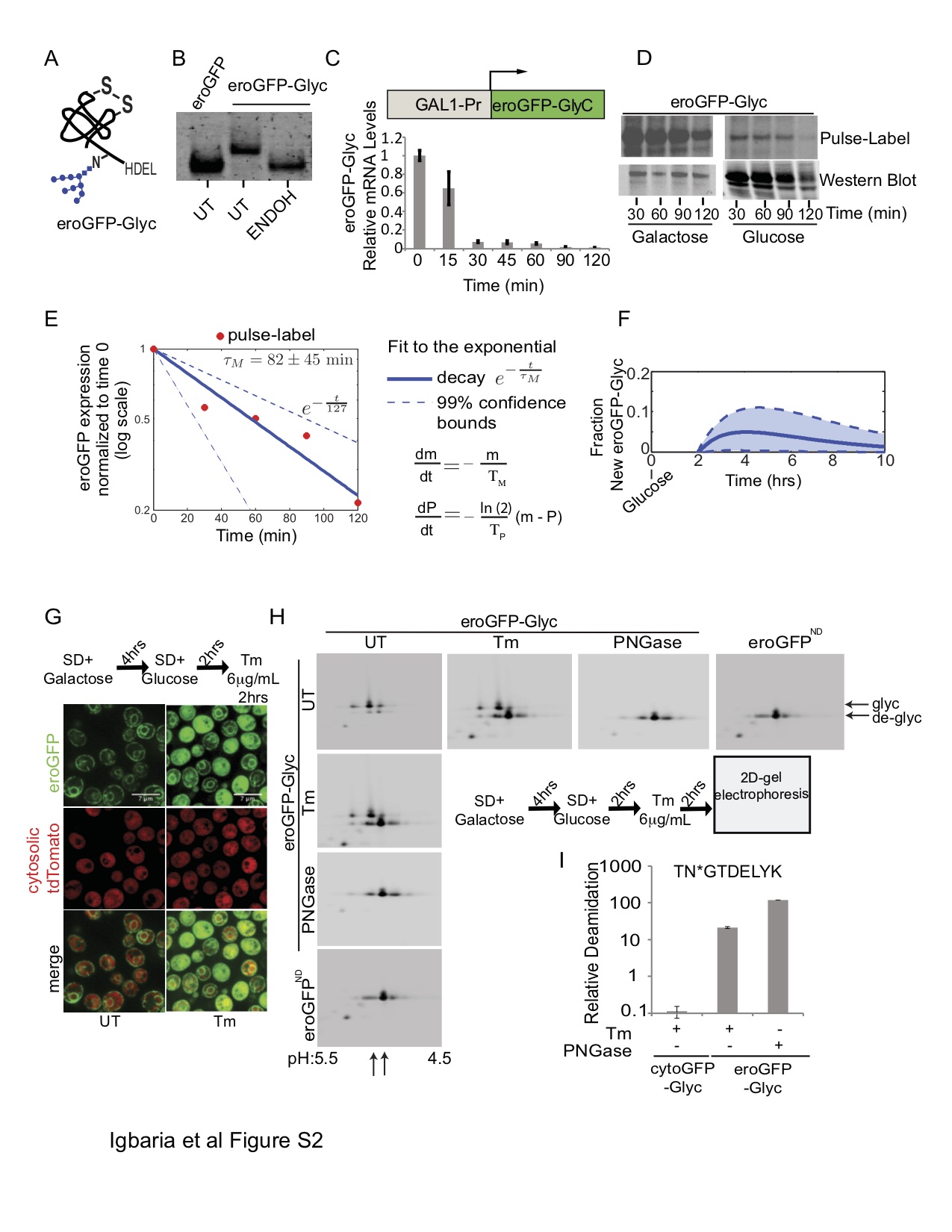

### Supplemental Figure 3

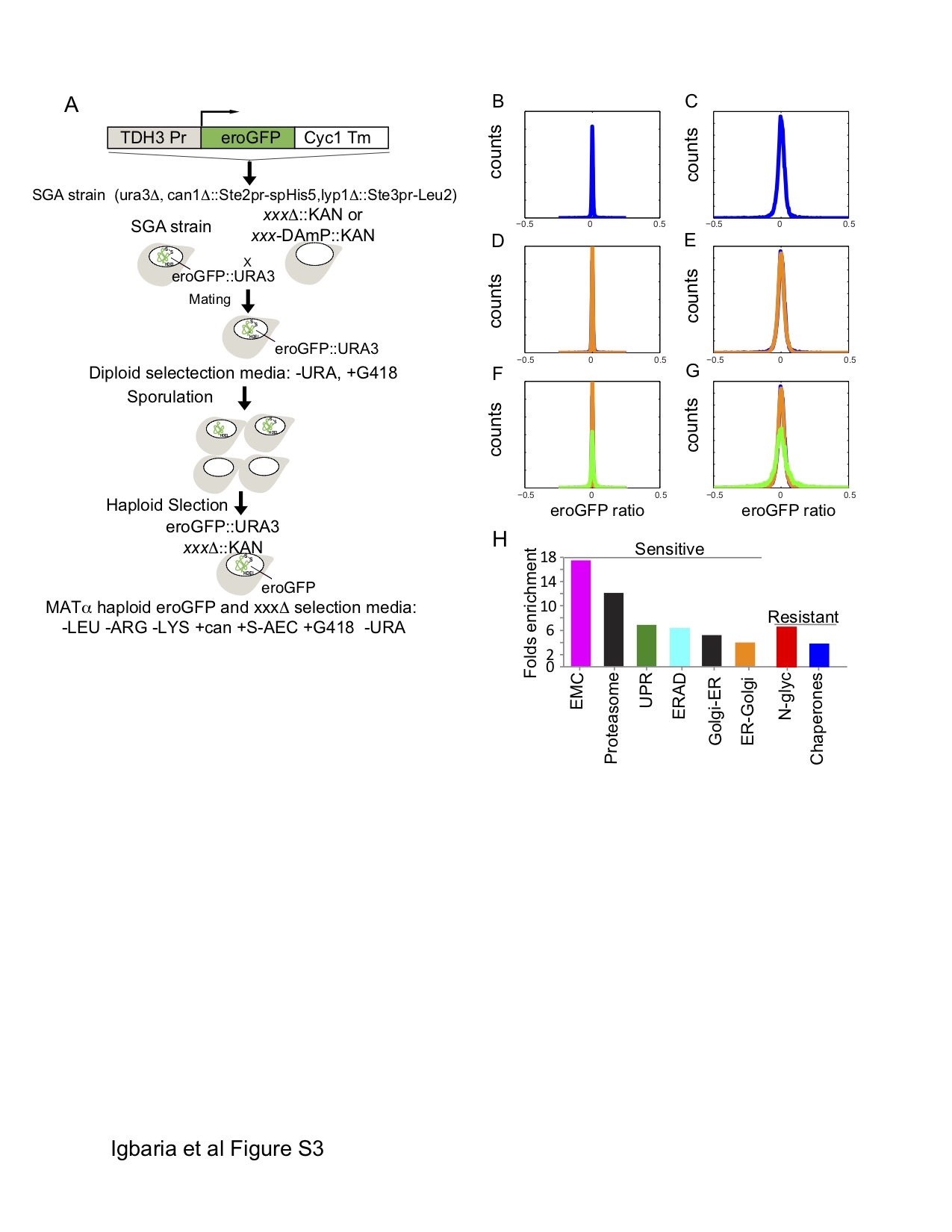

### Supplemental Figure 4

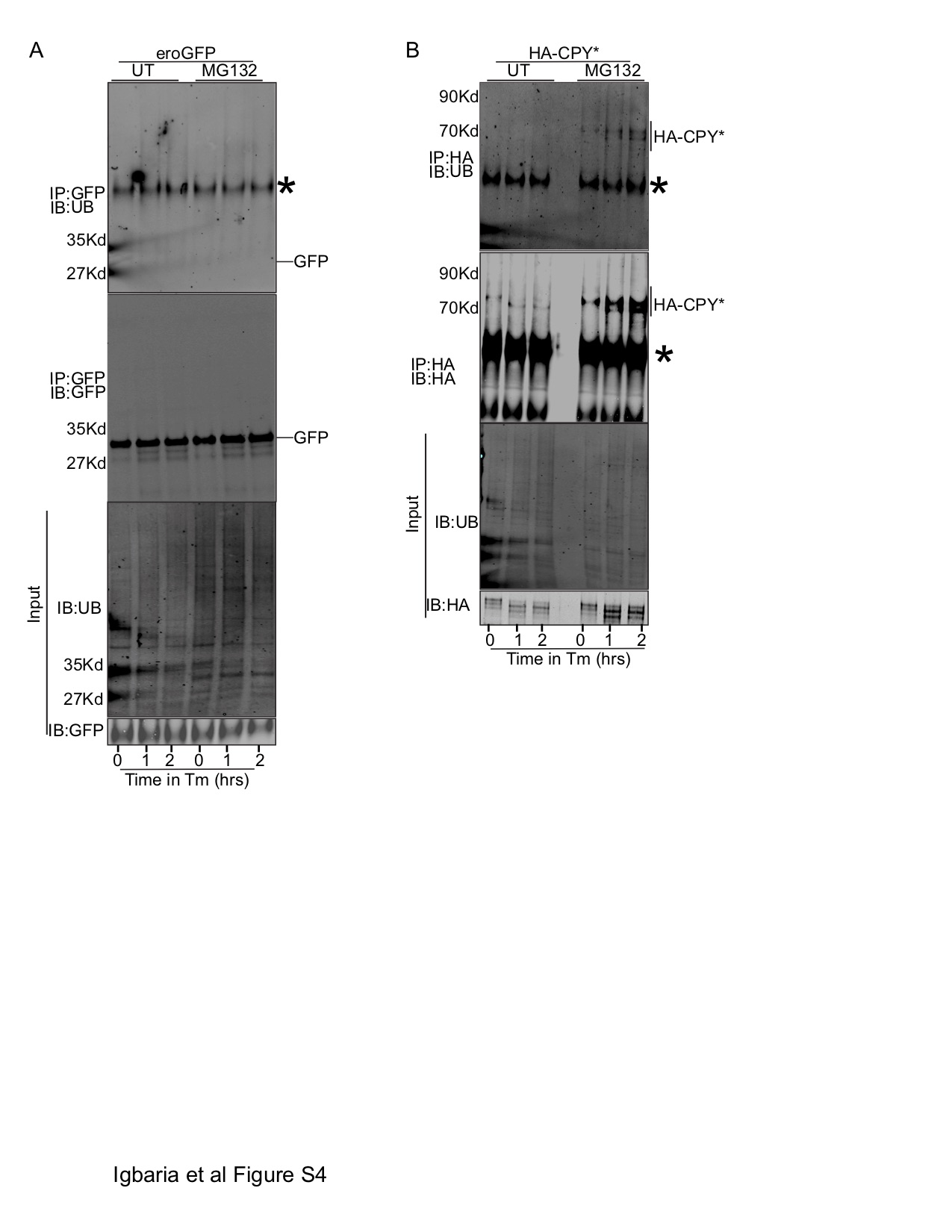

### Supplemental Figure 5

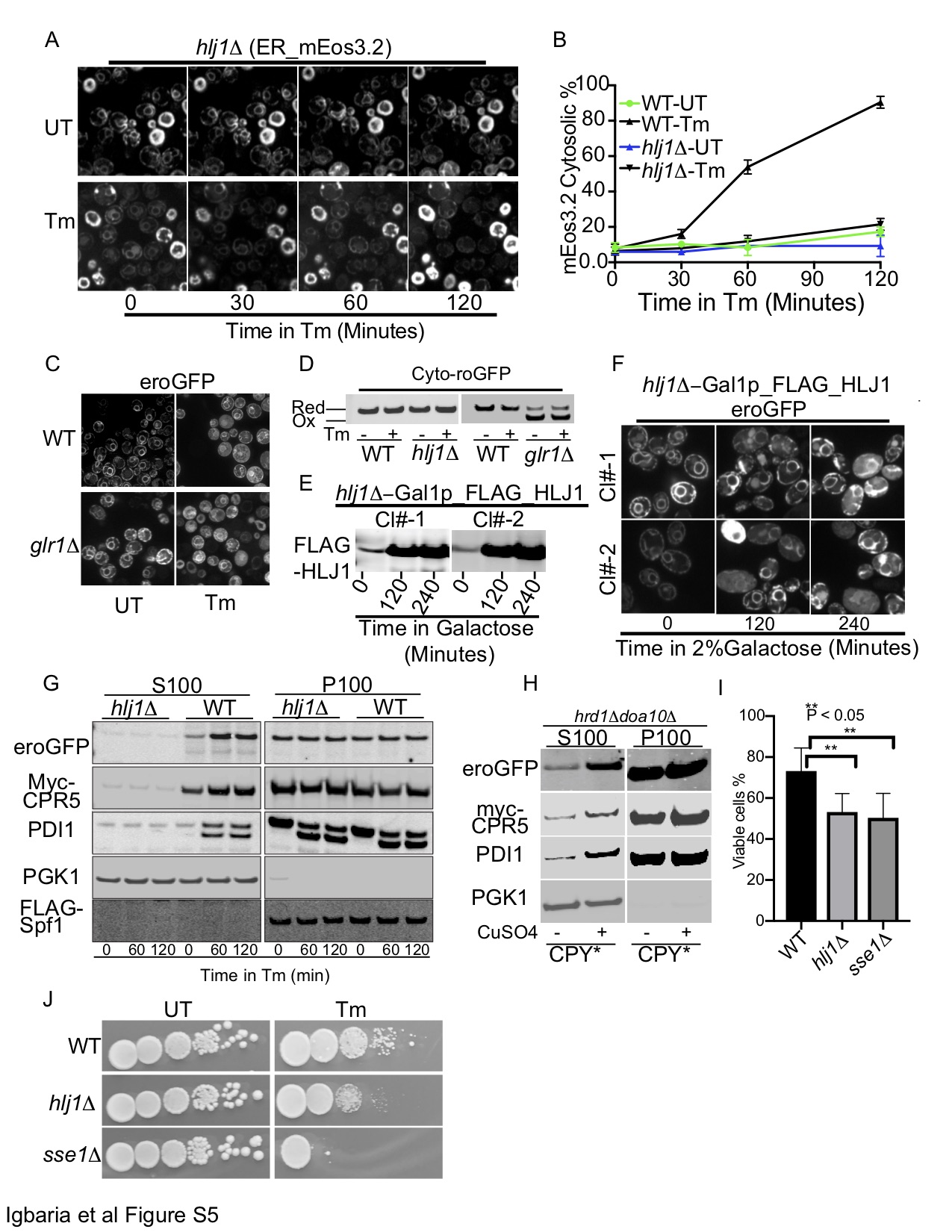
